# Cellular intelligence: dynamic specialization through non-equilibrium multi-scale compartmentalization

**DOI:** 10.1101/2021.06.25.449951

**Authors:** Rémy Tuyéras, Leandro Z. Agudelo, Soumya P. Ram, Anjanet Loon, Burak Kutlu, Kevin Grove, Manolis Kellis

**Affiliations:** Computer Science and Artificial Intelligence Laboratory, Massachusetts Institute of Technology, 02139, Cambridge MA; Picower Institute for Learning and Memory, Massachusetts Institute of Technology, 02139, Cambridge MA; Broad Institute of MIT and Harvard, 02139, Cambridge MA; Novo Nordisk Research Center Seattle. 98109, Washington

## Abstract

Intelligence is usually associated with the ability to perceive, retain and use information to adapt to changes in one’s environment. In this context, systems of living cells can be thought of as intelligent entities. Here, we show that the concepts of non-equilibrium tuning and compartmentalization are sufficient to model manifestations of cellular intelligence such as specialization, division, fusion and communication using the language of operads. We implement our framework as an unsupervised learning algorithm, IntCyt, which we show is able to memorize, organize and abstract reference machine-learning datasets through generative and self-supervised tasks. Overall, our learning framework captures emergent properties programmed in living systems, and provides a powerful new approach for data mining.

**Structured abstract:** *Background:* Although intelligence has been given many definitions, we can associate it with the ability to perceive, retain, and use information to adapt to changes in one’s environment. In this context, systems of living cells can be thought of as intelligent entities. While one can reasonably describe their adaptive abilities within the realm of homeostatic mechanisms, it is challenging to comprehend the principles governing their metabolic intelligence. In each organism, cells have indeed developed as many ways to adapt as there are cell types, and elucidating the impetus of their evolutionary behaviors could be the key to understanding life processes and likely diseases.

*Advances:* The goal of this article is to propose principles for understanding cellular intelligence. Specifically, we show that the concepts of non-equilibrium tuning and compartmentalization are enough to recover cellular adaptive behaviors such as specialization, division, fusion, and communication. Our model has the advantage to encompass all scales of life, from organelles to organisms through systems of organs and cell assemblies. We achieve this flexibility using the language of operads, which provides an elegant framework for reasoning about nested systems and, as an emergent behavior, non-equilibrium compartmentalization. To demonstrate the validity and the practical utility of our model, we implement it in the form of an unsupervised learning algorithm, IntCyt, and apply it to reference machine learning datasets through generative and self-supervised tasks. We find that IntCyt’s interpretability, plasticity and accuracy surpass that of a wide range of machine learning algorithms, thus providing a powerful approach for data mining.

*Outlook:* Our results indicate that the nested hierarchical language of operads captures the emergent properties of programmed cellular metabolism in the development of living systems, and provide a new biologically-inspired, yet practical and lightweight, computational paradigm for memorizing, organizing and abstracting datasets.

## Introduction

Biology, the study of living organisms, has mainly been built on experimental knowledge. While this knowledge has often been abstracted mathematically and computationally, the field of theoretical biology is far from being as unified as theoretical physics (*1*). Although sets of principles have been drawn for sub-fields of biology (*2*), there is no consensus as to how one should write, model and reason about biology within the same logical language (*3*). This discrepancy partly lies in the difficulty of modeling the multi-scale and diverse nature of living systems within a single framework (*4*) and partly in the complexity of understanding the chemical interactions defining life (*3, 5*). Despite the absence of a unified logical framework, current work seems to largely advocate the emergence of specialized functions in multi-scale living systems as the principle unifying ground for better understanding life (*4, 6*).

It goes without saying that the concept of specialization of function underlies many concepts of biology such as evolution, adaptation, and learning (*7*), in which case it is often associated with the concept of compartmentalization (*8*). From a cognitive perspective, compartmentalized specialization could be the key to designing more-brain-like artificial intelligence algorithms (*9, 10*). Indeed, although artificial neural networks (ANNs) were inspired by biological neuronal assemblies, they do not process information in the same way as brains do (*11*). Notably, ANNs are static architectures that lack many of the dynamic specialization abilities possessed by biological neural networks such as synaptic plasticity (*12*) and structural plasticity (*13*). Importantly, structural plasticity is believed to reinforce learning and memory (*14, 15*) through re-compartmentalization of dendritic spines (*16–18*). Such a compartmentalization process has been shown to be more pronounced in humans compared to rats (*19*), suggesting that this process could play an important role for human cognitive abilities. Interestingly, dendritic spines are themselves functionally diversified through their compartmentalized compositions (*20*), which makes dendritic compartmentalization only an intermediate level of cognitive organization and shows that multi-scale compartmentalization is fundamental to the organization of functional specialization in the brain.

While lack of functional specialization can compromise survival, extreme specialization can also be a dead end for living systems (*21*). Yet, living systems seem to naturally tend towards specialization (*22*), which lets us think of specialization as an equilibrium state towards which living systems are compelled to converge. On the other hand, an absence of specialization could be compared to a diffused mix of several specialization capabilities, all available within a same space of action. From an entropic point of view (*23*), this means that non-specialization can also be seen as an equilibrium state. These analogies with equilibrium states suggest that the survival of living systems lies in the ability to stay out of equilibrium (*24*). Interestingly, the oscillation resulting from this non-equilibrium tuning is reminiscent of a metabolic process working towards maintaining life. To better appreciate this analogy from an entropic point of view, we can compare living systems as compartments containing specific chemical attributes, in which case the game of interactions happening in a system of living systems (*25*) can be likened to Maxwell’s demon thought experiment (*26*). In this theoretical experiment, the compartmentalized system is able^1^ to increase the entropy – essentially the specialization – of one of its compartments by letting specific types of chemical attributes migrate to other compartments. On the other hand, the migration of these attributes may disturb the specialization processes of the surrounding compartments. Overall, if all the compartments use Maxwell’s demon principle to optimize their specialization, then the system should reach a stable oscillatory configuration and be endowed with the homeostatic needs characteristic of a living system.

Importantly, such self-organized dynamics do not necessarily take place within obvious compartments and can occur within ambiguous and flexible boundaries. For example, we showed in parallel work that liquid-phase-separated organelles formed in nuclei of cells can cluster and regulate the transcription of trait-specific genes as a response to starvation and cold conditions (*27*). Furthermore, these liquid-phase-separated optimizations are recorded, through evolution, into region-specific compartments in the chromatin architecture, facilitating cell-specific and context-dependent responses to metabolic pressures. This suggests that multiscale compartmentalization should not be thought of as a physical delimitation, but more as a context-dependent hierarchical organization of dynamic specialization events.

Here, we show, both mathematically and computationally, that we can recover the concept of dynamic specialization from the concepts of multi-scale compartmentalization and nonequilibrium tuning in a way that pieces together the principles described above. While our mathematical model establishes a link between two so-far-unrelated domains of science, namely operad theory (*28,29*) and systems biology (*4*), our computational model shows how multi-scale compartmentalization engenders learning and how non-equilibrium tuning makes this learning flexible.

## Results

### Modeling multi-scale living systems as nested structures

We model multi-scale organizations of living systems as nested arrangements of 2-level structures. Specifically, these 2-level structures constitute the building blocks of the system at a given scale (**Fig. 1A**). Abstractly, we visualize each of these building blocks as a circular environment encompassing a collection of smaller circular environments meant to receive further nesting. Because the structure of these building blocks is reminiscent of that of a cell, for which the outer environment is the cytosol and the inner environments are the organelles, we call these 2-level structures “cells” – or sometimes “abstract cells”, to distinguish them from living cells.

**Fig. 1.**
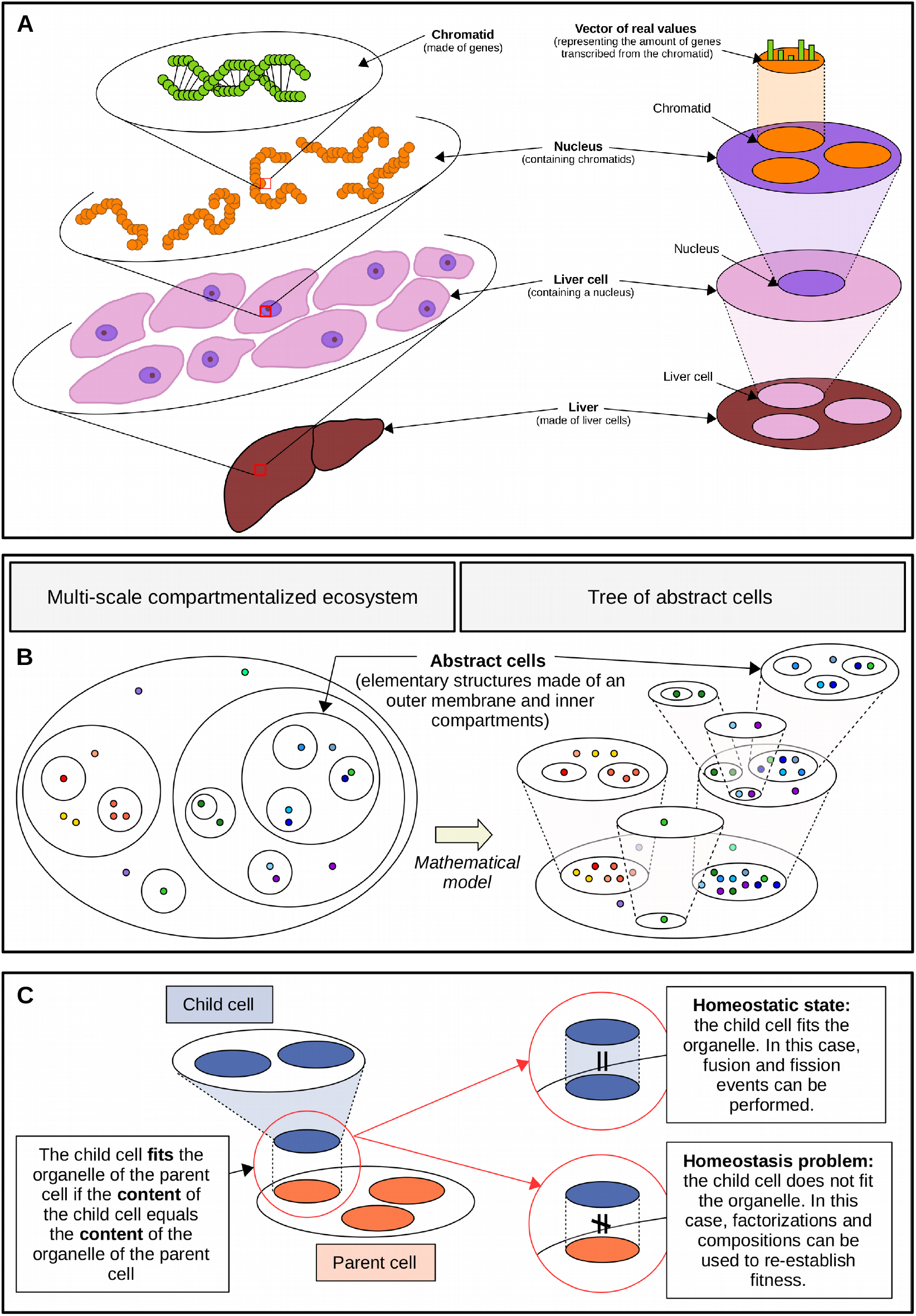
Modeling the inter- and intra-scale connections defining a multi-scale ecosystem. (**A**) Schematic of a multi-scale organization of a liver. On the left, successive scale enlargements zoom in the nested composition of the liver on three levels: a close-up on cells within the liver, a close-up on chromatids within a nucleus and a close-up on a piece of DNA within a chromatid. As illustrated on the right, these successive close-ups can be represented as a nesting of cell-like structures. These cell-like structures are made of a milieu and a collection of compartments meant to receive further nesting. The milieu and each compartment is equipped with a vector of real values representing its functional specialization within the system. A possible interpretation of the content of a chromatid is the amounts of proteins transcribed from the genes contained in the chromatid. (**B**) Schematic showing how we model a multiscale compartmentalized ecosystem as a tree of (abstract) cells. We can imagine visualizing the multi-scale compartmentalized ecosystem by looking at the tree of cells from above, as suggested by the descending transparent cones. In the tree shown in the picture, each cell compartment describes the content of its hovering child cell, without the compartmentalization structure: the tree has reached homeostasis (**C**) Illustrates the concept of fitness: a cell is said to fit an organelle of another cell if the content of the first cell is equal to the vector encoding the organelle of the second cell. If all cells of a tree reach fitness, then we are allowed to use operations such as fussion and fission within the tree. When a cell does not fit its parent organelle, then we can always change the value of the cytosolic contents of the two cells in order to establish fitness.

Within such a formalism, a multi-scale compartmentalized ecosystem can be visualized as a tree whose root, junctions and leaves are each associated with an abstract cell (**Fig. 1B**). To model the functional specializations of the ecosystem, we equip the outer environment and the inner compartments of every cell with vectors of real numbers of a fixed dimension (see the definition below). Each component of a vector represents the degree of specialization of the corresponding environment towards a specific function. This means that a vector represents the global specialization of its associated environment (**Fig. 1A**). Formally, for a given non-negative integer *N*, we define a *cell c* of *dimension N* as a quadruple (*n*, res(*c*), cyt(*c*), org(*c*)) where:

1. *n* is a non-negative integer, called the *number of organelles*;
2. res(*c*) is a non-negative real number, called the *residual*;
3. cyt(*c*) is a *N* -dimensional vector of real values, called the *cytosolic content*; and
4. org(*c*) is a *n*-tuple (org(*c*)_1_, org(*c*)_2_, …, org(*c*)_*n*_) of *N* -dimensional vectors of positive real values whose *i*-th component org(*c*)_*i*_ is called the *i-th organelles*.

The previous definition allows us to model multi-scale compartmentalized ecosystems that can functionally specialize in *N* different ways as trees of abstract cells of dimension *N*. More specifically, this means that we consider trees whose roots, junctions and leaves are each labeled by an abstract cell of dimension *N* such that every cell associated with a junction possesses a number of organelles equal to the number of children possessed by the junction (**Fig. 1B**). By convention, we link the *i*-th organelle of the cell associated with a junction to the cell associated with *i*-th child of the junction. Visually, each branch of the tree represents a close-up of the organelle with which it is associated, as illustrated in **Fig. 1B**.

To globally assess the amount of specialization within a cell, we define the *content K*(*c*) of a cell *c* as the *N* -dimensional vector resulting from the sum of the cytosolic content cyt(*c*) and all the organelle vectors org(*c*)_1_, org(*c*)_2_, …, org(*c*)_*n*_. Intuitively, a content vector describes the “visible inside” of the cell by overlooking its compartmentalized structure as well as its residual, which we regard more as an energetic resource (see Supplementary text). We say that a cell *d fits* the *i*-th organelle of another cell *c* if the content *K*(*d*) is equal to the *i*-th organelle vector org(*c*)_*i*_. This concept of fitness allows us to specify whether the cell *d* is a perfect close-up description of the inside of the *i*-th organelle of the cell *c*, as illustrated by the small colored particles shown in **Fig. 1B**.

Finally, a tree of cells can be in two states: it is in either a homeostatic state or a non-homeostatic state. We say that a tree is in a *homeostatic state* when, for each cell *c* that is not a leaf of the tree, meaning that *c* possesses a collection of child cells *d*_1_, …, *d*_*n*_ in the tree, every organelle vector of the form org(*c*)_*i*_ is equal to the content *K*(*d*_*i*_) of the *i*-th child cells *d*_*i*_. On the other hand, the tree is said to be in a *non-homeostatic state* if it is not in a homeostatic state (**Fig. 1C**).

In what follows, we use fitness to establish “contracts” between child cells and their corresponding parent cells. This allows us to define operations that preserve consistency within trees of cells.

### Modeling dynamic behaviors in terms of operadic operations

While trees of cells model the architectural aspect of multi-scale ecosystems, they cannot model the dynamic interactive behaviors that one often associates with living systems, namely communication, collaboration, exchanges and reproduction. To model such behaviors, we need to borrow concepts belonging to operad theory. Indeed, operad theory also focuses on tree-like structures, but along with composition operations, from which we can derive a whole dictionary of dynamic operations. In our case, we introduce composition operations on abstract cells, which we subsequently extend to trees of abstract cells. Intuitively, composing two cells together means inserting one cell into the organelle of the other cell and removing the “membrane” of the inner cell so that the organelles and cytosolic content of both cells mix together (**Fig. 2A**). More formally, this means that the *composition* of a cell *d* with a cell *c*, at the *i*-th organelle of *c*, is the cell *c* ○_*i*_ *d* whose:

**Fig. 2.**
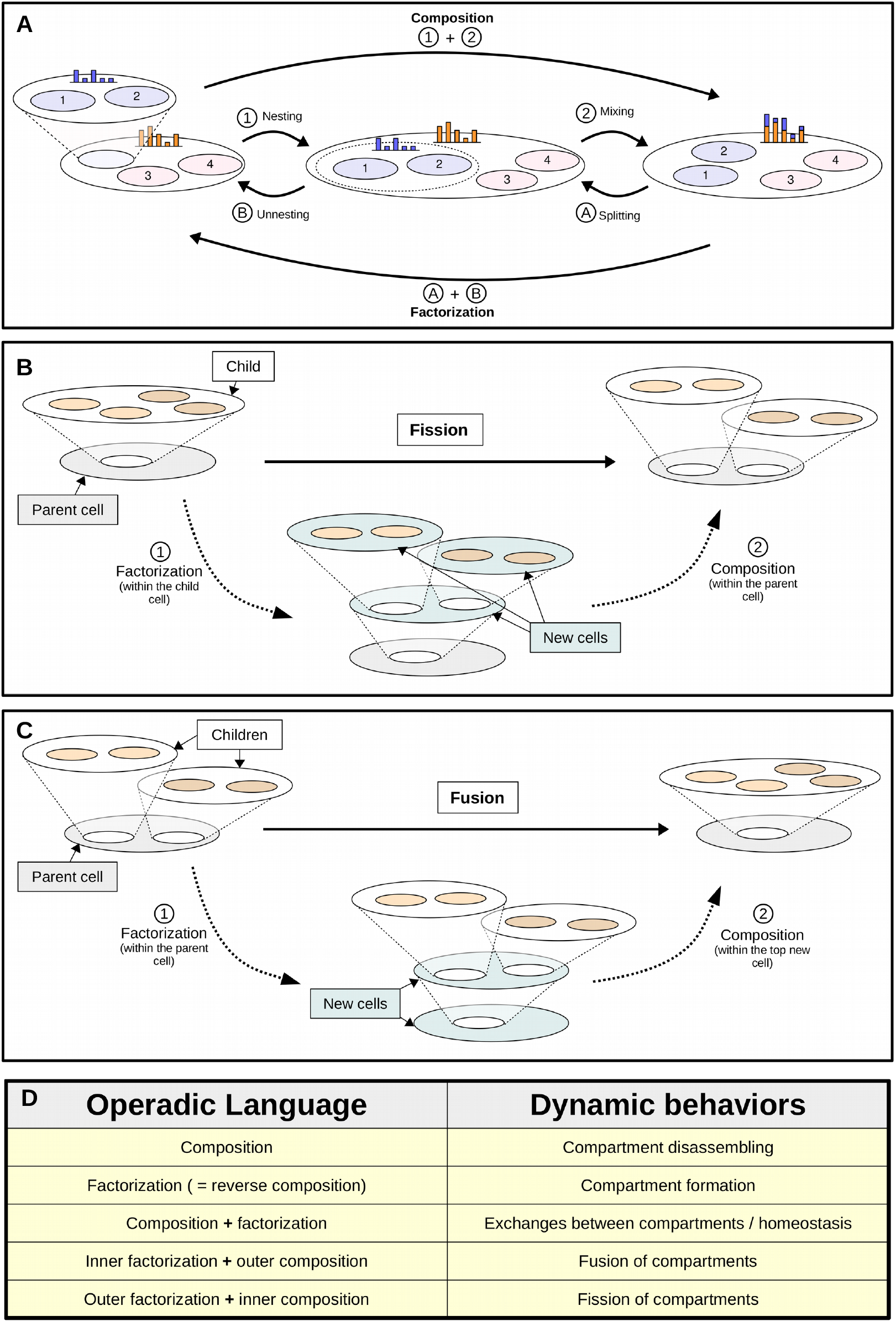
Modeling fundamental dynamic behaviors of compartmentalized ecosystems in terms of compositions. (**A**) Schematic of a composition of two cells, from left to right, and a factorization of a cell, from right to left. As can be visualized, composing two cells can be seen as the process of, first, nesting one cell (the inner cell) within an organelle of another cell (the outer cell) and, second, removing the membrane separating the inner cell from the outer cell so that the organelles and the cytosolic content of the inner cell can be mixed with the organelles and the cytosolic content of the outer cell. The residuals are summed along the way to give the residual of the resulting composition. The reverse process represents a factorization. More specifically, factorizing a cell can be seen as the process of, first, isolating a set of organelles, a part of the cytosolic content and a part of the residual within a compartment and, second, unnesting this compartment to create a new cell, as shown on the left. (**B**) Schematic showing how cell fission can be modeled from a factorization followed by a composition. Specifically, the panel shows how we can divide the hovering white cell of the top left complex into a pair of cells. First, we factorize the hovering child cell into a 2-level tree and we compose the middle cell of the resulting 3-level tree within the parent cell of the whole complex. (**C**) Schematic showing how cell fusion can be modeled from a factorization followed by a composition. Specifically, the panel shows how we can merge the two hovering white cells of the top left complex into a unique cell. First, we factorize the parent cell into a 2-level tree made of two superposed cells and we compose the two hovering white cells within the middle cell of the resulting 3-level tree. (**D**) Table translating the operadic terminology into a compartment ontology.

1. number of organelles is equal to the sum *n* + *m*;
2. collection of organelles org(*c* ○_*i*_ *d*) consists of all the organelles of *c*, except for the *i*- th organelle org(*c*)_*i*_, which is replaced with the collection of organelles of *d*, as shown below, where *n* and *m* denote the number of organelles of *c* and *d*, respectively:

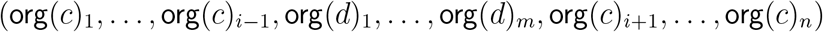
3. cytosolic content cyt(*c* ○_*i*_ *d*) is the sum of the vector cyt(*d*) with the vector cyt(*c*);
4. residual res(*c* ○_*i*_ *d*) is the sum of the quantity res(*d*) with the quantity res(*c*).

From the notion of composition, we derive the notion of factorization, which describes the reverse process of a composition (**Fig. 2A**). More specifically, the factorization of a cell *c* is a pair of cells (*c*′, *d*) as well as an index *i* for which the equation *c* = *c*′ ○_*i*_ *d* holds. Intuitively, a factorization can be thought of as the formation of a compartment within a cell *c*, which, once created, becomes the *i*-th organelle of a cell *c*′. The cell *d* can be seen as a close-up description of the inside of this compartment (**Fig. 2A**).

While it makes sense to use a composition at a specific organelle, it also makes sense to consider simultaneously compositions of cells within an ambient cell. Specifically, for a given cell *c* with *n* organelles and a given *n*-tuple of cells *d* = (*d*_1_, …, *d*_*n*_), we define the simultaneous composition of *c* with the collection of cells *d*_1_, …, *d*_*n*_ by the following equation:

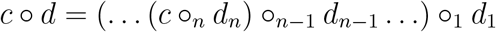

We show that the order in which we compose the cells *d*_*i*_ does not affect our results – therefore our composition operation is *operadic* (Supplementary text).

Interestingly, we can model exchanges between cells through loops of successive compositions and factorizations (**Fig. 2A** – loop of arrows). Specifically, if a cell *d* represents a close-up description of the *i*-th organelle of a cell *c*, then we can create an exchange between the cytosolic contents of *c* and *d* by first composing *d* at the *i*-th organelle of *c* and subsequently factorizing the resulting composite *c* ○_*i*_ *d* as a pair (*c*′, *d*′) such that *d*′ possesses the same residual and organelles as *d* and such that the relation *c* ○_*i*_ *d* = *c*′ ○_*i*_ *d*′ holds. The process of passing from the pair (*c, d*) to the pair (*c*′, *d*′) simulates an exchange between the cytosolic content of the cells *c* and *d*. Notably, such exchanges can be used to re-establish fitness between two cells. In Supplementary text, we draw a conceptual parallel between these types of tuning exchanges and the concept of homeostasis.

Another use of factorizations and compositions is the modeling of fission and fusion of cells in trees containing at least two levels of cells (a parent cell and a collection of child cells). Specifically, we can divide a child cell *d*, say associated with the *i*-th organelle of a parent *c*, by first factorizing the cell *d* as two child cells *d*_1_, *d*_2_ and a parent cell *c*′ and then composing the new parent cell *c*′ at the *i*-th organelle of the parent cell *c* (**Fig. 2B**). Conversely, we can merge two child cells *d*_1_, *d*_2_, respectively associated with the *i*_1_-th and *i*_2_-th organelles of a parent *c*, by first factorizing the parent *c* as a child cell *d* and a parent cell *c′* such that *d* specifically contains the two organelles of *c* associated with *d*_1_ and *d*_2_ and then composing the child cells *d*_1_ and *d*_2_ with the parent cell *c*, at the corresponding organelles (**Fig. 2C**).

Overall, the language of compositions provides a convenient and simplified language to recover dynamic behaviors that are usually associated with living cells and, more generally, living systems (**Fig. 2D**).

### Modeling interactions with an environment

Although compositions and factorizations allow the simulation of exchanges between abstract cells, these interactions are only internal to the system and do not reflect the interactions that the system may have with its surrounding environment. To simulate interactions with the environment, we make an analogy between the concept of external resource and the concept of information to be fed to the cell. As will be shown, an information intake results in an output that possesses the same structure as that of the input. This allows us to extend the feeding process to trees of abstract cells through forward-propagation.

Specifically, we model the environment of an abstract cell of dimension *N*, equipped with *n* organelles, as a collection of *n* × *N* -matrices. Each matrix represents a state of the environment offering potential resources to the cell. Conventionally, we regard these *n* × *N* -matrices as *n*-dimensional vectors *a* = (*a*_1_, *a*_2_, …, *a*_*n*_) whose *i*-th component *a*_*i*_ is a *N* -dimensional vector (*a*_*i*,1_, *a*_*i*,2_, …, *a*_*i,N*_). Each quantity *a*_*i,u*_ represents an amount of resource meant to be processed through the *u*-th function of the *i*-th organelle of the cell *c*.

As illustrated in **Fig. 3A**, we define the intake of a *n* × *N* -matrix *a* by the cell *c* in terms of a series of multiplications and additions between the coefficient of *a* and the parameters of *c*. Specifically, for each function *u* ∈ {1, …, *N*} and each organelle index *i* ∈ {1, …, *n*}, we define the action of the *i*-th organelle of *c* on the input *a*, relative to the *u*-th function, as the multiplication of the quantity *a*_*i,u*_ with the ratio org(*c*)_*i,u*_*/*𝒮*K*(*c*), where 𝒮*K*(*c*) denotes the sum of the components of *K*(*c*), namely *K*_1_(*c*) + *K*_2_(*c*) + … + *K*_*N*_ (*c*). This ratio quantifies the amount of specialization of the *i*-th organelle of *c* at the *u*-th function. We then define the action of the cell *c* on the input *a*, relative to the *u*-th function, by the sum of the actions done by each organelle, namely the following sum:

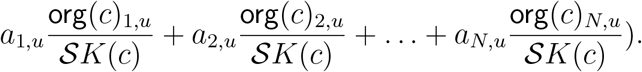

**Fig. 3.**
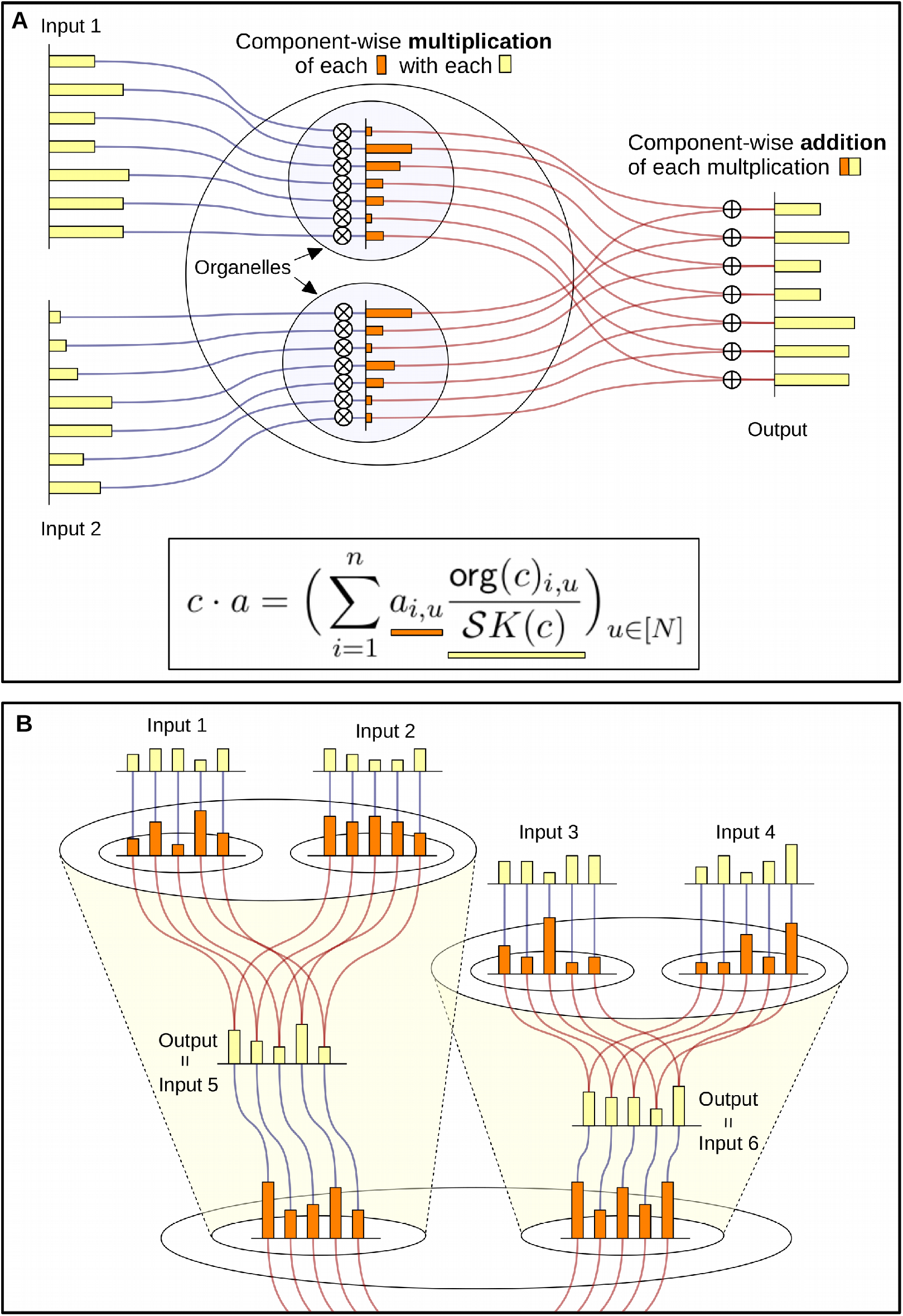
Modeling external exchanges through the underlying circuit structure of trees of cells. **(A)** Schematic showing how an abstract cell of dimension 7, equipped with 2 organelles, processes a 2 × 7-matrix of real numbers. The output, which is a vector of dimension 7, is the result of a component-wise multiplication of the columns of the matrix (*i*.*e*. called “input 1” and “input 2”) with the organelles of corresponding index, followed by a component-wise sum of these multiplication. (**B**) Schematic illustrating how a tree of cells of dimension 7, equipped with 4 leaf cells, processes a 4 × 7-matrix by propagating the actions of its cells from the leaves to the root. As can be seen, outputs returned by child cells are turned into inputs for the parent cells.

Finally, we define the *action* of the cell *c* on the input *a* as the vector of all actions, as shown below.

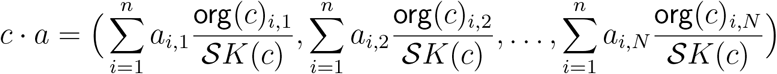

Extending cell actions to trees of cells essentially consists in transferring the actions of the child cells to their corresponding parent cells (**Fig. 3B**). More specifically, the organelles of the parent cell act on the outputs returned by the child cells to generate the parent cell’s action, which may then be used as a new input by one of the organelles of another parent cell, and so on. Note that the tree structure makes sure that the number of outputs created by a group of sibling cells matches the number of organelles of the corresponding parent cell.

### Modeling non-equilibrium tuning through the concept of operadic algebras

In the same way as a living system influences its internal exchanges through its interactions with the environment, the actions associated with a tree of cells can be used as a means to influence exchanges between its internal cells. To do so, we take inspiration from Maxwell’s demon thought experiment, which provides a non-equilibrium optimization principle. As explained below, we formalize this principle mathematically in terms of the canonical equation for operadic algebras.

According to Maxwell’s demon thought experiment, low-entropy configurations are associated with highly compartmentalized organizations, while high-entropy configurations are associated with moderately compartmentalized organizations. Using this analogy, in **Fig. 4A** we illustrate how living systems are able to both share and isolate their chemical contents by taking advantage of these low- and high-entropy configurations (allowing both communication and isolation). Building on this illustration, we want to think of living systems combining both low- and high-entropy configurations as minimizing the difference between the effects of their internal chemistry, when highly compartmentalized and when deprived of such a compartmentalization. Based on this principle, we evaluate the difference between low- and high-entropy configurations by comparing the actions of trees of cells with different compartmentalized structures.

**Fig. 4.**
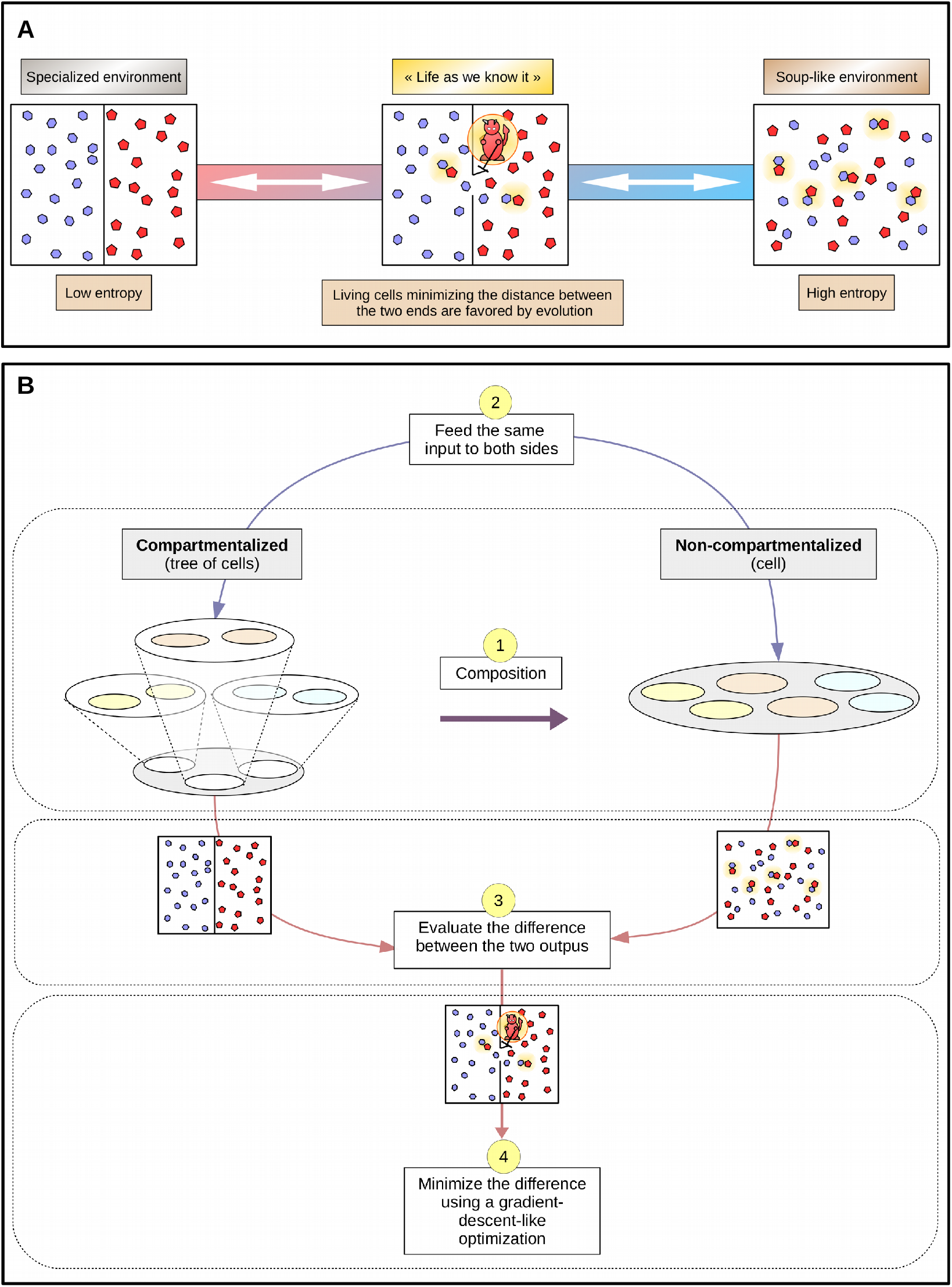
Tuning a tree of cells through a non-equilibrium-inspired loss function. **(A)** Schematic showing a refinement of Maxwell’s demon thought experiment. The panel shows three entropic state of a system. On the left, the system has a low-entropic configuration and is separated by a wall in two closed compartments containing dissociated types of chemical substances. On the right, the system has a high-entropic configuration and is mixture of two chemical substances spread throughout the system. In the middle, the system has an intermediate-entropic configuration and refers to the type of interaction promoted at proximity of the membrane of living cells, in which the boundary between low and high entropy is flexible. As illustrated, such flexibility allows both local chemical interactions and isolation of the chemical substances. (**B**) Schematic showing how each junction of a tree of cells is optimized during a 4 step procedure, which we design based on compartmentalization logics and non-equilibirium tuning. First, each junction of the tree is turned into a cell by composing the child cells with the parent cell, as shown in step 1. One of the main differences between the junction and its composition is their respective levels of compartmentalization (the junction having one more level). This difference allows us to compare the effect of the removed compartments in the same spirit as in Maxwells demon thought experiment. Specifically, we compare the two systems by giving them the same set of inputs (step 2) and we evaluate the difference of the returned outputs (step 3). We then use a gradient-descent-like optimization to deduce the changes to be made in the system in order to minimize this difference. This minimization is achieved along with various factorizations and compositions simulating formation and removal of compartments.

More specifically, for each junction of a tree of cells made of a parent cell *c* and a collection of child cells *d* = (*d*_1_, …, *d*_*n*_), we compare the action of the junction on a given input matrix *a* = (*a*_1_, …, *a*_*n*_) (**Fig. 4B** – left tree of step 1) with the action of the composition *c* ○ *d* on the same matrix *a* (**Fig. 4B** – right cell of step 1) via the following operation:

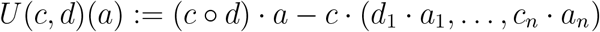

Note that the operation *U* is very well-known in operad theory and is used to define “algebras of operads” in terms of the equation *U* (*c, d*)(*a*) = 0, which is required to hold for every input *a*. In our context, solutions to such an equation would provide cells *c* and *d*_1_, …, *d*_*n*_ whose associated vectors contain zeros in all but one dimension. Because this suggests a type of extreme specialization, we take a more cautious approach by regarding the equation *U* (*c, d*)(*a*) = 0 more as an ‘ideal situation’ towards which the cells *c* and *d*_1_, …, *d*_*n*_ should be pressured to converge. In this respect, we are more interested in solutions (*c, d*) for which the scalar product *U* (*c, d*)(*a*)·*U* (*c, d*)(*a*) = *U* (*c, d*)(*a*)^2^ is as close as possible to 0. This idea of converging toward a ‘minimizing situation for a given environment’ matches Maxwell’s demon picture, in which finding the parameters that minimize the multivariate function (*c, d*) ⟼ *U*(*c, d*)(*a*)^2^ is to give us the metabolic states at which living systems tend to thrive (**Fig. 4B**).

In Supplementary text, we explain how we can minimize the multivariate function (*c, d*) ⟼ *U* (*c, d*)(*a*)^2^ through two types of optimizations: a gradient-descent-like optimization and a structural optimization using fission and fusion events on the cells *d*_1_, *d*_2_, …, *d*_*n*_. Our gradientdescent-like optimization cannot be seen as a conventional gradient descent optimization because we compute the differential of (*c, d*) ⟼ *U* (*c, d*)(*a*)^2^ on a manifold that is not the Euclidean plane. More specifically, our differential is computed as an infinitesimal variation of the function (*c, d*) ⟼ *U* (*c, d*)(*a*)^2^ relative to an infinitesimal cytosolic exchange between the cell variables *c* and *d*. We then use the resulting differential to change the parameters of the cells (*c, d*) to obtain a solution (*c*′, *d*′) with a smaller value *U* (*c*′, *d*′)(*a*)^2^. After a complete re-parametrization of the cells, the tree may not be in a homeostatic state and any fitness disagreement is corrected through cytosolic exchanges between parent cells and child cells. In addition to our gradient-descent-like optimization, we also show how the value of the function (*c, d*) ⟼ *U* (*c, d*)(*a*)^2^ can be reduced by merging or dividing the child cells *d*_1_, *d*_2_, …, *d*_*n*_. In Supplementary text, we give strategies to determine the set of cells to be merged or divided as well as methods to compute the division of a cell *d*_*i*_.

Overall, we succeeded in making a tree of cells learn, organize itself and specialize with respect to its environment by alternating between numerical and structural optimizations. The result of such an optimization process is shown in the following paragraph.

### Simulating dynamic specialization

In this paper, we propose IntCyt, a software that implements the mechanics discussed in the previous sections as an unsupervised machine learning algorithm.

The memory architecture on which our software IntCyt relies is a tree of abstract cells whose constituting organelle vectors learn abstractions of concepts given to the tree in the form of input vectors. All our inputs come from reference machine learning datasets such as `MNIST`, `fashion-MNIST` and `DREAM3` (see the documentation of IntCyt). For every input *a* and every junction (*c*, (*d*_1_, …, *d*_*n*_)) in the tree, we use a gradient-descent-like algorithm (together with factorization and composition operations) to update the memory stored in the organelles of the cell *c*. As illustrated in **Fig. 5A**, the information learned by the tree is contained in the organelles of each cell. Note that, when the tree is in a homeostatic state, the information contained in the organelle of each parent cell is the sum of the information contained in their corresponding child cells (**Fig. 5A**).

**Fig. 5.**
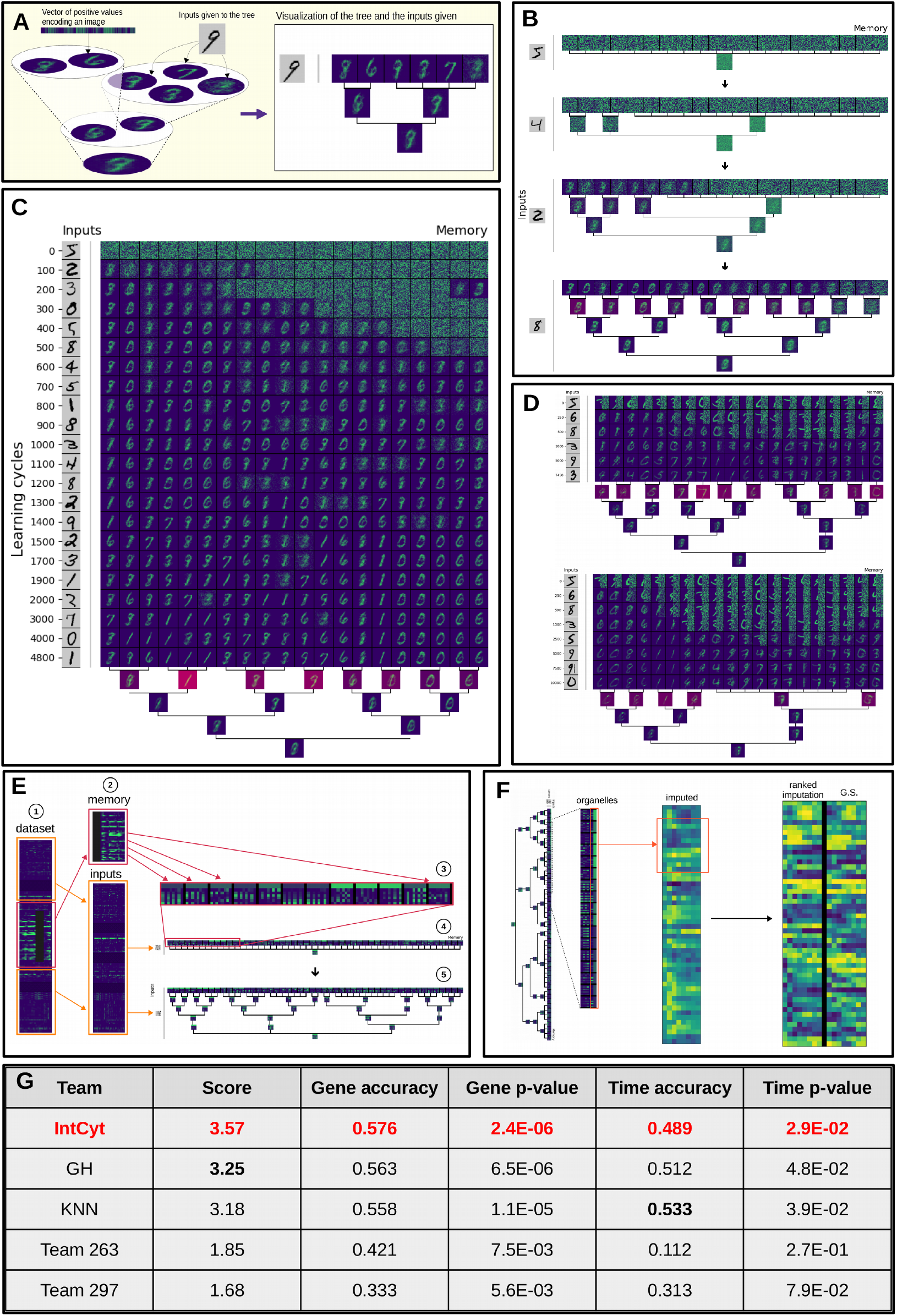
Learning reference machine learning datasets with IntCyt. (**A**) Schematic showing a translation of our usual tree representation into a computerized visualization of the memory of IntCyt. As suggested by the right-hand side tree, the computerized visualization shows the vector associated with the organelles of each cell. When the tree is in a homeostatic state, the organelles of the parent cells correspond to the contents of each child cell. The input given to IntCyt at a specific learning cycle is displayed on the left, in black and white. (**B**) Visualization showing the development of the architecture of IntCyt on the `MNIST` dataset from a single cell to a hierarchical network of cells. The figure shows the state of the tree for the cycles 0, 10, 100 and 500. During the development of the tree, IntCyt learns, abstracts and memorizes concepts from the inputs. The redder cells displayed in the tree, shown at the bottom, are associated with relatively high residuals. The amount of red displayed in each cell *c* is proportional to the quantity 1 +res(*c*)*/* 𝒮 *K*(*c*). (**C**) Visualization showing the evolution of the organelles of the leaf cells as well as the state of the tree for the last cycle. The inputs given to the tree are taken from the `MNIST` dataset. As can be seen, IntCyt first separates noise from information, and eventually clusters the different memorized pieces of information into hierarchical compartments. (**D**) Visualization showing the outputs of IntCyt on a set of `MNIST` images whose right sides are hidden with white noise. The panel shows two reconstructions of the set of images by IntCyt for different compartmentalization parameters (we use *ω* = 0 and *ω* = 30, respectively – see Supplementary text). (**E**) Schematic showing the use of IntCyt on the `DREAM3` dataset in order to impute partially-hidden gene expressions using the same type of technique as that used in panel **D**. (**F**) Schematic showing the extraction of learned information from the self-supervision of IntCyt on the `DREAM3` dataset. The extracted information gives imputed gene expression, which we rank and compare to the (ranked) golden standard provided with the `DREAM3` dataset. (**F**) Table showing the results of IntCyt on the `DREAM3` challenge and comparing them to the best results obtained during the challenge. As can be seen, IntCyt outperforms all the participants.

In the same fashion as differentiated cell tissues can be produced from a single cell through division and cellular communication, our implementation IntCyt starts its simulations with a single abstract cell, which it eventually develops into a tree of cells through fission and fusion operations (**Fig. 5B**). Importantly, we decompose these operations as successions of factorization and composition operations (Supplementary text). These two-step decompositions are particularly useful to avoid developmental incoherences. For instance, the root cell of a tree has no parent in which it can divide. As a result, a fission operation occurring at a root cell can only consist of a factorization step since the complementary composition step would require a parent cell in order to be completed. This asymmetric use of compositions and factorizations at the root cell forces the creation of new compartments, which eventually lead to the emergence of a hierarchical organization of the organelles of the original cell in the form of a tree of cells. In other parts of the tree, cells merge and divide properly, creating an exchange of compartments (*i*.*e*. of organelles) that contributes to the localization of the different pieces of information contained in the tree. More specifically, every fusion and fission of cells occurs as a way to minimize the algebra operator *U*. As explained in Supplementary text, this optimization process consists in merging cells that contain similar information together, and dividing cells with relatively distinct information into different compartments. Overall, these mechanisms lead to a local-to-global reinforcement of the different pieces of information contained in the tree of cells.

In broad terms, IntCyt is performing a dynamical integration of the inputs within a flexible hierarchical database. The ability of IntCyt to reshape its hierarchical architecture mainly lies in the use of composition operations, which aim to cancel out earlier factorizations in order to reorganize the organelles of the cells. As shown in **Fig. 5B** and **Fig. 5C**, IntCyt first starts to separate information from noise by using the information contained in the inputs. This separation tends to gather information, triggering what we could see as recruitment mechanisms, in which organelles containing information would couple with distant organelles that start to display non-random information of a similar type (**Fig. S1A**). These recruitment processes promote the reinforcement of the learned information across higher levels in the tree. This specialization process then influences the overall learning activity of the cell itself. For instance, certain organelles may be more representative of the content of the cell due to their higher values. As a result, the cell containing these organelles develops a preference towards these high functional specializations, which eventually influence all the organelles of the cell to specialize towards these functions (**Fig. S1B**).

Importantly, IntCyt is able to make abstractions of concepts learned at lower levels (*e*.*g*. the leaf cells) visible to higher levels (*e*.*g*. the parent cells) by ensuring that the tree is always in a homeostatic state after every update of the gradient descent. Importantly, this visibility is made possible through the removal of the cytosolic contents at every learning cycle. However, to avoid making the overall memory of IntCyt converge toward an empty memory through these removals, we make sure to eliminate the cytosolic contents in a way that preserves the overall numerical content of the system. To give more detail about this important procedure, take into account that, for every cell *c*, the vector cyt(*c*) is made of either negative or non-negative values, which account for past exchanges between organelles. If the sum of the components of cyt(*c*) is equal to zero, then turning the vector cyt(*c*) into a zero vector does not create any numerical loss in the system. On the other hand, if the sum 𝒮cyt(*c*) is not zero, then we prevent numerical loss by adding the amount 𝒮cyt(*c*) to the residual res(*c*).

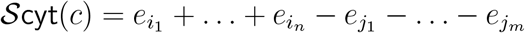

As a consequence, residuals can be used to measure the amount of learning stress under which the cell is, giving an intuitive way to perceive the progress of the system (amounts of residuals are indicated by the redder cells in **Fig. 5B, 5C** and **Fig. S1C**).

Interestingly, we can interpret the conversion of the cytosolic content cyt(*c*) into a zero vector as a chemical reaction of the form 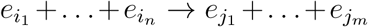, which, through the lenses of metabolism, can be viewed as a spontaneous chemical reaction releasing an amount of energy equal to res(*c*). These transformations work together with our gradient descent algorithm: while the minimization of the values of the algebra operator *U* increases specialization in the dimensions of the inputs that are the most prominent and decreases specialization in the dimensions with lesser presence, the chemical reactions ensure that loss or gain in certain functional specializations has a counter-effect on other functional specializations, creating a competition between every specialization of the system. This competition phenomenon eventually leads to only retaining the most prominent specializations for both the inputs and the memory of the system. Gradually, these selection mechanisms leads to evolutionary-like memorizations and abstractions of the concepts represented in the inputs. Along with this evolutionary-like learning, the system is also able to use the communication mechanisms occurring between the organelles (**Fig. S1B**) to smoothen the representation memorized across the whole tree and prevents IntCyt to overfit the input data (**Fig. S1C**).

As is customary in machine learning, we couple gradient descent increments with various update policies in order to stabilize and preserve learned information throughout the learning phase (Supplementary text). These policies are possible due to the interpretability of the information memorized by the organelles. More specifically, every organelle vector faithfully represents the images displayed in the visualizations of **Fig. 5** and **Fig. S1**. As a result, the organelles do not need to be preprocessed through heuristics in order to query their accumulated knowledge. This means that determining the type of information learned by IntCyt and the rate at which this information is learned is relatively straightforward.

Finally, we took advantage of the interpretability of IntCyt to resolve self-supervised learning challenges, namely challenges in which we desire to reconstruct data containing missing information. In **Fig. 5D**, we show two such reconstructions for images originating from the `MNIST` dataset. As shown in the panel, the output of IntCyt consists of a hierarchical clustering of the data as well as its full reconstruction. In **Fig. 5E, 5F, 5G**, we use this self-supervised learning approach to resolve the `DREAM3` challenge. This challenge relies on a dataset containing 9335 gene expressions and consists in reconstructing partially-hidden gene expressions for 50 genes of the dataset. To do so, we use the partially-hidden gene expressions as self-supervised data and we use the rest of the data as input (or “training”) data (**Fig. 5E**). After 3 runs of IntCyt on the training dataset, the missing data is imputed into hypothetical differential gene expressions, which we compare to the golden standard of the challenge after a ranking transformation (**Fig. 5F**) – see the documentation of IntCyt for more detail. As shown in **Fig. 5G**, we found that IntCyt was able to outperform the best algorithms of the challenge (*30*). Overall, our results show that IntCyt combines hierarchical clustering, concept abstractions and data reconstruction within a single paradigm, namely non-equilibrium compartmentalization, and is able to outperform classical unsupervised machine learning algorithms on reference machine learning challenges.

## Discussion

To our knowledge, this is the first work that has proposed a general mathematical framework for the field of multi-scale systems biology (*31*). More specifically, we showed that we can use this framework to reason across multiple scales through a set of fundamental operations. Furthermore, we also showed that these operations can be used to both model fundamental biophysical principles pertaining to the emergence of life and simulate intelligence-like behaviors present across all living beings. While the quantitative nature of our formalism gives us the freedom to integrate specific numerical models of systems biology within our framework, its hierarchical nature provides a multi-scale modeling logic giving the possibility of integrating such methods consistently across all scales of biology. From the point of view of the literature, our work tackles the longstanding need of ‘a generic theory or calculus of multi-scale modeling’ for systems biology, as explained in (*32*). Based on Castiglione *et al*. (*33*), our framework addresses the ‘ultimate goal of multi-scale modeling’ by proposing a mathematical language in which it is possible to ‘link [every scale] in a consistent manner so that the information from a lower scale can be carried into the simplified model of a higher scale’. Indeed, while frameworks for multi-scale systems biology have been put forth (*34*), these do not propose a fundamental language in which models can be designed within a single logic, but rather aim to integrate a network of already existing single-scale logics through ‘scale bridging techniques’ (*34*). The disadvantage of such approaches is the requirement to solve ‘bridging problems’, which may be far from obvious and quite challenging. By contrast, our framework can overcome such difficulties through factorization mechanisms allowing the refinement of a scale into a decomposition of intermediate scales, methodically inviting the user to add further information into the model. We would like to point out that the most valuable assets of our proposed framework all originate from operad theory. Interestingly, this field of science emerged from pure domains of mathematics, such as algebraic topology and category theory, which have rarely been used to model biology (*35*). Here, we hypothesize that future effort towards integrating these domains into biology can be the key to building a flexible and unified theory for life sciences. Indeed, we can already appreciate the unifying value of our framework by noting that the concepts and phenomena emerging from our model align with many of the proposed research directions for reaching a unified theory of biology. For instance, our method addresses the question of defining a ‘relational ontology’ (*36*) between scales through the language of compositions and factorizations. Because every behavior described by this ontology can be expressed in terms of compositions, any pair of behaviors can be compared and differentiated for the purpose of classification. Notably, this opens the door to a rigorous formalism for understanding the complexity of relations between living systems. Our formalism also recovers fundamental phenomena pertaining to biological systems such as functional specialization, local to global organization, global dynamics, emergence, flexibility, autonomy and robustness (*37*). In our model, all these phenomena emerge from two concepts that have been strongly anchored in the thinking of biology, the concept of compartmentalization, which fundamentally relates to the origin of life and the emergence of specialized functions in living systems (*38*), and the concept of non-equilibrium state, which has often been associated with the phenomenon of self-organization (*39*). While we can note that previous mathematical work has used single-scale compartmentalization to model reaction-diffusion pathways in cells (*40*), there still exists a gap in relating multi-scale compartmentalization and thermodynamics-inspired non-equilibrium principles. For instance, Kitano (*41*) pointed out, a decade ago, that theories on non-equilibrium dissipative systems had so far ‘not [taken] into account the heterogeneity and structured nature of biological systems’, which, to our knowledge, is still the case. By the present work, we hope to open the doors towards filling these gaps. However, further work is needed to thoroughly show that our framework can be a satisfying answer to unifying life sciences within a single language. In particular, the question of integrating specific single-scale models within our framework remains to be explored in detail. Additionally, our model could be further completed by incorporating other fundamental principles related to multi-scale ecosystems such as recombination of information and reproduction.

Even though our model does not treat the question of reproduction (or self-duplication), our implementation IntCyt still behaves in ways that remind us of early development in living beings. Notably, elementary cellular-like processes such as specialization, fusion and fission allow us to recover mechanisms that arise during development such as cell movement, cellular differentiation, dynamic compartmentalization and inter-cellular communication (*42*). In this paper, we showed that these mechanisms can be used to memorize, organize, abstract and reconstruct reference machine-learning datasets such as `MNIST`, `fashion-MNIST` and `DREAM3`. It is also worth noting that a byproduct of these mechanisms is the noticeable architectural plasticity of IntCyt, which is a feature that seems to be rarely associated with learning algorithms. Indeed, while concepts such as specialization, communication and compartmentalization often underlie current brain-inspired learning algorithms (*9, 10*), the concept of structural plasticity seems to have never been used in deep learning methods (*43*). One reason for this lack of use in that context could be that ANNs have so far been defined as static architectures, especially regarding the number of nodes and layers that they should contain (*44*). In contrast, advances in understanding the brain show that structural plasticity does occur to reinforce learning and memory (*14, 15*). Thus, our algorithm appears to be one of the unique learning algorithms that addresses the possible advantages of structural plasticity, which we could hypothesize as one of the main reasons why brain cognition is so different from current learning algorithms.

To better emphasize our point, the following two paragraphs show how structural plasticity, as exhibited by our proposed algorithm IntCyt, could be the key to solving challenging problems of deep learning, such as the lottery ticket hypothesis and the vanishing or exploding gradient problems (*45, 46*).

First, the *lottery ticket hypothesis* states that any dense^2^ randomly-initialized ANN contains a sub-network that performs similarly to its super-network (*45*). As of today, finding this sub-network is a difficult task because predicting the converging state of an ANN from its initial state amounts to making a series of guesses. On the other hand, living systems do not attempt to make such guesses and, instead, use developmental mechanisms in order for their brains to attain an efficient wiring configuration (*47*). If we were to copy brains to improve ANNs, we could imagine finding the sub-network of the lottery ticket hypothesis dynamically as an emergent property of development-like mechanisms, in much the same way as IntCyt works. Such a plasticity would surely reduce the training time as only the relevant cells would be taken into account, while letting the other cells be used for other tasks. Note that in the case of IntCyt, the tree-shaped architecture resulting from its structural plasticity has the advantage to organize learned concepts by categories. As shown in the results, this reorganization leads to reinforcing the learning of the concepts in the higher levels of the tree.

Second, the structural plasticity of IntCyt is strongly intertwined with the local nature of the dynamics governing its cells. Importantly, this localness arises both at a memory and tuning level in IntCyt. Here, we argue that such local properties can be beneficial to avoiding tuning problems such as the *vanishing or exploding gradient problems*. Recall that these two problems state that updates in the first layers of a deep ANN tend to be either too small or too large to really make the network learn effectively (*46*). In practice, this phenomenon arises due to the back-propagation algorithm, which computes update values as a function of previously computed update values in the outward layers. The drawback of such a computation lies in its global nature, which has the consequence of significantly amplifying or reducing the update values computed for the inward layers – thus leading to the so-called vanishing or exploding gradient problems. Even though certain machine learning architectures, such as LSTM (*48*), correct for the vanishing gradient problem, most, if not all, still rely on the back-propagation algorithm and continue to be susceptible to suffering from exploding gradients in the first layers (*49*). In contrast, brains are more compartmentalized and rather back-propagate information locally, independently of the overall performance of the whole network (*50*). In this sense, IntCyt is closer to the brain as it optimizes its cells locally through a loss function that only takes into account local performance. As a result, our local approach gives us a better control on the learning rate at a single-cell level, which can be used to prevent vanishing or exploding gradients.

Interestingly, the local and unsupervised nature of IntCyt tackles another challenge of deep learning called the *credit assignment problem* (*51*). This problem refers to the ability of understanding how a local change, within a system, can affect or be responsible for the learning of a given task. In the case of IntCyt, cells are tuned through local loss functions in an unsupervised manner. This allows us to understand the mechanisms by which a cell specializes independently of the performance of the whole network. In particular, this means that we can accurately evaluate the contribution of each cell with respect to the learning performance of the network. Note that the accuracy of such an evaluation can be further reinforced by the direct interpretability of the data memorized by each cell. Indeed, our model allows us to interpret the information stored in each cell of IntCyt as chemical substances whose variations positively correlate to the variability of the inputs. This makes the memory stored in the cells of IntCyt comparable to the inputs and allows us to extend any interpretation model holding for the inputs to the memory of the cells themselves.

Furthermore, the compatibility of the information stored in the cells with the inputs offers a wide range of possible refinements for IntCyt. For instance, such a compatibility suggests that we could make the memory of each cell interact with the inputs in much the same way as convolution neural network “filters” (CNN filters) interact with inputs through componentwise multiplication (*52*). However, in contrast to IntCyt, the information learned by CNN filters is usually not directly accessible and needs to be processed using heuristics before being visible. As a result, our model has the potential to provide more comprehensible filters, offering possibilities of direct interpretability and live interactions between the user and the machine. Importantly, our notion of filters would not be encoded by a single vector, as in CNNs, but by a collection of vectors integrated into a tree-shaped database, in which the vectors would be compartmentalized with respect to their filtering functions. Remarkably, this tree-shaped database can also be found at a structural level in the brain: while filters usually play the role of intra-neuronal connections, IntCyt instances^3^ seem to more appropriately match dendritic trees. A compelling reason for such a comparison is that IntCyt is endowed with structural plasticity, which is an inherent property of dendritic spines (*15*). In fact, because IntCyt instances also learn and abstract concepts, we could alternatively regard them as satisfiable models for whole neurons. If used as such in neural-like architectures, we could expect IntCyt instances to lead to much more flexible convolutional learning algorithms as their associated filters would contain a range of filtering concepts that could be dynamically compared, classified and restructured.

To conclude, the learning abilities of IntCyt are the result of a set of novel learning rules inspired from biology, a new type of unsupervised loss function borrowed from operad theory and a new set of architectural dynamics allowing structural plasticity. Interestingly, these three types of improvements also align with the types of ameliorations that Richards et al. (*53*) claim to be necessary to be addressed in order to tackle the challenging question of reducing the gap between deep learning algorithms and brain-like cognition. While we showed that IntCyt was able to outperform well-known unsupervised machine learning algorithms only from the point of view of the `DREAM3` challenge, we can envision countless extensions and additional applications of IntCyt to a plethora of machine learning datasets. We imagine that domain-specific tuning will be needed for many of these applications, similar to the tuning of neural networks over decades, which were greatly underperforming other types of machine learning methodologies, but have since come to dominate many domains of machine intelligence. While many new, exciting, and innovative architectures of deep neural networks have been proposed, and widely extended, the basic building blocks of neural networks and deep learning have seen relatively little change, hindering the overall space of models that they can be expanded into. By introducing Operads here, as a fundamentally novel operational unit for learning networks, we provide additional avenues of growth and innovation to the field of machine intelligence, of which we only scratch the surface in this first paper.

## Supporting information

Supplemental figure and mathematical method

Documentation (including computational method)

## Acknowledgments

This project was supported by A patent was filed through MIT for the implementation of Int - Cyt. We thank Anna Kondylis for her help in the process of creating preliminary visualizations for our software IntCyt and Patricia Purcell for proofreading an earlier version of the article. We also thank members of the Kellis lab for useful discussions and comments on metabolism and artificial intelligence.

The following contribution description is inspired from CRediT (Contributor Roles Taxonomy; see https://casrai.org/credit/). The contributors are listed in the alphabetical order. Project design: LZA, RT; Conceptualization: AL, LZA, RT; Mathematical methodology:

RT with feedback from LZA; Software: RT, SR with feedback from LZA & MK; Visualization: AL, MK, RT; Supervision: LZA, MK, RT; Writing of original draft: RT; Writing of review & editing: AL, BK, KG, LZA, MK; Funding acquisition: LZA, MK, RT; Project administration: LZA, MK, RT. All authors reviewed the manuscript. The authors declare no competing interest.

## Supplementary materials

Figure S1

Tables S1-S3

Materials and Methods

The code and the documentation for IntCyt is hosted at the following github address: https://github.com/remytuyeras/intcyt-library.

## Footnotes

1 Through the action of a demon.

2 whose successive layers of neurons are all fully connected

3 Here, an instance refers to the IntCyt algorithm seen as a distinct function.

