## Supplemental figure and mathematical method for "Cellular intelligence: dynamic specialization through non-equilibrium multi-scale compartmentalization"

#### This pdf file contains

Figure S1

Materials and Methods

The code and the documentation for INTCYT is hosted at the following github address: `https://github.com/remytuyeras/intcyt-library`.

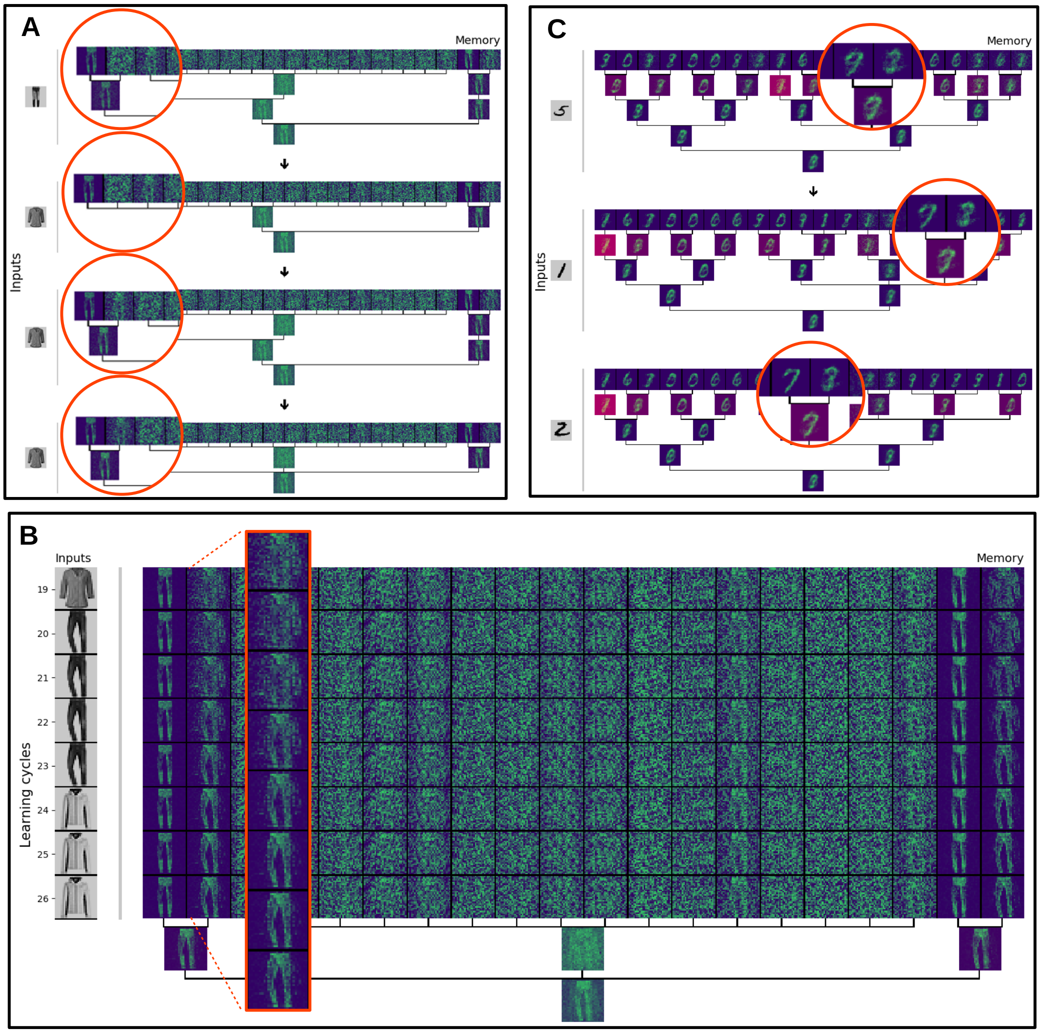
A
Memory
Inputs
C
Memory
Inputs
B
Inputs
Memory
Learning cycles
19
20
21
22
23
24
25
26

**Fig. S1.**

**Fig. S1 Learning reference machine learning datasets with IntCyt.**

(**A**) Visualization showing an example of a recruitment in the case where the inputs originate from the dataset `fashion-MNIST`. The leftmost organelle integrates a nearby cell and recruits the organelle of that cell with similar information in a new compartment. (**B**) Visualization showing an example of a communication between cells right after the cycles shown in panel **A**. Specifically, the organelle containing the image of a pair of pants influences its neighbor organelle to change its content and specialize towards a similar pair of pants. This communication is also influenced by the environment, which shows an image of a pair of pants to the two organelles. (**C**) Visualization showing that communication between organelles can lead to smoothening the abstractions made by IntCyt. Specifically, the memorized image of a 9 is slowly turned into a 7. The cycles shown in the figure correspond to the cycles 750, 1150 and 1250 of the simulation shown in **Fig. 5C**. By using such communication mechanisms, IntCyt avoids over-fitting its inputs.

### METHOD FOR: "CELLULAR INTELLIGENCE: DYNAMIC SPECIALIZATION THROUGH NON-EQUILIBRIUM MULTI-SCALE COMPARTMENTALIZATION"

#### 1. Model

1.1. **Compartmentalization.** In this section, we define the concept of a cell as a finite collection of real numbers representing various amounts of chemicals or energetic components. Note that the notations $\mathsf{res}(c)$, $\mathsf{org}(c)$ and $\mathsf{cyt}(c)$ used in the main text will only be introduced in this text from section 1.4. For the sections preceding section 1.4 – including the present one – we will use more explicit notations in order to facilitate the definition of certain operations such as compositions (Definition 1.6) and the statements of Theorem 1.8 and Theorem 1.17. These notations will also be relevant for the discussion of section 3.

**Convention 1.1** (Notation)**.** For every non-negative integer $n$, we will denote by $[n]$ the set of integers ranging from 1 to $n$. If $n$ is zero, then $[n]$ is empty. We will also denote by $\mathbb{R}$ the set of real numbers and by $\mathbb{R}_+$ the set of non-negative real numbers.

**Definition 1.2** (Cells)**.** Let $N$ be a positive integer. A *cell* of *dimension* $N$ consists of

- a non-negative integer $n \geq 0$, called the *number of organelles*;
- a non-negative real number $C$, called the *residual*;
- an element $x$ of $\mathbb{R}^N$, called the *cytosolic content*;
- a $n$-tuple $(x_i)_{i\in[n]}$ of elements in $\mathbb{R}_+^N$, where each $x_i$ is called the *i-th organelles*;

*Remark* 1.3 (Empty cell). By definition, any cell $c = (n, C, x, (x_i)_i)$ for which the natural number $n$ is equal to 0 must be equipped with an empty tuple $(x_i)_{i\in[n]}$ – meaning that the tuple does not contain any vector $x_i$.

**Example 1.4** (A model for cellular structures)**.** The following pictures represent two cells of dimension 3. The leftmost cell has 2 organelles encoded by the vectors $x_1 = (9, 7, 0)$ and $x_2 = (8, 0, 1.2)$ and its cytosolic content is encoded by the vector $x = (.2, 0, 1.4)$. The residual, indicated in brackets, is equal to 0.

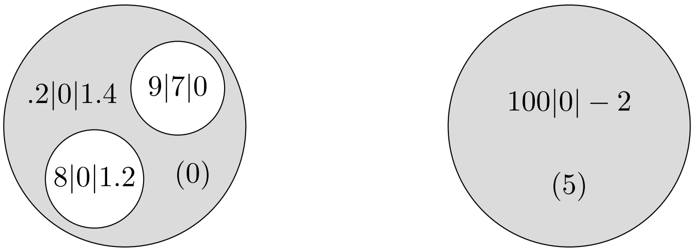


The rightmost cell does not possess any organelle (*i.e.* $n = 0$ and $(x_i)_i$ is empty). However, the cell has a non-trivial cytosolic content encoded by the vector $x = (100, 0, -2)$. The residual is equal to 5.

**Definition 1.5** (The set of cells as a topological space)**.** For convenience, we will denote by $\mathcal{C}^N(n)$ the set of cells of dimension $N$ with $n$ organelles. We deduce from Definition 1.2 that this set is in bijection with the topological space $\mathbb{R}_+ \times \mathbb{R}^N \times \mathbb{R}_+^{N\times n}$.

We now want to be able to nest cells within the organelles of other cells. To this end, we introduce a composition operation modeled on the type of composition used for operads, as shown in the picture below. Our interest in defining such a nesting operation comes from our desire to better conceptualize the sequence of mechanisms that allows cells to find optimal compartmentalization configurations through their life. As will be seen, we will model compartment formation in terms of a decomposition problem (*i.e.* a factorization problem) whose solutions are characterized in Theorem 1.8.

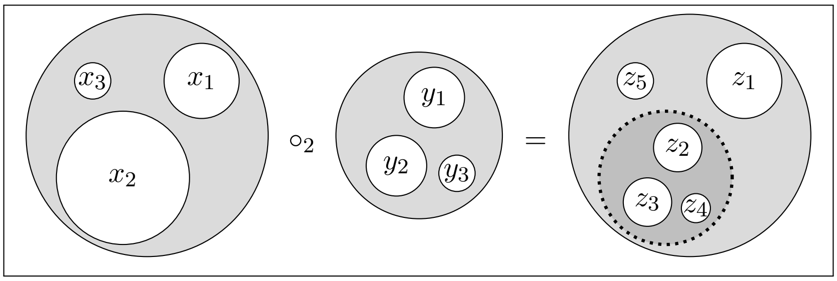


The previous picture, which is often the type of pictures used in operad theory, can also be given a more biological representation. Specifically, the dotted line shown in the previous picture can be seen as a disassembling cell membrane allowing its content to be mixed with the outer environment. This type of mechanisms, for instance, happens during cell division when the nucleus disassembles to permit the DNA material to interact with the cytoskeleton in order to be segregated in opposite parts of the cell.

**Definition 1.6** (Composition)**.** Let $c = (n, C, x, (x_i)_i)$ and $d = (m, D, y, (y_j)_j)$ be two cells of dimension $N$ and let $k \in [n]$. The composition of $c$ with $d$ at organelle $k \in [n]$ is defined by the following cell:

$$c \circ_k d = (n + m - 1, C + D, x + y, \overline{x} \circ_k \overline{y})$$

where $\overline{x} \circ_k \overline{y} = (x_1, \dots, x_{k-1}, y_1, y_2, \dots, y_m, x_{k+1}, \dots, x_n)$.

**Example 1.7** (Composition)**.** Equation (1.1) shows an example of a composition of two cells at the second organelle $x_2 = (11, 13)$ of the leftmost cell. The resulting cell, shown on the right-hand side, possesses the two organelles of the middle cell, which replace the second organelle of the leftmost cell, as well as the first organelle $x_1 = (0, 1)$ of the leftmost cell.

(1.1)

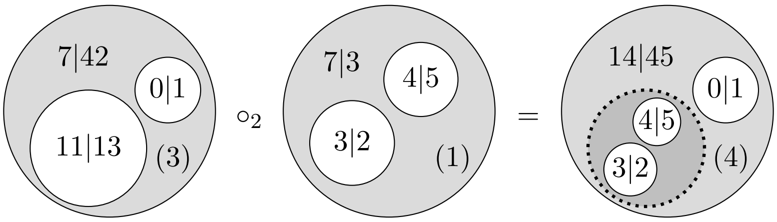


Note that composing cells does not require the cells to satisfy any condition. However, we will later see that the work produced by the cells can be affected by the way the cells are composed between each other. In section 1.4, we will introduce the concept of fitness to distinguish certain types of optimal compositions from others.

Definition 1.6 is simple enough to enable us to solve factorization problems (see Theorem 1.8). Factorizations will be used in section 1.6 to define homeostasis.

**Theorem 1.8** (Factorization problem)**.** *Let $c = (n, C, x, (x_i)_i)$ be a cell of dimension $N$ and let $k \in [n]$. The equation $X \circ_k Y = c$ holds, if and only if, there exist a non-negative integer $\mu$, a non-negative real number $\alpha$ and two vectors $\kappa$ and $\lambda$ in $\mathbb{R}_+^N$ such that the cells $Y$ and $X$ are of the following form:*

$$\begin{aligned} X &= \big(n + 1 - \mu, C - \alpha, x - \kappa, (x_1, \dots, x_{k-1}, \lambda, x_{k+\mu}, \dots, x_n)\big) \\ Y &= (\mu, \alpha, \kappa, (x_k, \dots, x_{k+\mu-1})) \end{aligned}$$

*Proof.* Directly follows from Definition 1.6. □

Below, we introduce the concept of identity cells, which will play an important role in section 2.1 (see Convention 2.10). The importance of these cells lies in their property with respect to compositions.

**Definition 1.9** (Identities)**.** For every positive integer $N$ and every vector $a$ in $\mathbb{R}^N$, we will denote the cell $(1, 0, 0, a)$ of $\mathcal{C}^N(n)$ as $\mathsf{id}_a$.

**Proposition 1.10** (Identities)**.** *For every cell $c = (n, C, x, (x_i)_i)$ be a cell of dimension $N$, the following equations hold for every vector $a$ in $\mathbb{R}^N$:*

$$c \circ_k \mathsf{id}_{x_k} = c \qquad\qquad \mathsf{id}_a \circ_1 c = c$$

*Proof.* By taking the parameters $\mu = 1$, $\alpha = 0$, $\kappa = 0$ and $\lambda = x_k$ in the factorization $X \circ_k Y = c$ of Theorem 1.8, we obtain an equation $c \circ_k \mathsf{id}_{x_k} = c$. Similarly, by taking $\mu = n$, $\alpha = C$, $\kappa = x$ and $\lambda = a$ in the factorization $X \circ_k Y = c$ of Theorem 1.8, we obtain the equation $\mathsf{id}_a \circ_1 c = c$ for any vector $a$ in $\mathbb{R}^N$. □

1.2. **Shuffling organelles.** In the sequel, we will prove results that are invariant up to shuffling the organelles of the cells. As a result, we will only prove the result for one configuration and all the other configuration will follow from the invariance by symmetries. To be able to say when certain results are invariant by symmetry, we define, below, the symmetric action of a bijection on a cell.

**Definition 1.11** (Symmetric action)**.** Let $c = (n, C, x, (x_1, x_2, \ldots, x_n))$ be a cell of dimension $N$. For every bijection $\sigma : [n] \to [n]$, we will denote by $\sigma \odot c$ the cell $(n, C, x, (x_{\sigma(1)}, x_{\sigma(2)}, \ldots, x_{\sigma(n)}))$.

**Proposition 1.12** (Symmetric action)**.** *Let $c \in \mathcal{C}^N(n)$ and $d \in \mathcal{C}^N(m)$. For every index $k \in [n]$ and every bijection $\sigma : [n] \to [n]$, the following identity holds: $(\sigma \odot c) \circ_k d = \sigma \odot (c \circ_{\sigma^{-1}(k)} d)$.*

*Proof.* This is a direct application of Definition 1.11 and Definition 1.6. □

**Convention 1.13** (Symmetric action on vectors)**.** In addition to acting on cells, we shall also need to make bijection act on vectors, as is already the case in Definition 1.11. In this respect, for every bijection $\sigma : [n] \to [n]$ and every vector $v = (v_1, v_2, \ldots, v_n)$, we will denote by $\sigma \odot v$ the vector $(v_{\sigma(1)}, v_{\sigma(2)}, \ldots, v_{\sigma(n)})$.

1.3. **Simultaneous compositions.** So far, our cell compositions have only dealt with one organelle at a time. However, it is possible to define a composition operation that simultaneously composes cells at multiple organelles. A very intuitive way to do so is shown in Definition 1.14.

**Definition 1.14** (Simultaneous composition)**.** Let $c$ be a cell in $\mathcal{C}^N(n)$ and let $d$ denote a $n$-tuple $(d_1, \ldots, d_n)$ of cells $d_k$ in $\mathcal{C}^N(m_k)$. We define the simultaneous composition of $c$ with $d$ as the following cell in $\mathcal{C}^N(m_1 + m_2 + \cdots + m_n)$:

$$c \circ d = ((\ldots(c \circ_n d_n) \circ_{n-1} d_{n-1} \circ_{n-2} \ldots) \circ_1 d_1)$$

**Example 1.15** (Composition)**.** Below, equation (1.2) shows an example of a simultaneous composition of cells. The cell given in the first component of the tuple is composed at the first organelle $x_1 = (0, 1)$ of the lefmost cell and the cell given in the second component of the tuple is composed at the second organelle $x_2 = (11, 13)$ of the leftmost cell. The resulting cell, shown on the right-hand side, possesses all the organelles shown in the tuple.

(1.2)
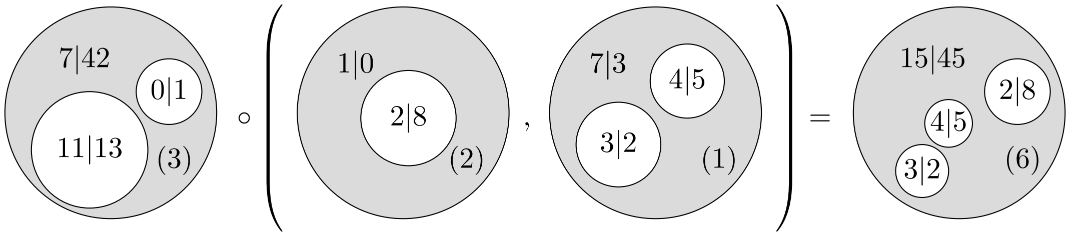


The reader may have noticed the composition of the second cell of the tuple does not affect the composition of the first cell of the tuple, which suggests that that cells could be composed in any order provided that we figure out the index at which the composition occur (see Theorem 1.17)

The following proposition uses the symmetric action on vectors defined in Convention 1.13.

**Proposition 1.16.** *Let $c$ be a cell in $\mathcal{C}^N(n)$ and let $d$ denote a $n$-tuple $(d_1,\dots,d_n)$ of cells $d_k$ in $\mathcal{C}^N(m_k)$. For every bijection $\sigma : [n] \to [n]$, the following identity holds: $(\sigma \odot c) \circ d = \sigma \odot (c \circ (\sigma^{-1} \odot d))$.*

*Proof.* This is a direct application of Definition 1.14 and Proposition 1.16. □

The order of the composition chosen in Definition 1.14 let us wonder whether a different order would provide a different composition. The answer is negative and is shown by Theorem 1.17, which states that the composed cells can be permuted, provided that the index of the composition is shifted accordingly.

**Theorem 1.17** (Compositional properties)**.** *Let $c \in \mathcal{C}^N(n)$, $d \in \mathcal{C}^N(m)$ and $e \in \mathcal{C}^N(p)$ be three cells of dimension $N$ and let $k \in [n]$ and $r \in [m+n-1]$. The following equations hold.*

$$(c \circ_k d) \circ_r e = \begin{cases} (c \circ_r e) \circ_{k+p-1} d & \textit{if } r < k \\ c \circ_k (d \circ_{r-k+1} e) & \textit{if } k \leq r \leq k+m-1 \\ (c \circ_{r-m+1} e) \circ_k d & \textit{if } k+m \leq r \end{cases}$$

*Proof.* Directly follows from Definition 1.6 since the addition of vectors in $\mathbb{R}^N$ is associative and commutative and so does the nesting of tuples. □

In section 1.6, simultaneous compositions will play a important role in defining homeostasis.

**Theorem 1.18** (Factorization problem)**.** *Let $c = (\ell, C, x, (x_i)_i)$ be a cell of dimension $N$. The equation $X \circ (Y_1,\dots,Y_n) = c$ holds, if and only if, for every $i \in [n]$, there exist a non-negative integer $\mu_i$, a non-negative real number $\alpha_i$ and two vectors $\kappa_i$ and $\lambda_i$ in $\mathbb{R}^N_+$ such that the equation $\ell = \sum_{i=1}^n \mu_i$ holds and the cells $Y_1,\dots,Y_n$ and $X$ are as follows:*

$$\begin{aligned} X &= \big(n, C - \textstyle\sum_{i=1}^n \alpha_i, x - \sum_{i=1}^n \kappa_i, (\lambda_i)_i\big) \\ Y_i &= (\mu_i, \alpha_i, \kappa_i, (x_{L(i)},\dots,x_{L(i)+\mu_i-1})) \end{aligned}$$

*where, for every $i \in [n]$, we let $L(i) := \sum_{k=1}^{i-1} \mu_k$.*

*Proof.* Follows from Definition 1.14 and $n$ applications of Theorem 1.8. □

**Convention 1.19** (Notations)**.** For the same notations as those used in the statement of Proposition 1.18, let us denote the vectors $(\mu_i)_i$, $(\alpha_i)_i$, $(\kappa_i)_i$ and $(\lambda_i)_i$ as $\mu$, $\alpha$, $\kappa$ and $\lambda$, respectively. From now on, we will denote the solutions $Y_1,\dots,Y_n$ and $X$ of the equation $X \circ (Y_1,\dots,Y_n) = c$ as follows:

$$\begin{aligned} \mathsf{left}(c)(\alpha,\kappa,\lambda) &:= X = \big(n, C - \textstyle\sum_{i=1}^n \alpha_i, x - \sum_{i=1}^n \kappa_i, (\lambda_i)_i\big) \\ \mathsf{right}(c)(\mu,\alpha,\kappa)_i &:= Y_i = (\mu_i, \alpha_i, \kappa_i, (x_{L(i)},\dots,x_{L(i)+\mu_i-1})) \end{aligned}$$

The vector $(Y_1,\dots,Y_n)$ will therefore be denoted as $\mathsf{right}(c)(\mu,\alpha,\kappa)$.

1.4. **Compositional fitness.** We introduce the notion of fitness to distinguish those cells that perfectly compose for the purpose of homeostasis (see section 1.6). The idea is that the organelle at which the composition is done should be equal to the content of the cell by which it is replaced.

**Convention 1.20** (Notation)**.** From now on, for more clarity, we will tend to refer to the structure of a cell $c = (n, C, x, (x_i)_i)$ through standard notations. Specifically, for such a cell $c$, we will denote the quantity $C$ as $\mathsf{res}(c)$, the vector $x$ as $\mathsf{cyt}(c)$ and the tuple $(x_i)_i$ as $\mathsf{org}(c)$ such that the vector $x_i$ will be referred to as $\mathsf{org}(c)_i$ for every $i \in [n]$.

**Definition 1.21** (Content)**.** For every cell $c \in \mathcal{C}^N(n)$, we define the *content of $c$* as the quantity:

$$K(c) := \begin{cases} \mathsf{cyt}(c) + \sum_{i=1}^n \mathsf{org}(c)_i & \text{if } n > 0 \\ \mathsf{cyt}(c) & \text{if } n = 0 \end{cases}$$

**Definition 1.22** (Fitness)**.** Let $c \in \mathcal{C}^N(n)$ and $d \in \mathcal{C}^N(m)$ be two cells and let $k \in [n]$. We define the *fitness* $\delta_k(c,d)$ of the cell $d$ for the $k$-th organelle of $c$ as the vector of $\mathbb{R}^N$ defined by the following formula.

$$\delta_k(c,d) := \mathsf{org}(c)_k - K(d)$$

**Convention 1.23** (Fitting)**.** A cell $d$ will be said to *fit* the $k$-th organelle of a cell $c$ if the fitness vector $\delta_k(c,d)$ is equal to the vector 0 in $\mathbb{R}^N$.

**Proposition 1.24** (Contents and fitness)**.** *Let $c \in \mathcal{C}^N(n)$ and $d \in \mathcal{C}^N(m)$ be two cells and let $k \in [n]$. The identity $\delta_k(c,d) = K(c) - K(c \circ_k d)$ holds.*

*Proof.* It suffices to verify that the equation of the statement holds. We will do so by simplifying the terms in the difference $K(c \circ_k d) - K(c)$. By definition, we have the following equation.

$$K(c \circ_k d) - K(c) = \mathsf{cyt}(c) + \mathsf{cyt}(d) + \sum_{i \in [n]-\{k\}} \mathsf{org}(c)_i + \sum_{j=1}^{m} \mathsf{org}(d)_j - (\mathsf{cyt}(c) + \sum_{i=1}^{n} \mathsf{org}(c)_i)$$

Simplifying the terms of the right-hand side gives the following series of equations

$$K(c \circ_k d) - K(c) = \mathsf{cyt}(d) + \sum_{j=1}^{m} \mathsf{org}(d)_j - \mathsf{org}(c)_k = K(d) - \mathsf{org}(c)_k = -\delta_k(c,d)$$

The last equality shows that the identity of the statement holds. □

*Remark* 1.25 (Content of simultaneous compositions)*.* In terms of a simultaneous composition, Proposition 1.24 implies that for every cell $c \in \mathcal{C}^N(n)$ and $n$-tuples $d = (d_1, \dots, d_n)$ of cells of dimension $N$, the following equation holds.

$$K(c \circ d) = K(c) - \sum_{k=1}^{n} \delta_k(c, d_k)$$

1.5. **Specialization theorem.** In this section, we prove our most important theorem (Theorem 1.38), which is explains the learning capacities of the algorithm presented in the main text (see section 2).

**Definition 1.26** (Sum)**.** For any vector $v = (v_1, \dots, v_N)$ in $\mathbb{R}^N$, we will denote the sum of all the components of $v$ as $\mathcal{S}(v)$ (as shown below).

$$\mathcal{S}(v) := \sum_{u=1}^{N} v_u$$

For the sake of conciseness, for any function $f : U \to \mathbb{R}^N$ sending an element $x \in U$ to a vector $(f_1(x), \dots, f_N(x))$, we will shorten the composition $\mathcal{S}(f(x)) = \sum_{u=1}^{N} f_u(x)$ as $\mathcal{S}f(x)$.

**Definition 1.27** (Inputs)**.** For every positive natural number $N$ and real number $A$, we will denote by $\Delta_N(A)$ the space defined by the following specification.

$$\{p \in \mathbb{R}_+^N \mid \mathcal{S}(p) \leq A\}$$

This space describes a higher dimensional pyramid in $\mathbb{R}_+^N$.

**Definition 1.28** (Empty and non-empty cells)**.** A cell $c \in \mathcal{C}^N(n)$ will be said to be *empty* if the sum $\mathcal{S}K(c)$ is equal to 0. Conversely, a cell $c \in \mathcal{C}^N(n)$ will be said to be *non-empty* if $\mathcal{S}K(c) \neq 0$.

*Remark* 1.29 (Entropy)*.* At the end of this section, we will interpret the concept of entropy in every non-empty cell $c \in \mathcal{C}^N(n)$ as a level of homogenization of its associated numerical values. To give an example, a cell in which the proportions $\mathsf{cyt}(c)_u/\mathcal{S}K(c)$ and $\mathsf{org}(c)_{i,u}/\mathcal{S}K(c)$ are relatively close to each other for every $u \in [N]$, as illustrated below, on the right, will be associated with a high entropy.

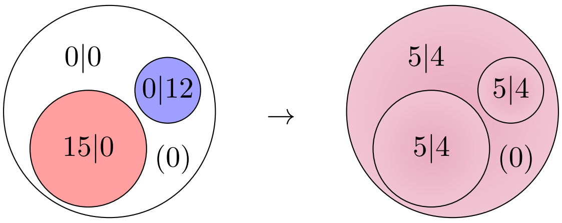

In what follows, we will use the organelles of a cell to reduce its associated entropy. Specifically, we will use the organelles of the cell to either keep specific quantities together or to isolate them from others (with which they should not interact). In Definition 1.30, we associate cells with an action operation that promotes this type of behavior through optimization mechanisms such as gradient descent, cell fusion and cell fission.

**Definition 1.30** (Action of a cell)**.** The *action of a non-empty cell* $c \in \mathcal{C}^N(n)$ on a vector $a = (a_i)_{i\in[n]}$ in $\Delta_N(A)^n$ is defined by the following vector:

$$c \cdot a = \Big(\sum_{i=1}^{n} a_{i,u}\frac{\mathsf{org}(c)_{i,u}}{\mathcal{S}K(c)}\Big)_{u\in[N]}$$

Note that, if the sum $\mathcal{S}\mathsf{cyt}(c)$ is positive, then the quantity $\mathsf{org}(c)_{i,u}$ is always less than or equal to $\mathcal{S}K(c)$ (see the formla given in Definition 1.21). As a result, the ratio of $\mathsf{org}(c)_{i,u}$ over $\mathcal{S}K(c)$ can be interpreted as a conditional probability, allowing us to interpret the action of a cell as a vector of conditional expected values. The following proposition formalizes this idea in terms of a stability property.

**Proposition 1.31** (Stability)**.** *For every non-empty cell* $c \in \mathcal{C}^N(n)$ *and vector* $a \in \Delta_N(A)^n$*, the action* $c\cdot a$ *belongs to* $\Delta_N(A - \lambda A)$ *where* $\lambda = \mathcal{S}\mathsf{cyt}(c)/\mathcal{S}K(c)$*.*

*Proof.* The proof follows from the series of relations displayed below.

$$\mathcal{S}(c\cdot a) = \sum_{u=1}^{N}\sum_{i=1}^{n} a_{i,u}\frac{\mathsf{org}(c)_{i,u}}{\mathcal{S}K(c)} \leq A\sum_{u=1}^{N}\sum_{i=1}^{n}\frac{\mathsf{org}(c)_{i,u}}{\mathcal{S}K(c)} = A\frac{\mathcal{S}K(c) - \mathcal{S}\mathsf{cyt}(c)}{\mathcal{S}K(c)}$$

□

*Remark* 1.32 (Spontaneous chemical reactions). For every non-empty cell $c$ of dimension $N$, the quantity $\mathcal{S}\mathsf{cyt}(c)$ may not always be positive. Indeed, the expression $\mathcal{S}\mathsf{cyt}(c)$ can be the sum of both positive and negative values, as illustrated below.

$$\mathcal{S}\mathsf{cyt}(c) = a + b - c + d - e - f + g$$

Our interest in modeling biology motivates the following interpretation: if the quantity $\mathcal{S}\mathsf{cyt}(c)$ is negative, then the cytosolic content of $c$ is missing resources. With such an interpretation, we could ask whether one can compensate this lack by looking for resources elsewhere in the cell. A solution could be the residual $\mathsf{res}(c)$, which we would like to see as an energetic resource.

More specifically, the idea is that, if the inequality $\mathsf{res}(c) + \mathcal{S}\mathsf{cyt}(c) \geq 0$ holds, there are enough positive values both in the residual and the cytosol to compensate the negative values in the cytosol. If we had to interpret this statement chemically, we could see this compensation as an overall chemical reactions happening in the cytosol, taking elements with positive values to create elements with negative values. The chemical reaction would go as follows:

$$\mathsf{res}(c)\mathsf{Energy} + a\mathsf{A} + b\mathsf{B} + g\mathsf{G} \to c\mathsf{C} + e\mathsf{E} + f\mathsf{F} + \mathcal{S}\mathsf{cyt}(c)\mathsf{Energy}$$

As shown above, the quantities $\mathsf{res}(c)$ and $\mathcal{S}\mathsf{cyt}(c)$ play the role of the energy being used and released by the reaction. In section 2.2, we will use this kind of logic to define the 'cleaning' of a cell (Definition 2.13), in which we will increment the variable $\mathsf{res}(c)$ by the value $\mathcal{S}\mathsf{cyt}(c)$ and set the variable $\mathsf{cyt}(c)$ to zero.

In the rest of this section, our goal is to define an operator that will allow us to control what we could see as the level of entropy of a cell (see Remark 1.29).

**Convention 1.33** (Notations)**.** For every collection $\{m_k\}_{k\in[n]}$ of non-negative integers and every collection $(a_k)_{k\in[n]}$ of vectors $a_k = (a_{k,i})_{i\in[m_k]} \in \Delta_N(A)^{m_k}$, we will denote by $a_1 \odot a_2 \odot \cdots \odot a_n$ the vector concatenation

$$(a_{1,1}, \dots, a_{1,m_1}, a_{2,1}, \dots, a_{2,m_2}, \dots, a_{n,m_n}, \dots, a_{n,m_n})$$

which lives in the space $\Delta_N(A)^L$ where $L = \sum_{k=1}^{n} m_k$

**Definition 1.34** (Algebra operator)**.** Let $c$ and $d_1,\dots,d_n$ be non-empty cells of dimension $N$. For every collection $a=(a_k)_{k\in[n]}$ of vectors $a_k\in\Delta_N(A)^{m_k}$, we will denote by $U(c,d)(a)$ the following difference:

$$(c\circ d)\cdot(a_1\odot a_2\odot\cdots\odot a_n)-c\cdot(d_1\cdot a_1\odot d_2\cdot a_2\odot\cdots\odot d_n\cdot a_n)$$

The quantity $U(c,d)(a)$ given in Definition 1.34 assesses the effect of creating compartments $d_1,\dots,d_n$ within a cell $c\circ d$ on the action operation. In the remainder of the article, we will refer to the function $a\mapsto U(c,d)(a)$ as the *algebra operator* of $c$ and $d$.

*Remark* 1.35 (Invariance by symmetry). Let $c\in\mathcal{C}^N(n)$, $a$ be a vector in $\Delta_N(A)^n$ and $\sigma:[n]\to[n]$ be a bijection. We can easily deduce from Definition 1.30 that the equation $(\sigma\odot c)\cdot a=c\cdot(\sigma^{-1}\odot a)$ holds. It then follows from Proposition 1.16 that the following identity holds.

$$\Big((\sigma\odot c)\circ(\sigma\odot d)\Big)\cdot(\sigma\odot a)=\Big(\sigma\odot(c\circ d)\Big)\cdot(\sigma\odot a)=c\cdot a$$

In the same fashion, we can deduce the following equation, which says that the algebra operator is invariant by symmetry.

$$U(\sigma\odot c,\sigma\odot d)(\sigma\odot a)=U(c,a),$$

This fact will be used in Theorem 1.58, in which we will show that the statement holds for one index configuration and deduce that the other index configurations hold after permutation along bijections $\sigma:[n]\to[n]$.

**Proposition 1.36** (Formula for the algebra operator)**.** *Let $c$ be a non-empty cell in $\mathcal{C}^N(n)$ and $d=(d_1,\dots,d_n)$ be a $n$-tuple of non-empty cells $d_k$ in $\mathcal{C}^N(m_k)$. For every collection $a=(a_k)_{k\in[n]}$ of vectors $a_k\in\Delta_N(A)^{m_k}$, the following identity holds:*

$$U(c,d)(a)=\left(\sum_{k=1}^{n}\Big(\frac{1}{\mathcal{S}K(c\circ d)}-\frac{\mathsf{org}(c)_{k,u}}{\mathcal{S}K(c)\mathcal{S}K(d_k)}\Big)\Big(\sum_{i=1}^{m_k}a_{k,i,u}\mathsf{org}(d_k)_{i,u}\Big)\right)_{u\in[N]}. \tag{1.3}$$

*If, for every $k\in[n]$, the identity $\mathcal{S}\delta_k(c,d_k)=0$ holds, then we obtain the following expression:*

$$U(c,d)(a)=\left(\sum_{k=1}^{n}\frac{(d_k\cdot a_k)_u}{\mathcal{S}K(c)}\Big(\mathcal{S}\mathsf{org}(c)_k-\mathsf{org}(c)_{k,u}\Big)\right)_{u\in[N]}. \tag{1.4}$$

*Proof.* By Definition 1.14 and Definition 1.30, we have the following formula.

$$(c\circ d)\cdot(a_1\odot a_2\odot\cdots\odot a_n)=\left(\sum_{k=1}^{n}\sum_{i=1}^{m_k}\frac{a_{k,i,u}\mathsf{org}(d_k)_{i,u}}{\mathcal{S}K(c\circ d)}\right)_{u\in[N]}$$

Similarly, by applying Definition 1.30 twice, we obtain the following formula.

$$c\cdot(d_1\cdot a_1\odot d_2\cdot a_2\odot\cdots\odot d_n\cdot a_n)=\left(\sum_{k=1}^{n}\frac{\mathsf{org}(c)_{k,u}}{\mathcal{S}K(c)}\sum_{i=1}^{m_k}\frac{a_{k,i,u}\mathsf{org}(d_k)_{i,u}}{\mathcal{S}K(d_k)}\right)_{u\in[N]}$$

The difference of the two vectors gives us expression (1.3). Factorizing by the inverse of $\mathcal{S}K(d_k)$ in each summand, we obtain the expression

$$U(c,d)(a)=\left(\sum_{k=1}^{n}\Big(\frac{\mathcal{S}K(d_k)}{\mathcal{S}K(c\circ d)}-\frac{\mathsf{org}(c)_{k,u}}{\mathcal{S}K(c)}\Big)(d_k\cdot a_k)_u\right)_{u\in[N]} \tag{1.5}$$

By Remark 1.25 and Definition 1.22, the identity $\mathcal{S}\delta_k(c,d_k)=0$ holding for every $k\in[n]$ gives us the identities $\mathcal{S}K(c\circ d)=\mathcal{S}K(c)$ and $\mathcal{S}\mathsf{org}(c)_k=\mathcal{S}K(d_k)$. Using these identities in expression (1.5) gives us expression (1.4). $\square$

*Remark* 1.37 (Lowerbound for the algebra operator)*.* While the formula given in (1.3) allows the quantity $U(c,d)(a)$ to be either negative, positive or zero, the formula given in (1.4) makes the quantity $U(c,d)(a)$ non-negative. The reason for this lowerbound comes from the fact that the quantity $\mathsf{org}(c)_{k,u}$ is always smaller than the sum $\mathcal{S}\mathsf{org}(c)_k$ of positive terms $\mathsf{org}(c)_{k,1},\dots,\mathsf{org}(c)_{k,N}$.

Hence, when each cell $d_i$ of the tuple $d$ fits the organelle of $c$ with which it is associated, the algebra operator satisfies the inequality $U(c,d)(a)\geq 0$ for every vector $a$. Theorem 1.38, given below, studies the upperbounds of $U(c,d)(a)$.

**Theorem 1.38** (Specialization theorem)**.** *Let $c$ be a non-empty cell in $\mathcal{C}^N(n)$ and $d=(d_1,\dots,d_n)$ be a $n$-tuple of non-empty cells $d_k$ in $\mathcal{C}^N(m_k)$ for which the identities $\mathcal{S}\delta_k(c,d_k)=0$ hold. For every collection $a=(a_k)_{k\in[n]}$ of vectors $a_k\in\Delta_N(A)^{m_k}$, the implication*

$$U(c,d)(a)_u\leq\eta \qquad\Rightarrow\qquad \mathcal{S}\mathsf{org}(c)_k-\eta\frac{\mathcal{S}K(c)}{(d_k\cdot a_k)_u}\leq \mathsf{org}(c)_{k,u}\leq\mathcal{S}\mathsf{org}(c)_k \tag{1.6}$$

*holds for every $u\in[N]$ and $k\in[n]$ if $(d_k\cdot a_k)_u\neq 0$.*

*Proof.* The upper bounds comes from the expression $\mathcal{S}\mathsf{org}(c)_k=\mathsf{org}(c)_{k,1}+\cdots+\mathsf{org}(c)_{k,N}$, which implies that $\mathcal{S}\mathsf{org}(c)_k$ must be greater than $\mathsf{org}(c)_{k,u}$ for every $u\in[N]$ since the organelles of $c$ are all vectors of non-negative values. The lower bound can be deduced from expression (1.4), given in Proposition 1.36, as follows. First, since the inequality $\mathsf{org}(c)_{k,u}\leq\mathcal{S}\mathsf{org}(c)_k$ holds, we can use the non-negativeness of the summands to deduce the following inequalities.

$$\frac{(d_k\cdot a_k)_u}{\mathcal{S}K(c)}\Big(\mathcal{S}\mathsf{org}(c)_k-\mathsf{org}(c)_{k,u}\Big)\leq\sum_{k=1}^{n}\frac{(d_k\cdot a_k)_u}{\mathcal{S}K(c)}\Big(\mathcal{S}\mathsf{org}(c)_k-\mathsf{org}(c)_{k,u}\Big)=U(c,d)(a)_u\leq\eta$$

Since $(d_k\cdot a_k)_u$ is non-negative and, in fact, by assumption, non-zero, a simple algebra argument between the extreme ends of the previous relation finally provides the lower bound. □

*Remark* 1.39 (Specialization against entropy)*.* Theorem 1.38 says that if one tries to minimize the values of the algebra operator of a non-empty cell $c$ as well as optimize its fitness with respect to a collection of non-empty cells $d=(d_1,\dots,d_n)$ at a dimension $u$ in which the input $a$ sends a strong signal $(d_k\cdot a_k)_u$, then the component of the organelle $\mathsf{org}(c)_k$ in that dimension $u$ tries to converge to the value $\mathcal{S}\mathsf{org}(c)_k$. Interestingly, this can create a 'competition' between the variables $\mathsf{org}(c)_{k,u}$, because the quantity $\mathcal{S}\mathsf{org}(c)_k$ is the sum of all variables $\mathsf{org}(c)_{k,u}$. More specifically, if there existed two variables $\mathsf{org}(c)_{k,u_0}$ and $\mathsf{org}(c)_{k,u_1}$ that were equal to $\mathcal{S}\mathsf{org}(c)_k$, then the following inequality would hold, which is impossible when $\mathcal{S}\mathsf{org}(c)_k\neq 0$.

$$\mathcal{S}\mathsf{org}(c)_k=\mathcal{S}\mathsf{org}(c)_{k,1}+\cdots+\mathcal{S}\mathsf{org}(c)_{k,N}\geq 2\mathcal{S}\mathsf{org}(c)_k$$

Hence, the only possible scenario is when there is only one index $u_0$ for which the identity $\mathsf{org}(c)_{k,u_0}=\mathcal{S}\mathsf{org}(c)_k$ holds. In addition, all the other variables $\mathsf{org}(c)_{i,u}$, for which $u\neq u_0$ and $i\in[n]$, must then be equal to zero. In other words, the variable $\mathsf{org}(c)_{k,u_0}$ takes over all the other variables $\mathsf{org}(c)_{i,u}$.

To conclude, reducing the values of the algebra operator promotes specialization in the components $u$ whose associated signals $(d_k\cdot a_k)_u$ are the highest. Interestingly, if the values of the algebra operator can be reduced in more than one dimension, then this specialization process can occur at various components simultaneously.

1.6. **Homeostasis as a back propagation mechanism.** In the preent section, the concept of homeostasis mainly refers to the ability of a system to maintain fitness between its components.

**Definition 1.40** (Homeostasis problem)**.** Let $c\in\mathcal{C}^N(n)$ and let $d=(d_1,\dots,d_n)$ be a tuple of $n$ cells $d_k$ in $\mathcal{C}^N(m_k)$. Below, we denote $m=(m_i)_{i\in[n]}$ and $\mathsf{res}(d)=(\mathsf{res}(d_i))_{i\in[n]}$.

▷ We define a *homeostasis problem* for $(c,d)$ as a collection $\lambda=(\lambda_i)_{i\in[n]}$ of vectors in $\mathbb{R}^N_+$.

▷ We define a *homeostasis solution* for the previous homeostasis problem as a collection $\kappa = (\kappa_i)_{i\in[n]}$ of vectors in $\mathbb{R}^N$ such that, for every $i \in [n]$, the cells

$$\begin{aligned} c' &= \mathsf{left}(c \circ d)(\mathsf{res}(d), \kappa, \lambda) \\ d'_i &= \mathsf{right}(c \circ d)(m, \mathsf{res}(d), \kappa)_i \end{aligned}$$

exist and the equation $\delta_i(c', d'_i) = 0$ holds. Such a solution will be denoted as a triple $(\kappa, c', d')$ where $d'$ denotes the tuple $\mathsf{right}(c \circ d)(m, \mathsf{res}(d), \kappa) = (d'_1, \dots, d'_n)$.

**Example 1.41** (Homeostasis)**.** At a conceptual level, homeostasis ensures that the fitness requirement imposed by the outer cell to its inner cells holds. If there is underfitting of an inner cell, then the outer cell makes sure to complete what is missing with new resources. If there is overfitting of an inner cell, then the outer cell makes sure to take the generated surplus. Let us consider the triple of cells $(c, d_1, d_2)$ shown below, where $c$ plays the role of the outer cell and $d_1$ and $d_2$ play the role of inner cells nested in the first organelle $x_1 = (0, 1)$ and the second organelle $x_2 = (11, 13)$ of $c$, respectively.

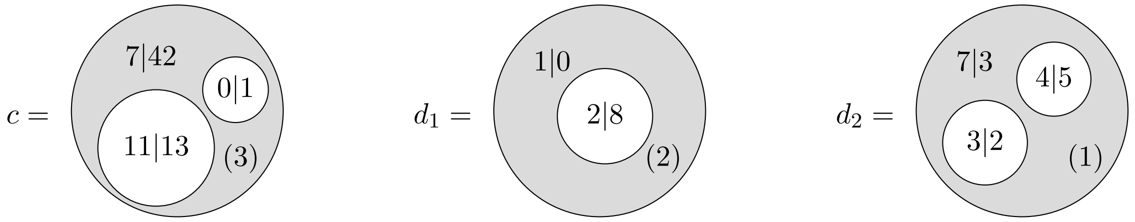


The first organelle of $c$ sets a state requirement of $(0, 1)$ to the inner cell $d_1$. Similarly, the second organelle of $c$ sets a state requirement of $(11, 13)$ to the inner cell $d_2$. The states of the inner cells $d_1$ and $d_2$ are given by their contents $K(d_1) = (3, 8)$ and $K(d_2) = (14, 10)$. Here we see that $d_1$ works too much in both components while $d_2$ works too much in the first component, but not enough in the second component.

Let us now consider the homeostasis problem that consists of the two 2-tuples $\lambda_1 = (0, 1)$ and $\lambda_2 = (11, 13)$. Solving the homeostasis problem for this transformation would amount to determining the state taken by the cell $c$ in order for the inner cell to fit the underlying composition. Intuitively, this means that information is exchanged by the two cells through the inner cell membrane. Letting the information go through the membrane is translated into the composition $c \circ d$ and closing the flux of information is translated into a factorization problem of the form $c' \circ (d'_1, d'_2) = c \circ (d_1, d_2)$. A solution of this problem is given below.

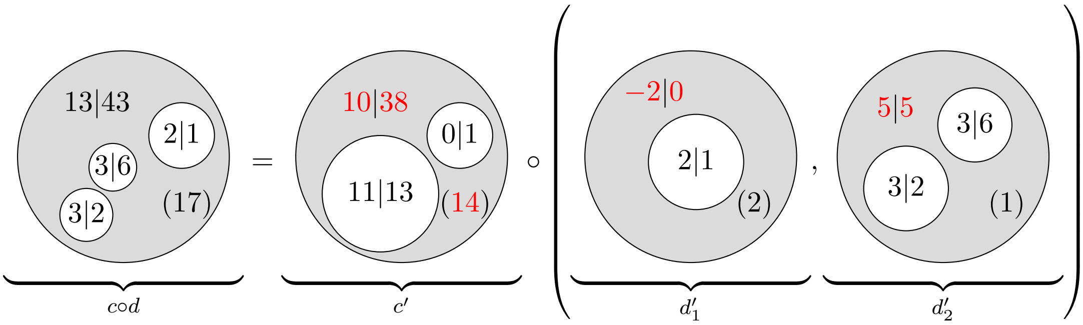


Compared to its previous state $c$, the outer cell $c'$ has taken 1 unit from its second organelle, in its first component, and has given 4 units to its second organelle, in its second component.

**Theorem 1.42** (A unique solution)**.** *The solution $(\kappa, c', d')$ of any homeostasis problem $(c, d, \lambda)$, as described in Definition 1.40, exists, is unique and is determined by the identity*

$$\kappa_i = \lambda_i - K(d_i) + \mathsf{cyt}(d_i)$$

*for every $i \in [n]$. In addition, the equations $\mathsf{res}(c') = \mathsf{res}(c)$ and $c' \circ d' = c \circ d$ hold.*

*Proof.* Let $L(i)$ denote the sum $\sum_{k=1}^{i-1} m_k$. By Definition 1.22, the condition $\delta_i(c', d'_i) = 0$ can be reformulated as follows:

$$\lambda_i = \mathsf{org}(c')_i = K(d'_i) = \kappa_i + \sum_{i=1}^{m_i} \mathsf{org}(c \circ d)_{L(i),i}$$

Since $K(d_i)$ is equal to $\mathsf{cyt}(d_i) + \sum_{i=1}^{m_i} \mathsf{org}(c \circ d)_{L(i),i}$, the previous relation can be derived into the formula given in the statement. In particular, this shows that uniqueness of the solution. To show that the solution exists, we need to check that the parameters defining $c'$ and $d'_i$ that are all well-defined. It turns out that we only need to check that the residual of $c'$ is non-negative. By Theorem 1.18, we have the following equation:

$$\mathsf{res}(c') = \mathsf{res}(c \circ d) - \sum_{i=1}^{n} \mathsf{res}(d_i) = \mathsf{res}(c) + \sum_{i=1}^{n} \mathsf{res}(d_i) - \sum_{i=1}^{n} \mathsf{res}(d_i) = \mathsf{res}(c)$$

Since the residual of $c$ is non-negative, the existence of the solution is proven. Finally, to show that the equation $c' \circ d' = c \circ d$ holds, we can use Theorem 1.18. For this, we need to make sure that the number of organelles of $c \circ d$ is $\sum_{i=1}^{n} m_i$, which directly follows from Definition 1.14. ☐

In the rest of this section, we study the variations of the algebra operator in terms of a differential over infinitesimal homeostatic exchanges (Definition 1.44). To do so, we will need Proposition 1.43 and Proposition 1.45, which will help us compute useful formulas for these variations.

**Proposition 1.43** (Properties of homeostasis solutions)**.** *Let $(\kappa, c', d')$ be the solution of a homeostasis problem $(c, d, \lambda)$ as described in Definition 1.40. The following equations hold:*

1) $K(c' \circ d') = K(c \circ d)$;
2) $\mathsf{org}(d'_k) = \mathsf{org}(d_k)$, *for every* $k \in [n]$.
3) $K(c') = K(c' \circ d')$;
4) $K(c) = K(c \circ d) + \sum_k \delta_k(c, d_k)$;
5) $K(d'_k) = \mathsf{org}(c')_k$, *for every* $k \in [n]$;
6) $K(d_k) = \mathsf{org}(c)_k - \delta_k(c, d_k)$;

*Proof.* Item 1 comes from the second statement of Theorem 1.42. Item 2 is a direct consequence of Definition 1.19. Item 3 directly follows from the equation $\delta_k(c', d'_k) = 0$, which, by Definition 1.40, holds for every $k \in [n]$. Item 4 follows from the equation given in Remark 1.25 and the fact that $\delta_k(c', d'_k) = 0$ for every $k \in [n]$. Finally, Item 5 is shown in Remark 1.25 and item 6 directly follows from Definition 1.22. ☐

**Definition 1.44** (Infinitesimal homeostatic solutions)**.** Let $(c, d, \lambda)$ be a homeostasis problem as described in Definition 1.40. For every real number $\varepsilon$, integer $v \in [N]$ and integer $j \in [n]$, a homeostasis solution $(\kappa, c', d')$ will be said to be $(\varepsilon, j, v)$-*infinitesimal* if

- ▷ $\lambda_{j,v} = \mathsf{org}(c)_{j,v} + \varepsilon$;
- ▷ $\lambda_{k,u} = \mathsf{org}(c)_{k,u}$ for every pair $(k, u) \neq (j, v)$ such that $k \in [n]$ and $u \in [N]$;
- ▷ the identity $\mathcal{S}\delta_k(c, d_k) = 0$ holds for every $k \in [n]$.

**Proposition 1.45.** *Let $(\kappa, c', d')$ be an $(\varepsilon, j, v)$-infinitesimal solution for a homeostasis problem $(c, d, \lambda)$ as described in Definition 1.40. The following equations hold:*

1) $\mathcal{S}K(c' \circ d') - \mathcal{S}K(c \circ d) = 0$;
2) $\mathcal{S}K(c') - \mathcal{S}K(c) = 0$;
3) $\mathcal{S}K(d'_j) - \mathcal{S}K(d_j) = \varepsilon$;
4) $\mathcal{S}K(d'_k) - \mathcal{S}K(d_k) = 0$ *for every* $k \in [n]$ *that is not* $j$.

*Proof.* Item 1 follows from the first item of Proposition 1.43. To show item 2, use the third and fourth items of Proposition 1.43 to deduce the equation $\mathcal{S}K(c') - \mathcal{S}K(c) = \sum_k \mathcal{S}\delta_k(c, d_k)$, and use the last item of Definition 1.44 to conclude. To show item 3 and item 4, use the fifth and sixth items of Proposition 1.43 to deduce the equation $\mathcal{S}K(d'_j) - \mathcal{S}K(d_j) = \mathcal{S}\mathsf{org}(c')_j - \mathcal{S}\mathsf{org}(c)_j + \mathcal{S}\delta_k(c, d_k)$ and use the items of Definition 1.44 with the identity $\lambda_j = \mathsf{org}(c')_j$ (Definition 1.40) to conclude. ☐

**Proposition 1.46** (Effects)**.** *Let $(\kappa, c', d')$ be an $(\varepsilon, j, v)$-infinitesimal solution for a homeostasis problem $(c, d, \lambda)$ where $c$ and $c'$ are non-empty cells in $\mathcal{C}^N(n)$ and, for every $k \in [n]$, the $k$-th components $d_k$ and $d'_k$ of $d$ and $d'$ are non-empty cells in $\mathcal{C}^N(m_k)$. For every collection $a = (a_k)_{k\in[n]}$ of vectors $a_k \in \Delta_N(A)^{m_k}$, the following equations hold.*

$$U(c', d')(a)_u - U(c, d)(a)_u = \frac{\varepsilon}{\mathcal{S}K(c)} \times \begin{cases} (d_j \cdot a_j)_u \dfrac{\mathsf{org}(c)_j}{\mathcal{S}\mathsf{org}(c)_j + \varepsilon} & \text{if } u \neq v \\ -(d_j \cdot a_j)_v \displaystyle\sum_{u=1,\neq v}^{N} \dfrac{\mathsf{org}(c)_{j,u}}{\mathcal{S}\mathsf{org}(c)_j + \varepsilon} & \text{if } u = v \end{cases}$$

*Proof.* By Definition 1.40 and Definition 1.44, the identities $\mathcal{S}\delta_k(c', d'_k) = 0$ and $\mathcal{S}\delta_k(c, d_k) = 0$ hold for every $k \in [n]$. By Proposition 1.36, this means that we have the following formulas for every $u \in [N]$:

$$U(c', d')(a)_u = \sum_{k=1}^{n} \frac{(d'_k \cdot a_k)_u}{\mathcal{S}K(c')} \Big(\mathcal{S}\mathsf{org}(c')_k - \mathsf{org}(c')_{k,u}\Big) \qquad U(c, d)(a)_u = \sum_{k=1}^{n} \frac{(d_k \cdot a_k)_u}{\mathcal{S}K(c)} \Big(\mathcal{S}\mathsf{org}(c)_k - \mathsf{org}(c)_{k,u}\Big)$$

According to the third and fourth items of Proposition 1.45 and the second item of Proposition 1.43, the following equations hold:

$$(d'_k \cdot a_k)_u = (d_k \cdot a_k)_u \times \begin{cases} 1 & \text{if } k \neq j \\ \dfrac{\mathcal{S}K(d_j)}{\mathcal{S}K(d_j) + \varepsilon} & \text{if } k = j \end{cases}$$

According to Definition 1.44 and Definition 1.40, the following identities hold:

$$\mathsf{org}(c')_{k,u} = \mathsf{org}(c)_{k,u} + \begin{cases} 0 & \text{if } (k,u) \neq (j,v) \\ \varepsilon & \text{if } (k,u) = (j,v) \end{cases} \qquad \mathcal{S}\mathsf{org}(c')_k = \mathcal{S}\mathsf{org}(c)_k + \begin{cases} 0 & \text{if } k \neq j \\ \varepsilon & \text{if } k = j \end{cases} \tag{1.7}$$

We can use the equations and the second item of Proposition 1.45 to show that the difference $U(c', d')(a)_u - U(c, d)(a)_u$ is equal to the following expression for every $u \in [N]$:

$$\frac{(d_j \cdot a_j)_u}{\mathcal{S}K(c)} \frac{\mathcal{S}K(d_j)}{\mathcal{S}K(d_j) + \varepsilon} \Big(\mathcal{S}\mathsf{org}(c)_j + \varepsilon - \mathsf{org}(c')_{j,u}\Big) - \frac{(d_j \cdot a_j)_u}{\mathcal{S}K(c)} \Big(\mathcal{S}\mathsf{org}(c)_j - \mathsf{org}(c)_{j,u}\Big)$$

Let us factorize the previous expression by the inverse of $\mathcal{S}K(d_j) + \varepsilon$ and let us replace the quantity $\mathcal{S}K(d_j)$ with $\mathcal{S}\mathsf{org}(c)_j$ since $\mathcal{S}\delta_j(c, d_j) = \mathcal{S}K(d_j) - \mathcal{S}\mathsf{org}(c)_j = 0$ to obtain the following expression:

$$\frac{1}{\mathcal{S}K(c)} \frac{(d_j \cdot a_j)_u}{\mathcal{S}\mathsf{org}(c)_j + \varepsilon} \Big(\mathcal{S}\mathsf{org}(c)_j(\mathcal{S}\mathsf{org}(c)_j + \varepsilon - \mathsf{org}(c')_{j,u}) - (\mathcal{S}\mathsf{org}(c)_j + \varepsilon)(\mathcal{S}\mathsf{org}(c)_j - \mathsf{org}(c)_{j,u})\Big)$$

We can simplify the right-hand side bracket of the previous expression as follows:

$$\frac{1}{\mathcal{S}K(c)} \frac{(d_j \cdot a_j)_u}{\mathcal{S}\mathsf{org}(c)_j + \varepsilon} \Big(\mathcal{S}\mathsf{org}(c)_j(\mathsf{org}(c)_{j,u} - \mathsf{org}(c')_{j,u}) + \varepsilon\mathsf{org}(c)_{j,u}\Big)$$

Using the left-hand side equations of (1.7) and the fact that $\mathcal{S}\mathsf{org}(c)_j$ is the sum $\sum_{u=1}^{N} \mathsf{org}(c)_{j,u}$, we can show that the previous expression is equal to the one given in the statement. This finishes the proof. □

**Definition 1.47** (Squared algebra operator)**.** Let $c$ and $d_1, \dots, d_n$ be non-empty cells of dimension $N$. For every collection $a = (a_k)_{k\in[n]}$ of vectors $a_k \in \Delta_N(A)^{m_k}$, we will denote by $U^2(c, d)(a)$ the scalar product of the vector $U(c, d)(a)$ with itself.

**Definition 1.48** (Allostasis)**.** Let $(\kappa, c', d')$ be an $(\varepsilon, j, v)$-infinitesimal solution for a homeostasis problem $(c, d, \lambda)$ where $c$ and $c'$ are non-empty cells in $\mathcal{C}^N(n)$ and, for every $k \in [n]$, the $k$-th components $d_k$ and $d'_k$ of $d$ and $d'$ are non-empty cells in $\mathcal{C}^N(m_k)$. For every collection $a = (a_k)_{k\in[n]}$ of vectors $a_k \in \Delta_N(A)^{m_k}$, we will denote:

$$\frac{\partial_{j,v} U^2(c, d)}{\partial(c, d)}(a) := \lim_{\varepsilon \to 0} \frac{U^2(c', d')(a) - U^2(c, d)(a)}{2\varepsilon}.$$

We will call the previous quantity the *allostatic differential* at $(j,v)$.

**Proposition 1.49** (Formula for allostatic differentials)**.** *Let $(\kappa,c',d')$ be an $(\varepsilon,j,v)$-infinitesimal solution for a homeostasis problem $(c,d,\lambda)$ where $c$ and $c'$ are non-empty cells in $\mathcal{C}^N(n)$ and, for every $k\in[n]$, the $k$-th components $d_k$ and $d'_k$ of $d$ and $d'$ are non-empty cells in $\mathcal{C}^N(m_k)$. For every collection $a=(a_k)_{k\in[n]}$ of vectors $a_k\in\Delta_N(A)^{m_k}$, the following equation holds.*

$$\frac{\partial_{j,v}U^2(c,d)}{\partial(c,d)}(a)=\frac{-1}{\mathcal{S}K(c)}\sum_{u=1,\neq v}^{N}\frac{\mathsf{org}(c)_{j,u}}{\mathcal{S}\mathsf{org}(c)_j}\Big((d_j\cdot a_j)_v\times U(c,d)(a)_v-(d_j\cdot a_j)_u\times U(c,d)(a)_u\Big)$$

*Proof.* By Definition 1.48 and Definition 1.47, the allostatic differential of the statement is equal to the limit

$$\frac{\partial_{j,v}U^2(c,d)}{\partial(c,d)}(a)=\lim_{\varepsilon\to 0}\sum_{u=1}^{N}\frac{U(c',d')(a)_u^2-U(c,d)(a)_u^2}{2\varepsilon}. \tag{1.8}$$

It is easy to check that the following ientity holds:

$$\frac{U(c',d')(a)_u^2-U(c,d)(a)_u^2}{2\varepsilon}=U(c,d)(a)_u\frac{U(c',d')(a)_u-U(c,d)(a)_u}{\varepsilon}+\frac{(U(c',d')(a)_u-U(c,d)(a)_u)^2}{2\varepsilon}$$

By Proposition 1.36, we can see that the difference $U(c',d')(a)_u-U(c,d)(a)_u$ takes the form of a multiplication $\varepsilon f(\varepsilon)$ where the limit of $f(\varepsilon)$ exists when $\varepsilon\to 0$. In particular, this gives us the following equation:

$$\lim_{\varepsilon\to 0}\frac{U(c',d')(a)_u^2-U(c,d)(a)_u^2}{2\varepsilon}=U(c,d)(a)_u\lim_{\varepsilon\to 0}f(\varepsilon)$$

It follows from the expression of $f(\varepsilon)$ given in Proposition 1.36 that the following identities hold:

$$\lim_{\varepsilon\to 0}\frac{U(c',d')(a)_u^2-U(c,d)(a)_u^2}{2\varepsilon}=U(c,d)(a)_u\frac{1}{\mathcal{S}K(c)}\times\begin{cases}(d_j\cdot a_j)_u\dfrac{\mathsf{org}(c)_j}{\mathcal{S}\mathsf{org}(c)_j} & \text{if } u\neq v\\ -(d_j\cdot a_j)_v\displaystyle\sum_{u=1,\neq v}^{N}\frac{\mathsf{org}(c)_{j,u}}{\mathcal{S}\mathsf{org}(c)_j} & \text{if } u=v\end{cases} \tag{1.9}$$

Since equation (1.8) is a limit of a finite sum whose terms admit limits, the allostatic differential of (1.8) is equal to the sum of these limits, all described in (1.9). Computing this sum gives us the equation of the statement. ☐

1.7. **Cell fission and cell fusion.** The goal of this section is to show that we can regulate the variations of the algebra operator though composition and factorization of cells (see Theorem 1.58).

**Definition 1.50** (Tensor operator)**.** For every positive integer $n$ greater than 2 and collection $\lambda=(\lambda_i)_{i\in[n]}$ of vectors in $\mathbb{R}_+^N$, we will denote the cell $(n,0,0,\lambda)$ as $\xi_n^N(\lambda)$. The map $\lambda\mapsto\xi_n^N(\lambda)$ will be called the *$n$-tensor operator.*

**Convention 1.51** (Tensors)**.** Let $n$ be a positive integer and $d=(d_1,\dots,d_n)$ be a $n$-tuple of cells $d_k$ in $\mathcal{C}^N(m_k)$. By Theorem 1.18, for every collection $\lambda=(\lambda_i)_{i\in[n]}$ of vectors in $\mathbb{R}_+^N$, the composition $\xi_n(\lambda)\circ d$, living in $\mathcal{C}^N(m_1+\dots+m_n)$, does not depend on $\lambda$ and yields a unique cell whose collection of organelles is the concatenation of the organelles of the cells $d_1,\dots,d_n$, as shown below.

$$\xi_n(\lambda)\circ d=\left(\sum_{k=1}^{n}m_k,\sum_{k=1}^{n}\mathsf{res}(d_k),\sum_{k=1}^{n}\mathsf{cyt}(d_k),\Big(\mathsf{org}(d_1),\mathsf{org}(d_2),\dots,\mathsf{org}(d_n)\Big)\right) \tag{1.10}$$

We will denote this cell as $d_1\otimes d_2\otimes\dots\otimes d_n$ and call it the *$n$-tensor of $d$.*

**Example 1.52** (Using tensors to model fission and fusion of cells)**.** A tensor of cells can be seen as a fusion of the two cells, were the cytosol are added together but the organelles are put next to each other. Alternatively, expressing a given cell as a tensor is a way to model cell division, were the residual and the organelles of the cell are split between the two child cells.

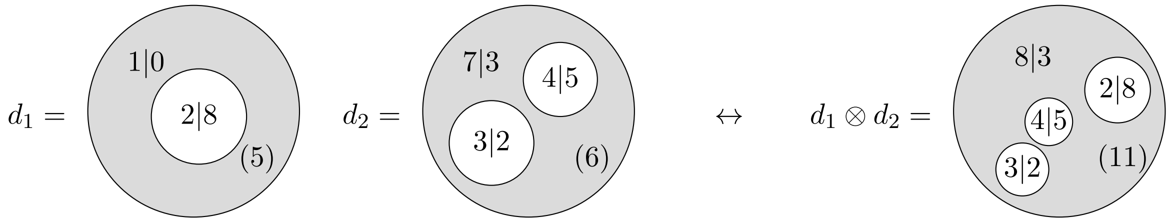


In the remainder of this section, we use tensors to study how cell division and fusion can decrease the values of the algebra operator.

**Example 1.53** (Fission and fusion via compartment formation)**.** The point of defining cell fission and cell fusion by using a tensor operator (Definition 1.50) is that it emphasizes compartment formation, where the created compartment is the tensor operator cell $\xi_n^N(\lambda)$. Of course, as seen in Example 1.52, we cannot explicitly see the formation of this compartment, because this cells is directly composed with other cells. Nevertheless, we can visualize the compartment formation if we make the factorization and composition operations defining (1.10) more explicit (through a two-step operation). More specifically, a fusion can be represented by

- an outer factorization (in the environment) to create the future merged cell;
- an inner composition (in the appeared compartment) to pop-up the inner membranes;

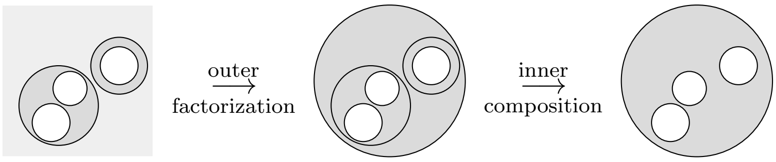


Similarly, a fission can be represented by

- an inner factorization (in the cell) to create the future divided cells;
- an outer composition (in the environment) to pop-up the old membrane;

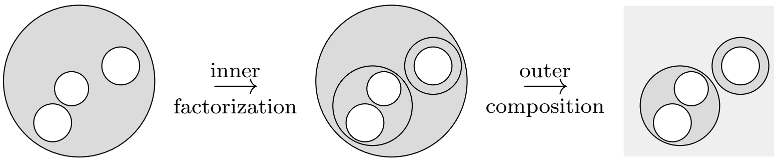


These types of decomposition will be used in section 2.1 to trigger compartment formation while minimizing the values of algebra operator via fission and fusion of cells.

**Proposition 1.54** (Content and tensors)**.** *Let* $d_1 \in \mathcal{C}^N(m_1)$ *and* $d_2 \in \mathcal{C}^N(m_2)$ *be two cells. The identity* $K(d_1 \otimes d_2) = K(d_1) + K(d_2)$ *holds.*

*Proof.* We can deduce the following identities from formula (1.10) given in Convention 1.51.

$$K(d_1 \otimes d_2) = \mathsf{cyt}(d_1 \otimes d_2) + \sum_{i=1}^{m_1+m_2} \mathsf{org}(d_1 \otimes d_2)_i = \mathsf{cy}(d_1) + \mathsf{cyt}(d_2) + \sum_{i=1}^{m_1} \mathsf{org}(d_1)_i + \sum_{i=1}^{m_2} \mathsf{org}(d_2)_i$$

We can then rearrange the terms of the right-hand side to express the sum $K(d_1) + K(d_2)$. □

**Proposition 1.55** (Action and tensors). *Let $d_1 \in \mathcal{C}^N(m_1)$ and $d_2 \in \mathcal{C}^N(m_2)$ be two cells such that their 2-tensor $d_1 \otimes d_2$ is a non-empty cell. For every pair of vectors $a_1 \in \Delta_N(A)^{m_1}$ and $a_2 \in \Delta_N(A)^{m_2}$, the following identity holds for every $u \in [N]$.*

$$((d_1 \otimes d_2) \cdot (a_1 \odot a_2))_u = (d_1 \cdot a_1)_u \frac{\mathcal{S}K(d_1)}{\mathcal{S}K(d_1 \otimes d_2)} + (d_2 \cdot a_2)_u \frac{\mathcal{S}K(d_2)}{\mathcal{S}K(d_1 \otimes d_2)}$$

*Proof.* By Definition 1.30, we have the equation

$$((d_1 \otimes d_2) \cdot (a_1 \odot a_2))_u = \sum_{i=1}^{m_1+m_2} (a_1 \odot a_2)_{i,u} \frac{\mathsf{org}(d_1 \otimes d_2)_{i,u}}{\mathcal{S}K(d_1 \otimes d_2)}$$

Because the vector $a_1 \odot a_2$ is the concatenation of $a_1$ and $a_2$ and the vector $\mathsf{org}(d_1 \otimes d_2)$ is the concatenation of $\mathsf{org}(d_1)$ and $\mathsf{org}(d_2)$, we can rewrite the previous expression as follows.

$$((d_1 \otimes d_2) \cdot (a_1 \odot a_2))_u = \sum_{i=1}^{m_1} (a_1)_{i,u} \frac{\mathsf{org}(d_1)_{i,u}}{\mathcal{S}K(d_1 \otimes d_2)} + \sum_{i=1}^{m_2} (a_2)_{i,u} \frac{\mathsf{org}(d_2)_{i,u}}{\mathcal{S}K(d_1 \otimes d_2)}$$

Factorizing the two sums, on the right-hand side, by $\mathcal{S}K(d_1)/\mathcal{S}K(d_1 \otimes d_2)$ and $\mathcal{S}K(d_2)/\mathcal{S}K(d_1 \otimes d_2)$, we obtain the following expression.

$$((d_1 \otimes d_2) \cdot (a_1 \odot a_2))_u = \frac{\mathcal{S}K(d_1)}{\mathcal{S}K(d_1 \otimes d_2)} \left( \sum_{i=1}^{m_1} (a_1)_{i,u} \frac{\mathsf{org}(d_1)_{i,u}}{\mathcal{S}K(d_1)} \right) + \frac{\mathcal{S}K(d_2)}{\mathcal{S}K(d_1 \otimes d_2)} \left( \sum_{i=1}^{m_2} (a_2)_{i,u} \frac{\mathsf{org}(d_2)_{i,u}}{\mathcal{S}K(d_2)} \right)$$

Finally, by Definition 1.30, this last expression is equivalent to the one given in the statement. □

*Remark* 1.56 (Barycenter). Proposition 1.55 and Proposition 1.54 say that the action of a tensor $(d_1 \otimes d_2)$ on a concatenation $(a_1 \odot a_2)$ is the barycenter of the actions $d_1 \cdot a_1$ and $d_1 \cdot a_2$ where the weights are given by the contents of $d_1$ and $d_2$. More generally, when fitness is assumed and the values of the cytosol are ignored, the action of a cell can be seen as a barycenter of the actions of its organelles. Interestingly, the barycenter of a set of real values can be used as a 'pivot' to divide the space of these real values into two parts: the sets of real values that are below it and those that are above it. In section 2.5, we will use this criterion to model mergings and divisions of cells: specifically, we will look at the barycenter of the organelles of a cell and gather together those organelles that tend to be grouped in the same parts (see Definition 2.30).

**Proposition 1.57** (Fitness and tensor cells). *Let $c \in \mathcal{C}^N(n+1)$ and $c' \in \mathcal{C}^N(n)$ be two cells such that there exits two cells $d_1$ and $d_2$ of dimension $N$ for which the equation $c = c' \circ_n \xi_2(K(d_1), K(d_2))$ holds. In this case, the following identity holds:*

$$K(c) - K(c') = \delta_n(c', d_1 \otimes d_2)$$

*Proof.* First, we apply Proposition 1.24 to the composition $c = c' \circ_n \xi_2(K(d_1), K(d_2))$ to deduce the expression of $K(c')$ in terms of $K(c)$: this gives us the equation $K(c) - K(c') = \mathsf{org}(c')_n - K(d_1) - K(d_2)$. Then, Proposition 1.54 allows us to turn this equation into the equation $K(c) - K(c') = \mathsf{org}(c')_n - K(d_1 \otimes d_2)$. Now, since Definition 1.22 gives us the identity $\delta_n(c', d_2 \otimes d_2) = \mathsf{org}(c')_n - K(d_1 \otimes d_2)$, the statement follows. □

The following theorem assesses by how much the values of the algebra operator are reduced and enhanced when compartment fission or fusion happens in a cell. Since the algebra operator is invariant by symmetry (see Remark 1.35), we will consider only one index configuration for the fusion (or fission) of the organelles, namely only the last two organelles $d_n$ and $d_{n+1}$ will be tensored. See Remark 1.59 for further discussion.

**Theorem 1.58** (Compartment theorem). *Let $c \in \mathcal{C}^N(n+1)$ and $c' \in \mathcal{C}^N(n)$ be two non-empty cells, let $d = (d_1, \dots, d_{n+1})$ be a $(n+1)$-tuples of non-empty cells where, for every $k \in [n+1]$, the cells $d_k$ is in $\mathcal{C}^N(m_k)$ and let $a = (a_k)_{k \in [n+1]}$ be a collection of vectors $a_k \in \Delta_N(A)^{m_k}$. Let us denote*

$$d' = (d_1, \dots, d_{n-1}, d_n \otimes d_{n+1}) \qquad \text{and} \qquad a' = (a_1, \dots, a_{n-1}, a_n \odot a_{n+1}).$$

*If the equation $c = c' \circ_n \xi_2(K(d_n), K(d_{n+1}))$ holds and if we assume that $\mathcal{S}\delta_k(c, d_k) = 0$ for every $k \in [n+1]$ and $\mathcal{S}\delta_k(c', d'_k) = 0$ for every $k \in [n]$, then the following identity holds for every $u \in [N]$.*

$$U(c', d')(a')_u - U(c, d)(a)_u = \Big((d_n \cdot a_n)_u - (d_{n+1} \cdot a_{n+1})_u\Big)\left(\frac{\mathsf{org}(c)_{n,u}}{\mathcal{S}\mathsf{org}(c)_n} - \frac{\mathsf{org}(c)_{n+1,u}}{\mathcal{S}\mathsf{org}(c)_{n+1}}\right)\frac{\mathcal{S}\mathsf{org}(c)_n \mathcal{S}\mathsf{org}(c)_{n+1}}{\mathcal{S}K(c)\mathcal{S}\mathsf{org}(c')_n}$$

*Proof.* According to Proposition 1.36, since $\mathcal{S}\delta_k(c, d_k) = 0$ for every $k \in [n+1]$ and $\mathcal{S}\delta_k(c', d'_k) = 0$ for every $k \in [n]$, we have the following expressions:

$$U(c, d)(a)_u = \sum_{k=1}^{n} \frac{(d_k \cdot a_k)_u}{\mathcal{S}K(c)}\Big(\mathcal{S}\mathsf{org}(c)_k - \mathsf{org}(c)_{k,u}\Big) \quad U(c', d')(a')_u = \sum_{k=1}^{n} \frac{(d'_k \cdot a'_k)_u}{\mathcal{S}K(c')}\Big(\mathcal{S}\mathsf{org}(c')_k - \mathsf{org}(c')_{k,u}\Big)$$

Before evaluating the difference between the previous terms, let us make some observations. First, Proposition 1.57 gives us the equation $K(c) - K(c') = \delta_n(c', d_n \otimes d_{n+1})$. We can couple this equation with the fact that the summed fitness $\mathcal{S}\delta_n(c', d_n \otimes d_{n+1})$ is zero to deduce that the equation $\mathcal{S}K(c) = \mathcal{S}K(c')$ holds.

Second, it follows from Definition 1.19 and the equation $c = c' \circ_n \xi_2(K(d_n), K(d_{n+1}))$ that the identity $\mathsf{org}(c')_k = \mathsf{org}(c)_k$ holds for every $k \in [n-1]$. As a result, the difference $U(c', d')(a')_u - U(c, d)(a)_u$ is equal to the following expression:

$$\begin{aligned} U(c', d')(a')_u - U(c, d)(a)_u = \frac{(d'_n \cdot a'_n)_u}{\mathcal{S}K(c)}\Big(\mathcal{S}\mathsf{org}(c')_n - \mathsf{org}(c')_{n,u}\Big) \\ -\frac{(d_n \cdot a_n)_u}{\mathcal{S}K(c)}\Big(\mathcal{S}\mathsf{org}(c)_n - \mathsf{org}(c)_{n,u}\Big) - \frac{(d_{n+1} \cdot a_{n+1})_u}{\mathcal{S}K(c)}\Big(\mathcal{S}\mathsf{org}(c)_{n+1} - \mathsf{org}(c)_{n+1,u}\Big) \end{aligned} \tag{1.11}$$

We now want to factorize expression (1.11). First, according to Proposition 1.54, the identity $\mathcal{S}K(d'_n) = \mathcal{S}K(d_n) + \mathcal{S}K(d_{n+1})$ holds. Since the quantities $\mathcal{S}\delta_n(c, d_n)$, $\mathcal{S}\delta_{n+1}(c, d_{n+1})$ and $\mathcal{S}\delta_n(c', d'_n)$ are all zeros, Definition 1.22 implies that the previous identity is equivalent to the identity

$$\mathcal{S}\mathsf{org}(c')_n = \mathcal{S}\mathsf{org}(c)_n + \mathcal{S}\mathsf{org}(c)_{n+1}. \tag{1.12}$$

We can use equation (1.12) in each of the brackets of (1.11) as well as the identity of Proposition 1.55 in the topmost summand of (1.11), where $d'_n = d_n \otimes d_{n+1}$ and $a'_n = a_n \odot a_{n+1}$, to obtain the following new expression:

$$\begin{aligned} U(c', d')(a')_u - U(c, d)(a)_u = \frac{(d_n \cdot a_n)_u}{\mathcal{S}K(c)}\left(\frac{\mathcal{S}K(d_n) - \mathcal{S}K(d'_n)}{\mathcal{S}K(d'_n)}\right)\Big(\mathcal{S}\mathsf{org}(c)_n - \mathsf{org}(c)_{n,u}\Big) \\ +\frac{(d_{n+1} \cdot a_{n+1})_u}{\mathcal{S}K(c)}\left(\frac{\mathcal{S}K(d_{n+1}) - \mathcal{S}K(d'_n)}{\mathcal{S}K(d'_n)}\right)\Big(\mathcal{S}\mathsf{org}(c)_{n+1} - \mathsf{org}(c)_{n+1,u}\Big) \\ +\frac{(d_n \cdot a_n)_u}{\mathcal{S}K(c)}\frac{\mathcal{S}K(d_n)}{\mathcal{S}K(d'_n)}\Big(\mathcal{S}\mathsf{org}(c)_{n+1} - \mathsf{org}(c)_{n+1,u}\Big) + \frac{(d_{n+1} \cdot a_{n+1})_u}{\mathcal{S}K(c)}\frac{\mathcal{S}K(d_{n+1})}{\mathcal{S}K(d'_n)}\Big(\mathcal{S}\mathsf{org}(c)_n - \mathsf{org}(c)_{n,u}\Big) \end{aligned} \tag{1.13}$$

Again, we can use the fact that $\mathcal{S}\delta_{n+1}(c, d_{n+1})$, $\mathcal{S}\delta_n(c, d_n)$ and $\mathcal{S}\delta_n(c', d'_n)$ are zeros to further simplify the dividends of expression (1.13). Specifically, we use the identities $\mathcal{S}\mathsf{org}(c)_{n+1} = \mathcal{S}K(d_{n+1})$, $\mathcal{S}\mathsf{org}(c)_n = \mathcal{S}K(d_n)$ and $\mathcal{S}\mathsf{org}(c')_n = \mathcal{S}K(d'_n)$ as well as equation (1.12) to turn each of the dividend of (1.13) whose expressions use terms of the form $\mathcal{S}K(\_)$ into expressions using terms of the form $\mathcal{S}\mathsf{org}(\_)_\_$, as shown below.

$$\begin{aligned} -\frac{(d_n \cdot a_n)_u}{\mathcal{S}K(c)}\left(\frac{\mathcal{S}\mathsf{org}(c)_{n+1}}{\mathcal{S}\mathsf{org}(c')_n}\right)\Big(\mathcal{S}\mathsf{org}(c)_n - \mathsf{org}(c)_{n,u}\Big) - \frac{(d_{n+1} \cdot a_{n+1})_u}{\mathcal{S}K(c)}\left(\frac{\mathcal{S}\mathsf{org}(c)_n}{\mathcal{S}\mathsf{org}(c')_n}\right)\Big(\mathcal{S}\mathsf{org}(c)_{n+1} - \mathsf{org}(c)_{n+1,u}\Big) \\ +\frac{(d_n \cdot a_n)_u}{\mathcal{S}K(c)}\frac{\mathcal{S}\mathsf{org}(c)_n}{\mathcal{S}\mathsf{org}(c')_n}\Big(\mathcal{S}\mathsf{org}(c)_{n+1} - \mathsf{org}(c)_{n+1,u}\Big) + \frac{(d_{n+1} \cdot a_{n+1})_u}{\mathcal{S}K(c)}\frac{\mathcal{S}\mathsf{org}(c)_{n+1}}{\mathcal{S}\mathsf{org}(c')_n}\Big(\mathcal{S}\mathsf{org}(c)_n - \mathsf{org}(c)_{n,u}\Big) \end{aligned}$$

Observe that the first and third summands as well as the second and fourth summands of the previous expression possess common factors, namely $(d_n \cdot a_n)_u/\mathcal{S}K(c)$ or $(d_{n+1} \cdot a_{n+1})_u/\mathcal{S}K(c)$. We can factorize these summands with respect to their common factors and obtain the following expression after minor simplifications.

$$\frac{(d_n \cdot a_n)_u}{\mathcal{S}K(c)} \frac{\mathcal{S}\mathsf{org}(c)_n \mathcal{S}\mathsf{org}(c)_{n+1}}{\mathcal{S}\mathsf{org}(c')_n} \left( \frac{\mathsf{org}(c)_{n,u}}{\mathcal{S}\mathsf{org}(c)_n} - \frac{\mathsf{org}(c)_{n+1,u}}{\mathcal{S}\mathsf{org}(c)_{n+1}} \right)$$
$$+ \frac{(d_{n+1} \cdot a_{n+1})_u}{\mathcal{S}K(c)} \frac{\mathcal{S}\mathsf{org}(c)_n \mathcal{S}\mathsf{org}(c)_{n+1}}{\mathcal{S}\mathsf{org}(c')_n} \left( \frac{\mathsf{org}(c)_{n+1,u}}{\mathcal{S}\mathsf{org}(c)_{n+1}} - \frac{\mathsf{org}(c)_{n,u}}{\mathcal{S}\mathsf{org}(c)_n} \right)$$

Finally, factorizing the previous expression by the obvious common factor gives the following expression:

$$U(c',d')(a')_u - U(c,d)(a)_u = \Big( (d_n \cdot a_n)_u - (d_{n+1} \cdot a_{n+1})_u \Big) \left( \frac{\mathsf{org}(c)_{n,u}}{\mathcal{S}\mathsf{org}(c)_n} - \frac{\mathsf{org}(c)_{n+1,u}}{\mathcal{S}\mathsf{org}(c)_{n+1}} \right) \frac{\mathcal{S}\mathsf{org}(c)_n \mathcal{S}\mathsf{org}(c)_{n+1}}{\mathcal{S}K(c)\mathcal{S}\mathsf{org}(c')_n}$$

This finishes the proof. □

*Remark* 1.59 (Increasing specialization through compartment flexibility). Theorem 1.58 assesses the variations of the algebra operator when an organelle of a cell is divided or, conversely, when two organelles are merged. More specifically, the formula of Theorem 1.58 implies that the values of the algebra operator are reduced at a given dimension $u$ through compartment division, meaning that the inequality

$$U(c',d')(a')_u - U(c,d)(a)_u > 0$$

holds, if the division of the cell $d'_n = d_n \otimes d_{n+1}$ into the two cells $d_n$ and $d_{n+1}$ is such that either condition (1.14) or condition (1.15) is satisfied:

$$(1.14) \qquad (d_n \cdot a_n)_u > (d_{n+1} \cdot a_{n+1})_u \qquad \frac{\mathsf{org}(c)_{n,u}}{\mathcal{S}\mathsf{org}(c)_n} > \frac{\mathsf{org}(c)_{n+1,u}}{\mathcal{S}\mathsf{org}(c)_{n+1}}$$

$$(1.15) \qquad (d_n \cdot a_n)_u < (d_{n+1} \cdot a_{n+1})_u \qquad \frac{\mathsf{org}(c)_{n,u}}{\mathcal{S}\mathsf{org}(c)_n} < \frac{\mathsf{org}(c)_{n+1,u}}{\mathcal{S}\mathsf{org}(c)_{n+1}}$$

In other words, the variation between the signals sent by $d_n$ and $d_{n+1}$ to $c$ at dimension $u$ agrees with the variation between the proportions of the $u$-th component of the content between $d_{n+1}$ and $d_n$. Similarly, the values of the algebra operator is reduced at a given dimension $u$ through compartment merging, meaning that the inequality

$$U(c',d')(a')_u - U(c,d)(a)_u < 0$$

holds, if the merging of the two cells $d_n$ and $d_{n+1}$ into the cell $d'_n = d_n \otimes d_{n+1}$ is such that either condition (1.16) or condition (1.17) is satisfied:

$$(1.16) \qquad (d_n \cdot a_n)_u > (d_{n+1} \cdot a_{n+1})_u \qquad \frac{\mathsf{org}(c)_{n,u}}{\mathcal{S}\mathsf{org}(c)_n} < \frac{\mathsf{org}(c)_{n+1,u}}{\mathcal{S}\mathsf{org}(c)_{n+1}}$$

$$(1.17) \qquad (d_n \cdot a_n)_u < (d_{n+1} \cdot a_{n+1})_u \qquad \frac{\mathsf{org}(c)_{n,u}}{\mathcal{S}\mathsf{org}(c)_n} > \frac{\mathsf{org}(c)_{n+1,u}}{\mathcal{S}\mathsf{org}(c)_{n+1}}$$

In other words, the variation between the signals sent by $d_n$ and $d_{n+1}$ to $c$ at dimension $u$ is opposed to the variation beween the proportions of the $u$-th component of the content between $d_{n+1}$ and $d_n$.

#### 2. Implementation: the algorithm and why it works

The goal of the present section is to describe, from a mathematical standpoint, the learning algorithm described in the result section of the main text. To facilitate the reader's understanding, we will also use pseudo code to describe the main steps of the learning algorithm. We will refer to the documentation of the code, available at `https://github.com/remytuyeras/intcyt-library`, for further details. Note that this documentation thoroughly explains the content of the present section, from a programming standpoint, through a tutorial. Therefore, we strongly encourage the reader who desires to fully understand the results shown in the main text of this article to have the documentation at hand.

The learning algorithm that we are about to describe is a combination of four main steps, shown in red, in the following diagram.

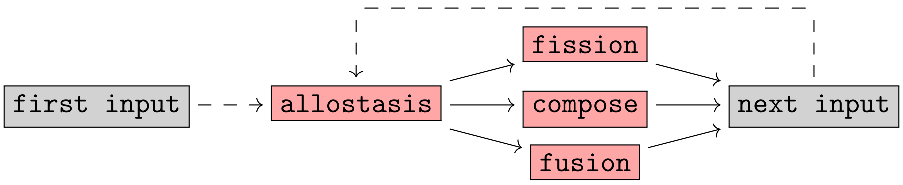


Every input given to the algorithm goes through the step `allostasis` (see diagram above), but depending on the iteration at which the algorithm is, the algorithm can go through either the step `fusion`, the step `fission` or the step `compose`. After any of these steps, the algorithm switches to the next input and restarts the same process from the step `allostasis`, as shown above.

The idea behind using the steps `fusion`, `fission` and `compose` at different iteration lies in Remark 1.53, in which we notice that fusion and fission of cells can be seen as two-step processes. Specifically, for a given *super cell* $\hat{c}$ (see Definition 2.1), the steps `fusion` and `fission` initiates fusion and fission events in $\hat{c}$, while the role of `compose` is to complete these steps. Even though there are actual formal reasons for using such a two-step process (see section 2.4), delaying the completion of fusions and fissions in $\hat{c}$ can also let the algorithm decides whether the chosen fusion or fission events are actually advantageous for $\hat{c}$. When fission and fusion events are not completed through `compose`, new compartments are created in $\hat{c}$. These new compartments can represent a new advantage to organize and classify the information learned. On the other hand, if these compartments turn out to be too disadvantageous for $\hat{c}$, the algorithm will eventually make them disappear.

2.1. **Super cells.** The following definition formalizes the concept of a tree of cells, which was used in the main text of this article, in terms of an object called a 'super cell'. In a few words, the concept of a super cell is an extension of the concept of a cell through a tree-like structure whose leaves and junctions are each associated with a cell.

**Definition 2.1** (Super cells)**.** For every every positive integer $N$ and every non-negative integers $q$ and $n$, we define the set $\mathcal{C}_q^N(n)$ by the recursive formulas given below, on the left (in which $\coprod$ and $\prod$ denote a coproduct and a product of sets, respectively):

$$\begin{cases} \mathcal{C}_0^N(n) = \mathcal{C}^N(n) \\ \mathcal{C}_{q+1}^N(n) = \coprod_{p\geq 1} \coprod_{\substack{(q_1,\dots,q_p)\,: \\ \max(q_i)_{i=1}^p = q}} \coprod_{\substack{(m_1,\dots,m_p)\,: \\ \sum_{i=1}^p m_i = n}} \left( \mathcal{C}_0^N(p) \times \prod_{i=1}^{p} \mathcal{C}_{q_i}^N(m_i) \right) \end{cases}$$

tree representation of an element in $\mathcal{C}_{q+1}^N(n)$

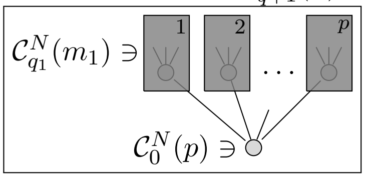


For every pair of non-negative integers $(q,n)$, an element $\hat{c} \in \mathcal{C}_q^N(n)$ is referred to as a *super cell* and said to be *of dimension* $N$, *of height* $q$ and to *possess* $n$ *organelles*. As shown above, on the right, a super cell in $\mathcal{C}_q^N(n)$ can be viewed as a tree-like structure whose leaves and junctions are labeled by cells of dimension $N$. This is further discussed in Convention 2.2.

**Convention 2.2** (Tree structure)**.** Let $\hat{c}$ be a super cell in $\mathcal{C}_q^N(n)$ for which $q \geq 1$. If we unpack the recursive structure of $\hat{c}$ specified in Definition 2.1, then we can express $\hat{c}$ in terms of a collection of tuples of cells and super cells (of lower height) as follows:

$$(2.1) \quad \begin{cases} \hat{c} = (c, (\hat{c}_1, \hat{c}_2, \dots, \hat{c}_p)) & \text{where } c \in \mathcal{C}^N(p) \text{ and } \hat{c}_i \in \mathcal{C}_{q_i}^N(m_i) \\ \hat{c}_{i_1\dots i_s} = (c_{i_1\dots i_s}, (\hat{c}_{i_1\dots i_s 1}, \dots, \hat{c}_{i_1\dots i_s p_{i_1\dots i_s}})) & \text{where } c_{i_1\dots i_s} \in \mathcal{C}^N(p_{i_1\dots i_s}) \text{ and } \hat{c}_{i_1\dots i_s i} \in \mathcal{C}_{q_i}^N(m_{i_1\dots i_s i}) \end{cases}$$

To simplify the previous relations, we can replace any long sequence of indices $i_1 \dots i_s$ with the corresponding collection $I = \{i_1, \dots, i_s\}$. Note that the collection $I$ can be empty, as in the case of the super cell $\hat{c}$ and the cell $c$ given above. Using the notation $I$, we can rewrite the relations of (2.1) more concisely, as shown below, on the left.

$$\text{(2.2)} \qquad \begin{cases} \hat{c} = (c, (\hat{c}_1, \hat{c}_2, \dots, \hat{c}_p)) \\ \hat{c}_I = (c_I, (\hat{c}_{I1}, \hat{c}_{I2}, \dots, \hat{c}_{Ip_I})) \end{cases}$$

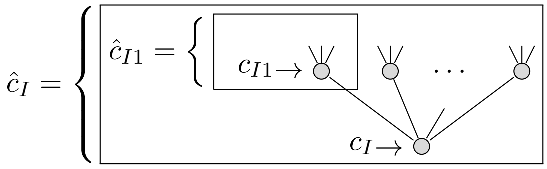


Each of the equations given in (2.2) can be visualized as a branching from a cell $c_I$ to a collection of super cell $\hat{c}_{I1}, \dots, \hat{c}_{Ip_I}$, as shown above, on the right. As a result, the total structure encoded by $\hat{c}$ can be represented as a tree of $q$ levels, where each of the equations of the form $\hat{c}_I = (c_I, (\hat{c}_{I1}, \dots, \hat{c}_{Ip_I}))$ represents a junction in the tree. On the other hand, the super cells $\hat{c}_I$ that cannot be expressed as a tuple of super cells must be of height 0 and correspond to the leaves of the tree structure associated with $\hat{c}$.

Later, given any super cell $\hat{c}$, we will refer to any associated super cell $\hat{c}_I$ of height greater than 1 as a *junction of* $\hat{c}$ and any associated super cell $\hat{c}_I$ of height 0 as a *leaf of* $\hat{c}$.

**Definition 2.3** (Indexing collections for junctions and leaves)**.** For every $\hat{c} \in \mathcal{C}_q^N(n)$, we denote by $\mathsf{Junc}(\hat{c})$ the set of collections indexing the junctions of $\hat{c}$ and by $\mathsf{Leaf}(\hat{c})$ the set of collections indexing the leaves of $\hat{c}$. More formally, these sets are defined by the following specifications:

$$\begin{cases} \mathsf{Junc}(\hat{c}) = \{I \mid \hat{c}_I \text{ is a super cell of } \hat{c} \text{ whose height is not } 0\} \\ \mathsf{Leaf}(\hat{c}) = \{I \mid \hat{c}_I \text{ is a super cell of } \hat{c} \text{ whose height is } 0\}. \end{cases}$$

We will denote by $\mathsf{Index}(\hat{c})$ the union $\mathsf{Junc}(\hat{c}) \cup \mathsf{Leaf}(\hat{c})$.

**Proposition 2.4** (Number of leaves)**.** *For every* $\hat{c} \in \mathcal{C}_q^N(n)$*, the set* $\mathsf{Leaf}(\hat{c})$ *possesses* $n$ *elements.*

*Proof.* Follows from an induction using the recursive structure given in Definition 2.1. □

**Definition 2.5** (Linear order)**.** For every super cell $\hat{c}$ in $\mathcal{C}_q^N(n)$, we define a linear order on $\mathsf{Index}(\hat{c})$ by taking the lexicographical order on integers, namely:

$$I = \{i_1, \dots, i_s\} \prec J = \{j_1, \dots, j_t\} \Leftrightarrow \exists u < \mathsf{min}(s, t) \text{ such that } i_k = j_k \text{ for every } k \in [u] \text{ and } i_{u+1} < j_{u+1}$$

**Convention 2.6** (Indexing shifts)**.** For every $\hat{c} \in \mathcal{C}_q^N(n)$ and every indexing collection $I \in \mathsf{Leaf}(\hat{c})$, we denote as $m(I)$ the number of organelles of $\hat{c}_I$. If $I \in \mathsf{Junc}(\hat{c})$, then we denote the number of organelles of $\hat{c}_I$ as $p(I)$. Finally, if we denote by $I_1 \prec \cdots \prec I_i$ the ordered sequence of elements of $\mathsf{Leaf}(\hat{c})$ that are less than $I$, then we denote by $L(I)$ the sum $m(I_1) + \cdots + m(I_i)$.

**Definition 2.7** (Organelles for a super cell)**.** For every $\hat{c} \in \mathcal{C}_q^N(n)$, any vector of the form $\mathsf{org}(\hat{c}_I)_i$, where $I \in \mathsf{Leaf}(\hat{c})$ and $i \in [m(I)]$, will be called an *organelle* of $\hat{c}$. If we denote by $I_1 \prec \cdots \prec I_n$ the ordered sequences of the elements of $\mathsf{Leaf}(\hat{c})$, then we denote by $\mathsf{org}(\hat{c})$ the concatenation $\mathsf{org}(\hat{c}_{I_1}) \odot \cdots \odot \mathsf{org}(\hat{c}_{I_n})$, which comprises all the organelles of $\hat{c}$.

**Proposition 2.8** (Retrieval)**.** *For every* $\hat{c} \in \mathcal{C}_q^N(n)$*, let* $(v_1, v_2, \dots, v_n)$ *denote the tuple* $\mathsf{org}(\hat{c})$ *of the organelles of* $\hat{c}$*. For every* $I \in \mathsf{Leaf}(\hat{c})$*, the vector* $(v_{L(I)+1}, v_{L(I)+1}, \dots, v_{L(I)+m(I)})$ *corresponds to* $\mathsf{org}(\hat{c}_I)$*.*

*Proof.* Follows from the construction of $\mathsf{org}(\hat{c})$ given in Definition 2.7 and the definition of $L(I)$ given in Convention 2.6. □

**Definition 2.9** (Pre-actions and actions)**.** Let $\hat{c}$ be a super cell in $\mathcal{C}_q^N(n)$ and let $a$ be a vector in $\Delta_N(A)^n$. For every $I \in \mathsf{Index}(\hat{c})$, we define the *pre-action* $\mathsf{preact}(\hat{c}|a)_I$ and the *action* $\mathsf{act}(\hat{c}|a)_I$ of the super cell $\hat{c}$ on the vector $a$ at the indexing collection $I$ by the following recursive formulas:

- ▷ if $I \in \mathsf{Index}(\hat{c})$, then $\mathsf{act}(\hat{c}|a)_I := c_I \cdot \mathsf{preact}(\hat{c}|a)_I$;
- ▷ if $I \in \mathsf{Leaf}(\hat{c})$, then $\mathsf{preact}(\hat{c}|a)_I = (a_{L(I)+1}, \dots, a_{L(I)+m(I)})$;

▷ if $I \in \mathsf{Junc}(\hat{c})$, then $\mathsf{preact}(\hat{c}|a)_I = \mathsf{act}(\hat{c}|a)_{I1} \odot \cdots \odot \mathsf{act}(\hat{c}|a)_{Ip(I)}$;

The leaves of any super cell are special super cells because their height is 0, meaning that they are proper cells. As a result, these super cells do not have enough structure to allow us to interpret fission and fusion events, because these events require two layers of cells, hence a super cell of height 1 (see section 1.6 and section 1.7). A solution to this problem will be to interpret each leaf of $\hat{c}$ as a super cell of height 1 by replacing their organelles with identity cells (see Definition 1.9).

**Definition 2.10** (Direct children and basal pruning)**.** For every $\hat{c} \in \mathcal{C}_q^N(n)$, we define the *direct children* of $\hat{c}$ as the following tuple of cells:

$$d(\hat{c}) = \begin{cases} (\mathsf{id}_{\mathsf{org}(c)_1}, \dots, \mathsf{id}_{\mathsf{org}(c)_n}) & \text{If } \hat{c} \text{ is a leaf, meaning that } \hat{c} = c \\ (c_1, \dots, c_p) & \text{If } \hat{c} \text{ is a junction, meaning that } \hat{c} = (c, (\hat{c}_1, \dots, \hat{c}_p)) \end{cases}$$

We define the *basal pruning* of $\hat{c}$ as the super cell $\mathsf{base}(\hat{c}) := (c, d(\hat{c}))$ where $c$ is the obvious root cell associated with $\hat{c}$.

**Definition 2.11** (Homeostatic state)**.** A super cell $\hat{c}$ in $\mathcal{C}_q^N(n)$ will be said to be *in a homeostatic state* (or to have *reached homeostasis*) if for every $I \in \mathsf{Junc}(\hat{c})$ such that $\hat{c}_I = (c_I, (\hat{c}_{I1}, \dots, \hat{c}_{Ip(I)}))$, each cell of the form $c_{Ii}$ (associated with the super cell $\hat{c}_{Ii}$) fits the $i$-th organelle of the cell $c_I$ (see Convention 1.23). In other words, the equation $\delta_i(c_I, c_{Ii}) = 0$ holds for every $I \in \mathsf{Junc}(\hat{c})$ and every index $i \in [p(I)]$.

Motivated by Convention 2.10, we diversify the pre-action of a super cell (Definition 2.9) in two ways, namely as an 'expected signal' and a 'specialized signal' (see Definition 2.12). In the spirit of the discussion following Definition 1.30, we think of the former as an expected value of the input, while we think of the latter as a measurement of how well the input matches each of the organelles of the super cell.

**Definition 2.12** (Expected and specialized signal)**.** Let $\hat{c}$ be a super cell in $\mathcal{C}_q^N(n)$ and let $a$ be a vector in $\Delta_N(A)$. For every $I \in \mathsf{Index}(\hat{c})$, we define

▷ the *expected signal* $\mathsf{esgn}(a|\hat{c})_I$ of the vector $a$ through $\hat{c}$ at the indexing collection $I$ as the pre-action $\mathsf{preact}(a'|\hat{c})_I$ on the concatenation $a' = a \odot a \odot \cdots \odot a$ of $n$ copies of $a$.

▷ the *specialized signal* $\mathsf{ssgn}(a|\hat{c})_I$ of the vector $a$ through $\hat{c}$ at the indexing collection $I$ as the pre-action $\mathsf{preact}(a'|\hat{c})_I$ on the concatenation $a' = (\mathsf{id}_{\mathsf{org}(\hat{c})_1} \cdot a) \odot (\mathsf{id}_{\mathsf{org}(\hat{c})_2} \cdot a) \odot \cdots \odot (\mathsf{id}_{\mathsf{org}(\hat{c})_n} \cdot a)$.

2.2. **Spontaneous reactions and homeostatic state.** Before describing the three steps of our algorithm, we must describe an intermediate step that is essential to the three steps. This intermediate step consists in making sure that a given super cell has reached homeostasis (Definition 2.11). This is done by going through successive steps of normalization called 'cleanings'. These cleaning procedures essentially implements the 'overall chemical reaction' described in Remark 1.32.

**Definition 2.13** (Cleaning)**.** For every cell $c \in \mathcal{C}^N(n)$, we call the *cleaning of* $c$ the cell $\mathsf{clean}(c)$ of $\mathcal{C}^N(n)$ determined by the following equations:

1) *Energy storage:* $\mathsf{res}(\mathsf{clean}(c)) = \mathsf{res}(c) + \mathcal{S}\mathsf{cyt}(c)$
2) *Disorder cleaning:* $\mathsf{cyt}(\mathsf{clean}(c)) = (0, 0, \dots, 0)$
3) *Compartmentalization:* $\mathsf{org}(\mathsf{clean}(c)) = \mathsf{org}(c)$

If we interpret the cleaning of a cell as the result of an 'overall chemical reaction', then we can interpret the three items of Definition 2.13 as follows: item 1 states that the cytosolic content is converted into energetic content, which is to be used in later similar chemical reactions; item 2 states that the cytosolic content is cleaned from its components, which have been turned into energetic content; and item 3 states that the organelles of the cells are untouched, mainly because they are separated from the cytosol by membranes.

By Definition 1.2, the cleaning of a cell is defined if, and only if its residual is non-negative. This means that a cell admits a cleaning if, and only if, it is well-defined according to the following definition.

**Definition 2.14** (Well-definedness)**.** A cell $c \in \mathcal{C}^N(n)$ will be said to be *well-defined* if the inequality $\mathsf{res}(c) + \mathcal{S}\mathsf{cyt}(c) \geq 0$ is satisfied.

The previous notion of well-definedness can be extended to super cells as follows.

**Definition 2.15** (Well-definedness)**.** A super cell $c \in \mathcal{C}_q^N(n)$ will be said to be *well-defined* if it satisfies the following inductive conditions:

1) If $\hat{c}$ is of height 0, then $\hat{c}$ is well-defined as a cell (Definition 2.14);
2) If $\hat{c} = (c, (\hat{c}_1, \dots, \hat{c}_p))$ is of height $q > 0$, then $c$ is well-defined as a cell and each super cell $\hat{c}_i$ is a well-defined super cell of height $q-1$ by induction.

We also define the cleaning of a super cell by induction. In our implementation, the cleaning of a super cell is implemented through the method `spontaneous_reaction` (see the documentation).

**Definition 2.16** (Cleaning)**.** For every well-defined super cell $\hat{c} \in \mathcal{C}_q^N(n)$, we define the *cleaning of* $\hat{c}$ as the super cell $\hat{e}$ of $\mathcal{C}_q^N(n)$ that possesses the same indexing structure as $\hat{c}$ and is determined by the following equations:

1) If $I \in \mathsf{Leaf}(\hat{c})$, then $\hat{e}_I = e_I$ is equal to the cell $\mathsf{clean}(c_I)$ (in the sense of Definition 2.13);
2) If $I \in \mathsf{Junc}(\hat{c})$, then $\hat{e}_I$ is equal to the super cell $(e_I, (\hat{e}_{I1}, \dots, \hat{e}_{Ip(I)}))$ where each super cell $\hat{e}_{Ii}$ is defined by induction and where the cell $e_I$ is determined by the following equations:
   2.1) *Energy storage:* $\mathsf{res}(e_I) = \mathsf{res}(c_I) + \mathcal{S}\mathsf{cyt}(c_I)$
   2.2) *Disorder cleaning:* $\mathsf{cyt}(e_I) = (0, 0, \dots, 0)$
   2.3) *Homeostatic state:* $\mathsf{org}(e_I) = (K(e_{I1}), \dots, K(e_{Ip(I)}))$

From now on, the cleaning $\hat{e}$ of $\hat{c}$ will be denoted as $\mathsf{clean}(\hat{c})$.

*Remark* 2.17 (Hierarchical learning and data integration). In our implementation, the ability to clean a super cell is essential to making the algorithm to learn and classify data. More specifically, cleaning allows a cell to communicate their learning progress to their parent cell in the tree structure. To understand why this is the case, we will use the following schematic, which represents various update states through which a cell and its parent go during learning.

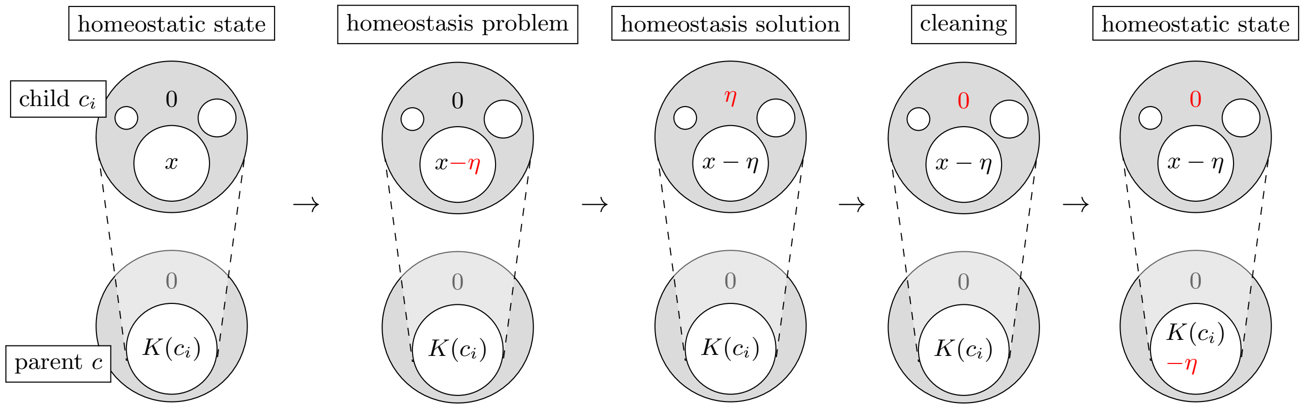


The first stage, from the left, represents a cell and its parent whose associated ambient super cell (not represented) is in a homeostatic state (Definition 2.11). Because the cytosolic content of the child cell $c_i$ is 0, the corresponding organelle $\mathsf{org}(c)_i$ of the parent cell $c$ is equal to the content $K(c_i)$ of the child cell (Definition 1.22). The second stage, from the left, shows a situation in which the organelle $x$ of the child cell $c_i$ undergoes a small change $-\eta$ (this kind of modification will be used in section 2.3 to make a super cell learn). After such a modification, the super cell might no longer be in a homeostatic state, so we re-establish fitness between the cells by solving the homeostasis problems induced by these modifications (see Definition 1.40). In the third stage, we can see that solving the homeostasis problems changes the values of the cytosolic contents. While these added values ensure fitness, they prevent the organelle $\mathsf{org}(c)_i$ of the parent cell $c$ to report what has been learned in its child cell $c_i$. We therefore proceed to a cleaning of the super cell, which we illustrate in the last two stages: the second last stage illustrates the cleaning of the child cell $c_i$ (Definition 2.13) and the last stage ensures that the child cell $c_i$ fits the organelle $\mathsf{org}(c)_i$. As a result, the organelle $\mathsf{org}(c)_i$ accounts for the information learned by its associated child cell $c_i$.

2.3. **Allostasis.** The first step of our algorithm, called `Allostasis`, is strongly inspired from gradient descent techniques. The goal of this step is to *minimize* the values of the algebra operators (Definition 1.34) associated with each of the junctions of a super cell. To do so, we use homeostasis problems as back propagation mechanisms (section 1.6). The main difference with the usual gradient descent is that we use a different concept of differentials, namely the concept of allostatic differential given in Definition 1.48. As with usual gradient descent, we will use a "gamma parameter". In our case, the gamma parameter is a function of the following form:

$$\gamma : \mathcal{C}^N(m) \times \Delta_N(A)^p \times [p] \times [N] \to \mathbb{R}$$

In Definition 2.18, given below, we use a parameter gamma whose third argument is the pre-action associated with an expected signal (Definition 2.12). In the spirit of gradient descent, the output of the gamma parameter should be viewed as an intensity, while the allostatic differential associated with it (see Definition 2.12) should be viewed as a directionality (through its positive and negative values).

**Definition 2.18** (Allostatic matrix)**.** Let $\hat{c}$ be a super cell in $\mathcal{C}_q^N(n)$ and let $a$ be a vector in $\Delta_N(A)^n$. For every $I \in \mathsf{Index}(\hat{c})$ such that $\mathsf{base}(\hat{c}_I) = (c_I, (d_1, d_2, \dots, d_p))$, we denote by $\mathsf{All}(\hat{c}|a)_I$ the $p \times N$-matrix whose $(j, v)$-coefficient is equal to the following multiplication:

$$\gamma(d_j, \mathsf{esgn}(\hat{c}|a')_I, j, v) \times \left( \frac{\partial_{j,v} U^2(\mathsf{base}(\hat{c}_I))}{\partial(\mathsf{base}(\hat{c}_I))} \right). \tag{2.3}$$

The following definition formalizes the tuning procedure illustrated in Remark 2.17.

**Definition 2.19** (Allostatic tuning)**.** Let $\hat{c}$ be a super cell in $\mathcal{C}_q^N(n)$, let $a$ be a vector in $\Delta_N(A)^n$ and let $I$ be in $\mathsf{Index}(\hat{c})$. Suppose that we are given a homeostasis problem $\lambda$ for $\mathsf{base}(\hat{c}_I) = (c_I, d(\hat{c}_I))$ such that:

$$\lambda_{i,u} \text{ is equal to either } \mathsf{org}(c_I)_{i,u} \text{ or } \mathsf{org}(c_I)_{i,u} - \mathsf{All}(\hat{c}|a)_{I,i,u}. \tag{2.4}$$

If we let $(\kappa, e_I, d_I)$ denote the homeostasis solution of $\lambda$ (see the equations below), then we define the *allostatic tuning* $\mathsf{tune}(\hat{c}|\lambda)_I$ of $\hat{c}$ *relative to* $(\lambda, I)$ as the super cell $(e_I, d_I)$ if $I \in \mathsf{Index}(\hat{c})$ and as the cell $e_I$ if $I \in \mathsf{Leaf}(\hat{c})$.

$$\begin{cases} e_I &= \mathsf{left}\big(c_I \circ d(\hat{c}_I)\big)\big(\mathsf{res}(d(\hat{c}_I)), \kappa, \lambda\big) \\ d_I &= \mathsf{right}\big(c_I \circ d(\hat{c}_I)\big)\big(m(\hat{c}_I), \mathsf{res}(d(\hat{c}_I)), \kappa\big) \end{cases}$$

*Remark* 2.20 (Homeostatic state). In our implementation, we compute formula (2.3) by using the formula of Proposition 1.49. In order to use Proposition 1.49, we need to make sure that the super cell $\hat{c}$ is in a homeostatic state (in our implementation, this is done through the methods `spontaneous_reaction` associated with the classes `Cell` and `SuperCell`). Such a condition is necessary because Proposition 1.49 relies on formula (1.4), which requires the super cell $\hat{c}$ to be in a homeostatic state. Note that this homeostatic state is always maintained through allostatic tuning due to the fitness properties satisfied by homeostasis solutions (see Definition 1.40).

*Remark* 2.21 (Existence of a tuning). While Theorem 1.42 states that homeostasis solutions always exist and are unique, it is important to note that a homeostasis problem $\lambda$ may be ill-defined when some of the coefficients $\lambda_{i,u}$ are negative (which is forbidden by Definition 2.19). As a result, the allostatic tuning of a super cell (Definition 2.19) may not always exist. This conflict can be solved algorithmically by setting the corresponding coefficients $\mathsf{All}(\hat{c}|a)_{I,i,u}$ to 0 so that $\lambda_{i,u}$ stays non-negative (see formula (2.4)). In our implementation, we use this kind of practice in order to find a well-defined allostatic tuning (in the sense of Definition 2.15); for more detail, see the methods `compute_variable`, `homeostasis` and `allostasis` associated with the class `SuperCell`.

In our implementation, we use gamma parameters of the form shown below, where $c \in \mathcal{C}^N(n)$ and $a \in \Delta_N(A)^n$, where the function $\mathsf{agr}$ is defined in Definition 2.22 (in our implementation, the function $\mathsf{agr}$ is implemented through the method `agreement`) and where the function $\mathsf{magr}$ is defined in Definition 2.23.

$$\gamma(c,a,j,u) = \begin{cases} 0 & \text{if learning should not occur according to specific criteria;} \\ 10^E \times \left( \dfrac{\mathsf{agr}(c,j,a_j)}{\mathsf{magr}(c,a)} \right)^F & \text{otherwise.} \end{cases}$$

The idea behind the previous formula is that the quantity $\mathsf{agr}(c,j,a_j)$ measures a correlation between $c$ and $a_j$ (a value between 0 and 1), the quantity $\mathsf{magr}(c,a)$ makes sure that the correlation measured by $\mathsf{agr}(c,j,a_j)$ is not biased towards a certain type of input $a$, the exponent $F$ attenuates the influence of the normalized correlation $\mathsf{agr}(c,j,a_j)/\mathsf{magr}(c,a)$ when its values are low and the power $10^E$ makes sure that the values $\gamma(c,j,a)$ land within a specific numerical interval. Note that the scenarios in which $\gamma(c,a,j,u)$ is set to 0 are relatively easy to formalize in our case due to the interpretability of the information learned by the cells (see the tutorial of the documentation).

**Definition 2.22** (Agreements)**.** Below, for every vector $x = (x_1, \dots, x_N)$, we let $\|x\|$ denote the norm $\sqrt{\sum x_u^2}$ of $x$. Let $c$ be a cell in $\mathcal{C}^N(n)$ and $a$ be a vector in $\Delta_N(A)$. For every integer $j \in [N]$, we define *the $j$-th agreement* $\mathsf{agr}(c,j,a)$ *of $c$ with $a$* as the following normalized scalar product

$$\mathsf{agr}(c,j,a) := \sum_{u=1}^{N} \left( \frac{\mathsf{org}(c)_{j,u}}{\|\mathsf{org}(c)_j\|} \times \frac{a_u}{\|a\|} \right)$$

The agreement of $c$ with $a$ can be interpreted as the cosine of the angular distance between $\mathsf{org}(c)_j$ and $a$ (also known as "cosine similarity").

**Definition 2.23** (Max-agreements)**.** Let $c$ be a cell in $\mathcal{C}^N(n)$ and $a$ be a vector in $\Delta_N(A)^n$. We define *the max-agreement* $\mathsf{magr}(c,a)$ *of $c$ with $a$* as the maximum of the values $\mathsf{agr}(c,j,a_j)$ for every $j \in [N]$.

$$\mathsf{magr}(c,a) := \mathsf{max}\{\mathsf{agr}(c,j,a_j) \mid j \in [N]\}$$

We finally give the pseudo code for the step `Allostasis`. In our implementation, this code is implemented through the methods `compute_variable`, `homeostasis`, and `allostasis` (see the documentation).

**`Allostasis`**

**Input:** $(\hat{c}, a, \gamma)$ where $\hat{c}$ is in a homeostatic state
**For** every indexing collection $I \in \mathsf{Index}(\hat{c})$ taken in the decreasing order **do:**
  **Compute** $\mathsf{All}(\hat{c}|a)_I$
  **Set** $\lambda_{I,i,u} := \mathsf{org}(c_I)_{i,u} - \mathsf{All}(\hat{c}|a)_{I,i,u}$
**For** every indexing collection $I \in \mathsf{Index}(\hat{c})$ taken in the decreasing order **do:**
  **If** $I \in \mathsf{Leaf}(\hat{c})$ **do:**
    **Update** $(c_I, (c_{I1}, \dots, c_{Ip(I)})) \leftarrow \mathsf{tune}(\hat{c}|\lambda_I)_I$
  **If** $I \in \mathsf{Leaf}(\hat{c})$ **do:**
    **Update** $c_I \leftarrow \mathsf{tune}(\hat{c}|\lambda_I)_I$
  **While** the cells of $\hat{c}_I$ are not well-defined (Definition 2.15) **do:**
    **Update** the conflicting $\lambda_{I,i,u}$ to $\mathsf{org}(c_I)_{i,u}$ (see Remark 2.21)
    **If** $I \in \mathsf{Leaf}(\hat{c})$ **do:**
      **Update** $(c_I, (c_{I1}, \dots, c_{Ip(I)})) \leftarrow \mathsf{tune}(\hat{c}|\lambda_I)_I$
    **If** $I \in \mathsf{Leaf}(\hat{c})$ **do:**
      **Update** $c_I \leftarrow \mathsf{tune}(\hat{c}|\lambda_I)_I$
**Return** $\hat{c}$

2.4. **Composition of cells.** The present section describes the step `Compose` of our algorithm. This step is used to complete the other two steps `Fusion` and `Fission` described in section 2.5. The main reason for this two-step completion is to avoid composing both leaf and non-leaf super cells within the same cell, because this operation would not return a super cell (see Definition 2.1).

**Definition 2.24** (Binary configurations)**.** A *binary configuration* for a super cell $\hat{c}$ in $\mathcal{C}_q^N(n)$ is a function $\beta : \mathsf{Index}(\hat{c}) \to \{0,1\}$. We organize the binary values $\beta_I$ of $\beta$ as a tree structure of the same form as the underlying tree structure encoding $\hat{c}$, namely we define by induction:

1) $\hat{\beta}_I := \beta_I$ for every $I \in \mathsf{Leaf}(\hat{c})$.
2) $\hat{\beta}_I := (\beta_I, (\hat{\beta}_{I1}, \dots, \hat{\beta}_{Ip(I)}))$ for every $I \in \mathsf{Junc}(\hat{c})$;

Each number 1 and 0 of the binary configuration specifies whether the outer cell of the super cell is to be composed within its outer environment or not. For instance, if a cell $c_I$ is associated with a number $\beta_I = 1$, then this means that we intend to compose the cell $c_I$ within its parent cell (see Definition 2.25).

As mentioned at the beginning of this section, the main reason for completing fusion and fission events in a separate composition step is that fusion and fission events may require to handle simultaneous compositions of leaf and non-leaf cells within the same cells. These mixed compositions can conflict with the definition of a super cell, in which the children of a parent cell are either all proper super cells (all of height greater than 0) or all leaf super cells (all of height equal to 0). We handle these potential conflicts through Definition 2.25, in which all-leaf-compositions are treated in item 2.1 and mixed-compositions are treated in item 2.2.

**Definition 2.25** (configured composition)**.** Let $\beta$ be a binary configuration for a super cell $\hat{c}$ in $\mathcal{C}_q^N(n)$. We define the *configured composition of $\hat{c}$ relative to $\beta$* as the super cell $\mathsf{comp}(\hat{c}|\beta)$ defined recursively as follows:

1) If $\hat{c}$ is a leaf (*i.e.* a cell), then $\mathsf{comp}(\hat{c}|\hat{\beta}) = \hat{c}$;
2) If $\hat{c} = (c, (\hat{c}_1, \dots, \hat{c}_p))$ is a junction and:
   2.1) all super cells $\hat{c}_1, \dots, \hat{c}_p$ are leaves for which $\beta_1 = \cdots = \beta_p = 1$, then we define:
   $$\mathsf{comp}(\hat{c}|\hat{\beta}) = c \circ (\hat{c}_1, \dots, \hat{c}_p)$$
   2.2) otherwise, we proceed as follows: for every $i \in [p]$, we define the numerical variable
   $$\nu_i = \begin{cases} 0 & \text{if } \mathsf{comp}(\hat{c}_i|\hat{\beta}_i) \text{ is a cell } c'_i \\ 1 & \text{if } \mathsf{comp}(\hat{c}_i|\hat{\beta}_i) \text{ is a super cell } (c'_i, (\hat{c}'_{i1}, \dots, \hat{c}'_{ip(i)})) \end{cases}$$
   and we define
   $$\begin{cases} \mathsf{comp}(\hat{c}|\hat{\beta}) = (c \circ \mathsf{val}(\hat{c}|\hat{\beta}), (\mathsf{coval}(c|\hat{\beta})_1, \dots, \mathsf{coval}(c|\hat{\beta})_p)) \\ \mathsf{val}(\hat{c}|\hat{\beta}) = (\mathsf{val}(\hat{c}|\hat{\beta})_1, \dots, \mathsf{val}(\hat{c}|\hat{\beta})_p) \end{cases}$$
   where we denote:
   $$\mathsf{val}(\hat{c}|\hat{\beta})_i = \begin{cases} c'_i & \text{if } \beta_i = 1 \\ \mathsf{id}_{K(c'_i)} & \text{if } \beta_i = 0 \end{cases} \qquad \mathsf{coval}(c|\hat{\beta})_i = \begin{cases} \hat{c}'_{i1}, \dots, \hat{c}'_{ip(i)} & \text{if } \beta_i = 1 \text{ and } \nu_i = 1 \\ (\mathrm{id}_{\mathsf{org}(c'_i)_1}, \dots, \mathrm{id}_{\mathsf{org}(c'_i)_p}) & \text{if } \beta_i = 1 \text{ and } \nu_i = 0 \\ \mathsf{comp}(\hat{c}_i|\hat{\beta}_i) & \text{if } \beta_i = 0 \end{cases}$$

The pseudo-code of the step `compose` is straightforward and mainly relies on Definition 2.25. In our implementation, this code is implemented through the method `compose` of the class `SuperCell` (see the documentation).

| `Compose` |
| --- |
| **Input:** $(\hat{c}, \hat{\beta})$ |
| **Return** $\mathsf{comp}(\hat{c}|\hat{\beta})$ |

2.5. **Merging and dividing cells.** The present section describes the steps `fusion` and `fission`. Our implementation of these steps mainly relies on the statement of Theorem 1.58 and the interpretation given in Remark 1.59. To give an intuition, we illustrate an example of the step `fission` below, on the left, and an example of the step `fusion` on the right. As can be seen, both steps consists in adding new compartments (in darker gray) to the original cell. The integers 0 and 1 next to each compartment represent the values of an associated binary configuration (Definition 2.24), which we construct according to the intuition described in Example 1.53.

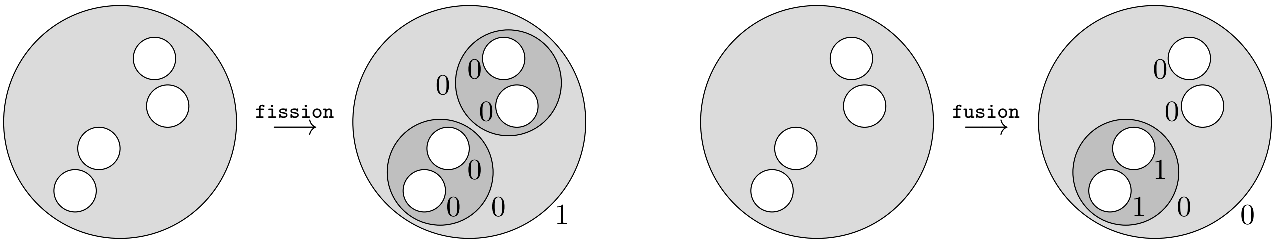


For `fusion`, we separate the organelles of the cell in two groups, distributed within two new compartments, and we compose the outer membrane of the super cell within the environment – when is possible. For `fission`, we also separate the organelles of the cell in two groups, but we only create a compartment for one of the groups, and we compose the super cells associated with the organelles of that group within the new compartment.

The following definition simulates a cell division at the root of a super cell.

**Definition 2.26** (Division)**.** Let $\hat{c}$ be a super cell in $\mathcal{C}_q^N(n)$ in a homeostatic state such that $q > 0$ and $\beta$ be a binary configuration for $\hat{c}$. Supposing that $\hat{c} = (c, (\hat{c}_1, \dots, \hat{c}_p))$, we let $S = \{i_1 < i_2 < \cdots < i_k\}$ be a subset of $[p]$. We define the *basal division* of $(\hat{c}, \hat{\beta})$ *relative to* $S$ as a pair $(\hat{e}, \hat{\tau})$ where $\hat{\tau} = (\tau, (\hat{\tau}_1, \hat{\tau}_2))$ is a binary configuration for the super cell $\hat{e} = (e, (\hat{e}_1, \hat{e}_2))$ defined as follows:

1) the cell $e$ is in $\mathcal{C}^N(p - k + 1)$ and is defined as the tensor cell
$$\xi_2\Big(\sum_{j \in S} \mathsf{org}(c)_j, \sum_{j \notin S} \mathsf{org}(c)_j\Big)$$
while we have the identity $\tau = 1$;
2) the super cells $\hat{e}_1$ and $\hat{e}_2$ are determined by the following relations
$$\hat{e}_i = \begin{cases} (e_1, (\hat{c}_j)_{j \in S}) & \text{if } i = 1 \\ (e_2, (\hat{c}_j)_{j \notin S}) & \text{if } i = 2 \end{cases}$$
where the cells $e_1$ and $e_2$ are determined by the equations
$$\begin{array}{lll} \mathsf{res}(e_1) = \mathsf{res}(c)/2, & \mathsf{cyt}(e_1) = 0, & \mathsf{org}(e_1) = (\mathsf{org}(c)_j)_{j \in S}, \\ \mathsf{res}(e_2) = \mathsf{res}(c)/2, & \mathsf{cyt}(e_2) = 0, & \mathsf{org}(e_2) = (\mathsf{org}(c)_j)_{j \notin S}, \end{array}$$
while we have the identity $\hat{\tau}_i = \hat{\beta}_i$ for every $i \in [p - k + 1]$.

The following definition simulates a cell merging at the root of a super cell.

**Definition 2.27** (Merging)**.** Let $\hat{c}$ be a super cell in $\mathcal{C}_q^N(n)$ in a homeostatic state such that $q > 0$ and $\beta$ be a binary configuration for $\hat{c}$. Supposing that $\hat{c} = (c, (\hat{c}_1, \dots, \hat{c}_p))$, we let $S = \{i_1 < i_2 < \cdots < i_k\}$ be a subset of $[p]$. We define the *basal merging* of $(\hat{c}, \hat{\beta})$ *relative to* $S$ as a pair $(\hat{e}, \hat{\tau})$ where $\hat{\tau} = (\tau, (\hat{\tau}_1, \dots, \hat{\tau}_{p-k+1}))$ is a binary configuration for the super cell $\hat{e} = (e, (\hat{e}_1, \dots, \hat{e}_{p-k+1}))$ defined as follows:

1) the cell $e$ is in $\mathcal{C}^N(p - k + 1)$ and is determined by the relations $\mathsf{res}(e) = \mathsf{res}(c)$, $\mathsf{cyt}(e) = 0$ and
$$\mathsf{org}(e)_i = \begin{cases} \mathsf{org}(c)_i & \text{if } i < i_1 \\ \sum_{j \in S} \mathsf{org}(c)_j & \text{if } i = i_1 \\ \mathsf{org}(e)_{i-q} & \text{if } i_q < i < i_{q+1} \end{cases}$$

while we have the identity $\tau = \beta$;

2) the super cells $\hat{e}_1, \dots, \hat{e}_p$ are determined by the following relations

$$\hat{e}_i = \begin{cases} \hat{c}_i & \text{if } i < i_1 \\ (\xi_k(K(c_{i_1}), K(c_{i_2}), \dots, K(c_{i_k})), \hat{c}_{i_1}, \dots, \hat{c}_{i_k}) & \text{if } i = i_1 \\ \hat{c}_{i-q} & \text{if } i_q < i < i_{q+1} \end{cases}$$

while we have the following identities:

$$\hat{\tau}_i = \begin{cases} \hat{\beta}_i & \text{if } i < i_1 \\ (0, (\hat{\tau}_{i_1}, \dots, \hat{\tau}_{i_k})) & \text{if } i = i_1 \\ \hat{\beta}_{i-q} & \text{if } i_q < i < i_{q+1} \end{cases} \quad \text{where} \quad \hat{\tau}_{i_j} = (1, (\hat{\beta}_{i_j 1}, \dots, \hat{\beta}_{i_j p(i_j)})).$$

Thus, for both steps `fusion` and `fission`, we divide the organelles in two groups with respect to a subset $S$ of the set of the organelles indices. In the rest of this section, our goal is to explain how we compute the set $S$ for our implementation. We proceed in two steps:

1) we first compute a weighted adjacency matrix on the set of organelle indices (Definition 2.30);
2) and we take $S$ to be one of the maximal connected component of the graph associated with the matrix (see Definition 2.33).

To do so, we use the idea of barycenters discussed in Remark 1.56, which we formalize through Definition 2.29. We start with a notation for tensors of child cells in a super cells.

**Convention 2.28** (Tensor of child cells)**.** Let $\hat{c}$ be a super cell in $\mathcal{C}_q^N(n)$. For every $I \in \mathsf{Junc}(\hat{c})$ such that $\hat{c}_I = (c_I, (\hat{c}_{I1}, \dots, \hat{c}_{Ip}))$ and every subset $S = \{i_1, i_2, \dots, i_k\}$ of $[p]$, we will denote by $c_{I,S}$ the tensor $c_{Ii_1} \otimes c_{Ii_2} \cdots \otimes c_{Ii_k}$.

Definition 2.29 introduces weighted sums of pre-actions, which we use to retrieve the type of actions computed in Proposition 1.55.

**Definition 2.29** (Tensored pre-action)**.** Let $\hat{c}$ be a super cell in $\mathcal{C}_q^N(n)$ and $a$ be a vector in $\Delta_N(A)$. For every $I \in \mathsf{Junc}(\hat{c})$ such that $\hat{c}_I = (c_I, (\hat{c}_{I1}, \dots, \hat{c}_{Ip}))$, let us denote by $(b_1, b_2, \dots, b_p)$ the specialized signal $\mathsf{ssgn}(\hat{c}|a)_I$. We define the *tensor of* $\mathsf{ssgn}(\hat{c}|a)_I$ as the following weighted sum (or barycenter).

$$\mathsf{center}(\hat{c}|a)_I := \sum_{i \in S} b_i \times \frac{\mathcal{S}K(c_{Ii})}{\mathcal{S}K(c_{I,[p]})}$$

The following definition implements the criterion discussed in Remark 1.56, namely the construction of an adjacency matrix that indicates which of the organelles are on the same side of the barycenter $\mathsf{center}(\hat{c}|a)_I$.

**Definition 2.30** (Adjacency matrices)**.** Let $\hat{c}$ be a super cell in $\mathcal{C}_q^N(n)$ and $a$ be a vector in $\Delta_N(A)$. For every $I \in \mathsf{Junc}(\hat{c})$ such that $\hat{c}_I = (c_I, (\hat{c}_{I1}, \dots, \hat{c}_{Ip}))$, let us denote by $(b_1, b_2, \dots, b_p)$ the specialized signal $\mathsf{ssgn}(\hat{c}|a)_I$. For every postive real number $\nu$, we define two $(p \times p)$-matrices $\pi_\nu^-(I)$ and $\pi_\nu^+(I)$ as follows (where $\#$ denotes the cardinality operation):

$$\pi_\nu^-(I,a)_{i,j} := \#\{u \in [N] \mid b_{i,u} \text{ and } b_{j,u} \text{ are less than } \nu \times \mathsf{center}(\hat{c}|a)_{I,u}\}$$
$$\pi_\nu^+(I,a)_{i,j} := \#\{u \in [N] \mid b_{i,u} \text{ and } b_{j,u} \text{ are greater than } \nu \times \mathsf{center}(\hat{c}|a)_{I,u}\}$$

From now on, the two weighted adjacency matrices $\pi_\nu^-(I,a)$ and $\pi_\nu^+(I,a)$ will be viewed as their associated weighted graphs. Below, we use the matrices $\pi_\nu^-(I,a)$ and $\pi_\nu^+(I,a)$ for cell fusion and cell fission, respectively.

**Definition 2.31** (Connected components)**.** Let $\hat{c}$ be a super cell in $\mathcal{C}_q^N(n)$ and $a$ be a vector in $\Delta_N(A)$. For every $I \in \mathsf{Junc}(\hat{c})$, every $\varepsilon \in \{+,-\}$ and non-negative integer $\omega$, we denote by $\mathsf{clust}_\nu^\varepsilon(I,a|\omega)$ the set of subsets $S \subseteq [p]$ such that

1) the maximum weight $m = \max\{\pi_\nu^\varepsilon(I,a)_{i,j} \mid i,j\}$ in the graph $\pi_\nu^\varepsilon(I,a)$ is greater than or equal to $\omega$;
2) there is a path linking the edges in $S$ through vertices of maximum weight $m$ in the graph $\pi_\nu^\varepsilon(I,a)$;
3) $S$ has a maximal cardinal among all subsets satisfying item 1.

While the elements of $\mathsf{clust}^{\varepsilon}_{\nu}(I, a|\omega)$ allow us to select groups of organelles with respect to one of the inequalities discussed in Remark 1.59, namely the 'signal comparison', it does not tell us anything about the second type of comparison, which compares the proportions in the organelles of the cell. We take care of this second step through the scoring system defined in Definition 2.32. In this definition, we replace the organelle vectors with content vectors, assuming that the underlying super cell in a homeostatic state (see Definition 2.11). Also, for every subset $S \subseteq [p]$, we will write $[p]\backslash S$ to denote the complement of $S$ within $[p]$.

**Definition 2.32** (Quality)**.** Let $\hat{c}$ be a super cell in $\mathcal{C}^N_q(n)$ and $a$ be a vector in $\Delta_N(A)$. For every $I \in \mathsf{Junc}(\hat{c})$, every $\varepsilon \in \{+,-\}$ and non-negative integer $\omega$, we define the *quality* of a set $S \in \mathsf{clust}^{\varepsilon}_{\nu}(I, a|\omega)$ as the sum given below, on the left.

$$\mathsf{qual}(S) := \sum_{u \in \mathbf{B}(I,S)} \frac{K(c_{I,S})_u}{\mathcal{S}K(c_{I,S})} \times \frac{\mathcal{S}K(c_{I,[p]\backslash S})}{K(c_{I,[p]\backslash S})_u} \quad \text{where} \quad \mathbf{B}(I,S) = \left\{ u \in [N] \;\middle|\; \frac{K(c_{I,S})_u}{\mathcal{S}K(c_{I,S})} > \frac{K(c_{I,[p]\backslash S})_u}{\mathcal{S}K(c_{I,[p]\backslash S})} \right\}.$$

**Definition 2.33** (Best-compartments)**.** Let $\hat{c}$ be a super cell in $\mathcal{C}^N_q(n)$ and $a$ be a vector in $\Delta_N(A)$. For every $I \in \mathsf{Junc}(\hat{c})$, every $\varepsilon \in \{+,-\}$ and non-negative integer $\omega$, we say that a set $S \in \mathsf{clust}^{\varepsilon}_{\nu}(I, a|\omega)$ is a *best choice* if the quality $\mathsf{qual}(S)$ is maximal among all the qualities of the elements of $\mathsf{clust}^{\varepsilon}_{\nu}(I, a|\omega)$.

We now give the pseudo-code of the steps `fusion` and `fission`. In our implementation, this code is implemented through the methods `best_compartment` and `proposed_clustering` associated with the class `Cell` and through the methods `merge_base`, `fusion`, `divide_base`, and `fission` associated with the class `SuperCell`.

| `Fusion` (if $\varepsilon = -$) / `Fission` (if $\varepsilon = +$) |
|---|
| **Input:** $(\hat{c}, \beta, a, \varepsilon, \nu, \omega)$ where $\beta$ is a binary configuration for $\hat{c}$ |
| **Update** $\hat{c} \leftarrow \mathsf{clean}(\hat{c})$ |
| **For** every indexing collection $I \in \mathsf{Index}(\hat{c})$ taken in the decreasing order **do:** |
| **Compute** $\mathsf{clust}^{\varepsilon}_{\nu}(I, a\|\omega)$ |
| **Select** a best compartment $S$ in $\mathsf{clust}^{\varepsilon}_{\nu}(I, a\|\omega)$ |
| **If** $\varepsilon = +$ **do:** |
| **Set** $(\hat{e}, \hat{\tau})$ as the basal division of $(\hat{c}_I, \hat{\beta}_I)$ relative to $S$ (Definition 2.26) |
| **If** $\varepsilon = -$ **do:** |
| **Set** $(\hat{e}, \hat{\tau})$ as the basal merging of $(\hat{c}_I, \hat{\beta}_I)$ relative to $S$ (Definition 2.27) |
| **Update** $\hat{c}_I \leftarrow \hat{e}$ |
| **Update** $\hat{\beta}_I \leftarrow \hat{\tau}$ |
| **Return** $(\hat{e}, \hat{\beta})$ |

#### 3. Link with operad theory

While we never really tried to formalize our model in categorical terms, it may have occurred to the learned reader that our model takes place within the world of 'colored operad' [28]. Specifically, for a given positive integer $N$, the collection of sets $\mathcal{C}^N(1)$, $\mathcal{C}^N(2)$, ... can be organized into a colored operad $\mathcal{D}^N$ on $\mathbb{R}^N_+$. This colored operad is given by a collection of sets $\mathcal{D}^N(x_1, \dots, x_n; y)$ containing all those cells $c = (n, C, x, (x_i)_i)$ for which $y = K(c)$ is in $\mathbb{R}^N_+$. These sets are equipped with identities $\mathsf{id}_a \in \mathcal{D}^N(a; a)$ (see Proposition 1.10) and satisfy the usual composition axioms for operads (see Theorem 1.17). From this point of view, we have used the operad $\mathcal{D}^N$ every time we composed a cell that fitted the organelle of another cell.

Ideally, we could use the operadic point of view to further simplify our model. For instance, the operad $\mathcal{D}^N$ can better be described by the colored operad $\mathcal{D}^N_n : (\mathbb{R}^N_+)^n \times \mathbb{R}^N_+ \to \mathbf{Set}$ mapping every tuple $(x_1, \dots, x_n, y)$ in $(\mathbb{R}^N_+)^n \times \mathbb{R}^N_+$ to the set $\mathbb{R}_+$ such that its identities are the 0 elements of the sets $\mathcal{D}^N_1(a, a) = \mathbb{R}_+$ and its compositions are given by the sums of real numbers of the following type:

$$\mathcal{D}_n^N(x_1, \ldots, x_n, y) \times \prod_{i=1}^{n} \mathcal{D}_{m_i}^N(z_{i,1}, \ldots, z_{i,m_i}, x_i) \to \mathcal{D}_{m_1+\cdots+m_n}^N(z_{1,1}, \ldots, z_{n,m_n}, y).$$

Interestingly, the tensor operations defined in Convention 1.51 can also be defined as sum operations of the following form:

$$\otimes : \mathcal{D}_n^N(x_1, \ldots, x_n, y) \times \mathcal{D}_m^N(x'_1, \ldots, x'_m, y') \to \mathcal{D}_{n+m}^N(x_1, \ldots, x_n, x'_1, \ldots, x'_m, y + y').$$

Such a reformulation could be useful to better understand the mechanisms pertaining to the evolution of super cells during a learning phase. Specifically, with this formalism, super cells can be defined in terms of a *free* colored operad on $\mathcal{D}^N$. This kind of formalization could be useful to clarify the relationships that exist between cell fission, cell fusion, and homeostasis. In particular, this clarification could be used to investigate potential improvements of our current implementation.
