## Supplementary material for "Cellular intelligence: dynamic specialization through non-equilibrium multi-scale compartmentalization": Documentation (including computational method)

### DOCUMENTATION FOR THE PYTHON LIBRARY INTCYT

A PROJECT FROM KELLIS LAB
COMPBIO, MIT

contact: Rémy Tuyéras


VERSION OF May 28, 2021

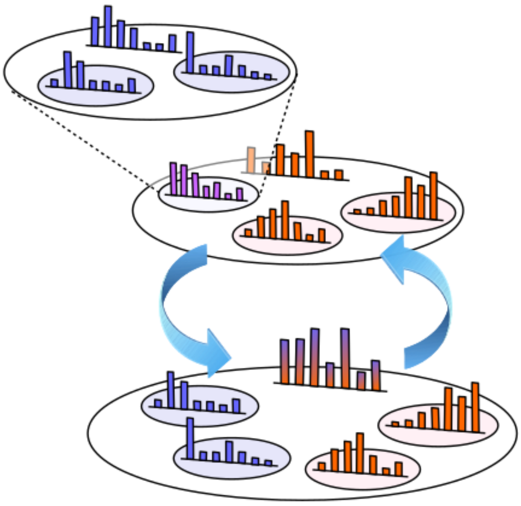

Typeset in $\LaTeX\,2_\varepsilon$.

*Chapter 1*

### Introduction

#### 1.1. About IntCyt

IntCyt is the result of an interdisplinary collaboration [2, 3] that took place at MIT, in Kellis lab (`http://compbio.mit.edu/`), from Fall 2019 to Summer 2020. Overall, IntCyt can be described as both an **unsupervised learning algorithm** and a **python library** whose functions and classes can be used to improve and refine the learning algorithm.

In a few words, the learning algorithm provided with IntCyt is inspired from living cell dynamics, biological compartmentalization principles and non-equilibrium tuning and can be classified as a hierarchical clustering on-the-fly gradient-descent-based algorithm. The main characteristics of the IntCyt learning algorithm are:

1) its ability to simultaneously learn and compartmentalize data through architectural reshaping of its network;
2) its high interpretability.

The library of functions and classes provided with intCyt, referred to as `intcyt`, is designed to permit the refinement and improvement of the original version of the IntCyt algorithm, as proposed in [2]. In this sense, the library `intcyt` is meant to be augmented through future updates.

#### 1.2. About this documentation

The present book contains a tutorial (see Chapter 2) showing how to use the various methods and classes contained in `intcyt` as well as a description of the importable functions and importable classes of its sub-modules. The library contains 3 sub-modules, which we describe in three distinct chapters::

- the sub-module `useful.py` is described in Chapter 3;
- the sub-module `celloperad.py` is described in Chapter 4;
- the sub-module `cellint.py` is described in Chapter 5.

The dependencies between the different chapters are shown in the following diagram.

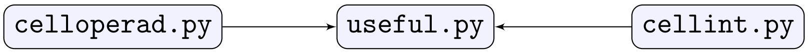

We shall use similar diagrams to specify dependencies between functions and classes. Specifically, each section describing a function or a class will start with a brief description of the item, followed by a dependency flow chart showing the different modules on which this item depend. Below, we give the example of such a flow chart.

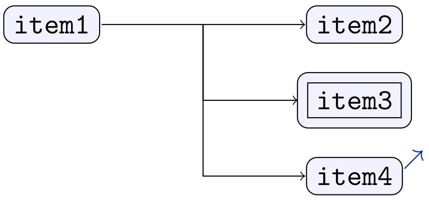

As can be seen above, our flow charts will use three different types of boxes:

1) boxes with no specific decoration will usually frame items (such as classes and functions) that belong to the module described by the chapter;
2) double boxes will frame the name of intermediate items that do not have dependencies with other items (final item);
3) boxes with clickable arrows in the top-right corner will frame items that are defined in external modules. The arrow should send the reader to that specific module;

We will illustrated each description with examples written in python code. There will be two modes: the editor mode and the console mode. The editor mode will mainly be used for the tutorial and will look as follows.

file.py

```
class MyClass:
  """
  This is a comment
  """
  def __init__(self,arg0,arg1):  #This is the constructor of the class
    """
    Another comment about the code
    """
```

The console mode will be used to illustrate the descriptions of classes and functions and will look as follows.

```
>>> P.obj
[0, 1, 2, 3, 4, 5, 6, 7, 8, 9, 10, 11, 13]
```

#### 1.3. Licensing and use of third party libraries

The library IntCyt is provided under the BSD license. Note that IntCyt is exclusively coded in python and uses third party libraries attached to the python package. Specifically, we use the libraries:

- `math` in the modules `celloperad.py` and `useful.py`;
- `random` in the module `celloperad.py`;
- `sys` in the modules `celloperad.py`, `useful.py`, and `cellint.py`;
- `time` in the module `useful.py`.

All the above are associated with the python package and are provided under the Open Source license. Similarly, the libraries used in the tutorial (section 2) are provided under either the Open Source license or the BSD license.

#### 1.4. Acknowledgments

All the following people have participated to the project in one way or another: Leandro Agudelo, Kevin Grove, Manolis Kellis, Anna Kondylis, Burak Kutlu, Anjanet Loon, Soumya P Ram, Rémy Tuyéras.

#### 1.5. Use of software and commands

The IntCyt unsupervised learning software is located in the directory `software` coming with the IntCyt package (see section 2.1) and consists of the following set of python scripts:

- ▷ `intcyt.py`,
- ▷ `challenge.py` (generates `load_initial.gz`),
- ▷ `data_processing.py` (generates `load_initial.gz` and `dream3_raining.gz`),
- ▷ `main.py` (generates `save_roo.gz`, `save_org.gz`, `save_tree.gz` and `c_info.txt`),
- ▷ `visualize.py`.

A detailed description of the possible uses of these scripts is specified below.

▷ `intcyt.py`: this script contains headers pointing to the sub-modules of the library. The presence of this file is required in any directory containing a python script that imports the library `intcyt`, as shown below.

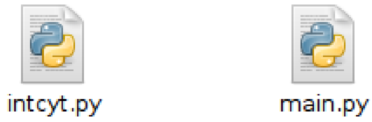

To import the library `intcyt`, use the code shown below in the main script and change the directory paths contained in `intcyt.py` adequately (see section 2).

```
1 from intcyt import *
```

▷ `challenge.py`: this script creates self-supervised learning challenges for IntCyt by using data contained in the `MNIST`, `fashion-MNIST` and `SVHN` datasets (see section 2.7 for more details on this matter). Its launching command possesses a set of options that allow the user to generate a collection of images some of whose parts have been hidden with white noise. The set of options that can be used with the script `challenge.py` is shown below.

```
[python challenge.py]   [fashion-MNIST]  [20]    [-right]
                        [MNSIT]                  [-right-noisy] [1   5]
                        [SVHN]                   [-left]
                                                 [-left-noisy]  [3   7]
```

Each call of the script `challenge.py` starts with `python challenge.py`, followed by the name of the dataset from which the images of the challenges are generated and the number of images generated. For instance, the two commands shown below would pick out 20 images from the `SVHN` dataset and save them in a file named `load_initial.gz`, located in a directory named `result-load` (this directory needs to be created by the user).

```
> mkdir result-load
> python challenge.py SVHN 20
```

To alter images generated by `challenge.py` with white noise, the user can use the other options associated with the script (see above). Below, we describe each of these options with examples.

▷ -right: keeps the right side of the image intact and hides the left side with white noise.

```
> python challenge.py MNIST 10 -right
```

▷ -right-noisy [a b]: keeps the two sides of the image intact and surrounds the picture of the right side with white noise, distributed according to a likelihood of a/b if a and b are specified and a likelihood of 1 otherwise.

```
> python challenge.py fashion-MNIST 10 -right-noisy 0 5
> python challenge.py fashion-MNIST 11 -right-noisy 1 5
> python challenge.py fashion-MNIST 17 -right-noisy 2 5
> python challenge.py fashion-MNIST 20 -right-noisy 3 5
> python challenge.py fashion-MNIST 30 -right-noisy 4 5
> python challenge.py fashion-MNIST 40 -right-noisy 5 5
> python challenge.py fashion-MNIST 50 -right-noisy
```

▷ -left: keeps the left side of the image intact and hides the left side with white noise.

```
> python challenge.py fashion-MNIST 15 -left
```

▷ -left-noisy [a b]: keeps the two sides of the image intact and surrounds the picture of the left side with white noise, distributed according to a likelihood of a/b if a and b are specified and a likelihood of 1 otherwise.

```
> python challenge.py MNIST 10 -left-noisy 0 2
> python challenge.py MNIST 11 -left-noisy 1 2
> python challenge.py MNIST 17 -left-noisy 2 2
> python challenge.py MNIST 50 -left-noisy
```

▷ data_processing.py: this script creates self-supervised learning challenges for IntCyt by using the hidden gene expressions contained in the DREAM3 dataset. It also contains functions to analyze the DREAM3 dataset as well as the results of IntCyt. The set of options that can be used with the script data_processing.py is shown below.

```
[python data_processing.py]    [method]
                               [training]
                               [result      [42000]]
```

Each call of the script data_processing.py starts with python data_processing.py, followed by one of the three options shown above. The first option, method, is meant to generate the analysis described in section 2.8, which the reader is referred to for more details. The second option, training, generates the training data and the initialization of IntCyt relative to the DREAM3 challenge. To use this option, create two directories named data and result-load (if they do not already exist)

```
> mkdir data
> mkdir result-load
```

and use the following command:

```
> python data_processing.py training
```

Finally, we can use the third option result to visualize the results generated by IntCyt. Specifically, to see the result of IntCyt at a certain learning cycle, say 42000, make sure that the file save_org.gz is present in the directory and use the following command:

```
> python data_processing.py result 42000
```

▷ main.py: this script is an implementation of the algorithm described in [1]. Its launching command possesses a set of options that allow the user to run the algorithm with a random initialization or load a pre-existing initialization. Loaded initialization should be stored in a directory named `result-load`, while the "training data" should be located in a directory named `data`. The set of options that can be used with the script `main.py` is shown below.

```
[python main.py]    [fashion-MNIST]   [-load]                 [14520]
                    [MNIST]           [-load-selfsup-right]
                    [SVHN]            [-load-selfsup-left]
```

Each call of the script `main.py` starts with `python main.py`, followed by the name of the dataset to be learned. The following command launches the IntCyt algorithm on the MNIST dataset for a random initialization of its cells (see [1]).

```
> python main.py MNIST
```

To load an existing initialization, which can be generated by either `challenge.py` or `data_processing.py`, use the option `-load<specifiy>`. The following command loads the images stored in the file `load_initial.py` located in `result-load`.

```
> python main.py fashion-MNIST -load
```

The files `save_org.gz` generated by `main.py` can also provide initializations after renaming them as `load_initial.py`. More specifically, a file `save_org.gz` contains a long list of potential initializations, which consist of the states of the organelles at every learning cycle of `main.py` for a given run. Our previous example, using `load` would only load the first of these initializations. To load an initialization located at a certain index, say 14520, use the following command:

```
> python main.py fashion-MNIST -load 14520
```

If the initialization stored in `initial_load.gz` was generated in the context of a self-supervised challenge use the following options instead of `-load`:

▷ `-load-selfsup-right` for challenges whose right side contains the guiding data;

```
> python challenge.py MNIST 20 -right
> python main.py MNIST -load-selfsup-right
> python main.py MNIST -load-selfsup-right 500
```

▷ `-load-selfsup-left` for challenges whose left side contains the guiding data.

```
> python challenge.py fashion-MNIST 20 -left
> python main.py fashion-MNIST -load-selfsup-left
> python main.py fashion-MNIST -load-selfsup-left 12500
```

▷ `visualize.py`: this script displays the memorized parameters of the algorithm in the form of a tree or a table. Its launching command possesses a set of options that allow the user to choose between a table, a series of trees or a table with a tree to display the memory of the algorithm. The set of options that can be used with the script `visualize.py` is shown below:

```
[python visualize.py]    [fashion-MNIST]    [-tree]   [0 5 260 261 ...]
                         [MNIST]            [-forest]
                         [SVHN]
```

Each call of the script `visualize.py` starts with `python visualize.py`, followed by the name of the dataset on which IntCyt was run. The following command displays the set of parameters characterizing the organelles of IntCyt from cycle 0 to cycle 4 and from cycle 100 to cycle 109.

```
> python visualize.py MNIST 0 5 100 110
```

In addition to displaying the organelles saved during a run of INTCYT, we can also display the tree structure associated with the last queried set of organelles by using the option `-tree`. The following command produce the same analysis as that of the previous command.

```
> python visualize.py fashion-MNIST -tree 0 5 100 110
```

Finally, we can request to see all the tree structures adopted by `IntCyt` during the learning stage by using the option `-forest`, as shown below.

```
> python visualize.py fashion-MNIST -forest 0 5 100 110
```

*Chapter 2*

### Tutorial

#### 2.1. Installation and preparation

To install the library, go to `https://github.com/remytuyeras/intcyt-library` and download the library package by clicking on the green download button.

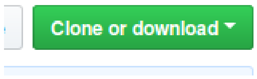

In your download directory, you should see a zip file named `intcyt-library-master.zip`. Copy this file in a new directory and extract its content with a zip extraction application. You should have the following display.

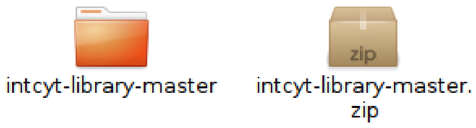

Now, enter the directory `intcyt-library-master`, in which you should see the following files.

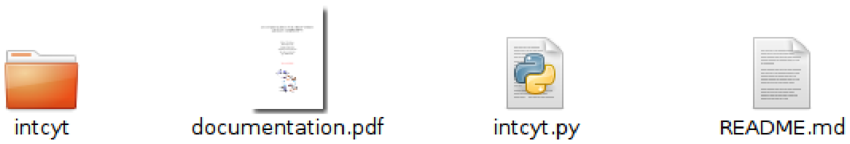

The directory named `intcyt` contains the codes of the functions of the library and the python file `intcyt.py` contains the headings referring to the different modules of the library. For the purpose of this tutorial, we will need three additional files:

- ▷ a python file `main.py` in which we will write an algorithm that learns a dataset of images;
- ▷ a python file `visualize.py` in which we will write a program that displays the progress of our learning algorithm;
- ▷ a python file `challenge.py` in which we will write a program that generates learning challenges for self-supervision tests;

To do this properly, create a new directory called `user` and copy the file `intcyt.py` in it.

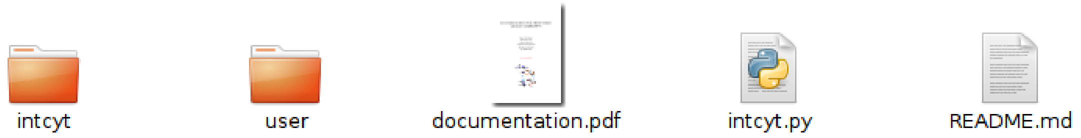

Then, go to `user` and open the file `intcyt.py`: you should see several instances of the function `sys.path.insert` containing directory paths in their second arguments, as shown below.

**intcyt.py**

```
import sys
sys.path.insert(0, "intcyt/useful/")
from useful import *
```

Add the text `../` at the beginning of every path passed to the function `sys.path.insert`, as shown below.

**intcyt.py**

```
import sys
sys.path.insert(0, "../intcyt/useful/")
from useful import *
```

Once the paths are all updated, go back to the directory `user` and create three files named `main.py`, `visualize.py`, and `challenge.py` as well as two directory `data` and `result-load`, as shown below.

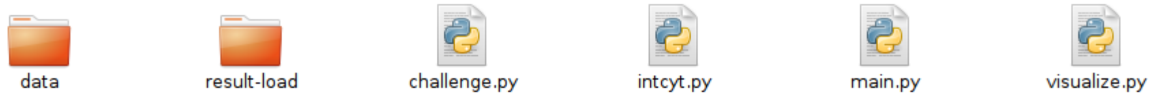

We are now ready to use the library – open the three files `main.py`, `visualize.py` and `challenge.py` and proceed to section 2.2.

#### 2.2. Third party libraries and overall organization

In this tutorial, we will use third party libraries to open and read third party machine learning datasets. Specifically, we will use

- `gzip` to read GZ files;
- `numpy` to read the `MNIST` dataset and the `fashion-MNIST` dataset;
- `scipy.io` to read the `SVHN` dataset;
- `matplotlib.pyplot` to display the progress of the algorithm (saved in GZ files).

In particular, one of the main roles of the scripts `main.py` and `challenge.py` will be to generate various types of outputs from the reading of the datasets `MNIST`, `fashion-MNIST`, or `SVHN`. In this respect, we start the scripts `main.py` and `challenge.py` with the following lines of code.

**main.py / challenge.py**

```
from intcyt import *
#------------------
#Libraries to open datasets
#------------------
import gzip
import numpy as np
import scipy.io as sio
```

In `visualize.py`, we will need the library `gzip` and `matplotlib` to open files and display their contents. Hence, we start the file `visualize.py` with the following lines of code.

visualize.py
```
from intcyt import *
import gzip
import matplotlib.pyplot as plt
```

The organization of this tutorial goes as follows: we first focus on writing the code for `main.py` in section 2.3 and section 2.8.5; we then write the code for `visualize.py` in section 2.6, followed by the code of `challenge.py` in section 2.7. Finally, in section 2.8, we conclude with a description of the method used in [2] to solve the `DREAM3` challenge. Below, the diagram shows the normal chronological use of three scripts `challenge.py`, `main.py` and `visualize.py`.

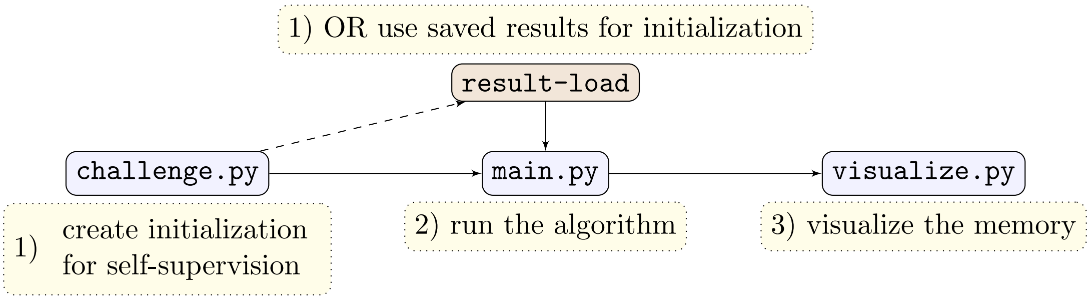

The main goal of the tutorial is to use the IntCyt library to design a light version of the machine learning algorithm presented in [2]. As will be seen, this algorithm can memorize, abstract, organize and infer the data of reference machine learning datasets such as `MNIST`, `fashion-MNIST`, `SVHN` and `DREAM3`.

#### 2.3. Loading datasets

In this section, we focus on the piece of code of `main.py` that allows us to load datasets into the memory. For convenience, we shall integrate the names of these datasets as options in the launch command of the script `main.py`. For example, the following launch command will instruct the script `main.py` to read and learn the data stored in the dataset `fashion-MNIST`.

```
> python main.py fashion-MNIST
```

Below, we present the datasets `MNIST`, `fashion-MNIST` and `SVHN` and explain how these can be downloaded and loaded in the memory.

First, the most challenging dataset that we will test in this tutorial can be found at the address http://ufldl.stanford.edu/housenumbers/ and is known as `SVHN`. This dataset contains RGB images of street numbers as shown below.

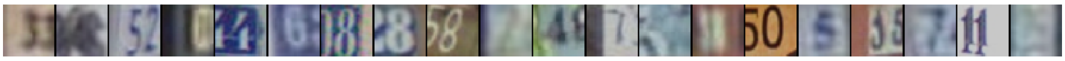

For the sake of this tutorial, download the file `train_32x32.mat` located at the clickable link given above and copy the downloaded file into the directory `data`. Once this is done, add the following piece of code to `main.py` – this will allow us to load the dataset `train_32x32.mat` into the RAM.

main.py
```
#-----------------------------------------------------------------------
#Loading the dataset for learning
#-----------------------------------------------------------------------
#Open the SVHN dataset
if len(sys.argv) > 1 and sys.argv[1] == "SVHN":
  dic_svhn = sio.loadmat("data/train_32x32.mat")
  #Access to the dictionary
  dicim = dic_svhn["X"] #shape:  32, 32, 3, 73257
  dictr = dic_svhn["y"] #shape:  73257, 1
  image_size = [32,32,3] #height, width, depth
```

Now, the easiest dataset that we will test in this tutorial can be found at the address http://yann.lecun.com/exdb/mnist/ and is known as MNIST. This dataset contains black and white images of hand-written numbers as shown below.

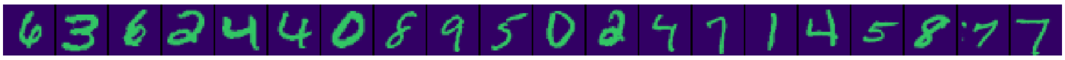

Download the files train-images-idx3ubyte.gz and train-labels-idx1-ubyte.gz linked therein, then copy these files into these directory data and add the following piece of code to main.py.

main.py
```
#------------------
#Open the MNIST dataset
elif len(sys.argv) > 1 and sys.argv[1] == "MNIST":
  fim = gzip.open("data/train-images-idx3-ubyte.gz","r")
  ftr = gzip.open("data/train-labels-idx1-ubyte.gz","r")
  ftr.read(8)
  fim.read(16)
  image_size = [28,28,1] #height, width, depth
```

Finally, an intermediate-level dataset that we will test in this tutorial can be found at the address https://github.com/zalandoresearch/fashion-mnist and is known as fashion-MNIST. This dataset contains black and white images of fashion clothes as shown below.

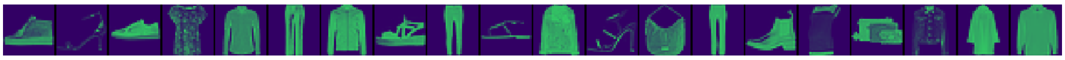

Download the files t10k-images-idx3-ubyte.gz and t10k-labels-idx1-ubyte.gz linked therein, then copy these files into the directory data and add the following piece of code to main.py.

main.py

```
#------------------
#Open the fashion-MNIST dataset
elif len(sys.argv) > 1 and sys.argv[1] == "fashion-MNIST":
  fim = gzip.open("data/t10k-images-idx3-ubyte.gz","r")
  ftr = gzip.open("data/t10k-labels-idx1-ubyte.gz","r")
  ftr.read(8)
  fim.read(16)
  image_size = [28,28,1] #height, width, depth
  categories = ["t-shirt","trousers","pullover", \
                "dress","jacket","sandal","shirt", \
                "sneaker","bag","ankle-boot"]
else:
  exit(0)
```

Once all the previous operations have been completed, the directory `data` should look as follows:

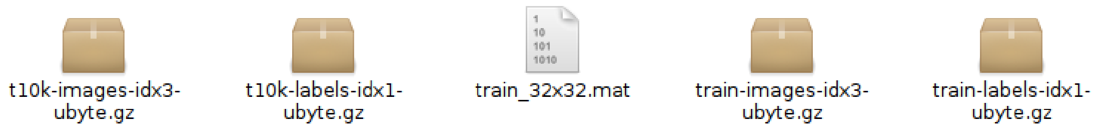

Let us now move on to section 2.4, which introduces the reader to the main tools of the IntCyt library. Even though we will take care of explaining the role of the various functions that we shall use in the code of this tutorial, we will often refer to the corresponding descriptions contained in this documentation for more details.

#### 2.4. Operads, Cells and Super Cells

This section demonstrates the use of the classes `Cell` (section 4.1) and `SuperCell` (section 4.2) and `Operad` (section 4.3), which belong to the module `celloperad.py` (section 4).

More specifically, we use these classes to construct a dynamic database for our algorithm. We will follow the construction of [1], which means that our database will take the form of a *super cell* [1, Def. 2.1]. Throughout each run of our algorithm, the structure of this super cell changes: it starts as one of the simplest types of super cell called a *cell* [1, Def. 1.2] and can develop into a complex hierarchical structure defining a proper super cell.

Mathematically, the evolution of a cell into a proper super cell takes place within an operadic environment. As a result, we shall start this section by calling an operad item. To do so, we call the constructor of the class `Operad` (section 4), which takes the dimensions of the data we want to analyze as an input (*e.g.* the dimensions of the images contained in the datasets). In our case, for any of our datasets, we can compute the dimensions of the associated data by using the lists named `image_size` defined in the previous section. More specifically, the dimensions are given the multiplication of the components of the lists `image_size`. In this respect, we write the following lines of code in the file `main.py`.

main.py

```
#----------------------------------------------------------------------
#Initializing the super cell
#----------------------------------------------------------------------
#Main parameters defining the initial super cell
#------------------
dim = image_size[0] * image_size[1] * image_size[2]
operad = Operad(dim)
```

Viewed from a mathematical standpoint, the item `operad` is supposed to be a map sending any tuple of the form $((x_1, x_2, \dots, x_{\texttt{ary}}), y)$, where $x_i$ and $y$ are lists of length `dim` containing `float` values, to a set of `SuperCell` items – this map is shown below.

$$\mathcal{O}^{\texttt{dim}}_{\texttt{ary}} : \left( \begin{array}{ccccc} \left(\texttt{float}^{\texttt{dim}}\right)^{\texttt{ary}} & \times & \texttt{float}^{\texttt{dim}} & \to & \texttt{Set}(\texttt{SuperCell}) \\ (x_1, \dots, x_{\texttt{ary}}) & , & y & \mapsto & \mathcal{O}^{\texttt{dim}}_{\texttt{ary}}(x_1, \dots, x_{\texttt{ary}}; y) \end{array} \right)$$

Because the encoding of the previous map is absolutely not practical – at least computationally, an `Operad` item is instead designed to only recover the algebraic properties of this mapping. However, the mathematical definition of an operad is still useful conceptually.

Now, with the previous picture in mind, we want encode the database on which our algorithm will rely as one of the elements of the sets

$$\mathcal{O}^{\texttt{dim}}_{\texttt{ary}}(x_1, \dots, x_{\texttt{ary}}; y).$$

Because the parameters `ary` and $x_1, \dots, x_{\texttt{ary}}$ can vary for the same operad $\mathcal{O}^{\texttt{dim}}$, we specify these parameters when initializing the super cell and not when initializing the operad item. In our case, we will consider a super cell whose parameter `ary`, which represents its number of "organelles" [1, Def. 2.7], is equal to 20. For more clarity and flexibility, we will set up variable of the same name, namely `ary`, as shown below.

main.py

```
46 | ary = 20
```

Regarding the initialization of the parameters $x_1, \dots, x_{\texttt{ary}}$, we give two options: either we reload previously constructed super cells, saved in a directory named `result-load`, or we create a new super cell, which will saved in a collection of output files.

To control which of the previous scenarios the user would like to use, let us add a second option to our launch command. Specifically, the system variable `sys.argv[2]` will be assumed to be either empty or contain the string `"-load"`. For example, we will use the following launch command to call our program on the dataset `fashion-MNIST` relative to the parametrization of the super cell saved in the directory `result-load`.

```
> python main.py fashion-MNIST -load
```

Here, the option `-load` will instruct the script `main.py` to initialize the collection of organelles $x_1, \dots, x_{\texttt{ary}}$ with the vectors specified in a file named `load_initial.gz` located in the directory `result-load`. On the other hand, if `sys.argv[2]` is not allocated, then the parameters $x_1, \dots, x_{\texttt{ary}}$ are randomly initialized.

We take care of the option `-load` as follows. First, we open the file `load_initial.gz` in `result-load` by using the module `gzip`. Then, we use the function `usf.get_memory` (see section 3.1.10) to load the first `ary` vectors stored in the file `load_initial.gz`. These vectors play the role of our organelles $x_1, \dots, x_{\texttt{ary}}$ and can be used to construct a `Cell` item (see section 4.1). The resulting `Cell` item can then be turned it into a `SuperCell` item (see section 4.2) in order to permit its growth into a hierarchical structure – this structure will constitute the database of our learning algorithm.

main.py
```
#------------------
#Initializing the organelles and the cytosolic content (either new ones
or saved ones)
#------------------
if len(sys.argv) > 2 and sys.argv[2][:5] == "-load":
  #Load a previously generated tree (height 0)
  fload = gzip.open("result-load/load_initial.gz","r")
  initial_organelles = usf.get_memory(fload,ary,[0])
  fload.close()
  c = Cell(dim,0,[0]*dim,initial_organelles[0])
  sc = SuperCell(c)
```

On the other hand, if the system variable `sys.argv[2]` is not allocated or does not contain the string `"-load"`, then we use the function `operad.generate` (see section 4.3.5) to generate a `SuperCell` item of height 0 whose organelles are randomly initialized.

main.py
```
else:
  #Generate a random tree of height 0
  sc = operad.generate(levels = 0, arity = ary, key = lambda:
random.randint(0,30))
```

It is now a good time to recall a few fact about the overall structure of a super cell. As explained in [1, Supp. text], a super cell is a tree structure whose leaves and junctions are each associated with a `Cell` item. As a consequence, we adapt the terminology of trees to super cells. In this respect, we can say that a super cell possesses: a *root cell*, a (flexible) number of *leaf cells*, and a (flexible) number of *junction cells*, which are each made of a *parent cell* and a collection of *child cells* (see [1, Conv. 2.2]). According to [1, Def. 2.7], the collection of organelles $x_1, x_2, \dots, x_{\mathtt{ary}}$ of a super cell is defined as the concatenation of the collections of organelles associated with the leaf cells.

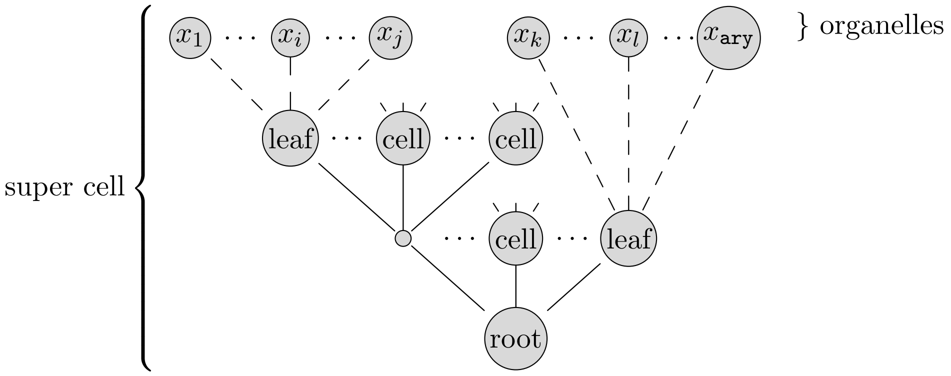

While the organelles of the leaf cells give us the collection of parameters $x_1, x_2, \dots, x_{\mathtt{ary}}$, the root cell determines the parameter $y$ through its 'content' (see [1, Def. 1.21]). In order to visualize or analyze the evolution of a super cell, we want to save all this information in various files. Specifically, we will have three types of files:

- `"save_roo.gz"` will contain vectors describing the evolution of the parameter $y$;
- `"save_org.gz"` will contain vectors describing the evolution of the organelles;
- `"save_tre.gz"` will contain vectors describing the evolution of the tree structure.

Note that the file `"save_tre.gz"` will mostly be used for visualizations, while the other files are more likely to be used for quick access to re-usable information.

In the piece of code given below, we use the three print functions `usf.print_root`, `usf.print_organelles`, and `usf.print_tree` to print the parameters defining the root, the organelles and the tree structure of the super cell in their corresponding files, respectively.

main.py

```
#------------------
#The initial sate of the super cell is saved in the memory
#------------------
fsave_roo = gzip.open("save_roo.gz","w")
fsave_org = gzip.open("save_org.gz","w")
fsave_tre = gzip.open("save_tre.gz","w")
usf.print_root(sc,image_size,fsave_roo,option="save")
usf.print_organelles(sc,image_size,fsave_org,option="save")
usf.print_tree(sc,image_size,fsave_tre,option="save")
fsave_roo.flush()
fsave_org.flush()
fsave_tre.flush()
```

#### 2.5. Learning datasets with IntCyt

In this section, we use the IntCyt algorithm (see [1, Sec. 2]) to memorize, abstract and organize data originating from the datasets presented in section 2.3. Our algorithm will run on two concatenated loops: the outer one will contain 10,000 iterations (for simplicity), while the inner one can be given a flexible number of iterations (called "epochs"). Note that, at each iteration of the outer loop, our algorithm will take a new image from the dataset. On the other hand, the inner loop will iterate our algorithm on the same image.

We read each dataset through two variables: `label` and `inputs`. At every iteration, the variable `inputs` contains a list of `float` values representing an image and the variable `label` indicates the type of image contained by `inputs`.

We start with the dataset `SVHN` as shown below. The variable `label` is a `numpy.ndarray` item containing an integer and the variable `inputs` is a `numpy.ndarray` containing exactly `dim` `float` values.

main.py

```
#--------------------------------------------------------------------
#Running the Learning algorithm
#--------------------------------------------------------------------
iterations = 10000
for i in range(iterations):
  #------------------
  #Get labels and inputs
  #------------------
  if sys.argv[1] == "SVHN":
    label = dictr[i]
    inputs = dicim[:,:,:,i].reshape(dim)
```

In the case of `MNIST`, we read an image and its label as shown below. Again, the variables `label` and `inputs` are `numpy.ndarray` items containing numerical values.

main.py

```
  #------------------
  if sys.argv[1] == "MNIST":
    buf_lab = ftr.read(1)
    buf_inp = fim.read(dim)
    label = np.frombuffer(buf_lab, dtype=np.uint8).astype(np.int64)
    inputs = np.frombuffer(buf_inp, dtype=np.uint8).astype(np.float32)
```

Finally, the following panel shows how we read the dataset `fashion-MNIST`, which is very similar to the dataset `MNIST`, except that the labels here refer to the indices of the strings contained in the list `categories` defined in section 2.3.

main.py

```
  #------------------
  if sys.argv[1] == "fashion-MNIST":
    buf_lab = ftr.read(1)
    buf_inp = fim.read(dim)
    label = map(lambda x:  categories[x], np.frombuffer(buf_lab,
dtype=np.uint8).astype(np.int64))
    inputs = np.frombuffer(buf_inp, dtype=np.uint8).astype(np.float32)
```

The next piece of code is introduced for optimization purposes. Specifically, because we would like to avoid both underfitting and overfitting the datasets, we want our algorithm to be somewhat “cautious” and “lazy”. We do so by minimizing the influence of every input data through rescaling. In our case, we will make sure that all the input data are rescaled so that their maximum values is equal to `0.0001` (note that this number is suited for implementations in which the super cell is initialized with non-negative integers ranging from 0 to $\tilde{3}00$).

main.py

```
  #------------------
  #Normalization of the input vector
  #------------------
  inputs = 0.0001*inputs/max(inputs)
  vector = inputs.tolist()
```

For debugging purposes, we use the function `debug_time.set` (section 3.2) to display the iteration of the loop and the type of the analyzed data in the standard output.

main.py

```
  #------------------
  #The algorithm
  #------------------
  debug_time.set("Learning data labeled as " + ",
".join(map(str,label)))
```

We can now proceed to the main part of our code, in which we use of the method `intcyt` (section 5.1). Before calling this function, we will need to fix a few parameters, which we are going to discuss in the following paragraphs.

The parameters that we are about to set up will help us control the various intermediate steps that constitute our algorithm. Let us recall from the introduction of [1, Sec. 2] that our algorithm is made of four main steps, shown in red, in the following diagram.

(2.1)

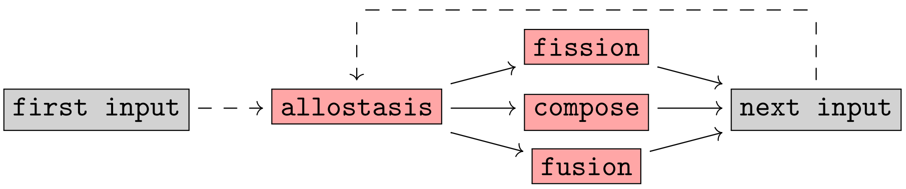

Every input given to the algorithm goes through the step `allostasis` (see diagram above), but depending on the iteration at which the algorithm is, the algorithm can proceed to either the step `fusion`, the step `fission` or the step `compose`. After one of these steps, the algorithm switches to the next input and the algorithm restarts the same process from `allostasis`. Here, it is important to note that the step `next input` does not necessarily refer to the data coming next in the dataset. Indeed, there are instances of our algorithm in which the next input is actually equal to the same data. The number of iterations done on the same data will be determined by an `epoch` number. This number determines how many times our algorithm needs to go through the loop shown in (2.1), with the same input data, before being able to switch to the next data in the dataset. The interested reader can find more details about the reasons why the algorithm is decomposed as shown in (2.1) in [1, Sec. 2].

For this tutorial, we will focus on a version of `IntCyt` in which:

- the `epoch` number is equal to 4, which means that there are four epochs before proceeding to the next iteration of the outer loop;
- during the first 4 iterations of the outer loop (*i.e.* a total 16 epochs): we go through the step `allostatis` with no extra step `fission`, `fusion`, or `compose`, as shown below for the first iteration;

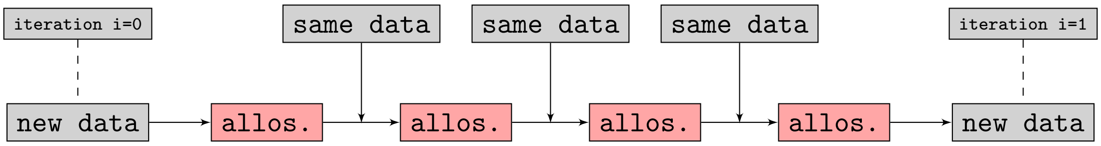

- after the fourth iteration: the step `fission` occurs at the first epoch;
- after the fourth iteration: the step `fusion` occurs at the third epoch;
- after the fourth iteration: the step `compose` occurs at the second and fourth epochs;

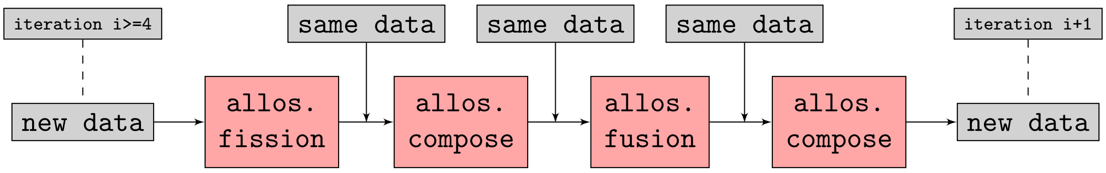

In the piece of code given below, the variable `start` specifies the iteration number at which the steps `fission`, `fusion`, and `compose` will start being used by our algorithm. The variable `epoch` contains the epoch number. The variables `fission_events`, `fussion_events` an `compose_events` contain the epochs at which the events `fission`, `fusion`, and `compose` occur within the inner loop (made of 4 epochs). Finally, the variable `events` contains the list of all these parameters and will later be given to the function `intcyt` as an argument.

main.py
```
#------------------
start = 4
epoch = 4
fission_events = [0]
fusion_events = [2]
compose_events = [1,3]
events = [start,epoch,fission_events,fusion_events,compose_events]
```

We also create a `list` item `filtering` that plays the role of the parameters $\nu$ and $\omega$ used in [1, Def. 2.30]. These parameters allow us to control fusion events and fission events occurring in the super cell by limiting these events to particular groups of cells. Giving high values to `filtering` (*e.g* values greater than `1` for the first two parameters and values above 5% of the value `dim` for the last one) will prevent cells that are too different from merging and will prevent cells that are similar enough from dividing. In our case, we chose the following parameterization.

main.py
```
#------------------
filtering = [1.5, #To control fission
             1.5, #To control fusion
             0]   #To control fusion and fission
```

We are now ready to code the inner loop. We start the loop by displaying, on the standard output, the state of the super cell `sc` by using the method `.stdout`. This method displays an ASCII tree that shows us about the architecture of super cell at the current iteration.

main.py
```
#------------------
for k in range(epoch):
  debug_time.set("TREE")
  sc.stdout(vector)
```

Before calling `intcyt`, we are going to set up a collection of parameters that will control the learning rate of our algorithm. These parameters depend on the type of data contained in our dataset. From the point of view of [1], these parameters help us define what is called the "gamma parameter", defined in [1, Sec. 2.3]. Recall that the formula for the gamma parameter is of the following form:

$$\gamma(c,a,j,u) = \begin{cases} 0 & \text{if "learning should not occur according to specific criteria";} \\ \text{see [1, Sec. 2.3]} & \text{otherwise.} \end{cases} \tag{2.2}$$

In what follows, the collection of parameters that we define allows us to give a meaning to the sentence "learning should not occur according to specific criteria". The idea is to take advantage of the interpretability of the data memorized by the cells of the super cell to determine whether these cells have learned any information from the data. If information has been learned, then we only want to update the content of the corresponding cells if the input is similar to their content and has a chance to make the memorized information more robust (see the diagram below).

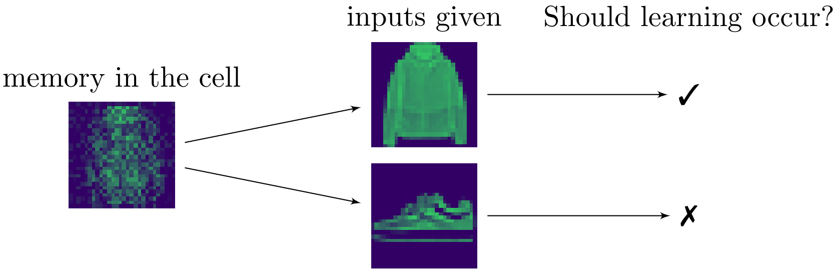

Because our data represent images, we employ a relatively simple approach, which measures the amount of contrast in the data. A strong contrast is likely to be linked to an abstraction of a concept rather than random data (as will be seen below).

To measure contrast in our data, we define three variables: `brightness`, `profiles` and `scores`. The variable `brightness` contains a list of brightness levels: the higher, the brighter. In our case, we will consider 5 levels of brightness, each expressed in percent with respect to the maximum value contained in the data: 10%, 25%, 50%, 75% and 90%. For instance, a random input tends to approximately have

- ▷ 90% of its pixels above 10% of the value of its brightest pixel;
- ▷ 25% of its pixels above 75% of the value of its brightest pixel;
- ▷ 50% of its pixels above 50% of the value of its brightest pixel;
- ▷ 75% of its pixels above 25% of the value of its brightest pixel;
- ▷ 10% of its pixels above 90% of the value of its brightest pixel.

In other words, for a random image, the rule of thumb is that $(100 - x)\%$ of the pixels is greater than or equal to $x\%$ of the value of its brightest pixel. For non-random images, the former percentage tend to be decreased. For instance, in the case of `MNIST`, we were able to notice that each data satisfied the following profile (on average):

- ▷ under 17.5% of its pixels were above 10% of the value of its brightest pixel;
- ▷ under 15.6% of its pixels were above 75% of the value of its brightest pixel;
- ▷ under 13.3% of its pixels were above 50% of the value of its brightest pixel;
- ▷ under 10.6% of its pixels were above 25% of the value of its brightest pixel;
- ▷ under 8.8% of its pixels were above 90% of the value of its brightest pixel.

In our implementation, we will define various contrast profiles corresponding to various learning stages (*e.g.* student and expert). These stages will be specified in terms of pairs of lower bounds and upper bounds and will be stored in the list `profiles`. For instance, a relation `profiles[i] = (0,.175)` means that the percentage of pixels above `brightness[i]` percent of the brightest pixel value should be between 0% and 17.5%.

In order to know whether it is reasonable to update the memory of a cell while it is learning, we use agreement levels. Here, we define the agreement of two vectors, say $v$ and $w$, as the cosine of their angular distance. Alternatively, the agreement of $v$ and $w$ can be computed as the normalized scalar product of $v$ and $w$, as shown below.

$$\mathsf{argeement}(v,w) = \frac{v \cdot w}{\|v\|\|w\|} = \frac{\sum v_i w_i}{\|v\|\|w\|} = \cos(\theta)$$

Because, in our case, the components of $v$ and $w$ are non-negative, the agreement of $v$ and $w$ is a value between 0 and 1. Intuitively, this quantity can be seen as measuring a correlation between the two vectors $v$ and $w$.

In the context of our algorithm, we only want to update an organelle of a cell if the agreement score of the organelle with the input is high enough for a given contrast profile.

As shown in the following diagram, this extends our filtering procedure to a two-step check.

| | | |
|---|---|---|
| `contrast`(`organelle`) passes the threshold ✓ | $\longrightarrow$ | is `argeement`(`organelle`, `input`) good enough ? |

To control the quality of agreements between organelles and inputs, we use the list `scores`. Specifically, this list will contain pairs of lower and upper bounds between which an agreement score needs to be in order for the update of the organelle to be authorized. Technically, this means that an organelle will be modified through a gradient descent method if the following property is satisfied:

$(P)$ : there exists an index `i` for which

1) the organelle satisfies the contrast profile `profiles[i]`;

2) the agreement of the organelle with the input is greater than or equal to `scores[i][0]` and less than or equal to `scores[i][1]`.

To conclude our discussion, the sentence "learning should not occur according to specific criteria" corresponds to the logical negation of the property $(P)$.

If we look at the formula of the gamma parameter given in [1, Sec. 2.3], we still need to define two more values $E$ and $F$, which appear in the case where $(P)$ is true, as shown below.

$$\gamma(c, a, j, u) = \begin{cases} 0 & \text{if } \mathsf{not}(P) \\ (\text{a formula depending on } E \text{ and } F) & \text{otherwise.} \end{cases}$$

While the value $E$ allows to increase the speed of the learning process, the value $F$ allows us to slow down the learning rate of the organelles that possess a relatively low agreement with the input. Note that, in our case, these values where determined heuristically.

Enough explanation and let us now proceed to the code! We start with dataset `SVHN`, for which our method does not work as well as with the datasets `MNIST` and `fashion-MNIST`, mainly because of the variability of RGB codes. A possible correction for the `SVHN` data would be to adapt our contrast profile method to these RGB codes.

main.py

```
    #------------------
    #brightness:  intensity levels (length m)
    #profiles:  intervals for intensity counts (length n*m)
    #scores:  agreement scores for a given profile (length n)
    if sys.argv[1] == "SVHN":
      #------------------
      brightness = [.1,.25,.5,.75,.9]
      profiles = [[(0,.6),(0,.375),(0,.15),(0,.05),(0,.005)]] #expert
      scores = [(.82,1)] #expert
      E = 13
      F = 25
```

Contrary to `SVHN`, the images stored in the `MNIST` dataset are bicolor. This makes the learning process of an image much more apprehensible for our contrast profiles. As a result, we were able to distinguish two learning stages in the evolution of the cells of IntCyt: a student phase and an expert phase. In the following lines of code, we can see that the expert profile has relatively low upper bounds.

main.py

```
    #------------------
    if sys.argv[1] == "MNIST":
      #------------------
      brightness = [.1,.25,.5,.75,.9]
      #the average profile for MNIST is [(0,.175), (0,.156), (0,.133), (0,.106), (0,.088)]
      profiles = [[(0,.6),(0,.25),(0,1),(0,1),(0,.1)], #student
                  [(0,.2),(0,.15),(0,.1),(0,.1),(0,.1)]] #expert
      scores = [(.7,1), #student
                (.8,1)] #expert
      E = 12.5
      F = 20
```

The `fashion-MNIST` dataset is very similar to the `MNIST` dataset, so it should not be surprising that we also have a student phase and an expert phase. However, the contrast profiles are a little bit different due to the nature of the data contained in the `fashion-MNIST` dataset, which involve greater amounts of bright pixels.

main.py

```
    #------------------
    if sys.argv[1] == "fashion-MNIST":
      #------------------
      brightness = [.1,.25,.5,.75,.9]
      profiles = [[(0,.7),(0,.5),(0,.3),(0,.1),(0,.01)], #student
                  [(0,.5),(0,.3),(0,.2),(0,.15),(0,.1)]] #expert
      scores = [(.5,1), #student
                (.8,1)] #expert
      E = 13.5
      F = 20
```

Before calling the function `intcyt`, let us extend the option `-load` of our launch command to two additional values `-load-selfsup-left` and `-load-selfsup-right`. These values will allow us to parameterize self-supervised challenges, as those shown in [1, Results]. Specifically, these options will instruct the script `main.py` to adapt the learning strategy of the gamma parameter to self-supervised challenges (this will be discussed in section 2.7 in more detail). Typical examples of such challenges would make the algorithm start with a cell whose organelles contain halves of images whose other halves are completed with white noise, as shown below. The goal of the challenge for our algorithm would then be to reconstruct the images entirely.

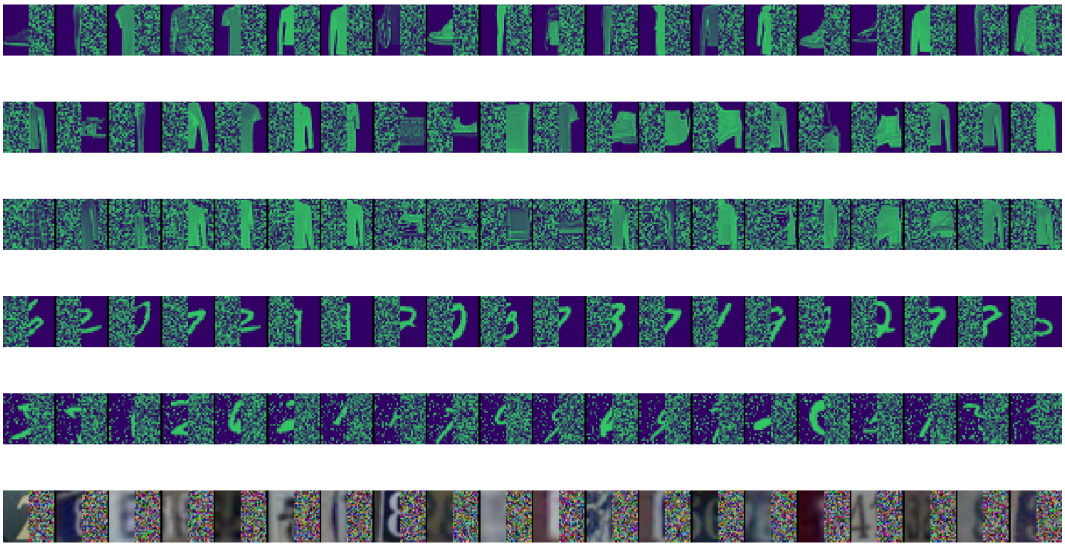

As will be seen, we will generate such images through the script `challenge.py` by calling one of the following options: `-left`, `-right`, `-left-noisy` or `-right-noisy`. The generated images are then saved in the directory `result-load` in a file named `load_initial.gz`.

Because, in such circumstances, the main goal for the algorithm is to be able to reconstruct the missing half of the image, one may want to not alter the side of the image that contains information during the learning (unless this is desired by the user). For this purpose, we add the two options `-load-selfsup-left` and `-load-selfsup-right` to our launch command in order to specify which of the sides of the image should not be modified during the learning phase. These two options create a list `selfsup` containing a parameterization of the area of the image that should not be altered. To do so, we add following piece of code to the file `main.py`.

main.py

```
    #-------------------
    selfsup = list()
    if len(sys.argv) > 2 and sys.argv[2] = "-load-selfsup-right":
      selfsup.append([image_size[1]/2, #middle of image
                      image_size[1]-1, #right of image
                      image_size[1]])  #width
    elif len(sys.argv) > 2 and sys.argv[2] = "-load-selfsup-left":
      selfsup.append([0,                 #left of image
                      image_size[1]/2-1, #middle of image
                      image_size[1]])    #width
```

Note that by not calling the options `-load-selfsup-left` or `-load-selfsup-right`, we let the variable `selfsup` be an empty list. As a result, the algorithm can alter both sides of the images during the learning process.

It is now time to use the function `intcyt` (section 5.1), which constitutes the heart of our learning algorithm. First, we create a variable `gamma_parameter`, which we use to store the output of the function `usf.gamma` (section 3.1.4) applied to our contrast profiles, as shown below. Then, we call the function `intcyt` on the variable `gamma_parameter` as well as the parameters defined before the inner loop.

main.py
```
    #-------------------
    gamma_parameter = usf.gamma(E,F,brightness,profiles,scores,*selfsup)
    intcyt(operad,sc,epoch*i+k,events,vector,gamma_parameter,filtering)
```

Before ending the outer loop, we want to save important parameters defining the state of the super cell, namely the root cell, the organelles and the tree structure of the super cell, in the files `save_roo.gz`, `save_org.gz` and `save_tre.gz`, respectively.

main.py
```
    #-------------------
    #The sate of the super cell is saved in the memory
    #-------------------
    #usf.print_root(sc,image_size,sys.stdout,option="display")
    #usf.print_organelles(sc,image_size,sys.stdout,option="display")
    #usf.print_tree(sc,image_size,sys.stdout,option="display")
    usf.print_root(sc,image_size,fsave_roo,option="save")
    usf.print_organelles(sc,image_size,fsave_org,option="save")
    usf.print_tree(sc,image_size,fsave_tre,option="save")
    fsave_roo.flush()
    fsave_org.flush()
    fsave_tre.flush()
```

We finish the code by closing any open file pointers, as shown below.

main.py
```
#-------------------
fsave_roo.close()
fsave_org.close()
fsave_tre.close()
if sys.argv[1] in ["MNIST","fashion-MNIST"]:
  fim.close()
  ftr.close()
```

We are now done for the code of `main.py`. To conclude this section, let us summarize the different ways in which this script can be called. If we formalize the various options of `main.py` through a formal grammar description, then the script `main.py` can be launched via the following command

```
> python main.py <dataset-name> <loading-option>
```

where we can replace the two tags with one of the following values:

```
<dataset-name>:
MNIST | fashion-MNIST | SVHN
<loading-option>:
-load | -load-selfsup-right | -load-selfsup-right | ε (empty)
```

Note that forgetting to specify the name of the dataset will exit the program.

#### 2.6. Visualizing the results

The goal of this section is to program a set of functions that will allow us to visualize the progress of our algorithm during its learning phase. More specifically, we will have three functions, which we will call to display different types of parameters characterizing the state of the associated super cell (see the displays shown below).

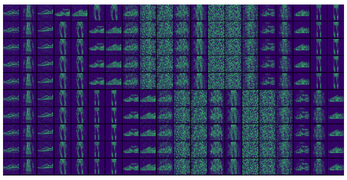
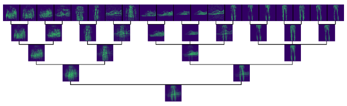
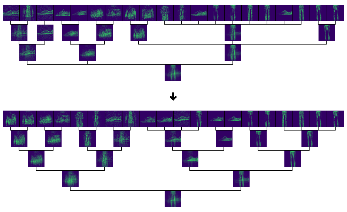

Our first function, `visualize`, will allow us to visualize the evolution of the root or the leaves of the super cell. Specifically, for a given dataset and a list of integers, call it `query`, the function `visualize` displays the collection of vectors stored in the dataset at the indices specified in `query`. Note that, in our case, the dataset will be either `save_roo.gz` or `save_org.gz`.

In the code given below, the variable `fload` is a file pointer reading data in either `save_roo.gz` or `save_org.gz`, the variable `batch` is the number of vectors supposed to be read and `image_size` corresponds to a list of integers describing the dimensions of the images encoded by the vectors of the dataset – the lists `image_size` correspond to the lists `image_size` used in section 2.8.5.

visualize.py

```
#------------------------------------------------------------------------
def visualize(fload,batch,query,image_size):
  memory = usf.get_memory(fload,batch,query)
  image = usf.make_rgb_grid(memory,image_size)
  fig, ax = plt.subplots(1)
  #remove axis
  ax.axis("off")
  #display the image
  ax.imshow(image)
  plt.show()
```

Our second function, `visualize_tree`, combines the output of `visualize` with the display of a tree representing the structure of the super cell associated with the greatest index contained in `query`. This time, the variable `fload` is supposed to read the file `save_tre.gz`. The function `visualize_tree` starts with a loop that collects the leaves of all but the last tree returned by the function `usf.get_trees` (see section 3.1.11) and stores the last tree in a variable `tree` (as shown below).

visualize.py

```
#------------------------------------------------------------------------
def visualize_tree(fload,query,image_size):
  trees = usf.get_trees(fload,query)
  memory = list()
  tree = list()
  for i in range(len(trees)):
    if i != len(trees)-1:
      memory.append(trees[i][0])
    else:
      tree = trees[i]
```

For clarity, we store the images of the leaves and the image of the tree in two different variables `image1` and `image2`, respectively.

visualize.py

```
  image1 = usf.make_rgb_grid(memory,image_size)
  image2 = usf.make_rgb_tree(tree,image_size)
```

We then concatenate the two image as one image and finish in the same way as in `visualize`, as shown below.

```
visualize.py
    image = image1 + image2
    fig, ax = plt.subplots(1)
    #remove axis
    ax.axis("off")
    #display the image
    ax.imshow(image)
    plt.show()
```

Last but not least, our third function, `visualize_forest`, displays the tree structures of all the super cells associated with the indices of the list `query`. The code of the function `visualize_forest` is very similar to that of `visualize_tree`.

```
visualize.py
#---------------------------------------------------------------------
def visualize_forest(fload,query,image_size):
    trees = usf.get_trees(fload,query)
    image = list()
    for i in range(len(trees)):
        image = image + usf.make_rgb_tree(trees[i],image_size)
        if i != len(trees)-1:
            image = image + usf.make_arrow_panel(panel_width=len(image[0]),
length=6)
    fig, ax = plt.subplots(1)
    #remove axis
    ax.axis("off")
    #display the image
    ax.imshow(image)
    plt.show()
```

As in `main.py`, we want to equip the script `visualize.py` with a collection of options calling our three functions. Specifically, we shall define three option-tags, which will allow us to decide which of functions `visualize`, `visualize_tree` and `visualize_forest` we want to call and set up the arguments needed for these functions.

The launch command for `visualize.py` will be of the following form:

```
> python visualize.py <dataset-name> <display-option> <query-intervals>
```

where we can replace the three tags with one of the following values:

`<dataset-name>:`
`MNIST | fashion-MNIST | SVHN`
`<display-option>:`
`-forest | -tree |` $\varepsilon$ *(empty)*
`<query-intervals>:`
$i_1$ $j_1$ $i_2$ $j_2$ ... $i_n$ $j_n$ *(a list of non-negative integers where* $i_k < j_k$*)*

Because the tag `<display-option>` can be empty, the first value of `<query-intervals>` can be contained by either `sys.argv[2]` or `sys.argv[3]`. As a result, we will have to use an index re-adjustment `r` to collect the potential arguments $i_k$ and $j_k$ so that these are contained in the system variables `sys.argv[2*(k-1)+r]` and `sys.argv[2*(k-1)+r+1]`. The value stored in `r` is determined as follows.

visualize.py

```
#------------------------------------------------------------------------
if sys.argv[2][0] != "-":
  r = 2
else:
  r = 3
```

An example of a launch command for `visualize.py` is given below.

```
> python visualize.py fashion-MNIST -forest 200 203 50 51 100 101 0 1
```

As can be seen above, each pair of integer is made of successive integers. However, the pairs need not be given in an increasing order. In the piece of code given below, we use the list of integers associated with the tag `<query-intervals>` to construct the list `query` required by our three functions. In the previous example, the list `query` would be of the form `[200,201,202,50,100,0]`.

visualize.py

```
#------------------------------------------------------------------------
query = list()
for i in range((len(sys.argv)-2)/2):
  query = query + range(int(sys.argv[2*i+r]),int(sys.argv[2*i+r+1]))
```

We now take care of the value of the tag `<dataset-name>`, through which we parametrize three variables `image_size`, `batch_roo` and `batch_org`. As shown above, the values of `<dataset-name>` are the name of the different datasets on which we will run our algorithm.

First, calling the option `SVHN` will set up the following parameters.

visualize.py

```
#------------------------------------------------------------------------
#To visualize learning from SVHN dataset
if sys.argv[1] == "SVHN":
  image_size = [32,32,3] #rgb
  batch_roo = 1
  batch_org = 20
```

Alternatively, we use the same parameters for either options `MNIST` and `fashion-MNIST`.

visualize.py

```
#------------------------------------------------------------------------
#To visualize learning from MNIST dataset
if sys.argv[1] in ["MNIST","fashion-MNIST"]:
  image_size = [28,28,1] #bicolor
  batch_roo = 1
  batch_org = 20
```

Finally, we take care of the values of the tag `<display-option>` as follows. For no value given, we apply the function `visualize` to the data stored in `save_roo.gz` and `save_org.gz`. The result of such an operation displays the roots and the organelles of the super cell associated with the iteration indices stored in the list `query`.

visualize.py
```python
#-----------------------------------------------------------------------
if sys.argv[2][0] != "-":
  fload_roo = gzip.open("save_roo.gz","r")
  fload_org = gzip.open("save_org.gz","r")
  #--------------
  visualize(fload_roo,batch_roo,query,image_size)
  visualize(fload_org,batch_org,query,image_size)
  #--------------
  fload_roo.close()
  fload_org.close()
```

If the option `-tree` is given, then we display the leaves of all the super cells associated with the iteration indices stored in the list `query`, together with the tree structure of the last super cell.

visualize.py
```python
#-----------------------------------------------------------------------
elif sys.argv[2] == "-tree":
  fload_tre = gzip.open("save_tre.gz","r")
  #--------------
  visualize_tree(fload_tre,query,image_size)
  #--------------
  fload_tre.close()
```

If the option `-forest` is given, then we display the tree structures of all the super cells associated with the iteration indices stored in the list `query`.

visualize.py
```python
#-----------------------------------------------------------------------
elif sys.argv[2] == "-forest":
  fload_tre = gzip.open("save_tre.gz","r")
  #--------------
  visualize_forest(fload_tre,query,image_size)
  #--------------
  fload_tre.close()
#-----------------------------------------------------------------------
```

This finalizes the code of the file `visualize.py`. Before moving on to the code of our next script, `challenge.py`, we conclude this section by showing examples of outputs returned by the scripts `main.py` and `visualize.py`.

We begin with the dataset SVHN. Because we do not have any initialization for this dataset, we generate one through the script `main.py`. We start with the following command.

```
> python main.py SVHN
```

Right after taping the carriage return key ↩ on the keyboard, you should see, in the standard output, a display similar to the one shown below. The label `[Allostasis 0]` shown towards the end of the display indicates that the algorithm is currently at cycle 0.

```
[Learning data labeled as 1]

[TREE]
[0] -> Cell[0]oooooooooooooooooooo [0]:0.7978860358 [1]:0.7938557942
[2]:0.8118207372 [3]:0.8014380039 [4]:0.8095886724 [5]:0.8055258661
[6]:0.8026621764 [7]:0.8046829853 [8]:0.7997212603 [9]:0.8068208694
[10]:0.7960182403 [11]:0.8028378799 [12]:0.7931610043 [13]:0.7953383775
[14]:0.8070709252 [15]:0.7935919636 [16]:0.8007175106 [17]:0.8096277577
[18]:0.7965840558 [19]:0.801800826

[Allostasis 0]
[brightness] 0.8675 0.7382 0.4902 0.2587 0.096
False
```

After some time, the algorithm will reach the label `[Allostasis 1]`. Form then on, we can start to visualize the past states of the super cell. For instance, to visualize the initialization of the super cell, we can launch the scrip `visualize.py` as follows.

```
> python visualize.py SVHN -tree 0 1
```

This has the result of creating a `matplotlib` window containing the right-hand side of the display shown below (see below for more explanation). Note that the image given below is more detailed than the output produced by our script and corresponds to the output of a more sophisticated version of the file `visualize.py`, which can be found in the directory `software`.

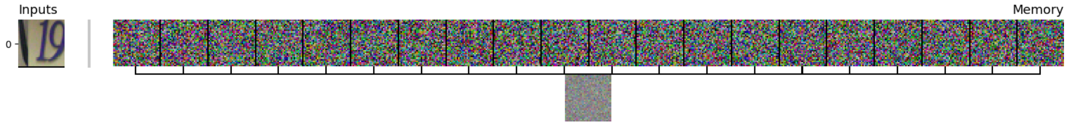

The left-hand side of the previous display shows the input taken by our algorithm on its first learning cycle. The right-hand side shows the output of our command, which corresponds to the memory of the underlying super cell prior to taking the input shown on the left. The memory of the super cell is organized as follows: the collection of images arranged in a row represents the information contained by all the organelles associated with the super cell (which here turns out to be a cell), while the image shown underneath the row represents the vector sum of all the organelles (which corresponds to the *content* of the cell, as defined in [1, Def. 1.21]).

After the label `[Allostasis 200]` has passed, we can call the following command.

```
> python visualize.py SVHN -tree 0 1 200 201
```

The output associated with the previous command corresponds to the grid of images shown below, on right-hand side, where the top row shows the organelles of the initial cell, the second row shows the organelles of the super cell at cycle 200 and the tree, below it, shows the state and structure of the super cell at cycle 200.

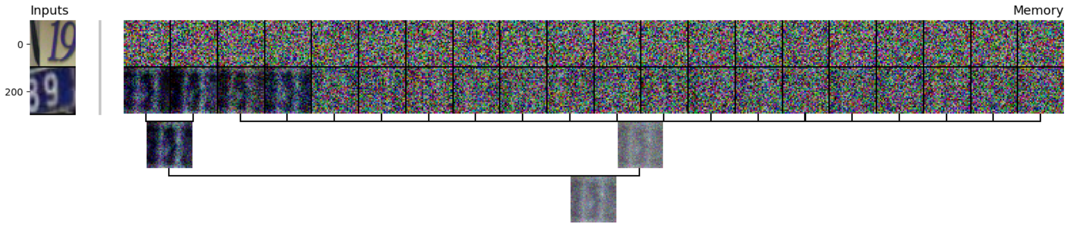

Despite the presence of learned concepts in the leftmost leaves of the previous tree, our parameterization for `SVHN` is not optimal and needs to be adapted to a RGB code colors. This becomes more obvious later during the learning phase, where the concepts learned by `IntCyt` remains hardly indistinguishable numbers. For instance, if we wait until cycle 1159 and use the following command, where we replace the option `-tree` with the option `-forest`, then our output looks as shown after the command.

```
> python visualize.py SVHN -forest 0 1 200 201 1159 1160
```

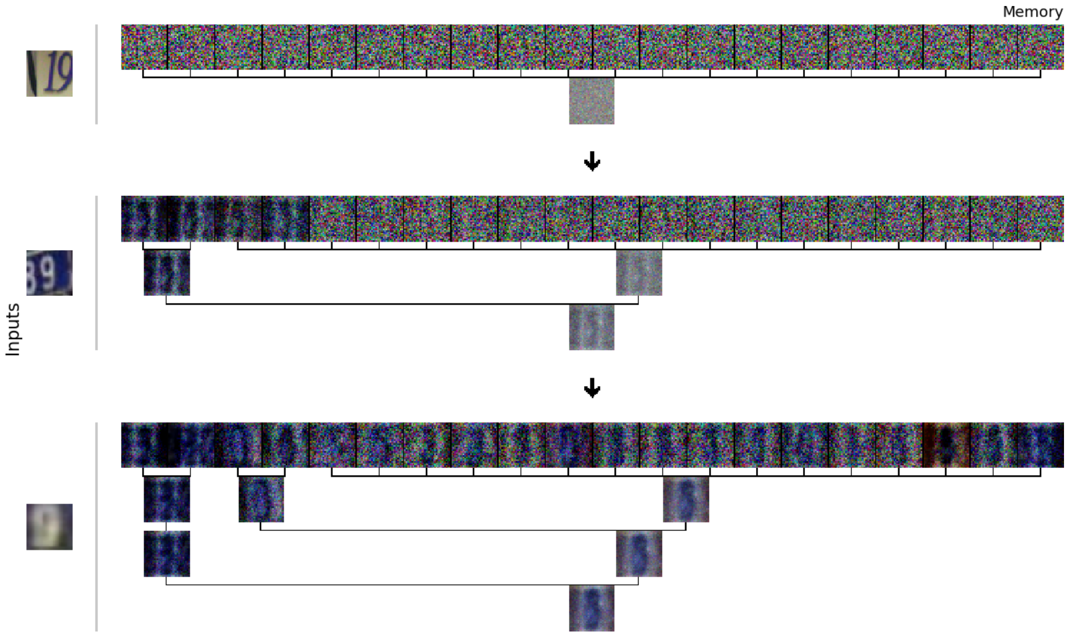

As can be seen, the concepts learned by the super cell are still quite blurry. However, to put things in context, the dataset `SVHN` contains a lot of blurry images itself. This tells us more about what `intcyt` does: it tries to construct abstractions of the concepts contained in a dataset at a quality level that approximates that of the data contained in the dataset.

Note that the organelles of the super cells (shown above) all seem to describe the same abstraction. Such a limited learning can be explained by the current parametrization of the function `usf.gamma` (section 3.1.4), which is not adapted to RGB code colors. The interested reader is invited look into the code of the module `useful.py` to experiment better learning strategies for `SVHN`.

From now on, we shall focus on our two bicolor datasets, which is to say `MNIST` and `fashion-MNIST`. We start with the dataset `MNIST` and launch the script `main.py` with the following command.

```
> python main.py MNIST
```

After a few hundreds cycles, say 2001, we can call the following command to visualize the shape of the super cell at cycle 2000 and the evolution of its organelles.

```
> python visualize.py MNIST -tree 0 1 100 101 200 201 300 301 400 401 500
501 600 601 700 701 800 801 900 901 1000 1001 1100 1101 1200 1201 1300 1301
1400 1401 1500 1501 1700 1701 1900 1901 2000 2001
```

In my case, the returned output looks as shown below, on the right-hand side (as usual, the inputs taken by the algorithm are specified on the left-hand side of each row).

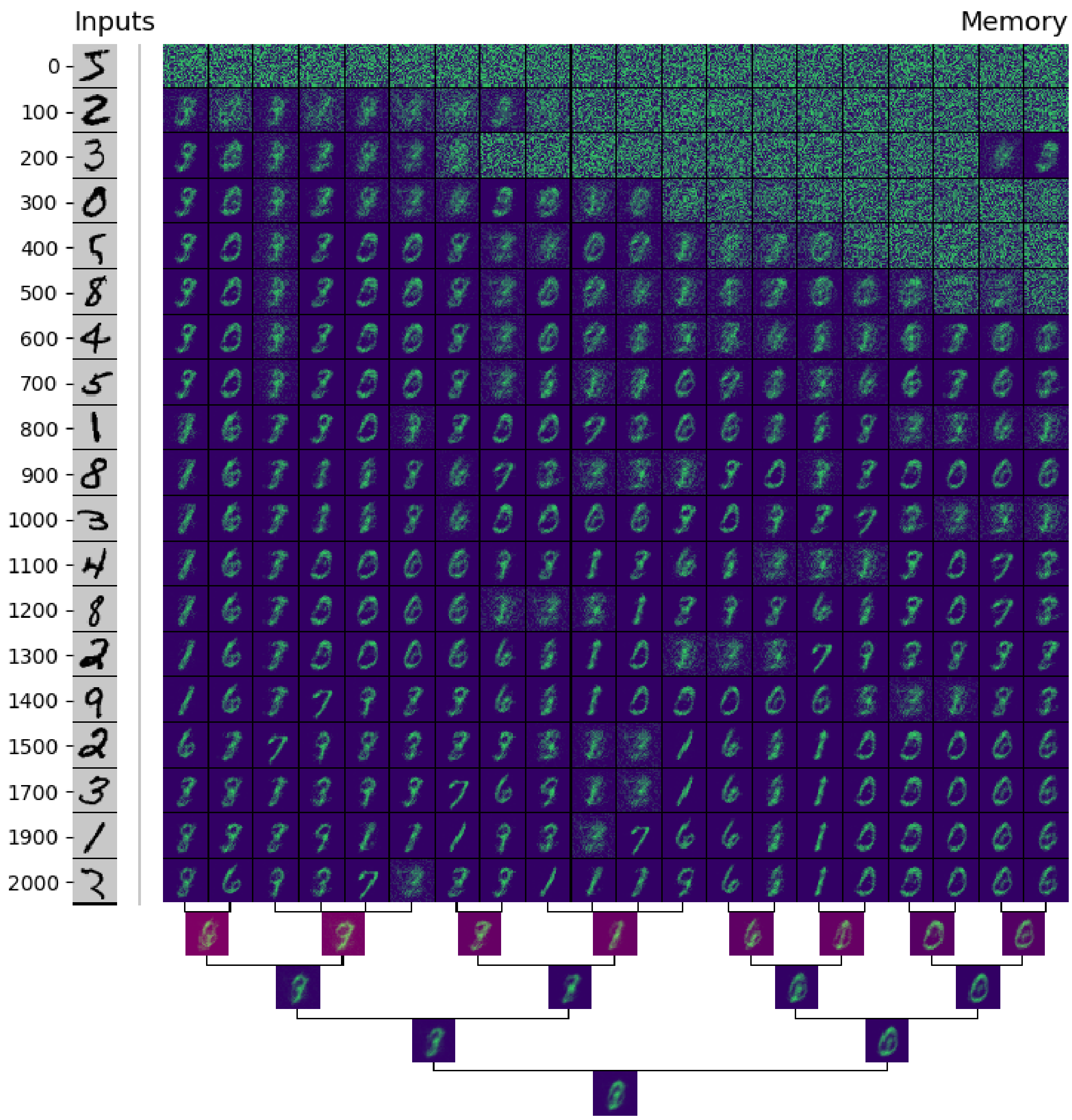

Interestingly, we can see that the algorithm first focuses on separating the information from the white noise, in the first cycles, and then moves on to gathering together pieces of information that look similar. The reddish (or purple) color used on the junctions of the tree displayed at the bottom indicates a high learning activity at these junctions.

Alternatively, we can call the following command to see how the architecture of the super cell evolved during the learning phase.

```
> python visualize.py MNIST -forest 0 1 1000 1001 2000 2001
```

In my case (see the picture displayed below), we can see that the architecture of the super cell evolved quite significantly throughout the learning phase. In particular, at cycle 2001, the super cell may have still been in the process of evolving, meaning that the super cell could end up being more organized than the one shown below.

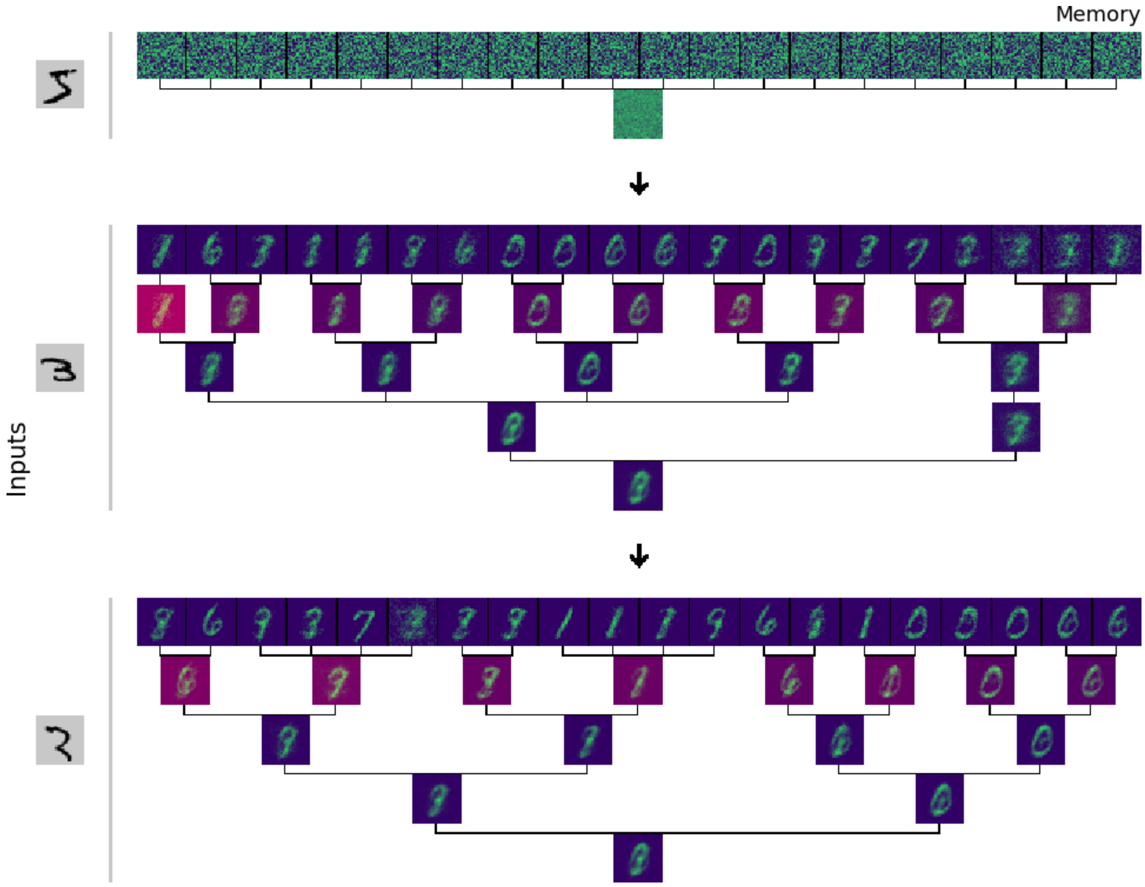

We proceed similarly for the dataset `fashion-MNIST`. We start by calling the script `main.py` as follows.

```
> python main.py fashion-MNIST
```

After a few hundred cycles, we can we call the script `visualize.py`. For instance, we can use the following command, which, in my case, gives the display shown below, after the command.

```
> python visualize.py fashion-MNIST -forest 0 1 400 401 938 939
```

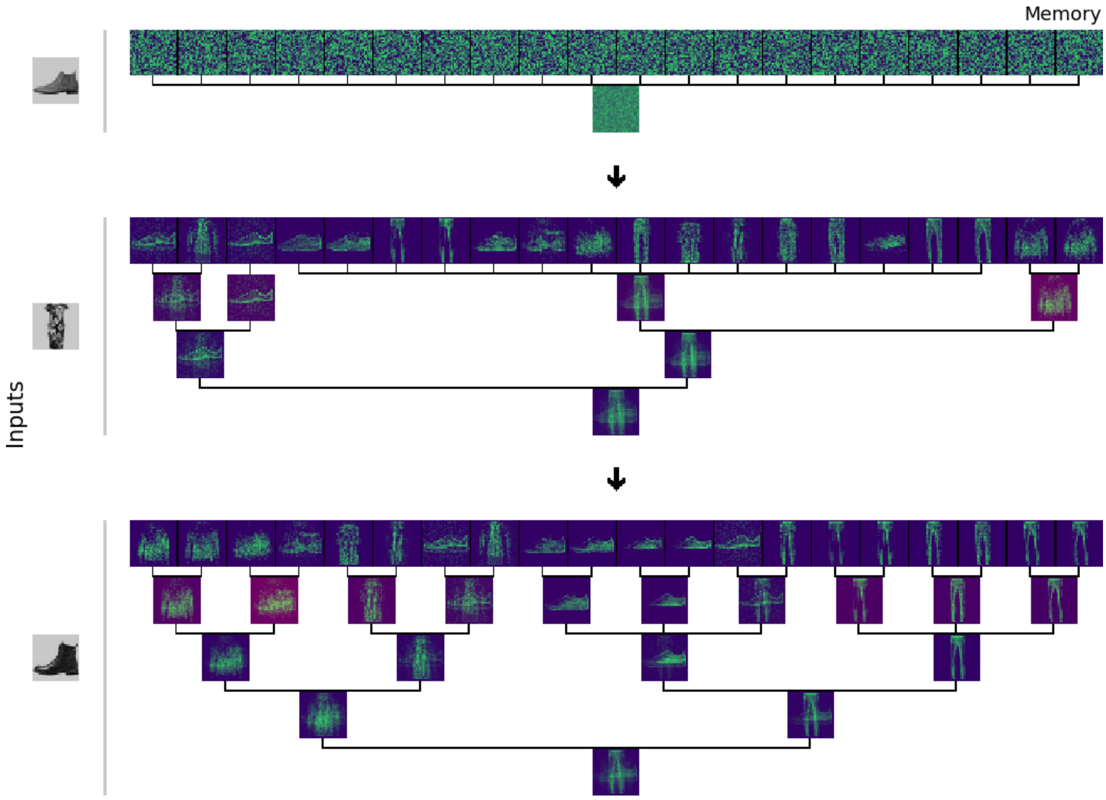

In the following section, we will see how our script `main.py` performs on initializations that are not everywhere random and contains partial information regarding the dataset.

#### 2.7. Self-supervision

The goal of the present section is to write a script that will generate semi-random initializations containing halves of images from our datasets. Given the learning and abstraction abilities seen in section 2.6, we expect our algorithm to be able to reconstruct the missing halves. We will have 5 types of challenges, which we will call through the following options:

- `-right`: hides the left part of an image with white noise;

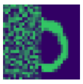

- `-right-noisy`: hides the left part of an image with white noise and fills the background of the image with similar white noise;

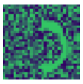

- `-left`: hides the right part of an image with white noise;
- `-left-noisy`: hides the right part of an image with white noise and fills the background of the image with similar white noise;
- $\varepsilon$ (`empty`): takes the full image;

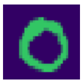

The code of our script will be saved in the file `challenge.py` and will be very similar to that of `main.py`. As in `main.py`, we start by opening the dataset files, which we can control by passing the options `MNIST`, `fashion-MNIST`, or `SVHN` to the launch command of `challenge.py`.

challenge.py

```
#-----------------------------------------------------------------------
#Loading the dataset for learning
#-----------------------------------------------------------------------
#Open the SVHN dataset
if len(sys.argv) > 1 and sys.argv[1] == "SVHN":
  dic_svhn = sio.loadmat("data/train_32x32.mat")
  #Access to the dictionary
  dicim = dic_svhn["X"] #shape:  32, 32, 3, 73257
  dictr = dic_svhn["y"] #shape:  73257, 1
  image_size = [32,32,3] #height, width, depth
#------------------
#Open the MNIST dataset
elif len(sys.argv) > 1 and sys.argv[1] == "MNIST":
  fim = gzip.open("data/train-images-idx3-ubyte.gz","r")
  ftr = gzip.open("data/train-labels-idx1-ubyte.gz","r")
  ftr.read(8)
  fim.read(16)
  image_size = [28,28,1] #height, width, depth
#------------------
#Open the fashion-MNIST dataset
elif len(sys.argv) > 1 and sys.argv[1] == "fashion-MNIST":
  fim = gzip.open("data/t10k-images-idx3-ubyte.gz","r")
  ftr = gzip.open("data/t10k-labels-idx1-ubyte.gz","r")
  ftr.read(8)
  fim.read(16)
  image_size = [28,28,1] #height, width, depth
  categories = ["t-shirt","trousers","pullover", \
                "dress","jacket","sandal","shirt", \
                "sneaker","bag","ankle-boot"]
else:
  exit(0)
```

We will also need three parameters, which we call `ary`, `dim` and `flatten_width`, to control the dimensions of our initialization. The parameter `dim` contains the "volume" of every image in a given dataset, the parameter `flatten_width` contains the multiplication between the width and the depth of the images of the dataset (as shown below)

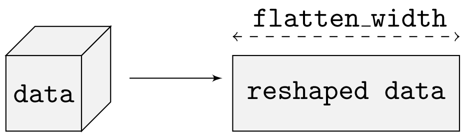

and `ary` is the number of organelles possessed by the super cell.

challenge.py
```
#-----------------------------------------------------------------------
#Parameters
#-----------------------------------------------------------------------
dim = image_size[0] * image_size[1] * image_size[2]
flatten_width = image_size[1] * image_size[2]
ary = 20
```

As previously said, we intend to initialize the organelles of our super cell with halves of images. These images will be picked randomly within the 1000 first images of a given dataset. We implement this random picks through the following loop, which creates the list `init` containing the indices of those images to be used in the initialization.

challenge.py
```
#-----------------------------------------------------------------------
#Randomly select data in the dataset.
#-----------------------------------------------------------------------
init = list()
while len(init) < ary:
  n = random.randint(0,999)
  if not(n in init):
    init.append(n)
init = sorted(init)
print init
```

We can classify our challenges in two groups: those that hide the left half of the images and those that hide the right half. Because we do not intend to alter the part of the image containing the information, we store, in a variable `random_init`, the position (modulo `flatten_width`) of the pixels that can be altered with white noise.

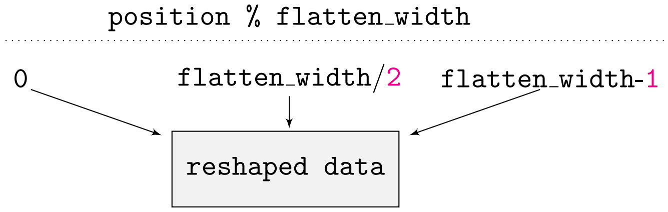

We do so by adding the following lines of code to `challenge.py`.

challenge.py
```
#-----------------------------------------------------------------------
random_init = list()
if len(sys.argv) > 2 and sys.argv[2][:6] == "-right":
  random_init.extend([0, flatten_width/2-1])
elif len(sys.argv) > 2 and sys.argv[2][:5] == "-left":
  random_init.extend([flatten_width/2, flatten_width-1])
#-----------------------------------------------------------------------
```

Before entering the main part of the code, we generate a file `load_initial.gz` in which we will store the initialization. We create it using the module `gzip` as follows.

challenge.py
```
#-----------------------------------------------------------------------
#Save the randomly picked data in a file
#-----------------------------------------------------------------------
fself = gzip.open("result-load/load_initial.gz","w")
```

We are now ready to collect the images whose indices belong to the list `init` and alter them according to the options specified in the launch command. This launch command will take the following form.

```
> python challenge.py <dataset-name> <initialization-option> <pair>
```

The different values for each of the tags is given below.

```
<dataset-name>:
MNIST | fashion-MNIST | SVHN
<initialization-option>:
-right | -left |
-right-noisy | -left-noisy
<pair>:
i j (a pair of non-negative integers) | ε (empty)
```

The tag `<pair>` is meant to refine the two options `-right-noisy` and `-left-noisy` by specifying the probability with which the white noise should be added to the half of the image that contains data. Specifically, if the tag `<pair>` is given two non-negative integers `i` and `j`, then the pixels of null intensity that are in the half containing the image will have a probability `i/j` to be completed with white noise, as shown below.

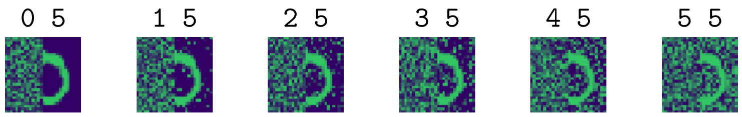

Let us now go through the code for collecting the images whose indices belong to the list `init`. To do so, we use a loop that goes through the first 1000 images of the datasets. The following piece of code is the same as the one we used in `main.py`.

challenge.py

```
for i in range(1000):
  #-------------------
  #Get labels and inputs
  #-------------------
  if sys.argv[1] == "SVHN":
    label = dictr[i]
    inputs = dicim[:,:,:,i].reshape(dim)
#-------------------
  if sys.argv[1] == "MNIST":
    buf_lab = ftr.read(1)
    buf_inp = fim.read(dim)
    label = np.frombuffer(buf_lab, dtype=np.uint8).astype(np.int64)
    inputs = np.frombuffer(buf_inp, dtype=np.uint8).astype(np.float32)
#-------------------
  if sys.argv[1] == "fashion-MNIST":
    buf_lab = ftr.read(1)
    buf_inp = fim.read(dim)
    label = map(lambda x:  categories[x], np.frombuffer(buf_lab, dtype=np.uint8).astype(np.int64))
    inputs = np.frombuffer(buf_inp, dtype=np.uint8).astype(np.float32)
```

The rest of the code is now dedicated to the situation in which the loop reads an image whose associated iteration index (in the loop) belongs to the list `init`. First, we display its corresponding label on the standard output by using the function `debug_time.set`.

challenge.py

```
#------------------
  if i in init:
    #------------------
    debug_time.set("Data picked:  " + ", ".join(map(str,label)))
    #------------------
```

Second, if the list `random_init` is non-empty, then we construct the initialization according to the specification given in the launch command, as shown below.

challenge.py

```
    if len(random_init) == 2:
      for k in range(len(inputs)):
        if random_init[0] <= k % flatten_width <= random_init[1]:
          inputs[k] = random.randint(0,250)
        elif len(sys.argv) > 2 and sys.argv[2] in ["-right-noisy",
"-left-noisy"] and inputs[k] == 0:
          if len(sys.argv) == 3:
            inputs[k] = random.randint(0,250)
          elif len(sys.argv) > 4:
            if 1 <= random.randint(0,int(sys.argv[4])) <=
int(sys.argv[3]):
            inputs[k] = random.randint(0,250)
```

We finish the code by displaying the initialization on the standard output and saving it in the file `load_initial.gz`.

challenge.py

```
    #------------------
    linputs = inputs.tolist()
    usf.print_data(linputs,image_size,sys.stdout,option = "display")
    usf.print_data(linputs,image_size,fself,option = "save")
    fself.write("\n")
#----------------------------------------------------------------------
fself.close()
if sys.argv[1] in ["MNIST","fashion-MNIST"]:
  fim.close()
  ftr.close()
```

The rest of this section is dedicated to showing examples of outputs of `main.py` relative to initializations generated by the script `challenge.py`.

We start with `MNIST`. First, we generate a self-supervised challenge made of left halves of images by calling the following command.

```
> python challenge.py MNIST -left
```

In my case, I obtain the following initialization:

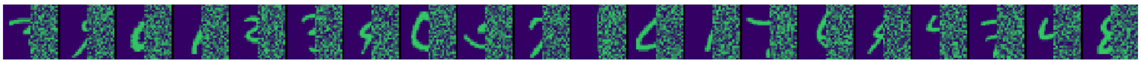

Before calling the script `main.py`, we shall change the contrast profile associated with `MNIST`. Indeed, note that the previous initialization contains valuable information about the data and it would makes sense to take advantage of this information to alter the halves of the images that contain white noise. As a result, our strategy will be to use the contrast profiles only on the left halves of the images. To do this properly, we want to adapt the contrast profiles associated with `MNIST` to these halves as best as possible. For the present challenge, we found that the following contrast profile work well.

main.py

```
    #------------------
    if sys.argv[1] == "MNIST":
      #------------------
      brightness = [.1,.25,.5,.75,.9]
      #the average profile for MNIST is [(0,.175), (0,.156), (0,.133), (0,.106), (0,.088)]
      profiles = [[(0,.45),(0,.25),(0,.1),(0,.05),(0,.01)], #student
                  [(0,.3),(0,.3),(0,.1),(0,.05),(0,.01)]] #expert
      scores = [(.7,1), #student
                (.8,1)] #expert
      E = 14
      F = 25
```

After making these changes in `main.py`, we start the learning phase through the following command, in which we use the option `-load-selfsup-left` to instruct the script to not alter the left halves of the images.

```
> python main.py MNIST -load-selfsup-left
```

After a few thousands cycles (in my case, 5001 cycles), I called the script `visualize.py` with the command shown below and I obtained the image shown after the command.

```
> python visualize.py MNIST -tree 0 1 500 501 1000 1001 1359 1360 4579 4580 5000 5001
```

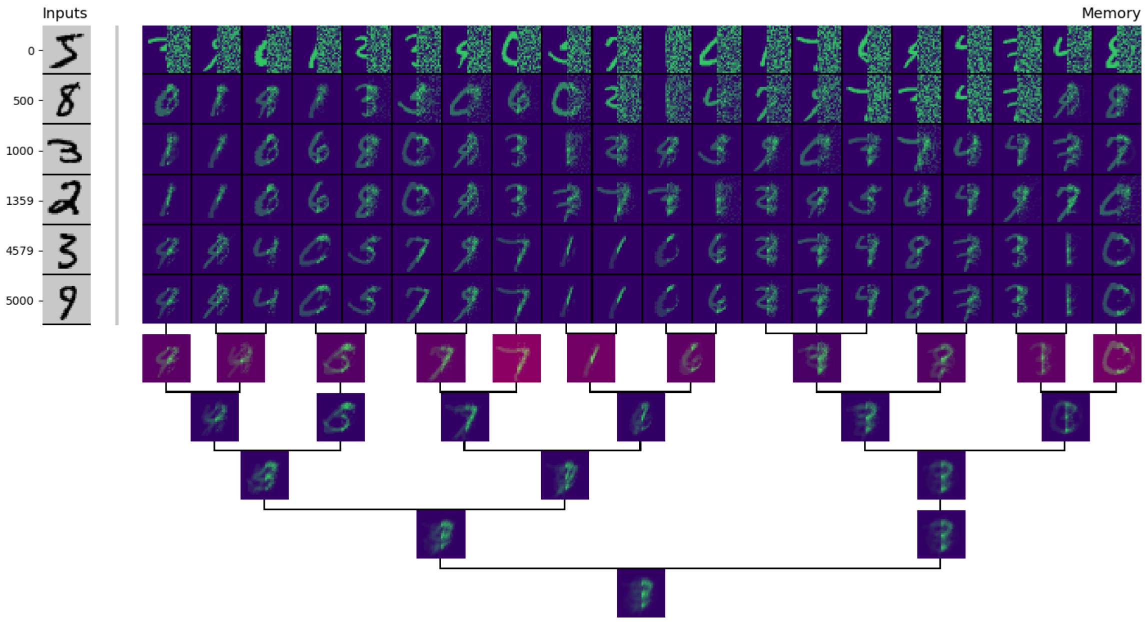

We can proceed similarly for `fashion-MNIST`. First, let us generate a new challenge as follows.

```
> python challenge.py fashion-MNIST -right
```

In my case, I obtain the following initialization.

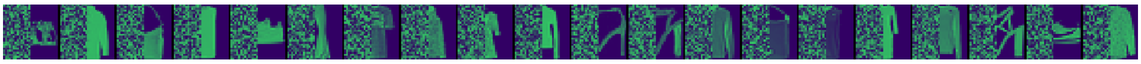

As with `MNIST`, we should also update the contrast profile of `fashion-MNIST`. For our current challenge, we apply the following changes.

main.py

```
        #-------------------
        brightness = [.1,.25,.5,.75,.9]
        profiles = [[(0,1),(0,1),(0,1),(0,1),(0,1)], #student
                    [(0,.7),(0,.6),(0,.5),(0,.4),(0,.3)]] #expert
        scores = [(.85,1), #student
                  (.95,1)] #expert
        E = 13.5
        F = 22.5
```

In addition, to prevent the compartmentalization of concepts too early, we change the code of `main.py` as follows:

main.py

```
    #-------------------
    step = i/float(iterations)
    interval = lambda s, a, b:  a*(1-s)+b*s if s < 1 else b
    #-------------------
```

main.py

```
    #-------------------
    filtering = [1.5, #To control fission
                 1.5, #To control fusion
                 interval(step,10,2)] #To control fission and fusion
```

After a few thousands cycles (in my case, 35000 cycles), I called the script `visualize.py` with the command shown below and I obtained the image displayed below, after the command.

```
python visualize.py fashion-MNIST -tree 0 1 5000 5001 10000 10001 20000
20001 30000 30001 35000 35001
```

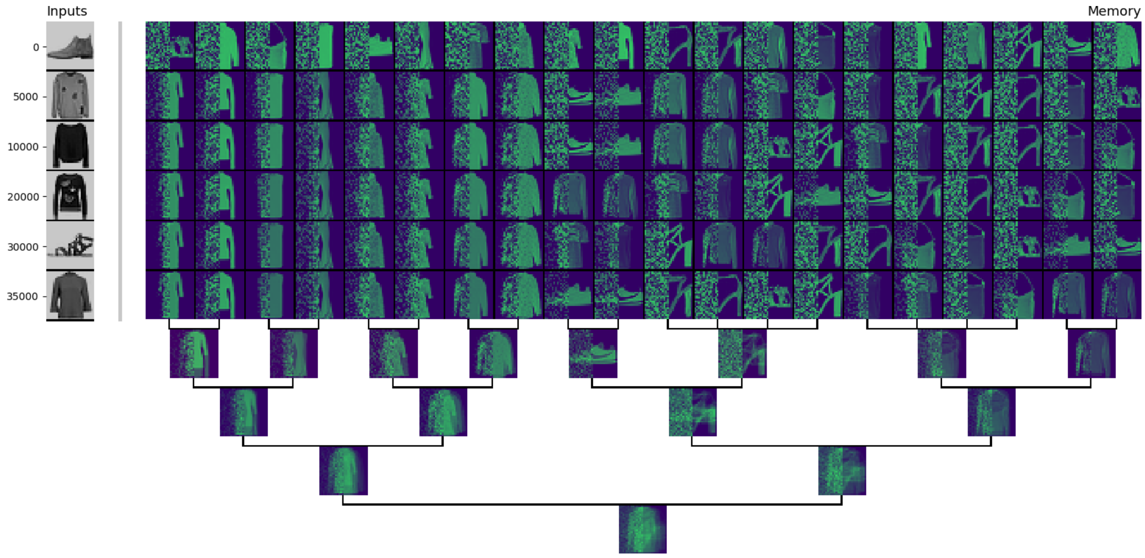

I will let the reader experiment on the other types of challenges. Note that this may require the design of new contrast profiles depending on the dataset chosen.

#### 2.8. Method for the DREAM3 challenge

The present section describes the method used in [2] to obtain the results of IntCyt (shown therein) on the `DREAM3` dataset. Recall that the `DREAM3` dataset can be represented as a differential gene expression matrix, as shown below, whose vertical dimension consists of 8 time points in 4 different strains of yeasts and whose horizontal dimension consists of 9335 genes, all shared by the four strains (in the image given below, a significant amount of gene expressions is not shown and is instead represented by the darker blue boxes, on the left and right).

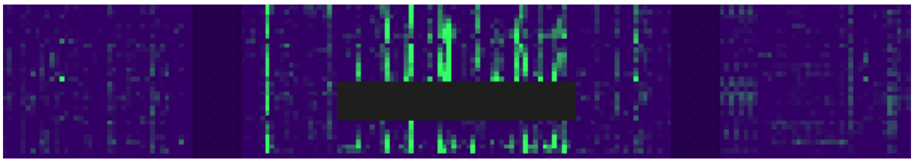

The four strains of yeast will be referred to as `wt` (short for *wild type*), and as `strain1`, `strain2`, and `strain3`. The last three strains are different from `wt` in that they underwent a deletion of one of their regulatory genes, therefore providing different overall gene expression profiles.

Among the 9335 genes, 50 genes are missing their expressions for `strain1` (represented by the dark box in the previous picture). In this section, we explain how we can use IntCyt to recover these genes expressions using the self-supervised learning approach described in section 2.7. In the following, we will call the 50 genes “hidden genes”, even though we know their expressions for three of the strains, and we will call the rest of the genes “known genes”.

**2.8.1. Principle of the method.** In this section, we present the principle of the method used in [2] to address the `DREAM3` challenge.

As shown in the schematic below, we use the expression of the 50 hidden genes as the initial self-supervised memory for IntCyt. Specifically, the missing expression are all attributed a value of 100, while the non-missing gene expression are all amplified by a factor of 100. On the other hand, the expression of the known genes are used as *training data* and stored in a file named `dream3_training.gz`, in the directory `data`. As with the self-supervised initialization of IntCyt, the gene expressions contained in `dream3_training.gz` are all amplified by a factor of 100 (compared to the original expressions).

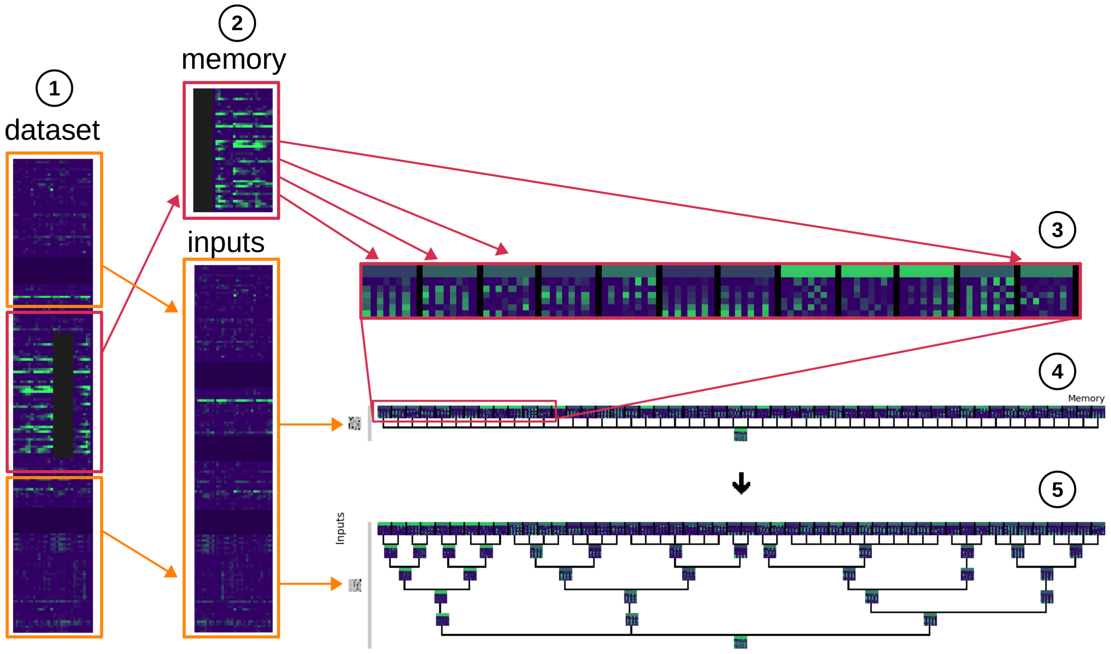

Also, because IntCyt only takes vectors of non-negative values, all the expression vectors are turned into vectors of non-negative values by doubling their dimensions, associating odd components with negative values and even components with non-negative values.

Once the initialization of IntCyt is set up and the training dataset `dream3_training.gz` is prepared, our strategy is to go through the training dataset three times with different agreement and brightness criteria to filter different levels of information.

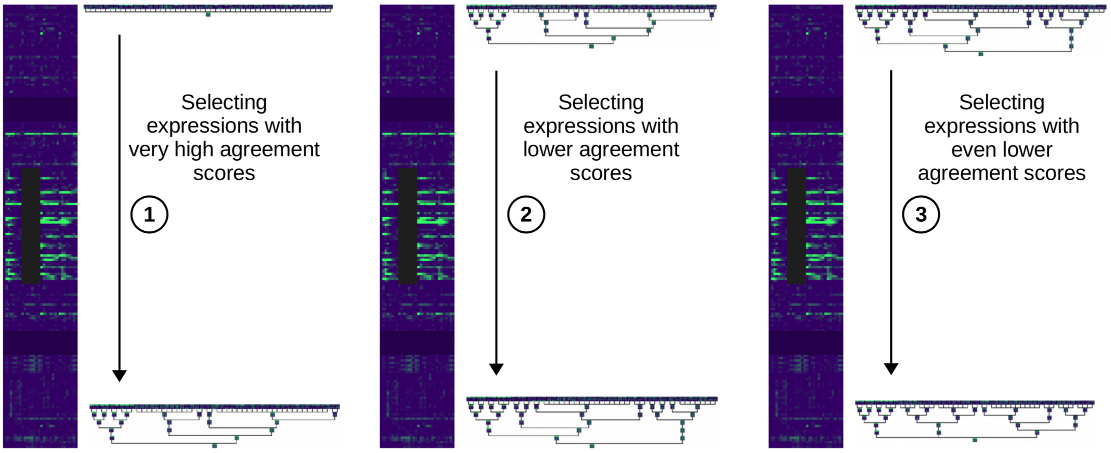

To be more specific, our algorithm will go as follows: during a first run through the dataset `dream3_training.gz`, IntCyt will take care of only using gene expressions that possess relatively high agreements scores with, at least, one of the hidden gene expressions. Note that these levels of agreement will be defined through collection of thresholds, which we will discuss in section 2.8.4.

After the first run, a certain proportion of the organelles will likely show, at the missing gene expression components, differentiated values that may capture the true missing expression profiles. On the other hand, some of the hidden gene may still be associated with non-differentiated missing gene expressions. These expressions need to be updated by using less selective criteria during a second run. Ideally, this second run should not alter the expressions that have been learned during the first run. To prevent these potential alterations to happen, we will use brightness levels (see section 2.8.5) to prevent the update of already-learned gene expressions. In our case, we will do three runs with less and less selective agreement thresholds together with brightness levels that prevent the update of sufficiently-differentiated missing gene expressions.

As will be seen later in the method, the significance associated with the prediction accuracy of our algorithm will take either one of the following forms:

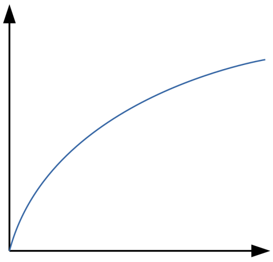

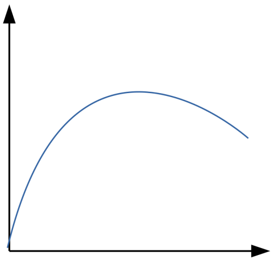

The left graph suggests a scenario in which the data selected by our algorithm has mainly contributed to the overall recovering of the true missing expression. On the other hand, the right graph suggests a scenario in which the data selected by our algorithm contained expression profiles that were too far from the true missing expressions. In section 2.8.4, we will discuss a general strategy to find selective criteria that are conducive to the first type of scenarios.

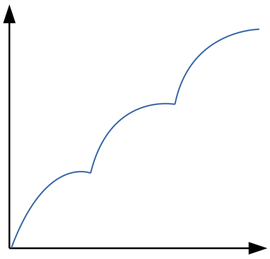

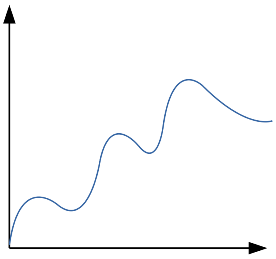

Throughout consecutive runs of our algorithm on the dataset `dream3_training.gz` (for which we expect the accuracy significance to be as shown above, on the left), we will be able to impute the missing expressions associated with the hidden genes. The obtained gene expressions will then be ranked and compared to a golden standard (GS) to assess the prediction accuracy of our algoirthm, as discussed earlier.

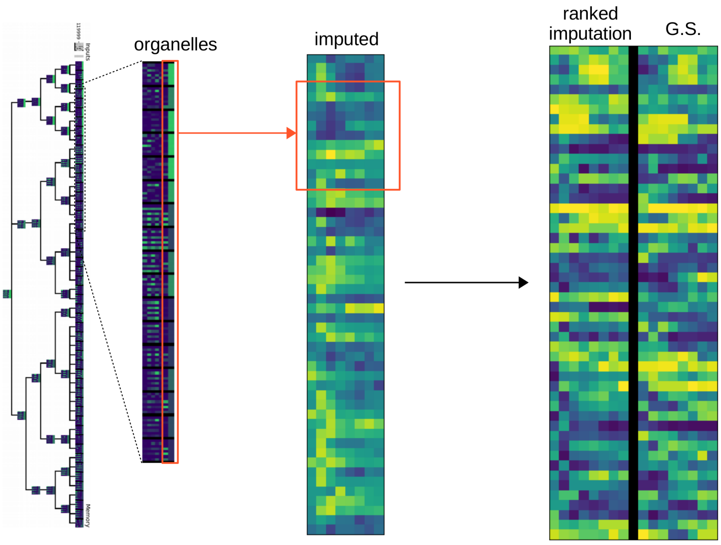

The comparison between ranked imputation and the golden standard is done through two types of $p$-value-based scores. Specifically, we assess the accuracy of our result

1) through the geometric mean $p_{\mathsf{col}}$ of the $p$-values resulting from the Spearman correlations between every column of same index in the ranked imputation and in the golden standard.

2) through the geometric mean $p_{\mathsf{row}}$ of the $p$-values resulting from the Spearman correlations between every row of same index in the ranked imputation and in the golden standard.

We can combine these two $p$-values into an accuracy score given by the formula

$$s = -\frac{1}{2}\mathsf{log}(p_{\mathsf{col}} \cdot p_{\mathsf{row}})$$

At the time of the `DREAM3` challenge, the best scores obtained by the participants of the challenge are shown in the following table [4], in which we also display the score obtained by IntCyt.

| Team | Score | Gene accuracy | Gene p-value | Time accuracy | Time p-value |
|---|---|---|---|---|---|
| **IntCyt** | **3.57** | **0.576** | **2.4E-06** | **0.489** | **2.9E-02** |
| GH | **3.25** | 0.563 | 6.5E-06 | 0.512 | 4.8E-02 |
| KNN | 3.18 | 0.558 | 1.1E-05 | **0.533** | 3.9E-02 |
| Team 263 | 1.85 | 0.421 | 7.5E-03 | 0.112 | 2.7E-01 |
| Team 297 | 1.68 | 0.333 | 5.6E-03 | 0.313 | 7.9E-02 |

Below, the pieces of code proposed for the parameterization of IntCyt are applied to the implementation of IntCyt contained in the file `main.py` located in the directory `software`.

**2.8.2. Restricting fission and fussion of cells.** Recall that the list `filtering` introduced in the tutorial (see section 2.8.5) was used to control the rate of fusion and fission events of our algorithm, which also holds in the case of IntCyt. The higher the components of `filtering` are, the more difficult it is for IntCyt to shuffle information through hierarchical compartmentalization.

While the component `filtering[2]` was set to `0` throughout the whole section 2.8.5 in order to promote information sharing between organelles, it is more advantageous, for the `DREAM3` challenge, to make this component take positive values in order to prevent any misleading commutation between these organelles. Such precaution is necessary because the `DREAM3` challenge requires precise predictions of differential gene expression.

Hence, we set the components of `filtering` as in the last example of section 2.7, in which fusion and fission events are permitted to occur more easily overtime.

main.py

```
#------------------
step = i/float(iterations)
interval = lambda s, a, b:  a*(1-s)+b*s if s < 1 else b
#------------------
  start = 4
  epoch = 4
  fission_events = [0]
  fusion_events = [2]
  compose_events = [1,3]
  events = [start,epoch,fission_events,fusion_events,compose_events]
#------------------
filtering = [1.5, #To control fission
             1.5, #To control fusion
             interval(step,10,2)] #To control fission and fusion
```

The reason for making the parameter `filtering[2]` decrease overtime is explained by the following facts:

1) early compartmentalization of cells that are not specialized (*i.e.* differentiated) can create conflicts of learning interest in later stages, because the specialization of a cell may be too different from the cells with which it possess an early apparent similarity;

2) cell communication is desirable when specialization has already been initiated because it accounts more for the information contained in the dataset and less for the guiding information used for self-supervision.

**2.8.3. Setting up brightness levels and contrast profiles.** In the tutorial (see section 2.8.5), our brightness levels were defined through ranges of percentages reflecting the variability of the learning rate. However, in the case of the `DREAM3` challenge, the learning rate of IntCyt will be relatively slow (see the discussion at the end of the section). As a result, our brightness levels for the `DREAM3` challenge will consist of a unique percentage. More specifically, we consider the following contrast profiles to decide whether a cell needs to be updated or not:

1) *the expression profile is still in the process of being learned* if 85% of the missing expression values are greater than or equal to 98% of the maximum value of the expression profile;

2) *the expression profile is considered sufficiently-learned* if 85% of the missing expression values are less than or equal to 98% of the maximum value of the expression profile;

main.py

```
  #------------------
  if sys.argv[1] == "DREAM3":
    #------------------
    brightness = [.98]
    profiles = [ [(.85,1)], [(0,.85)] ]
```

The fact that the brightness level and the contrast profiles are relatively high reflects the fact that the expression values being learned will not change much compared to the default value of `100` initially given to all the missing gene expressions. This can be explained by the high agreement thresholds that we will consider in section 2.8.4 for the parameterization of IntCyt, as the higher these thresholds are, the less influence each input has on the overall learning of IntCyt, and the slower the learning is.

**2.8.4. Setting up the agreement thresholds.** In this section, we explain how we determine agreement thresholds (see section 2.8.5) for defining contrast profiles suited to the learning of the `DREAM3` dataset by IntCyt. In a few words, we find the range of the best agreement scores between the hidden genes and the known genes and decompose this range into equal sub-intervals to find accumulation points, which allow us to determine agreement thresholds that we can use with the contrast profiles. We achieve this analysis through with script `data_processing.py`, located in the directory `software`, by calling the option `method` (see section 1.5 for a description of the other options) .

We shall now proceed in three steps:

**(1)** we associate every hidden gene with a distribution accounting for the agreement scores that they possess with the known genes;

**(2)** we use these distributions to determine an approximate range of agreements to be used to reconstruct the expressions of the hidden genes;

**(3)** we decompose this range into intervals of equal lengths to evaluate the distribution of the agreements and find adequate thresholds for the contrast profiles.

**Step 1**: Throughout this section, we will only need to use the gene expressions of the strains that do not miss any data, namely `wt`, `strain2`, and `strain3`. First, we decompose their associated gene expressions into two matrices `hidden` and `known` accounting for the expressions of the known genes and the hidden genes, respectively (see below). Each of these matrices possess $8 \times 3 = 24$ columns, describing the expression values associated with the three strains `wt`, `strain2`, and `strain3`.

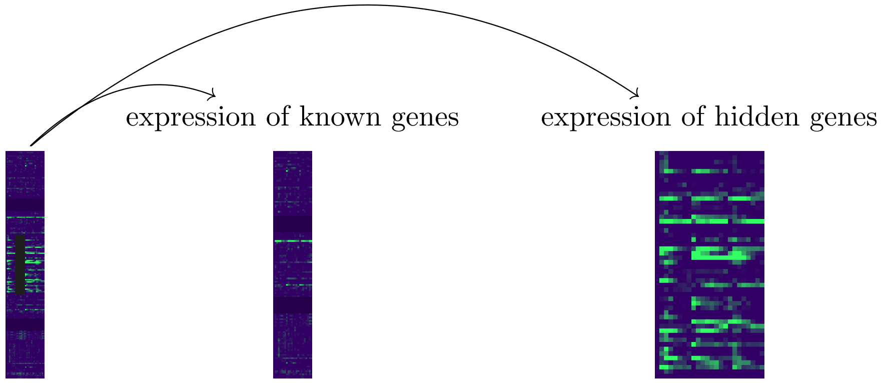

For every hidden gene `i` of expression `hidden[i]`, we use normalized scalar products, which we also called *agreements* in section 2.8.5, to assess the closeness of the hidden gene `i` with each of the known genes `j` of expression `known[j]`. Specifically, we define a matrix, call it `intensities`, such that, for every hidden gene `i`, the `j`-th elements of the list `intensities[i]` is the following agreement score.

$$\mathsf{argeement}(\texttt{hidden[i]},\texttt{known[j]}) = \frac{\texttt{hidden[i]} \cdot \texttt{known[j]}}{||\texttt{hidden[i]}||\,||\texttt{known[j]}||}$$

Below, we show the lists `intensities[i]`, for each of the 50 genes, in the form of heat maps:

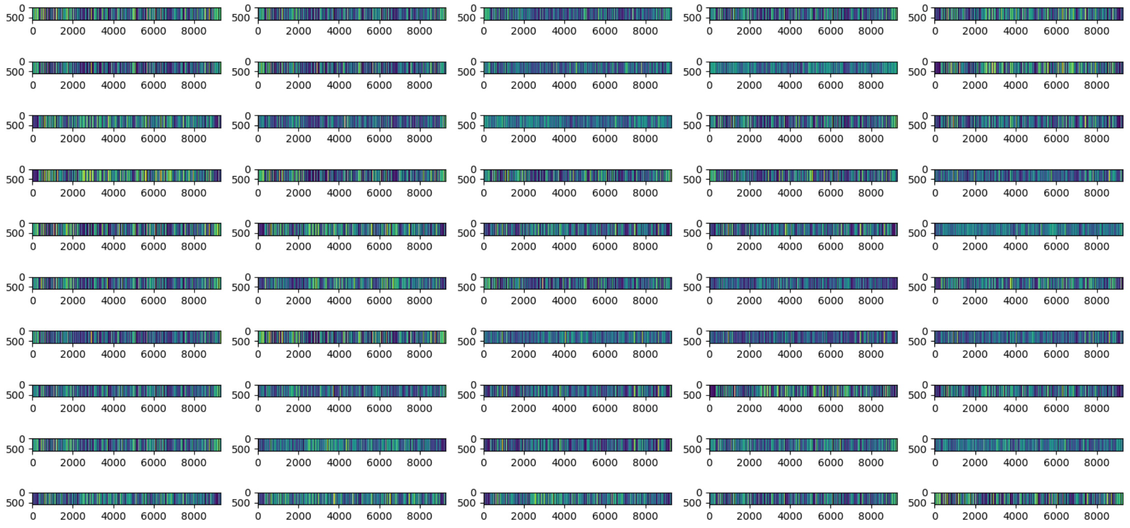

We now divide each of the lists `intensities[i]` in 100 bins of equal length and map each bin to the number of agreements scores belonging to the bin. This process gives us a non-normalized probability distribution for every hidden gene. Below, we plot these distributions from the highest agreement scores to the lowest – the bins with zero counts are displayed in red (mainly located at the beginning of the distribution, where agreement scores are the highest).

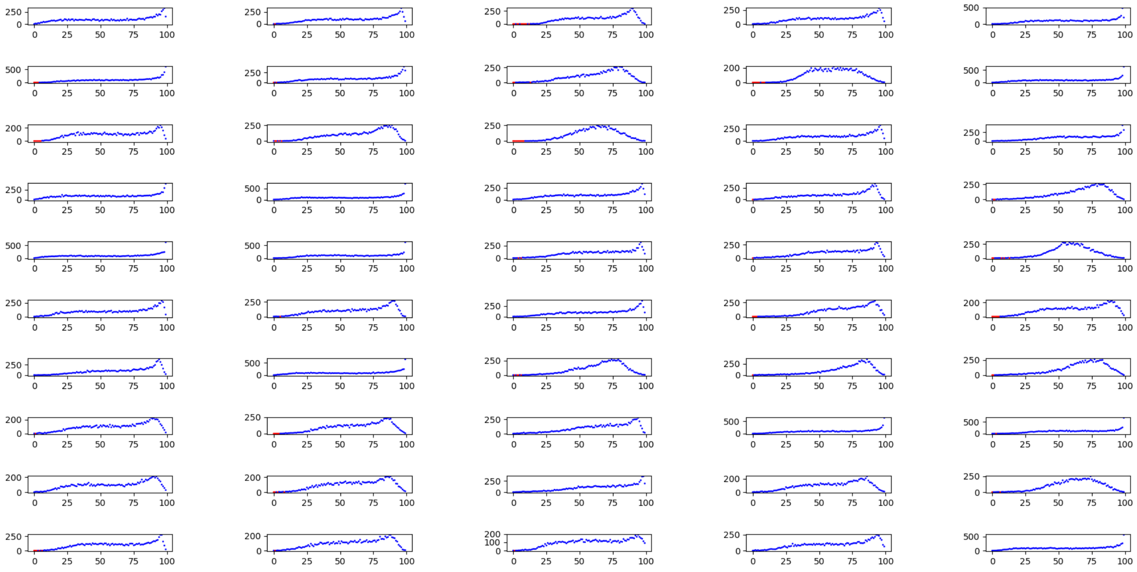

**Step 2**: For this second step, we select, for each hidden gene, the first bin of each distribution that includes non-zero counts and consider the agreement scores associated with this bin. These agreement scores give us an assessment of the highest scores associated with each hidden gene. We can therefore use these agreement scores to detect the most relevant gene expressions needed to reconstruct the missing expressions of the hidden genes. Below,

we display these scores for the 50 hidden genes.

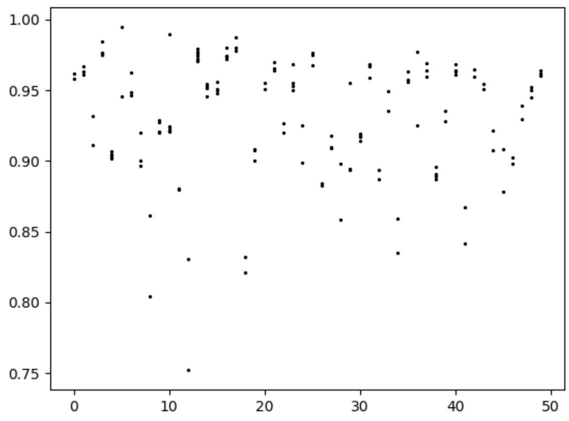

As can be seen, these agreement scores range from 0.75 to 1.00. Also, we can see a clear accumulation of agreement scores around .95 or .9. This tells us that setting the agreement threshold of our contrast profiles to .9 will select most of the known gene expressions agreeing with the hidden gene expression. Unfortunately, doing so will also prevent the selection of known gene expressions whose highest agreement scores are lower than 0.9. We could reduce the threshold to 0.75 for all genes, but this threshold may also cause the inclusion of known gene expressions that are too different from the hidden gene expressions that we want to infer.

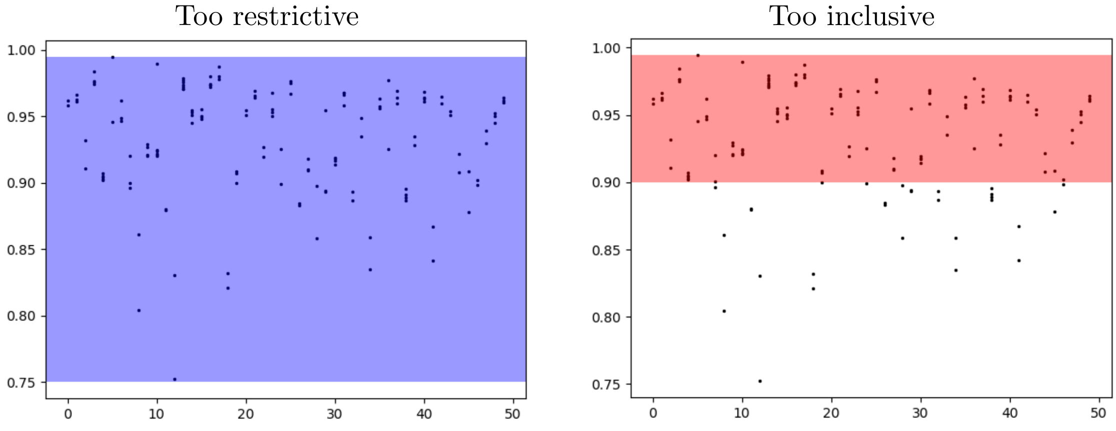

As a result, we want to use a collection of thresholds that we shall apply successively to first select the known gene expression with the highest scores, and then select those with lower scores while minimizing their influence on the information learned by IntCyt at higher agreement thresholds.

**Step 3**: To select our thresholds, we associate every hidden gene with the average of its previously selected agreements, and we bin these averages in various subdivisions, ranging from the minimum to the maximum averages, with intervals of equal length, as shown below.

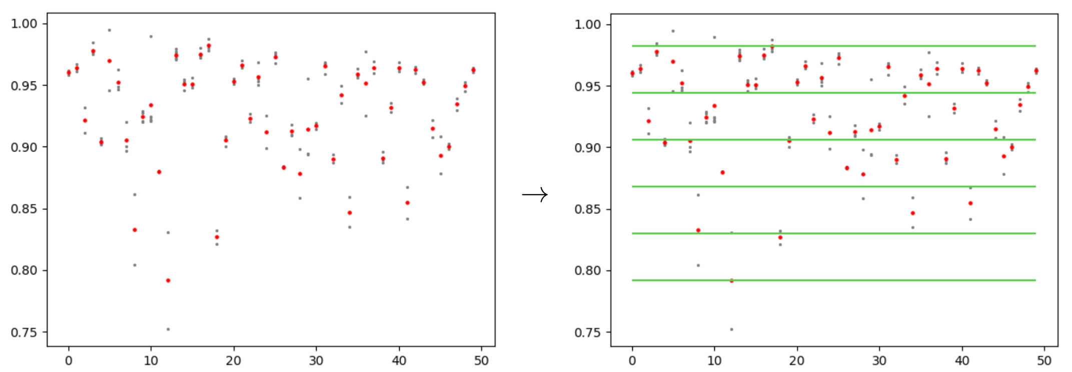

The lower lines of each of these bins define potential thresholds that we can use to select known gene expressions. Every subdivision of our set of agreement score induces a distribution mapping every bin to the number of agreement scores in the bin. We propose to find the most optimal subdivisions of our set of agreements scores by finding a distribution that shows clear accumulation areas (or clusters). More specifically, we analyze a set of subdivisions whose number of bins vary from 50 to 3, as shown below.

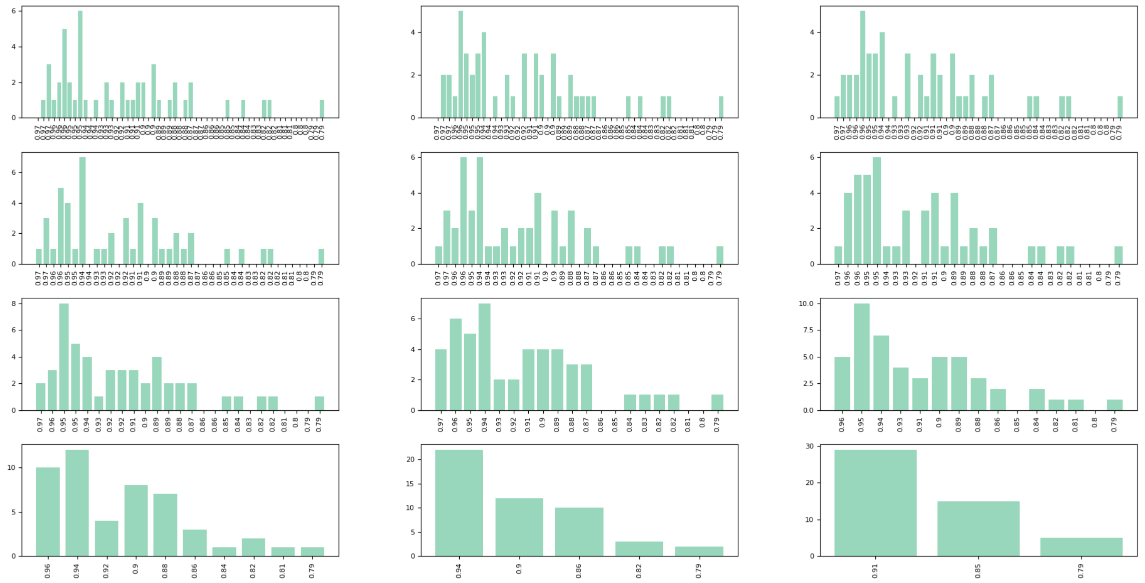

We can assess the different accumulation areas by smoothening these distributions through moving averages. We find that the distribution of our set of selected agreement scores almost always include three clusters (see the bumps of the purple curves).

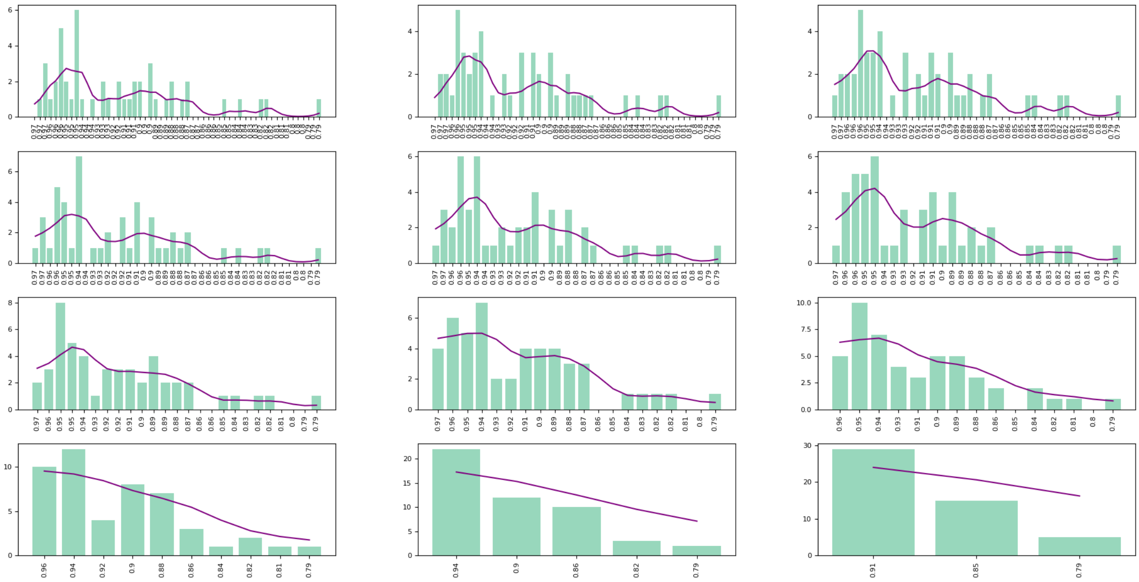

Note that determining how many clusters there are and where these clusters stars and ends on the x-axis is left to the discretion of the user. In our case, we decided that there were three clusters that could be described by Gaussian functions such that the start of these three clusters are approximately given by the scores `0.9`, `0.85` and `0.8` – this is illustrated below by the blue Gaussian curve, which we use to find the threshold `0.9`.

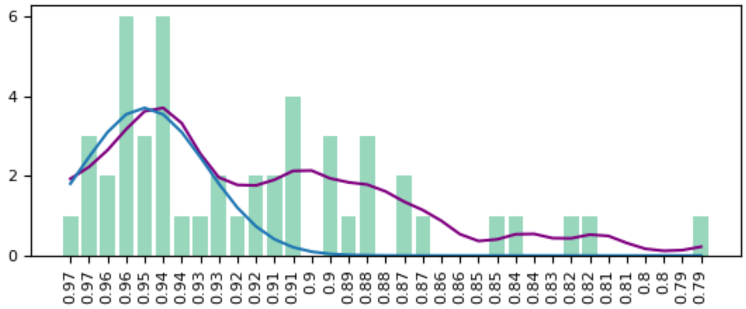

We will later use these agreement thresholds with tailored contrast profiles to analyze the three clusters detected above separately.

**2.8.5. Adjusting the gamma parameter.** In this section, we determine multiplicative factors to be applied to the gamma parameter of the gradient descent in order to correct for the lowering of the agreement thresholds (section 2.8.4) across the three runs of IntCyt on the dataset `dream3_training.gz`. Indeed, the lowering of the thresholds is likely to lower the quality of the updates operated through the gradient descent. Hence, we also want to reduce the intensity of the updates accordingly.

Recall that, at every update step, the gradient descent increments each of the parameters of IntCyt by a value of the following form, where the differential encodes a "directionality" and $\gamma$ encodes an intensity.

$$-\gamma \times \frac{\partial_{j,v} U^2(c)}{\partial c}$$

Within each run of IntCyt on the dataset `dream3_training.gz`, the parameter $\gamma$ should be kept approximately constant, because the agreement threshold does not change. However, the parameter should be reduced by a factor less than 1 between runs. The value of the factor is left to the discretion to the user. In our case, we reduce the order of the gamma parameter by a factor of 0.6 between each run, as shown below.

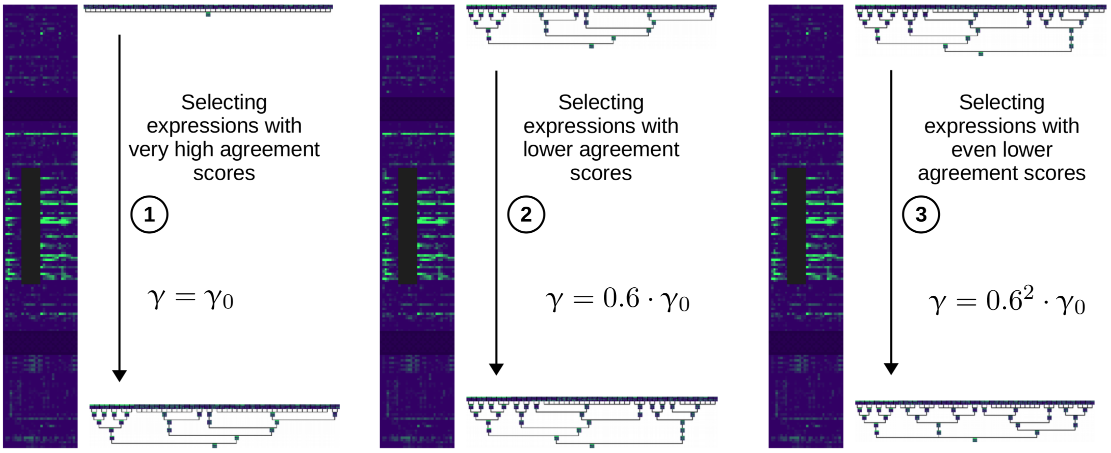

Note that the gamma parameter is reduced by a factor of $0.6^2 = 0.36$ from the first run to the last run. The different values of $\gamma$ will be used in section 2.8.6 to determine the values of two parameters, namely the parameters `E` and `F` that were used in the tutorial (section ).

**2.8.6. Using IntCyt to impute missing gene expressions.** In this section, we summarize our previous analyses and give the full parametrization of IntCyt for the `DREAM3` challenge. This parametrization can be found in the file named `main.py` located in the directory `software`. Note that, in this file, the dataset is loaded through the following piece of code, in which we set the variable `ary` is 50 and define the variable `training_images` to represent the number of images in the dataset `dream3_training.gz`.

main.py

```
#-------------------
elif len(sys.argv) > 1 and sys.argv[1] == "DREAM3":
#-------------------
fim = gzip.open("data/dream3_training.gz","r")
training_images = 9285
listim = np.array(usf.get_memory(fim,1,range(training_images)))
#-------------------
ary = 50
#-------------------
image_size = [64,1,1] #height, width, depth
categories = map(str,range(training_images))
#-------------------
```

We now propose the following parametrization of IntCyt for the `DREAM3` challenge (see below for more explanation).

main.py
```
    #------------------
    if sys.argv[1] == "DREAM3":
      #-------------------
      brightness = [.98]
      profiles = [ [(.85,1)], [(0,.85)] ]
      if 0 <= i < training_images:
        scores = [ (.9,1), (.98,1) ]
        E = 11
        F = 20
      if training_images <= i < 2*training_images-1:
        scores = [ (.85,1), (.98,1) ]
        E = 11.25
        F = 20
      if 2*training_images <= i < iterations:
        scores = [ (.8,1), (.98,1) ]
        E = 11.5
        F = 20
```

The values for the variables `brightness` and `profiles` were discussed in section 2.8.3 and the values for the lists `scores` were discussed in section 2.8.4. The value of `F` is arbitrary, but was – in our case – influenced by the values considered for previously tested databases such as `MNIST` and `fashion-MNIST` (see section 2.8.5). There only remains to explain the different values of the parameters `E` for each of the if-conditions shown above, in the code.

As explained in section 2.8.5, the gamma parameter $\gamma$ should approximately take the values: $\gamma_0$, $0.6 \times \gamma_0$ and $0.36 \times \gamma_0$. Also, recall that the gamma parameter $\gamma$ can be estimated by the following formula:

$$\gamma = 10^{\mathtt{E}} \times \left(\frac{x}{M}\right)^{\mathtt{F}} \tag{2.3}$$

where $x$ stands for an agreement score between a certain organelle and an input and $M$ is a maximal upper bound for all the agreements between the organelles and their associated inputs. Our strategy is to use the different values of $\gamma$ and formula (2.3) to determine the values of the parameter `E` accordingly.

According to formula (2.3), the parameter $\gamma_0$ can be estimated as follows during the first run of `IntCyt` on the dataset `dream3_training.gz`.

$$\gamma_0 = 10^{11} \times \left(\frac{.9}{M}\right)^{20}$$

This means that the parameter `E` used in the second if-condition (see above) satisfies the following equation:

$$10^{\mathtt{E}} \times \left(\frac{.85}{M}\right)^{20} = 0.6 \times 10^{11} \times \left(\frac{.9}{M}\right)^{20}$$

Solving this equation for `E` gives the following equation:

$$\mathtt{E} = 11 + 20 \times \mathsf{log}\left(\frac{.9}{.85}\right) + \mathsf{log}(0.6) \simeq 11.27$$

For our implementation, we took the value `11.25` as a representative approximation of the previous calculation. We proceeded similarly for the exponent `E` used in the third run. This time, we have the following estimate:

$$\mathtt{E} = 11 + 20 \times \mathsf{log}\left(\frac{.9}{.8}\right) + \mathsf{log}(0.36) \simeq 11.57$$

For our implementation, we took the value `11.5` as a representative approximation of the previous calculation.

**2.8.7. Integrating the results.** After applying the parametrization of section 2.8.6 to the code of the file `main.py` located in the directory `software` and running the resulting implementation on the dataset `dream3_training.gz`, we find the results shown in red in the following table – these results are obtained from the script `data_processing.py` located in the directory `software`.

| Team | Score | Gene accuracy | Gene p-value | Time accuracy | Time p-value |
|---|---|---|---|---|---|
| **IntCyt** | **3.57** | **0.576** | **2.4E-06** | **0.489** | **2.9E-02** |
| GH | **3.25** | 0.563 | 6.5E-06 | 0.512 | 4.8E-02 |
| KNN | 3.18 | 0.558 | 1.1E-05 | **0.533** | 3.9E-02 |
| Team 263 | 1.85 | 0.421 | 7.5E-03 | 0.112 | 2.7E-01 |
| Team 297 | 1.68 | 0.333 | 5.6E-03 | 0.313 | 7.9E-02 |

These results show that IntCyt can be considered as a competitive learning machine algorithm for data science. We finish the present section with more in-depth analyses of the results shown in the previous table.

First, we show the gene accuracy scores, given by Spearman's correlations, followed by their corresponding $p$-values. Each of the values given in the following graphs represents one of the 8 time points associated with the expression measurements of the `DREAM3` dataset. The accuracy scores are shown in the following graph, in which the red line shows the overall gene accuracy score:

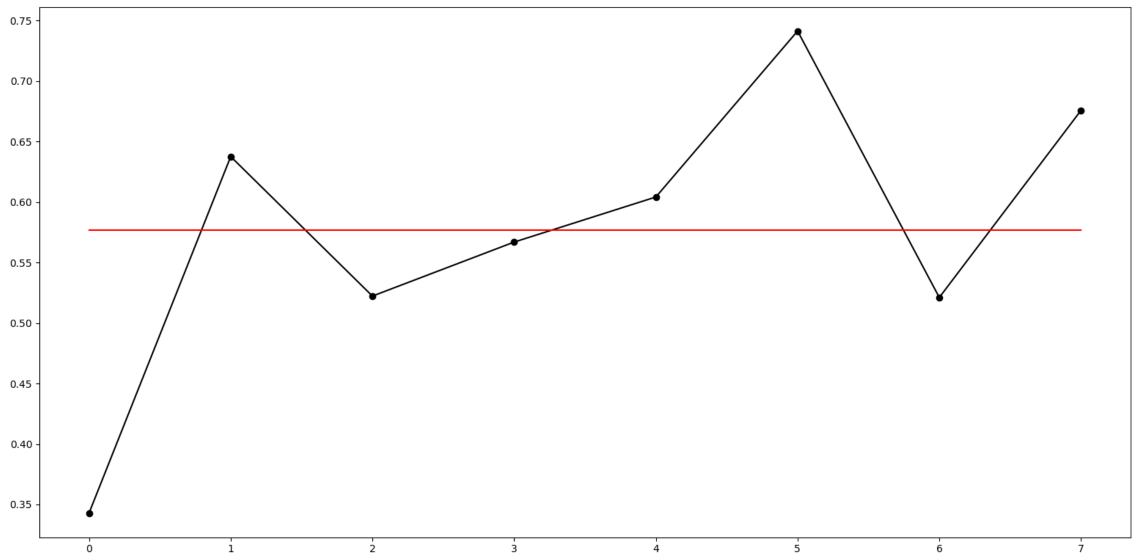

The $p$-values are given by the following graph:

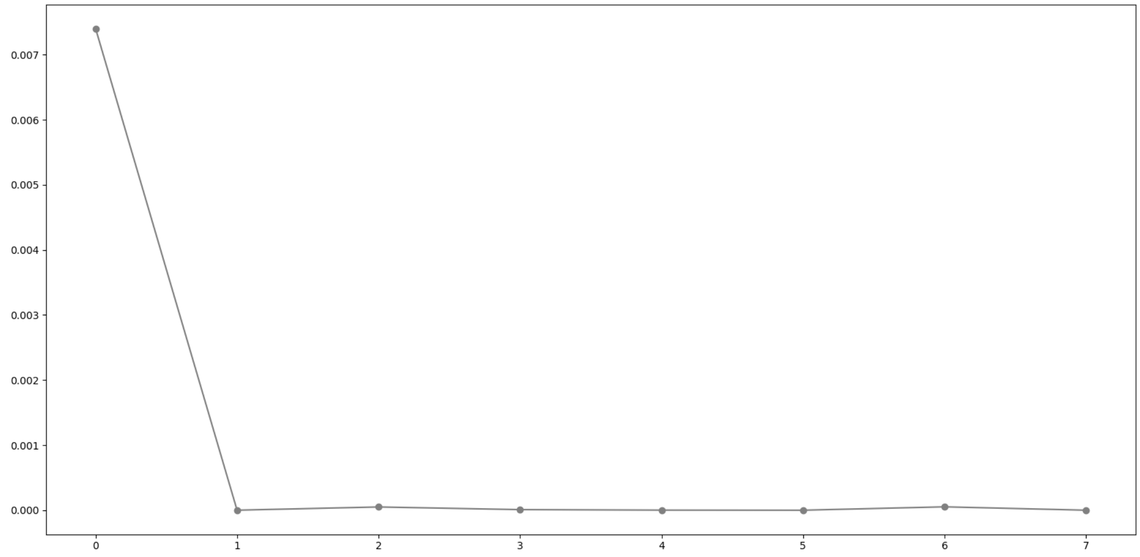

We now show the time accuracy scores, also given by Spearman's correlations, and their corresponding $p$-values. Each of the values given in the following graphs represents one of the 50 hidden genes associated with the `DREAM3` challenge. The accuracy scores are shown in the following graph, in which the red line shows the overall time accuracy score:

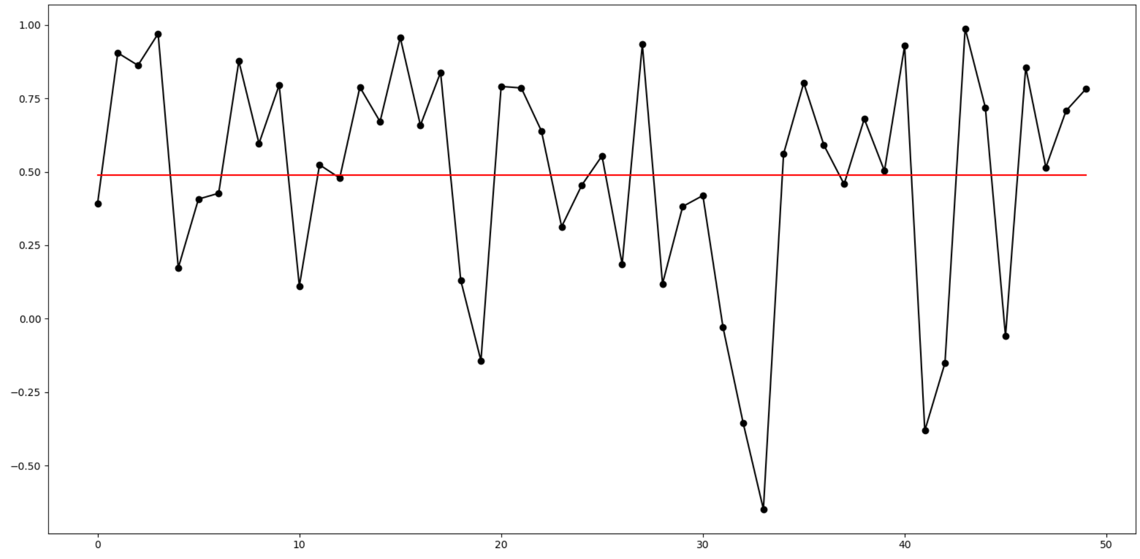

The $p$-values are given by the following graph:

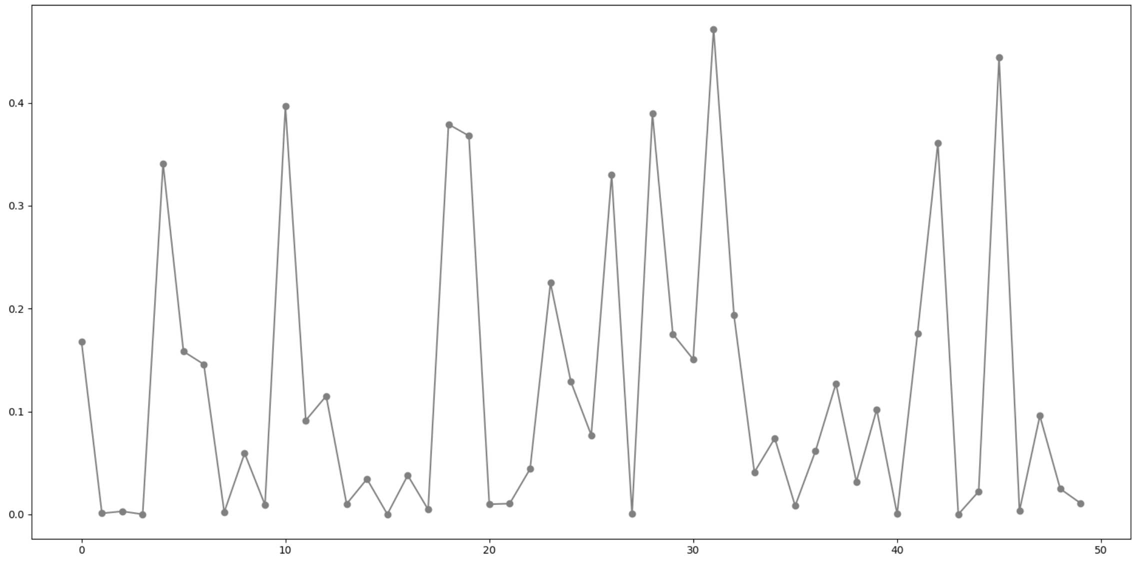

Finally, we show the change in the accuracy scores accross the three runs of IntCyt on the dataset `dream3_training.gz`. The three runs are separated by red dots, shown at the values 40000 and 80000 on the x-axis.

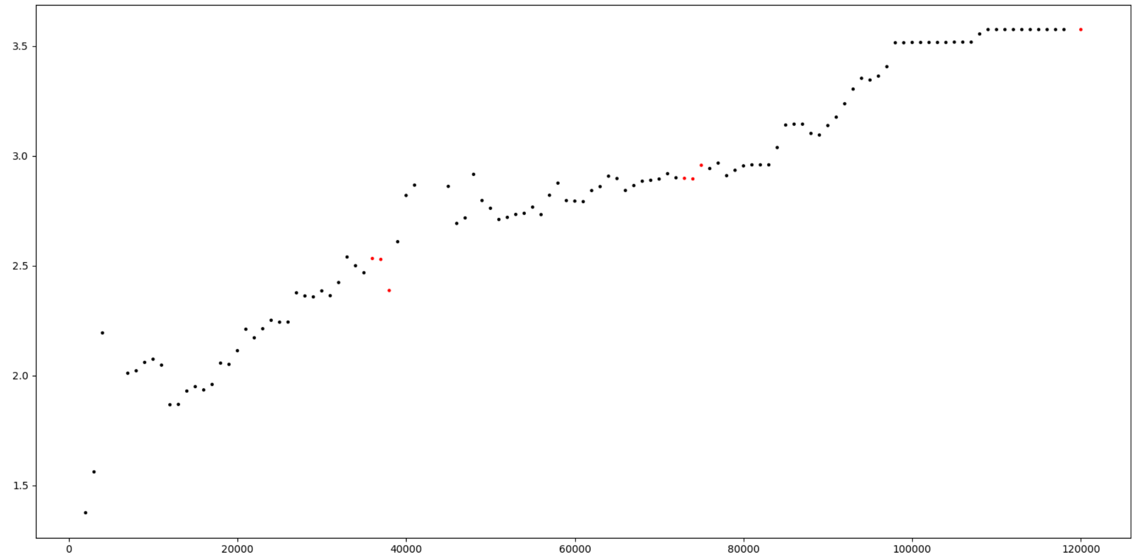

The previous graph can be compared to the types of graphs discussed in section 2.8.1 for accuracy scores. As can be observed, the type of graphs shown above corresponds to the anticipated (and desired) type of graphs discussed in section 2.8.1.

Chapter 3

### Presentation of the module useful.py

#### 3.1. Description of usf (class item)

**3.1.1. Introduction.** This section introduces the reader to the class item usf, which is a repository of functions used in the tutorial of section 2 as well as in the class Cell (section 4.1). The class item usf is also used in the file data_processing.py contained in the director software of the library.

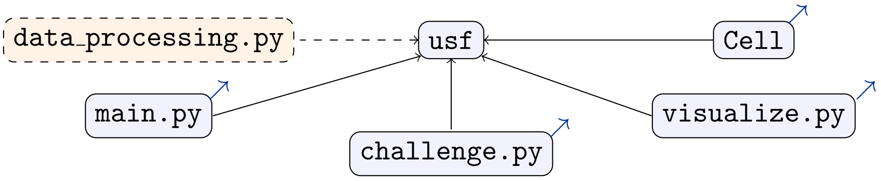

Note that usf is not a class, but a item of the non-importable class _Useful. The class _Useful is equipped with 5 types of methods, which are listed below:

▷ a set of 2 methods taking care of controling the learning rate of intcyt (section 5.1):

- contrast: indicates whether a vector satisfies a certain contrast profile;
- gamma: corresponds to the gamma parameter used in [1, Sec. 2.3].

▷ a set of 4 methods taking care of saving and displaying information stored in a super cell:

- print_data: saves or displays a vector;
- print_root: saves or displays the root of a super cell [1, Conv. 2.2];
- print_organelles: saves or displays the organelles of a super cell [1, Def. 2.7];
- print_tree: saves or displays the structure of a super cell [1, Conv. 2.2];

▷ a set of 4 methods loading information from saved data:

- parse_image: parses the components of a vector stored in a file;
- get_memory: loads a collection of vector data stored in a file;
- get_trees: loads a super cell structure stored in a file;
- get_last_cycle: returns the index of the last super cell stored in a file;

▷ a set of 6 methods displaying the evolution of a super cell from saved data:

- join_images: attach two images next to each other;

• rgb_colormap: returns an RGB code color for bicolor and RGB images;

• make_rgb_panel: displays a row of images representing memorized data in a super cell;

• make_rgb_grid: displays a grid of images representing memorized data in the organelles of a super cell;

• make_rgb_tree: displays a tree of images representing memorized data in a super cell;

• draw_arrow: displays an arrow going down.

▷ a set of 3 functions taking care of the combinatorics behind cell fusion and cel fission:

• zero_matrix: returns a square matrix initialized with zeros;

• join_fibers: computes the transitive closure of two partitions;

• cliques: returns one of the maximally weighted connected components of a weighted adjacency matrix.

**3.1.2. Structure.** The following tables give a preview of the class item usf. The table given below describes the various dependencies of the class item.

| Dependencies | |
|---|---|
| Class type | Module section |
| _Useful | N/A |
| Statistics | |
| ▷ Importable objects: 0 | |
| ▷ Non-importable objects: 0 | |
| ▷ Importable functions: 19 | |
| ▷ Non-importable functions: 0 | |

The following two tables give a description of the 16 importable functions of the class item. In the third column, we use the symbol ∼ to refer to a type that will be specified in the topmost section shown in the corresponding rightmost column.

| Functions | | | |
|---|---|---|---|
| Name | Input types | Output types | Related sections |
| .contrast | - list(float)<br>- list(list(float)) | - fun: ∼ -> ∼ | ▷ section 3.1.3 |
| .gamma | - float<br>- float<br>- list(float)<br>- list(list(float * float))<br>- list(float)<br>- list(list(float)) | - fun: ∼ -> ∼ | ▷ section 3.1.4 |
| .print_data | - list(float)<br>- list(int)<br>- file<br>- string | - self | ▷ section 3.1.5 |
| .print_root | - SuperCell<br>- list(int)<br>- file<br>- string | - self | ▷ section 3.1.6 |
| .print_organelles | - SuperCell<br>- list(int)<br>- file<br>- string | - self | ▷ section 3.1.7 |

In this second table, the type image refers to a sub-type of list(list(list(float))) consisting of lists of lists of RGB lists (lists of float of length 3).

| | | | |
|---|---|---|---|
| .print_tree | - SuperCell<br>- list(int)<br>- file<br>- string<br>- int<br>- int | - self | ▷ section 3.1.8 |
| .parse_image | - string<br>- list(string)<br>- list(float) | - list(string) | ▷ section 3.1.9 |
| .get_memory | - file<br>- int<br>- list(int) | - list(list(float)) | ▷ section 3.1.10 |
| .get_trees | - file<br>- list(int) | - list(list(float)) | ▷ section 3.1.11 |
| .get_last_cycle | - file<br>- int | - int | ▷ section 3.1.12 |
| .join_images | - image<br>- image<br>- int<br>- int<br>- bool | - image | ▷ section 3.1.13 |
| .rgb_colormap | - float<br>- bool<br>- float | - fun: ∼ -> ∼ | ▷ section 3.1.14 |
| .make_rgb_panel | - list(list(float))<br>- list(int)<br>- bool<br>- list(float) | - image | ▷ section 3.1.15 |
| .make_rgb_grid | - list(list(list(float)))<br>- list(int)<br>-bool | - image | ▷ section 3.1.16 |
| .make_rgb_tree | - list(list(float))<br>- list(int)<br>- bool | - image | ▷ section 3.1.17 |
| .draw_arrow | - int<br>- int | - image | ▷ section 3.1.18 |
| .zero_matrix | - int | - list(list(int)) | ▷ section 3.1.19 |
| .join_fibers | - list(list(int))<br>- list(list(int)) | - list(list(int)) | ▷ section 3.1.20 |
| .cliques | - list(list(int)) | - list(list(int)) | ▷ section 3.1.21 |

**3.1.3. Description of usf.contrast (function).** This section describes the code and the functionalities of the function usf.contrast. The method is equipped with the following inputs.

| .__init__ | | |
|---|---|---|
| Inputs | Types | Specifications |
| brightness | list(float) | necessary |
| profiles | list(list(float * float)) | necessary |

The method possesses one action, which we describe below through examples.

| Action | |
|---|---|
| Condition | Always |
| Description | The method returns a function `contrast_profiles` that takes a `float` item `vector` and an optional `list(list(float))` item `challenge` and returns a `list(bool)` item `contrast_tests` whose coefficient `contrast_tests[i]` indicates whether the input `vector` *satisfies the contrast profile* defined by the list `profiles[i]` and the value `brightness[i]` (see the tutorial or the explanation below). To explain what this means, let us denote by `percents[j]` the percentage of indices `u` (ranging from `0` to `self.dimension`) for which<br>1) the ratio `vector[u]/max(vector)` is greater than `brightness[j]`;<br>2) the following equation holds (when `challenge` is non-empty):<br>`challenge[0][0] <= u % challenge[0][2] <= challenge[0][1]`.<br>Then, the `bool` item `contrast_tests[i]` indicates whether there exists an index `j` for which the following inequality holds:<br>`profiles[i][j][0] <= percents[j] <= profiles[i][j][1]`<br>As a result, the output list `contrast_tests` indicates whether any of the contrast profiles are satisfied by the input `vector`. |

In the following examples, we illustrate the use of the function `usf.contrast` with no input `challenge` given. Note that giving such an input would amount to the same type of behavior, except that some of the components of the lists contained in `vectors` would not be considered in the computation.

```
>>> brightness = [.25,.5,.75]
>>> profiles = [[(.0,.9),(.0,.5),(.0,.25)],[(.0,.4),(.0,.6),(.0,.5)]]
>>> vectors = [[1,2,5,3,7], [5,0,0,0,4],[25,51,8,1,52]]
>>> contrast_profiles = usf.contrast(brightness,profiles)
>>> for j in range(len(vectors)):
...  print "contrast profiles:  " + str(contrast_profiles(vectors[j]))
...  print "----------"
...
[brightness] 0.8 0.4 0.2
True False

contrast profiles:  [True, False]
----------
[brightness] 0.4 0.4 0.4
False True

contrast profiles:  [False, True]
----------
[brightness] 0.6 0.4 0.4
False False

contrast profiles:  [False, False]
----------
```

The grey messages displayed in the previous example appear automatically in the standard output. They show the brightness profiles associated with each of the lists of the input `vectors` and indicate whether these profiles satisfy the contrast profiles defined by the pair `brightness` and `profiles`. In the context of the previous example, the messages inform us that the list `vectors[0]` satisfies the contrast profile `profiles[0]`, the list `vectors[1]` satisfies the contrast profile `profiles[1]`, and the list `vectors[2]` does not satisfy any of the contrast profiles of `profiles`.

**3.1.4. Description of `usf.gamma` (function).** This section describes the code and the functionalities of the function `usf.gamma`. The method is equipped with the following six inputs, including one that is optional.

| `.__init__` | | |
|---|---|---|
| Inputs | Types | Specifications |
| `E` | `float` | necessary |
| `F` | `float` | necessary |
| `brightness` | `list(float)` | necessary |
| `profiles` | `list(list(float * float))` | necessary |
| `scores` | `list(float * float)` | necessary |
| `*challenge` | `list(list(float))` | optional |

The method possesses one action, which we describe below through examples.

| Action | |
|---|---|
| Condition | Always |
| Description | The method returns a function `gamma_parameter` that plays the role of the "gamma parameter" discussed in [1, Sec. 2.3]. Specifically, the function `gamma_parameter` takes a `Cell` item `c` (see section 4.1.3) and a `list(list(float))` item `a` and returns a `list(list(float))` item `gamma_parameter(c,a)` whose coefficients `gamma_parameter(c,a)[i][u]` satisfies the following formula: $$\texttt{gamma\_parameter(c,a)[i][u]} = 10^{\texttt{E}} \times \left(\frac{\texttt{agreement[i]}}{\texttt{max(agreement[k]} \mid \texttt{k)}}\right)^{\texttt{F}}$$ in which the `list` item `agreement` is computed as $$\texttt{agreement[i]} = \begin{cases} 0 & \text{if } (*) \\ \texttt{c.agreement(i,a[i],*challenge)} & \text{else.} \end{cases}$$ where (∗) is the property that there exists an index `j` for which: 1) the *i*-th organelle of `c` satisfies the contrast profile defined by the list `profiles[j]` and the value `brightness[j]`, which means that the list returned by the function `self.contrast(brightness,profiles)(c.organelles[i],*challenge)` contains at least one `True` value (see section 3.1.3); 2) the agreement `c.agreement(i,a[i],*challenge)` (see section 4.1.16) is greater than or equal to `scores[j][0]` and less than or equal to `scores[j][1]`. |

In the following examples, we illustrate the use of the function `usf.gamma` with no input `challenge` given. Note that giving such an input would amount to the same type of behavior, except that some of the components of the organelles of the `Cell` item `c` (see section 4.1.3) would not be considered in the computation.

```
>>> E = 1
>>> F = 1
>>> brightness = [.25,.5,.75]
>>> profiles = [[(.0,.9),(.0,.5),(.0,.25)],[(.0,.4),(.0,.6),(.0,.5)]]
>>> scores = [(.5,1),(.8,1)]
>>> gamma_parameter = usf.gamma(E,F,brightness,profiles,scores)
>>> c = Cell(dimension = 5, residual = 0.5, cytosol = [-2,3,5,0,0],
organelles = [[1,2,5,3,7], [5,0,0,0,4],[25,51,8,1,52]])
>>> a = [[70,9,5,7,58],[1,5,65,2,44],[8,7,87,5,100]]
>>> g = gamma_parameter(c,a)
[brightness] 0.8 0.4 0.2
True False

[brightness] 0.4 0.4 0.4
False True

[brightness] 0.6 0.4 0.4
False False
```

Above, the grey messages inform us that

- the first organelle of `c` satisfies the contrast profile defined by `profiles[0]`,

- the second organelle of `c` satisfies the contrast profile defined by `profiles[1]`,

- the third organelle of `c` does not satisfy any of the contrast profiles of `profiles`.

However, the previous data is only half of the information needed by `gamma_parameter` to determine the values of the output `g`. Specifically, the function `gamma_parameter` also tests whether the agreements of the organelles with their respective inputs belong to the corresponding interval given in `scores`. If the agreement associated with the $j$-th organelle of `c` is less than the threshold `scores[j][0]` or greater than the threshold `scores[j][1]`, then the elements of the $j$-th list of `g` are zero values.

We illustrate the previous discussion through the following display, in which we show the agreements and the values for the gamma parameters together.

```
>>> for j in range(len(g)):
...   print "agreement:  "+str(c.agreement(j,a[j]))
...   print "gammas:  "+str(g[j])
...
```

Below, the gamma parameter associated with the first organelle of `c` is equal to `10.0 * 1.0` because that organelle is associated with the maximum agreement, which also is greater than `scores[0][0]` and less than `scores[0][1]`.

```
agreement:  0.627367719768
gammas:  [10.0, 10.0, 10.0, 10.0, 10.0]
```

Now, the gamma parameter associated with the second organelle of `c` is equal to `10.0 * 0.0` because its associated agreement is less than `scores[1][0]`.

```
agreement:  0.359257831799
gammas:  [0.0, 0.0, 0.0, 0.0, 0.0]
```

Finally, the gamma parameter associated with the third organelle of `c` is equal to `10.0 * 0.9990995982390753` because that organelle does not satisfy any of the contrast profile and hence does not need to be filtered through an agreement test.

```
agreement:  0.626802836768
gammas:  [9.990995982390753, 9.990995982390753, 9.990995982390753,
9.990995982390753, 9.990995982390753]
```

**3.1.5. Description of** `usf.print_data` **(function).** Writing in progress – see `cl_usf.py` for more information.

**3.1.6. Description of** `usf.print_root` **(function).** Writing in progress – see `cl_usf.py` for more information.

**3.1.7. Description of** `usf.print_organelles` **(function).** Writing in progress – see `cl_usf.py` for more information.

**3.1.8. Description of** `usf.print_tree` **(function).** Writing in progress – see `cl_usf.py` for more information.

**3.1.9. Description of** `usf.parse_image` **(function).** Writing in progress – see `cl_usf.py` for more information.

**3.1.10. Description of** `usf.get_memory` **(function).** Writing in progress – see `cl_usf.py` for more information.

**3.1.11. Description of** `usf.get_trees` **(function).** Writing in progress – see `cl_usf.py` for more information.

**3.1.12. Description of** `usf.get_last_cycle` **(function).** Writing in progress – see `cl_usf.py` for more information.

**3.1.13. Description of** `usf.join_images` **(function).** Writing in progress – see `cl_usf.py` for more information.

**3.1.14. Description of** `usf.rgb_colormap` **(function).** Writing in progress – see `cl_usf.py` for more information.

**3.1.15. Description of** `usf.make_rgb_panel` **(function).** Writing in progress – see `cl_usf.py` for more information.

**3.1.16. Description of** `usf.make_rgb_grid` **(function).** Writing in progress – see `cl_usf.py` for more information.

**3.1.17. Description of** `usf.make_rgb_tree` **(function).** Writing in progress – see `cl_usf.py` for more information.

**3.1.18. Description of** `usf.draw_arrow` **(function).** Writing in progress – see `cl_usf.py` for more information.

**3.1.19. Description of** `usf.zero_matrix` **(function).** Writing in progress – see `cl_usf.py` for more information.

**3.1.20. Description of** `usf.join_fibers` **(function).** Writing in progress – see `cl_usf.py` for more information.

**3.1.21. Description of** `usf.cliques` **(function).** Writing in progress – see `cl_usf.py` for more information.

#### 3.2. Description of debug_time (class item)

**3.2.1. Introduction.** This section introduces the reader to the class item `debug_time`, which gives an easy way to debug programs using this library (see examples in section 2).

**3.2.2. Structure.** The following tables give a preview of the class item `usf`. The table given below describes the various dependencies of the class item.

| Dependencies | |
|---|---|
| Class type | Module section |
| _DebugTime | N/A |
| Statistics | |
| ▷ Importable objects: 3 | |
| ▷ Non-importable objects: 0 | |
| ▷ Importable methods: 3 | |
| ▷ Non-importable methods: 0 | |

The following table gives a description of the 3 importable objects of the class:

| Objects | | |
|---|---|---|
| Name | Type | Related sections |
| .event_names | list(string) | ▷ section 3.2.3 |
| .iteration | int | ▷ section 3.2.3 |
| .global_time | string | ▷ section 3.2.3 |

Finally, the following table gives a description of the 3 importable methods of the class:

| Methods | | | |
|---|---|---|---|
| Name | Input types | Output types | Related sections |
| .__init__ | - self | - self | ▷ section 3.2.3 |
| .set | - string | - self | ▷ section 3.2.4 |
| .call | - self | - self | ▷ section 3.2.5 |

**3.2.3. Description of debug_time.__init__ (function).** Writing in progress – see `cl_dbg.py` for more information.

**3.2.4. Description of debug_time.set (function).** Writing in progress – see `cl_dbg.py` for more information.

**3.2.5. Description of debug_time.call (function).** Writing in progress – see `cl_dbg.py` for more information.

*Chapter 4*

### Presentation of the module `celloperad.py`

#### 4.1. Description of `Cell` (class)

**4.1.1. Introduction.** This section introduces the reader to the code of the class `Cell`, which possesses the following external dependencies.

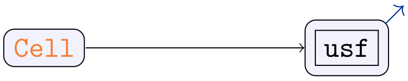

The main goal of the class `Cell` is to model the features of a cell, as defined in [1, Def. 1.2]. Before presenting the features of the class, recall that, mathematically, a cell of some dimension $N$, equipped with $n$ organelles, consists of a non-negative real number $\mathsf{res}(c)$, called the residual; a $N$-dimensional vector $\mathsf{cyt}(c) = (\mathsf{cyt}(c)_1, \dots, \mathsf{cyt}(c)_N)$ of real numbers, called the cytosolic content and a $n$-tuple $\mathsf{org}(c) = (\mathsf{org}(c)_1, \dots, \mathsf{org}(c)_n)$ of vectors in $\mathbb{R}_+^N$, where each vector $\mathsf{org}(c)_i$ is called the $i$-th organelles of $c$.

$c$ :

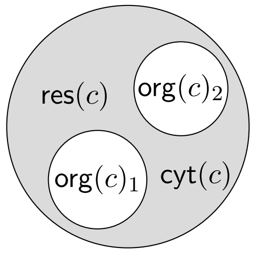

In this library, the class `Cell` is equipped with 9 objects, modeling various parameters pertaining to the previous definition, and 3 types of methods, which are listed below:

▷ a set of 3 methods meant to be used for testing and debugging:

- `check`: checks that the parameters of a cell are well-defined;
- `copy`: copies the data of a cell somewhere else in the memory;
- `stdout`: prints the list of parameters defining a cell on the standard output.

▷ a set of 9 methods meant to model the concepts developed in [1]:

- `__init__`: returns a cell for its associated set of parameters.
- `content`: returns the content $K(c)$ of a cell $c$ [1, Def. 1.21].
- `compose`: computes an operadic composition of cells [1, Def. 1.6].
- `compose_all`: computes a simultaneous operadic composition of cells [1, Def. 1.14].

- `left`: returns the cell $\mathsf{left}(c)(\alpha,\kappa,\lambda)$, as defined in [1, Conv. 1.19].
- `right`: returns the list $\mathsf{right}(c)(\mu,\alpha,\kappa)$ of cells, as defined in [1, Conv. 1.19].
- `spontaneous_reaction`: returns the *cleaning* of a cell – see [1, Rem. 1.31 & Def 2.13].
- `action`: returns the action of a cell on a vector, as defined in [1, Def. 1.29].
- `algebra_operator`: returns the values of the algebra operator $U(c,d)(a)$ as expressed in [1, Prop. 1.35 (1.4)], given that certain fitting conditions are satisfied.

▷ and a set of 8 methods engineering the ideas of [1] in order to produce the learning algorithm described therein:

- `allostasis`: generalizes the formula of the allostatic differential $\partial_{j,v}U^2(c,d)/\partial(c,d)$ given in [1, Prop. 1.48] to a more practical operator.
- `agreement`: returns the $k$-th agreement of a cell on a vector [1, Def. 2.22].
- `merge`: merges a subset of the organelles of a cell.
- `divide`: decomposes a cell $c$ into a pair $(c_1,c_2)$ of cells such that $c = c_1 \otimes c_2$, as illustrated in [1, Conv. 1.50 & Ex. 1.51], provided that the cytosol of $c$ is zero.
- `organelle_proportions`: returns the list of ratios $\mathsf{org}(c)_{n,u}/\mathcal{S}\mathsf{org}(c)_n$, which are used in the formula shown in [1, Th. 1.57].
- `content_proportions`: returns the list of ratios $\mathcal{S}\mathsf{org}(c)_n/(\sum_{k=1}^{n}\mathcal{S}\mathsf{org}(c)_k)$ appearing in the formula of [1, Prop. 1.54] provided that certain fitting conditions hold.
- `best_compartment`: returns the best partitioning of the organelles of a cell according to one of the criteria discussed in [1, Rem. 1.58].
- `proposed_clustering`: proposes an input cluster for the method `merge` or `divide`.

**4.1.2. Structure.** The following tables give a preview of the class `Cell`. The table given below describes the various dependencies of the class.

| Dependencies | |
|---|---|
| Superclass ancestry | Module section |
| `object` | N/A |
| Statistics | |
| ▷ Importable objects: 9<br>▷ Non-importable objects: 0<br>▷ Importable methods: 20<br>▷ Non-importable methods: 0 | |

The following table gives a description of the 9 importable objects of the class.

| Objects | | |
|---|---|---|
| Name | Type | Related sections |
| `.dimension` | `int` | ▷ section 4.1.3 |
| `.residual` | `float` | ▷ section 4.1.3 |
| `.cytosol` | `list(float)` | ▷ section 4.1.3 |
| `.organelles` | `list(list(float))` | ▷ section 4.1.3 |
| `.K` | `list(float)` | ▷ section 4.1.3 |
| `.SK` | `float` | ▷ section 4.1.3 |
| `.Sorg` | `list(float)` | ▷ section 4.1.3 |
| `.well_defined` | `bool` | ▷ section 4.1.3 |
| `.well_defined_cytosol` | `list(bool)` | ▷ section 4.1.3 |

Finally, the following table gives a description of the 20 importable methods of the class:

| Methods | | | |
|---|---|---|---|
| Name | Input types | Output types | Related sections |
| `.__init__` | - `int`<br>- `float`<br>- `list(float)`<br>- `list(list(float))` | - `self` | ▷ section 4.1.3 |
| `.check` | - `self` | - `self` | ▷ section 4.1.4 |
| `.copy` | - `self` | - `Cell` | ▷ section 4.1.5 |
| `.stdout` | - `self` | - `self` | ▷ section 4.1.6 |
| `.content` | - `self` | - `list(float)` | ▷ section 4.1.7 |
| `.compose` | - `int`<br>- `Cell` | - `Cell` | ▷ section 4.1.8 |
| `.compose_all` | - `list(Cell)` | - `Cell` | ▷ section 4.1.9 |
| `.left` | - `list(float)`<br>- `list(float)`<br>- `list(list(float))` | - `Cell` | ▷ section 4.1.10<br>▷ section 4.1.11 |
| `.right` | - `list(float)`<br>- `list(float)`<br>- `list(float)` | - `Cell` | ▷ section 4.1.11<br>▷ section 4.1.10 |
| `.spontaneous_reaction` | - `self` | - `self` | ▷ section 4.1.12<br>▷ section 4.1.7<br>▷ section 4.1.18<br>▷ section 4.1.20 |
| `.action` | - `list(list(float))` | - `list(float)` | ▷ section 4.1.13 |
| `.algebra_operator` | - `list(list(float))` | - `list(float)` | ▷ section 4.1.14 |
| `.allostasis` | - `list(list(float))`<br>- `list(float)`<br>- `int`<br>- `int` | - `float` | ▷ section 4.1.15 |
| `.agreement` | - `int`<br>- `list(float)`<br>- `list(float)` | - `float` | ▷ section 4.1.16 |
| `.merge` | - `list(list(float))`<br>- `string` | - `self` | ▷ section 4.1.17 |
| `.divide` | - `list(list(float))` | - `Cell * Cell` | ▷ section 4.1.18 |
| `.organelle_proportions` | - `self` | - `list(list(float))` | ▷ section 4.1.19<br>▷ section 4.1.21 |
| `.content_proportions` | - `self` | - `list(float)` | ▷ section 4.1.20<br>▷ section 4.1.22 |
| `.best_compartment` | - `list(list(int))` | - `list(int)` | ▷ section 4.1.21<br>▷ section 4.1.22 |
| `.proposed_clustering` | - `list(list(float))`<br>- `string`<br>- `float` | - `list(int)` | ▷ section 4.1.22<br>▷ section 4.1.21 |

**4.1.3. Description of `.__init__` (method).** This section describes the code and the functionalities of the method `.__init__`. The method is equipped with the following inputs.

| .__init__ | | |
|---|---|---|
| Inputs | Types | Specifications |
| dimension | int | necessary |
| residual | float | necessary |
| cytosol | list(float) | necessary |
| organelles | list(list(float)) | necessary |

The method possesses one conditional action, which we describe below through sevaral examples.

| Action | |
|---|---|
| Condition | If the length of the list cytosol and the lengths of the lists contained in organelles are all equal to dimension |
| Description | The method initializes the object self.dimension with the value of dimension; the object self.residual with the value of residual; the object self.cytosol with the list stored in cytosol; and the object self.organelles with the list stored in organelles.<br>The method also initializes the object self.K with the *content* [1, Def. 1.21] of the cell, which also corresponds to the output of the method self.content (see section 4.1.7). The method stores in the object self.SK the sum sum(self.K) and stores in the object self.Sorg a list containing the sums sum(self.organelles[i]) for every index i.<br>Finally, the method initializes the objects self.well_defined and self.well_defined_cytosol with boolean values. The former indicates whether the value of self.residual as well as the sum<br>self.residual + sum(self.cytosol),<br>and the values stored in every variable self.organelles[i][u] are non-negative. The latter indicates whether the values stored in every variable self.cytosol[u] are non-negative. |

In the example given below, we define a Cell item through the constructor and we check that the arguments passed to the constructor are stored in the appropriate variables.

```
>>> c = Cell(dimension = 2, residual = 0.5, cytosol = [-2,3], organelles =
[[1,2], [5,0], [8,7]])
>>> print c.dimension
2
>>> print c.residual
0.5
>>> print c.cytosol
[-2, 3]
>>> print c.organelles
[[1, 2], [5, 0], [8, 7]]
```

The values stored in the objects c.K, c.SK and c.Sorg are quantities whose values are updated whenever the parameters of the cell change. This is to allow a quick access to their values instead of going through their re-computation. Specifically,

▷ c.K contains the content of the cell c (see below), which can also be computed by the method c.content() (see section 4.1.7). More specifically, the object c.K contains a list whose element c.K[u], for any given u ranging from 0 to self.dimension-1, is the sum

$$\texttt{c.cytosol[u]} + \sum_{\texttt{i}} \texttt{c.organelles[i][u]}.$$

In [1], the quantity c.K corresponds to the quantity $K(c)$ introduced in [1, Def. 1.21];

- c.SK contains the sum sum(c.K) of the elements of the list c.K. The quantity c.SK corresponds to the quantity $\mathcal{S}K(c)$ used in [1, Def. 1.29];
- c.Sorg contains a list whose elements are the sums of each row of the matrix c.organelles, meaning that each c.Sorg[u] is the sum

$$\sum_{\mathtt{i}} \mathtt{c.organelles[i][u]}.$$

In [1], the quantity c.Sorg corresponds to the quantity $\mathcal{S}\mathrm{org}(c)$ used in [1, Prop. 1.45].

```
>>> print c.K
[12, 12]
>>> print c.SK
24
>>> print c.Sorg
[3, 5, 15]
```

In the display given above, we show the values of c.well_defined and c.well_defined_cytosol. We can see that the cell c is well defined, but the first component of c.cytosol is negative, which may be an important piece of information if one wants to update the parameters of c.cytosol without changing the value of c.well_defined.

```
>>> print c.well_defined
True
>>> print c.well_defined_cytosol
[False, True]
```

**4.1.4. Description of .check (method).** This section describes the code and the functionalities of the method .check, which does not take any input. The method possesses one action, which we describe below through an example.

| Action | |
|---|---|
| Condition | Always |
| Description | The method prints, on the standard output, a list of boolean values indicating whether each of the values given by the object self.residual, the expression<br>self.residual + sum(self.cytosol),<br>and the elements self.cytosol[u] and self.organelles[i][u] is non-negative. |

Here is an example that illustrates the previous description.

```
>>> c = Cell(dimension = 2, residual = 0.5, cytosol = [-4,3], organelles =
[[1,2], [5,0], [8,7]])
>>> c.check()
residual:  True
residual + sum(cytosol):  False
-> cytosol[0]:  False
-> cytosol[1]:  True
organelles[0][0]:  True
organelles[0][1]:  True
organelles[1][0]:  True
organelles[1][1]:  True
organelles[2][0]:  True
organelles[2][1]:  True
```

**4.1.5. Description of `.copy` (method).** This section describes the code and the functionalities of the method `.copy`, which does not take any input. The method possesses one action, which we describe below through an example.

| Action | |
|---|---|
| Condition | Always |
| Description | The method allocates, somewhere else in memory, a new `Cell` item whose parameters are the same as `self`. |

Below, we use the method `.stdout` (see section 4.1.6) to display the parameters of the cells created using `.copy`.

```
>>> c = Cell(dimension = 2, residual = 0.5, cytosol = [-2,3], organelles =
[[1,2], [5,0], [8,7]])
>>> c.stdout()
residual:  0.5
cytosol[0]:  -2
cytosol[1]:  3
1-th organelle:  [1, 2]
2-th organelle:  [5, 0]
3-th organelle:  [8, 7]
>>> d = c.copy()
>>> c.stdout()
residual:  0.5
cytosol[0]:  -2
cytosol[1]:  3
1-th organelle:  [1, 2]
2-th organelle:  [5, 0]
3-th organelle:  [8, 7]
```

**4.1.6. Description of `.stdout` (method).** This section describes the code and the functionalities of the method `.stdout`, which does not take any input. The method possesses one action, which we describe below through an example.

| Action | |
|---|---|
| Condition | Always |
| Description | The method prints the values stored in the variables `self.residual`, `self.cytosol[u]` and `self.organelles[i][u]` on the standard output. |

As shown in the following example, we can display the parameters of a cell by calling the method `.stdout`.

```
>>> c = Cell(dimension = 2, residual = 0.5, cytosol = [-2,3], organelles =
[[1,2], [5,0], [8,7]])
>>> c.stdout()
residual:  0.5
cytosol[0]:  -2
cytosol[1]:  3
1-th organelle:  [1, 2]
2-th organelle:  [5, 0]
3-th organelle:  [8, 7]
```

**4.1.7. Description of `.content` (method).** This section describes the code and the functionalities of the method `.content`, which does not take any input. The method possesses one action, which we describe below through sevaral examples.

| Action | |
|---|---|
| Condition | Always |
| Description | The method returns a list `the_content`, of length `self.dimension`, whose element `the_content[u]` is the sum of the value contained in `self.cytosol[u]` with the values contained in `self.organelles[0][u]`, `self.organelles[1][u]`, ... and `self.organelles[self.dimension-1][u]`. |

We illustrate the use of `.content` through the following three examples.

```
>>> c = Cell(dimension = 2, residual = 16.5, cytosol = [-2,3], organelles =
[[1,2], [5,0], [8,7]])
>>> print c.content()
[12, 12]
>>> c = Cell(dimension = 2, residual = 16.5, cytosol = [0,0], organelles =
[[0,1], [1,0], [9,2]])
>>> print c.content()
[9, 4]
```

Note that the content of a `Cell` item will contain negative values if the values of the list `.cytosol` are negative and large enough.

```
>>> c = Cell(dimension = 2, residual = 16.5, cytosol = [-18,2], organelles
= [[0,1], [1,0], [9,2]])
>>> print c.content()
[-8, 5]
```

One way to prevent such a scenario is to make sure that the values contained in `.cytosol` do not go below a certain threshold. This can be done through the method

`.spontaneous_reaction`

(see section 4.1.12), which sets the values contained in `.cytosol` to 0, as shown below.

```
>>> c.spontaneous_reaction()
>>> print c.content()
[10, 3]
>>> c.stdout()
residual:  0.5
cytosol[1]:  0
cytosol[2]:  0
1-th organelle:  [0, 1]
2-th organelle:  [1, 0]
3-th organelle:  [9, 2]
```

**4.1.8. Description of .compose (method).** This section describes the code and the functionalities of the method .compose. The method is equipped with two inputs.

| .compose | | |
|---|---|---|
| Inputs | Types | Specifications |
| index | int | necessary |
| a_cell | Cell | necessary |

The method possesses one conditional action, which we describe below through an example.

| Action 1 | |
|---|---|
| Condition | If a_cell.dimension is equal to self.dimension |
| Description | The method returns the operadic composition of the Cell item self with the Cell item a_cell at the index-th organelle (see [1, Def. 1.6]), namely the cell whose dimension is self.dimension, whose residual is the sum self.residual + a_cell.residual, whose cytosol is the list containing the sums self.cytosol[u] + a_cell.cytosol[u] and whose lists of organelles is the list obtained from replacing the (1+index)-th index of the list self.organelles with the entire list of organelles contained in a_cell.organelles. |

The following example shows how the method .compose can be used to compose cells at a given organelle. The idea is to mix the organelles and the cytosol of the "inner cell" with the organelles and the cytosol of the "outer cell".

```
>>> outer_cell = Cell(dimension = 2, residual = 0.5, cytosol = [-2,3],
organelles = [[1,2], [5,0], [8,7]])
>>> inner_cell = Cell(dimension = 2, residual = 0.5, cytosol = [0,0],
organelles = [[0,1], [1,0], [9,2]])
>>> comp = outer_cell.compose(1,inner_cell)
```

Below, we can use the method .stdout to see that the organelles of the inner cell constitutes the organelles of the composite cell from the 2nd organelle to 4-th organelle.

```
>>> comp.stdout()
residual:  1.0
cytosol[1]:  -2
cytosol[2]:  3
1-th organelle:  [1, 2]
2-th organelle:  [0, 1]
3-th organelle:  [1, 0]
4-th organelle:  [9, 2]
5-th organelle:  [8, 7]
```

**4.1.9. Description of .compose_all (method).** This section describes the code and the functionalities of the method .compose_all. The method is equipped with the following input variable.

| .compose_all | | |
|---|---|---|
| Inputs | Types | Specifications |
| cells | list(Cell) | necessary |

The method possesses one conditional action, which we describe below through an example.

| Action 1 | |
|---|---|
| Condition | If the length of the list `cells` is equal to `len(self.organelles)` |
| Description | The method returns the operadic composition of the `Cell` item `self` with each of the `Cell` items contained in `cells` for their corresponding indices in `cells`. The returned `Cell` item corresponds to a simultaneous composition of cells, as defined in [1, Def. 1.14] |

Below, we show an example of a simultaneous composition of cells.

```
>>> outer_cell = Cell(dimension = 2, residual = 0.5, cytosol = [-2,3],
organelles = [[1,2], [5,0], [8,7]])
>>> incell_1 = Cell(dimension = 2, residual = 0.5, cytosol = [0,0],
organelles = [[0,1], [1,0], [9,2]])
>>> incell_2 = Cell(dimension = 2, residual = 0.5, cytosol = [0,0],
organelles = [[10,1]])
>>> incell_3 = Cell(dimension = 2, residual = 0.5, cytosol = [0,0],
organelles = [[2,2],[22,0]])
>>> comp = outer_cell.compose_all([incell_1,incell_2,incell_3])
```

As shown below, the organelles of the "outer cell" were all replaced with the list of organelles of the "inner cells".

```
>>> comp.stdout()
residual:  2.0
cytosol[1]:  -2
cytosol[2]:  3
1-th organelle:  [0, 1]
2-th organelle:  [0, 1]
3-th organelle:  [9, 2]
4-th organelle:  [10, 1]
5-th organelle:  [2, 2]
6-th organelle:  [22, 0]
```

**4.1.10. Description of `.left` (method).** This section describes the code and the functionalities of the method `.left`. The method is equipped with the following input variables.

| `.left` | | |
|---|---|---|
| Inputs | Types | Specifications |
| `alpha_var` | `list(float)` | necessary |
| `kappa_var` | `list(list(float))` | necessary |
| `lambda_var` | `list(list(float))` | necessary |

The method possesses one conditional action, which we describe below through an example.

| Action | |
|---|---|
| Condition | If the lengths of the lists alpha_var, kappa_var, and lambda_var are equal to each other |
| Description | If we let $c$ represents the Cell item self and we let $\alpha$, $\kappa$ and $\lambda$ represent the three inputs lists, then the method returns the Cell item $\mathsf{left}(c)(\alpha,\kappa,\lambda)$, as defined in [1, Conv. 1.19]. This Cell item constitutes half of a solution for a factorization problem expressing self as a simultaneous composition of the form: $$c = \mathsf{left}(c)(\alpha,\kappa,\lambda) \circ \mathsf{right}(c)(\mu,\alpha,\kappa).$$ The other half $\mathsf{right}(c)(\mu,\alpha,\kappa)$ is accessible through the method self.right (see section 4.1.11) and returns a list of Cell items. The output of self.left(alpha_var,kappa_var,lambda_var) is a Cell item whose dimension is equal to self.dimension, whose residual is equal to the difference between self.residual and sum(alpha_var), whose cytosol is the difference between self.cytosol the total sum of the coefficients of the matrix kappa_var and whose list of organelles is lambda_var. |

The example given below uses the same Cell item as well as the same set of inputs alpha_var and kappa_var as those used in section 4.1.11, where we describe the other half of the factorization returned by the method .right.

```
>>> c = Cell(dimension = 2, residual = 0.5, cytosol = [-2,3], organelles =
[[1,2], [5,0], [8,7]])
>>> left = c.left([0,0.125],[[1,0],[0,2]],[[2,5],[0,7]])
>>> left.stdout()
residual:  0.375
cytosol[1]:  -3
cytosol[2]:  1
1-th organelle:  [2, 5]
2-th organelle:  [0, 7]
```

**4.1.11. Description of .right (method).** This section describes the code and the functionalities of the method .right. The method is equipped with the following input variables.

| .right | | |
|---|---|---|
| Inputs | Types | Specifications |
| mu_var | list(float) | necessary |
| alpha_var | list(float) | necessary |
| kappa_var | list(list(float)) | necessary |

The method possesses one conditional action, which we describe below through an example. Note that the first condition given in the following table is justified by the factorization theorem stated in [1, Th. 1.18].

| Action | |
|---|---|
| Condition | If the sum sum(mu_var) is equal to the length of the list self.organelles and the lengths of the lists lambda_var, alpha_var, and kappa_var are equal to each other |
| Description | If we let $c$ represents the Cell item self and we let $\mu$, $\alpha$ and $\kappa$ represent the three input lists, then the method returns the list $\mathsf{right}(c)(\mu,\alpha,\kappa)$ of Cell items, as defined in [1, Conv. 1.19]. This Cell item constitutes half of a solution for a factorization problem expressing self as a simultaneous composition of the form: $$c = \mathsf{left}(c)(\alpha,\kappa,\lambda) \circ \mathsf{right}(c)(\mu,\alpha,\kappa).$$ The other half $\mathsf{left}(c)(\alpha,\kappa,\lambda)$ is accessible through the method self.left (see section 4.1.10) and returns a Cell item.<br>Each element self.right(mu_var,alpha_var,kappa_var)[i] is a Cell item whose dimension is equal to self.dimension, whose residual is equal to alpha_var[i], whose cytosol is equal to the list kappa_var[i] and whose list of organelles is given by the sublist of self.organelles ranging from the index L = sum(mu_var[:i]) to index L+mu_var[i]-1. |

The example given below uses the same Cell item as well as the same set of inputs alpha_var and kappa_var as those used in section 4.1.10, where we describe the other half of the factorization returned by the method .left.

```
>>> c = Cell(dimension = 2, residual = 0.5, cytosol = [-2,3], organelles =
[[1,2], [5,0], [8,7]])
>>> right = c.right([1,2],[0,0.125],[[1,0],[0,2]])
>>> for r in right:
...     r.stdout()
...     print "--"
...
residual:  0
cytosol[1]:  1
cytosol[2]:  0
1-th organelle:  [1, 2]
--
residual:  0.125
cytosol[1]:  0
cytosol[2]:  2
1-th organelle:  [5, 0]
2-th organelle:  [8, 7]
--
```

**4.1.12. Description of .spontaneous_reaction (method).** This section describes the code and the functionalities of the method .spontaneous_reaction, which does not take any input. The method possesses one action, which we describe below through an example.

| Action | |
|---|---|
| Condition | Always |
| Description | The method returns the 'cleaning' [1, Def. 2.13] of the `Cell` item `self`. This cleaning is meant to simulate an overall chemical reaction occurring in the cytosol of the cell, as described in [1, Rem. 1.31].<br>Specifically, the method adds the value `sum(self.cytosol)` to `self.residual` while it subtracts it from `self.SK`. If the value of `self.residual` becomes negative after that change, then the function exits the program with an error message. In the case where no error message is returned, the method goes on and subtracts the value `self.cytosol[u]` from the value `self.K[u]` for every index `u`. Finally, the method sets the values of the list `self.cytosol` to 0. |

The idea behind the "overall chemical reaction" computed by `.spontaneous_reaction` is to convert the positive values of the cytosol, which can be seen as available resources, into numbers that would compensate the negative values of the list `.cytosol`. These negative values can be seen as used or missing resources that needs to be compensated in order to re-establish equilibirum inside the cell. In the case where the "available resources" are not enough to compensate the "missing resources", we want to use the value stored in the object `.residual` as an alternative resource. This value can be interpreted as the energetic surplus of the system. The product of this reaction is the sum `sum(self.cytosol)`, which can be seen as the energy remaining after the global reaction.

`residual+ positive_cytosol` → `sum(cytosol) + negative_cytosol`

In the following example, we show that the method `.spontaneous_reaction` changes the values of the content `.K` (see section 4.1.3), its sum `.SK` and the value of the object `.residual`.

```
>>> c = Cell(dimension = 2, residual = 0.5, cytosol = [-2,3], organelles =
[[1,2], [5,0], [8,7]])
>>> print c.SK
24
>>> print c.K
[12, 12]
>>> c.spontaneous_reaction()
>>> c.stdout()
residual:  1.5
cytosol[1]:  0
cytosol[2]:  0
1-th organelle:  [1, 2]
2-th organelle:  [5, 0]
3-th organelle:  [8, 7]
>>> print c.SK
23
>>> print c.K
[14, 9]
```

**4.1.13. Description of `.action` (method).** This section describes the code and the functionalities of the method `.action`. The method is equipped with the following input variable.

| `.action` | | |
|---|---|---|
| Inputs | Types | Specifications |
| `matrix` | `list(list(float))` | necessary |

The method possesses one conditional action, which we describe below through an example.

| Action | |
|---|---|
| Condition | If the length of matrix is equal to len(self.organelles) and the length of each lists in matrix is equal to self.dimension |
| Description | The method returns the action of the Cell item self on the matrix, as defined in [1, Def. 1.29]. More specifically, the method returns a list the_action of length self.dimension whose element the_action[u] is the sum of the values (self.organelles[i][u]/self.SK) * matrix[i][u] over every index i ranging from 0 to len(self.organelles)-1. By definition of the object self.SK (see section 4.1.3), the ratio self.organelles[i][u]/self.SK is smaller than 1 when sum(self.cytosol) is positive. In this case, we can interpret the output the_action as a vector of conditional expected values (see the formula given in [1, Def. 1.29]). |

The following example shows how to use the method .action.

```
>>> c = Cell(dimension = 2, residual = 0.5, cytosol = [-2,3], organelles =
[[1,2], [5,0], [8,7]])
>>> a = [[70,9],[1,5],[5,81]]
>>> print c.action(a)
[4.791666666666666, 24.375]
```

**4.1.14. Description of .algebra_operator (method).** This section describes the code and the functionalities of the method .algebra_operator. The method is equipped with the following input variable.

| .algebra_operator | | |
|---|---|---|
| Inputs | Types | Specifications |
| matrix | list(list(float)) | necessary |

The method possesses one conditional action, which we describe below through an example.

| Action | |
|---|---|
| Condition | If the length of matrix is equal to len(self.organelles) and the length of each lists in matrix is equal to self.dimension |
| Description | If we let $c$ represent self and we let $a$ represent the input matrix, the method returns the value of the algebra operator $U(c,d)(a)$ [1, Def. 1.33], where $d$ is inferred from the organelles of $c$. More specifically, we use the formula of [1, Prop. 1.35 (1.4)], for which we replace each coefficient $(d_k \cdot a_k)_u$ shown therein with the element matrix[k][u]. In other words, the output of the method .algebra_operator is the sum of the terms (dividend[k]/self.SK) * matrix[k][u], over every index k ranging from 0 to len(self.organelles)-1, where dividend[k] is equal to (self.Sorg[k]-self.organelles[k][u]). |

The following example shows how to use the method .algebra_operator.

```
>>> c = Cell(dimension = 2, residual = 0.5, cytosol = [-2,3], organelles =
[[1,2], [5,0], [8,7]])
>>> a = [[70,9],[1,5],[5,81]]
>>> print c.algebra_operator(a)
[7.291666666666666, 28.416666666666668]
```

As explained in the description above, the method `.algebra_operator` returns the values of a formula that is more general than the one given [1, Prop. 1.35 (1.4)]. To recover the formula given therein, we would need to give to the method `.algebra_operator`, the outputs of the method `.action` for a list of cells `d[1]`, . . . , `d[len(self.organelles)]`. For instance, the method `.allostasis` (section 4.2.12) associated with the class `SuperCell` uses the method `.algebra_operator` in this way.

**4.1.15. Description of `.allostasis` (method).** This section describes the code and the functionalities of the method `.allostasis`. The method is equipped with the following input variables.

| .allostasis | | |
|---|---|---|
| Inputs | Types | Specifications |
| `matrix` | `list(list(float))` | necessary |
| `weight` | `list(float)` | necessary |
| `org_index` | `int` | necessary |
| `dim_index` | `int` | necessary |

The method possesses one conditional action, which we describe below through an example.

| Action | |
|---|---|
| Condition | If the length of `matrix` is equal to `len(self.organelles)` and the length of the list `matrix[org_index]` is equal to `self.dimension` |
| Description | If we let $c$ represent `self`, we let $a$ represent the input `matrix` and we let $j$ and $v$ denote the inputs `org_index` and `dim_index`, the method returns the value of the "allostatic differential" $\partial_{j,v}U(c,a)/\partial(c,d)(a)$ [1, Def. 1.47], where $d$ is inferred from the organelles of $c$. More specifically, we use the formula given in [1, Prop. 1.48], for which we replace each coefficient $U(c,d)(a)_u$ shown in the formula therein with the element `weight[u]`. In other words, the output of the method `.allostasis` is the sum of the terms<br>`- (ratio[u] / self.SK) * (term1 - term2[u]),`<br>where `ratio[u]` is equal to the ratio<br>`self.organelles[org_index][u]/self.Sorg[org_index],`<br>where `term1` is equal to the multiplication `matrix[org_index][dim_index] * weight[dim_index]` and where `term2[u]` is equal to the multiplication `matrix[org_index][u] * weight[u]` over every index `u` ranging from 0 to `self.dimension-1`. |

In the following example, we illustrate the use of the method `.allostasis` in the case where the input variable `weight` is the vector $U(c,d)(a)_u$. In other words, the vector `weight` contains the output of the function `c.algebra_operator(a)` (see section 4.1.14). This vector corresponds to the list of values that one is supposed to use with `.allostasis` in order to satisfy the formula given in [1, Prop. 1.48].

```
>>> c = Cell(dimension = 2, residual = 0.5, cytosol = [-2,3], organelles =
[[1,2], [5,0], [8,7]])
>>> a = [[70,9],[1,5],[5,81]]
>>> for j in range(len(c.organelles)):
...     for v in range(c.dimension):
...       print (j, v), c.allostasis(a,c.algebra_operator(a),j,v)
(0, 0) -7.07407407407
(0, 1) 3.53703703704
(1, 0) -0.0
(1, 1) -5.61631944444
(2, 0) 44.047337963
(2, 1) -50.3398148148
```

**4.1.16. Description of `.agreement` (method).** This section describes the code and the functionalities of the method `.agreement`. The method is equipped with the following three input variables, including one that is optional.

| .agreement | | |
|---|---|---|
| Inputs | Types | Specifications |
| `index` | `int` | necessary |
| `vector` | `list(float)` | necessary |
| `*challenge` | `list(float)` | optional |

The method possesses two actions, which we describe below through sevaral examples.

| Action 1 | |
|---|---|
| Condition | If `challenge` is empty (absent from the list of given inputs) |
| Description | The method returns the agreement of the underlying `Cell` item `self` with the input `vector`, as defined in [1, Def. 2.22]. In other words, the method computes the normalized scalar product of the vectors `self.organelles[index]` and `vector`. When the list `vector` only contains non-negative values, the output of the method is a `float` item between `0.0` and `1.0`. |

The following example shows how to use the method `.agreement`.

```
>>> c = Cell(dimension = 2, residual = 0.5, cytosol = [-2,3], organelles =
[[1,2], [5,0], [8,7]])
>>> a = [[70,9],[1,5],[5,81]]
>>> for j in range(len(c.organelles)):
...     print j,"-->",c.agreement(j,a[j])
...
0 --> 0.557621357192
1 --> 0.196116135138
2 --> 0.703620697651
```

We now describe the second action of the method `.agreement`.

| Action 2 | |
|---|---|
| Condition | If `challenge` is given and contains a unique element (meant to be a list of length 3 or more, containing `int` values ) |
| Description | The method returns the agreement of the underlying `Cell` item `self` with the list `v` of length `len(self.dimension)` such that `v[u]` is equal to the coefficient `vector[u]` if the following condition is satisfied<br>`challenge[0][0] <= u % challenge[0][2] <= challenge[0][1]`<br>and `v[u]` is equal to 0 otherwise. |

The following example shows how to use the method `.agreement`.

```
>>> c = Cell(dimension = 2, residual = 0.5, cytosol = [-2,3], organelles =
[[1,2], [5,0], [8,7]])
>>> a = [[70,9],[1,5],[5,81]]
>>> challenge1 = []
>>> for j in range(len(c.organelles)):
...     print j,"-->",c.agreement(j,a[j],*challenge1)
...
0 --> 0.557621357192
1 --> 0.196116135138
2 --> 0.703620697651
>>> challenge2 = [[0,0,2]]
>>> for j in range(len(c.organelles)):
...     print j,"-->",c.agreement(j,a[j],*challenge2)
...
0 --> 1.0
1 --> 1.0
2 --> 1.0
>>> challenge3 = [[1,1,2]]
>>> for j in range(len(c.organelles)):
...     print j,"-->",c.agreement(j,a[j],*challenge3)
...
0 --> 1.0
1 --> 0
2 --> 1.0
```

**4.1.17. Description of `.merge` (method).** This section describes the code and the functionalities of the method `.merge`. The method is equipped with the following two inputs, including one that is optional.

| `.merge` | | |
|---|---|---|
| Inputs | Types | Specifications |
| `list_of_organelles` | `list(list(float))` | necessary |
| `order = "order-sorted"` | `string` | optional |

The method possesses one action, which we describe below through sevaral examples.

| Action | |
|---|---|
| Condition | Always |
| Description | The method merges the organelles of self whose indices are in list_of_organelles. The organelles are merged into a unique organelle, which, after the merging, can be found at the index min(list_of_organelles). The method gives the option to terminate faster if the list list_of_organelles is already sorted. We can do so by passing the string "sorted" to the second input variable order. |

The following two examples show how to use the method .merge.

```
>>> c = Cell(dimension = 2, residual = 0.5, cytosol = [-2,3], organelles =
[[1,2], [5,0], [8,7], [4,27]])
>>> c.merge([0,2],order = "sorted")
>>> c.stdout()
residual:  0.5
cytosol[1]:  -2
cytosol[2]:  3
1-th organelle:  [9, 9]
2-th organelle:  [5, 0]
3-th organelle:  [4, 27]
```

Note that the method directly acts on the underlying Cell item self, meaning that the original item is lost after application of .merge.

```
>>> c.merge([2,1])
>>> c.stdout()
residual:  0.5
cytosol[1]:  -2
cytosol[2]:  3
1-th organelle:  [9, 9]
2-th organelle:  [9, 27]
```

**4.1.18. Description of .divide (method).** This section describes the code and the functionalities of the method .divide. The method is equipped with the following two input variables, including one that is optional.

| .divide | | |
|---|---|---|
| Inputs | Types | Specifications |
| list_of_organelles | list(list(float)) | necessary |
| order = "order-sorted" | string | optional |

The method possesses one conditional action, which we describe below through an example.

| Action | |
|---|---|
| Condition | If the elements of the list self.cytosol are all zero values |
| Description | The method divides self into two Cell items c1 and c2 where c1 contains all the organelles of self whose indices are in list_of_organelles and c2 contains all the other organelles of self. The residual of both cells c1 and c2 is equal to half of self.residual and their cytosols are encoded by lists of zero values. The method gives the option to terminate faster if the list list_of_organelles is already sorted. We can do so by giving the string "sorted" to the second input variable order. |

The following example shows how to use the method .divide. Note that the method will output an error message if the cytosol of the cell is not made of zero values. As shown below, we can change the cytosol to a list of zero values by using the method `.spontaneous_reaction` (see section 4.1.12).

```
>>> c = Cell(dimension = 2, residual = 0.5, cytosol = [-2,3], organelles =
[[1,2], [5,0], [8,7], [4,27]])
>>> c1, c2 = c.divide([0,2],order = "order-sorted")
Error:  in Cell.divide:  cytosol is not zero -- cannot divide
>>> c.spontaneous_reaction()
>>> c1, c2 = c.divide([0,2],order = "order-sorted")
>>> c1.stdout()
residual:  0.75
cytosol[1]:  0
cytosol[2]:  0
1-th organelle:  [1, 2]
2-th organelle:  [8, 7]
>>> c2.stdout()
residual:  0.75
cytosol[1]:  0
cytosol[2]:  0
1-th organelle:  [5, 0]
2-th organelle:  [4, 27]
```

**4.1.19. Description of `.organelle_proportions` (method).** This section describes the code and the functionalities of the method `.organelle_proportions`, which does not take any input. The method possesses one action, which we describe below through an example.

| Action | |
|---|---|
| Condition | Always |
| Description | The method outputs a `list(list(float))` item `prop` whose element `prop[i][u]` is the ratio `self.organelles[i][u]/self.Sorg[i]`. |

The following example shows what the outputs of the method `.organelle_proportions` look like for a given cell.

```
>>> c = Cell(dimension = 2, residual = 0.5, cytosol = [-2,3], organelles =
[[1,2], [5,0], [8,7]])
>>> prop = c.organelle_proportions()
>>> for i in range(len(prop)):
...     print prop[i]
...
[0.3333333333333333, 0.6666666666666666]
[1.0, 0.0]
[0.5333333333333333, 0.4666666666666667]
[0.12903225806451613, 0.8709677419354839]
```

**4.1.20. Description of `.content_proportions` (method).** This section describes the code and the functionalities of the method `.content_proportions`, which does not take any input. The method possesses one action, which we describe below through an example.

| Action | |
|---|---|
| Condition | Always |
| Description | The method outputs a list(list(float)) item prop whose element prop[i] is the ratio self.Sorg[i]/sum(self.Sorg). This type of ratios corresponds to the type of ratios shown in [1, Prop. 1.54] provided that certain fitting conditions hold.<br>Specifically, if the organelles of the Cell item self were equal to the contents of a collection of cells $d_1, d_2, \dots, d_n$ and if the cytosol of self was only made of zero values, then the ratio prop[i] returned by the method .content_proportions would correspond to the ratio<br>$$K(d_i)/K(d_1 \otimes d_2 \otimes \cdots \otimes d_n).$$<br>Since the tensoring of cells is associative, the previous type of ratios corresponds to the type of ratios used in [1, Prop. 1.54].<br>This correspondence between the outputs of .content_proportions and the previous ratios is used by some of the methods associated with the class SuperCell (see section 4.2). Very often, these methods use .content_proportions with the method .spontaneous_reaction in order to clear the cytosol of the underlying cell and make the previous correspondence hold. |

The following example shows what the outputs of the method .content_proportions look like for a given cell.

```
>>> c = Cell(dimension = 2, residual = 0.5, cytosol = [-2,3], organelles =
[[1,2], [5,0], [8,7]])
>>> prop = c.content_proportions()
>>> for i in range(len(prop)):
...     print prop[i]
...
0.0555555555556
0.0925925925926
0.277777777778
0.574074074074
```

**4.1.21. Description of .best_compartment (method).** This section describes the code and the functionalities of the method .best_compartment. The method is equipped with the following input variable.

| .best_compartment | | |
|---|---|---|
| Inputs | Types | Specifications |
| cliques | list(list(int)) | necessary |

The method possesses one action, which we describe below through sevaral examples.

| Action | |
|---|---|
| Condition | Always |
| Description | The method `.best_compartment` implements the selection criterion described in [1, Def. 2.32]. The goal of this criterion is to choose among all the lists of the input variable `cliques` a list that contains indices of organelles whose gathering would theoretically minimize the values of the algebra operator (section 4.1.14). As described in [1, Def. 2.32], the idea is to return one of the lists `cliques[k]` for which a certain associated sum `result_k` of terms of the form<br>(4.1) `(dividend1 * divisor2)/float(dividend2 * divisor1)`<br>is maximal. More specifically, if we let `compl[k]` denote the complement of `cliques[k]` in<br>`range(len(self.organelles))`,<br>then we compute, for every index `u` ranging from `0` to `self.dimension-1`, the following quantities:<br>▷ `dividend1 = sum(self.organelles[i][u] for i in cliques[k])`<br>▷ `divisor1 = sum(self.Sorg[i] for i in cliques[k])`<br>▷ `dividend2 = sum(self.organelles[i][u] for i in compl[k])`<br>▷ `divisor2 = sum(self.Sorg[i] for i in compl[k])`<br>and we add ratio (4.1) to the variable `result_k` (initially equal to 0) whenever ratio (4.1) is greater than `1`. We then use the resulting value as a score to assess which of the list of organelles of `cliques` should be compartmentalized in a new cell in order to prominently minimize the values of the algebra operator (section 4.1.14).<br>To better understand the previous statement, first observe that the ratios `dividend1/divisor1` and `dividend2/divisor2` correspond to the ratios that the method `.organelle_proportions` (section 4.1.19) would return if `self` was a `Cell` item with two organelles such that:<br>1) one organelle is encoded by the vector sum of those organelles whose indices are in `cliques[k]`<br>2) the other organelle is the sum of all the remaining organelles of `self`.<br>Our interest in these two ratios lies in the statement of [1, Th. 1.57] and its interpretation given in [1, Rem. 1.58]. More specifically, the method is to be used when one wants to separate the organelles of `self` within two (potentially temporary) cells $d_1$ and $d_2$. The indices contained in the list `cliques[k]` are the indices of those organelles of `self` contained in the cell $d_1$ while the indices in `compl[k]` are the indices of those organelles of `self` contained in the cell $d_2$. In addition, if the two cells $d_1$ and $d_2$ were assumed to *fit* [1, Def. 1.23] the organelles of some third cell $c$ – meaning that the content of $d_1$ (section 4.1.7) equals the vector $\texttt{org}(c)_1$ and the content of $d_2$ equals the vector $\texttt{org}(c)_1$ – then the method `.best_compartment` can be understood as follows: |

| Description | $(i)$ it first computes the sum of real values resulting from the quotient of the quantity $\mathsf{org}(c)_{1,u}/\mathcal{S}\mathsf{org}(c)_1$ over the quantity $\mathsf{org}(c)_{2,u}/\mathcal{S}\mathsf{org}(c)_2$ whenever inequality (4.2) is satisfied for the index $u$; (4.2) $$\frac{\mathsf{org}(c)_{1,u}}{\mathcal{S}\mathsf{org}(c)_1} > \frac{\mathsf{org}(c)_{2,u}}{\mathcal{S}\mathsf{org}(c)_2};$$ $(ii)$ it then returns one of the lists `cliques[k]` whose associated sum of quotients, computed at step $(i)$, is maximal. Thus, the method is looking for a list `cliques[k]` for which inequality (4.2) seems to prominently hold over the list of indices $u$. The output of the method `.best_compartment` is used by the method `.proposed_clustering` (section 4.1.22) in order to know which of the proposed list `cliques[k]` is the best suited for creating a new compartment within the cell. |
|---|---|

Below, we give four examples illustrating the use of the method `.best_compartment`.

```
>>> c = Cell(dimension = 3,residual = 0.5,cytosol = [-2,3,0],organelles =
[[1,10,1],[1,1,10],[10,1,1],[20,1,1],[1,20,1]])
>>> print c.best_compartment([[0,1],[0,3],[1,2],[1,3],[2,3]])
[1, 2]
>>> print c.best_compartment([[0,1,3],[1,2,3],[2,3]])
[2, 3]
>>> c = Cell(dimension = 3,residual = 0.5,cytosol = [-2,3,0],organelles =
[[1,10,1],[1,1,10],[10,1,1],[20,1,1],[1,20,1]])
>>> print c.best_compartment([[0,4],[1,4],[2,4],[3,4]])
[0, 4]
>>> print c.best_compartment([[0,3],[0,4],[0,1],[0,2]])
[0, 4]
>>> print c.best_compartment([[2,3],[2,1],[2,4],[2,0]])
[2, 3]
```

**4.1.22. Description of `.proposed_clustering` (method).** This section describes the code and the functionalities of the method `.proposed_clustering`. The method is equipped with the following three input variables, including one that is optional.

| .proposed_clustering | | |
|---|---|---|
| Inputs | Types | Specifications |
| `matrix` | `list(list(float))` | necessary |
| `option` | `string` | necessary |
| `filtering = [1.5,0]` | `list(float)` | optional |

The method possesses one conditional action, which we describe below through sevaral examples.

| Action | |
|---|---|
| Condition | If the length of `matrix` is equal to `len(self.organelles)` and the lengths of the lists contained in `matrix` are equal to `self.dimension` |
| Description | The method implements the clustering algorithm discussed throughout [1, Sec. 2.5]. Depending on the value of the input `option` (which corresponds to the parameter $\varepsilon$ in [1, Sec. 2.5]), the method computes the lists of indices `i` for which the values `matrix[i][u]` are either greater or less (by a factor equal to the value of `filtering[0]`) than the value<br>$$\texttt{barycenter[u]} = \sum_{\texttt{k=0}}^{n} \texttt{matrix[k][u] * prop[k]}$$<br>where the list `prop` corresponds to the output of the method `self.content_proportions` (see section 4.1.20) and $n$ denotes the integer `len(self.organelles)`. Note that, because each weight `prop[k]` can be viewed as a probability, the resulting list `barycenter` can be interpreted as a barycenter of the lists `matrix[k]`; see [1, Rem. 1.55] for further explanation.<br>In more detail, the method goes through the following steps. First, it computes a `list(list(float))` item encoding a weighted adjacency $(n \times n)$-matrix `graph` whose coefficient `graph[i][j]` counts the number of indices `u`, ranging from `0` to `self.dimension-1`, for which the two values `matrix[i][u]` and `matrix[j][u]` are<br>(1) either greater than the multiplication `filtering[0] * barycenter[u]` if the input `option` is equal to the string `"division"`;<br>(2) or less than the multiplication `filtering[0] * barycenter[u]` if the input `option` is equal to the string `"merging"`.<br>Here, the reader following the development of [1] may have noticed that the factor `filtering[0]` corresponds to the factor $\nu$ used from [1, Def. 2.29] to [1, Def. 2.32].<br>If `graph` is the zero matrix, then the method `.proposed_clustering` returns the empty list. Otherwise, the method passes the matrix `graph` to the method `usf.cliques` (section 3.1.21), which returns a list `cliques` of the connected components of `graph` such that the weights of the vertices of these connected components are maximal and greater than or equal to `filtering[1]`. In this case, the method `.proposed_clustering` returns the output of the function `self.best_compartment(cliques)` (see section 4.1.21). |

In the spirit of [1, Rem. 1.58], the goal of the method `.proposed_clustering` is to find a set of organelles (specified by their indices) whose gathering in a new `Cell` item will decrease the values of `self.algebra_operator` (see section 4.1.14).

We illustrate the use of the method `.proposed_clustering` in the following examples, in which the method returns the indices of the organelles meant to be separated from the other organelles.

We start with an example illustrating the use of the option `"merging"` together with the value `filtering = [1,0]`. In this case, the method `.proposed_clustering` corresponds to the fusion algorithm of [1, Sec. 2.5] for the parameters $\varepsilon = -$, $\nu = 1$ and $\omega = 0$.

```
>>> c = Cell(dimension = 2,residual = 0.5,cytosol = [-2,3],organelles =
[[1,2],[5,0],[8,7],[4,27]])
>>> a = [[70,9],[1,5],[5,1],[12,45]]
>>> print "maximal clique = " + str(c.proposed_clustering(a,"merging",
filtering=[1,0]))
Searching cliques in the following graph:
[0, 1, 0, 1]
[0, 0, 1, 2]
[0, 0, 0, 1]
[0, 0, 0, 0]
Maximal weight = 2
Edges with maximal weight = [[1, 3]]
maximal clique = [1, 3]
```

Note that, contrary to the method best_compartment (section 4.1.21), both the values of the organelles and the values of the input influence the choice of the output of the method .proposed_clustering. This is illustrated below for the option "division" and the value filtering = [1,0]. In this case, the method proposed_clustering corresponds to the fission algorithm of [1, Sec. 2.5] for the parameters $\varepsilon = +$, $\nu = 1$ and $\omega = 0$.

```
>>> c = Cell(dimension = 3,residual = 0.5,cytosol = [-2,3,0],organelles =
[[1,10,1],[1,1,10],[10,1,1],[20,1,1],[1,20,1]])
>>> a = [[1,10,1],[1,1,10],[10,1,1],[20,1,1],[1,20,1]]
>>> print "maximal clique = " + str(c.proposed_clustering(a,"division",
filtering=[1,0]))
Searching cliques in the following graph:
[0, 0, 0, 0, 1]
[0, 0, 0, 0, 0]
[0, 0, 0, 1, 0]
[0, 0, 0, 0, 0]
[0, 0, 0, 0, 0]
Maximal weight = 1
Edges with maximal weight = [[0, 4], [2, 3]]
maximal clique = [0, 4]
>>> a = [[1,10,1],[1,1,10],[10,1,1],[20,1,1],[1,20,10]]
>>> print "maximal clique = " + str(c.proposed_clustering(a,"division",
filtering=[1,0]))
Searching cliques in the following graph:
[0, 0, 0, 0, 1]
[0, 0, 0, 0, 1]
[0, 0, 0, 1, 0]
[0, 0, 0, 0, 0]
[0, 0, 0, 0, 0]
Maximal weight = 1
Edges with maximal weight = [[0, 4], [1, 4], [2, 3]]
maximal clique = [0, 1, 4]
>>> a = [[1,10,1],[1,1,10],[10,1,1],[20,1,1],[10,20,10]]
>>> print "maximal clique = " + str(c.proposed_clustering(a,"division",
filtering=[1,0]))
Searching cliques in the following graph:
[0, 0, 0, 0, 1]
[0, 0, 0, 1, 0]
```

```
[0, 0, 0, 0, 1]
[0, 0, 0, 0, 1]
[0, 0, 0, 0, 0]
Maximal weight = 1
Edges with maximal weight = [[0, 4], [2, 3], [2, 4], [3, 4]]
maximal clique = [0, 2, 3, 4]
>>> a = [[1,10,1],[1,1,10],[10,1,1],[20,1,1],[15,20,10]]
>>> print "maximal clique = " + str(c.proposed_clustering(a,"division",
filtering=[1,0]))
Searching cliques in the following graph:
[0, 0, 0, 0, 1]
[0, 0, 0, 0, 1]
[0, 0, 0, 0, 0]
[0, 0, 0, 0, 1]
[0, 0, 0, 0, 0]
Maximal weight = 1
Edges with maximal weight = [[0, 4], [1, 4], [3, 4]]
maximal clique = [0, 1, 3, 4]
```

#### 4.2. Description of SuperCell (class)

**4.2.1. Introduction.** This section introduces the reader to the code of the class SuperCell whose main goal is to model the features of a super cell, as defined in [1, Def. 2.1]. Before presenting the features of the class, recall that, mathematically, a super cell of some dimension $N$, of some height $q$ and equipped with $n$ organelles, consists of a tree structure labeled by cells of dimension $N$. In this respect, the collection of cells $c, c_1, \dots, c_p, c_{11}, \dots$ defining a super cell $\hat{c}$ are either junctions, meaning nodes possessing child cells, or leaves, meaning nodes without children (see [1, Conv. 2.2 & Def. 2.3]).

$\hat{c}$ : 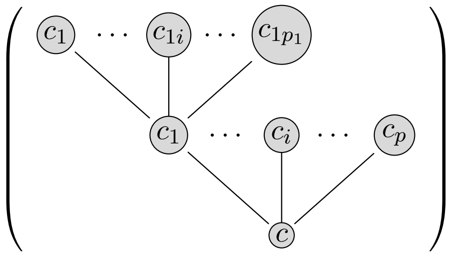

Except for the root cell, shown above at the bottom of the tree, every other cell has a parent cell. Other than the root and the leaves, all cells of a super cell are similar in structure. This similarity allows us to construct our methods as recursive extensions of simpler methods that only act on the *basal junction* of the super cell, namely the root and its junction. In this library, the class SuperCell is equipped with 8 objects, related to the various results of [1, Sec. 2], as well as 3 types of methods, which are listed below:

▷ a set of 2 methods meant to be used for initializing the tree structure of a super cell:

- `set_level`: increments the levels of each super cell/cell by a given value;
- `reset_depth`: recomputes the depth of each super cell/cell within a given super cell;

▷ a single debugging method:

- `stdout`: displays the tree structure of a super cell on the standard output;

▷ a set of 14 methods meant to implement the algorithmic steps of [1, Sec. 2]:

- `__init__`: returns a super cell for its associated set of parameters;
- `compress`: returns the total and recursive composition of all the cells of a super cell;

- `action`: returns the action of a super cell on a vector, as defined in [1, Def. 2.9];
- `base`: returns the basal pruning of the super cell, as defined in [1, Def. 2.10];
- `compute_variables`: computes parameters for the operations right and left; as defined in [1, Conv. 1.19];
- `homeostasis`: solve a homeostasis problem for the basal pruning of a super cell; for more information about this operation, see [1, Def. 1.39];
- `allostasis`: implements the algorithmic step of [1, Sec. 2.3];
- `spontaneous_reaction`: returns the cleaning of the super cell – see [1, Def. 2.16];
- `tensor_pre_action`: computes a barycenter of pre-actions – this is explained in more detail in [1, Rem 1.55 & Sec. 2.5];
- `merge_base`: merges the child super cells of the basal junction of a super cell, as described in [1, Sec. 2.5];
- `divide_base`: divides the child super cells of the root of a super cell, as described in [1, Sec. 2.5];
- `fusion`: implements the algorithmic step of [1, Sec. 2.5];
- `fission`: implements the algorithmic step of [1, Sec. 2.5];
- `compose`: implements the algorithmic step of [1, Sec. 2.4].

**4.2.2. Structure.** The following tables give a preview of the class `Cell`. The table given below describes the various dependencies of the class.

| Dependencies | |
|---|---|
| Superclass ancestry | Module section |
| `object` | N/A |
| Statistics | |
| ▷ Importable objects: 7 | |
| ▷ Non-importable objects: 0 | |
| ▷ Importable methods: 17 | |
| ▷ Non-importable methods: 2 | |

The following table gives a description of the 7 importable objects of the class.

| Objects | | |
|---|---|---|
| Name | Type | Related sections |
| `.depth` | `int` | ▷ section 4.2.3 |
| `.level` | `int` | ▷ section 4.2.3 |
| `.cell` | `Cell` | ▷ section 4.2.3 |
| `.is_leaf` | `bool` | ▷ section 4.2.3 |
| `.compose_state` | `bool` | ▷ section 4.2.3 |
| `.pre_action` | `list(float)` | ▷ section 4.2.3 |
| `.innercells` | `list(SuperCell)` | ▷ section 4.2.3 |

Finally, the following table gives a description of the 17 importable methods of the class. In the second column, we use the symbol $\sim$ to refer to a type that is specified in the topmost section shown in the corresponding rightmost column.

| Methods | | | |
|---|---|---|---|
| Name | Input types | Output types | Related sections |
| .__init__ | - Cell<br>- list(SuperCell) | - self | ▷ section 4.2.3 |
| .set_level | - int | - self | ▷ section 4.2.4 |
| .reset_depth | - self | - self | ▷ section 4.2.5 |
| .stdout | - list(float) | - self | ▷ section 4.2.6 |
| .compress | - self | - Cell | ▷ section 4.2.7 |
| .action | - list(float)<br>- fun: ∼ -> ∼<br>- string | - list(float) | ▷ section 4.2.8 |
| .base | - fun: ∼ -> ∼ | - SuperCell | ▷ section 4.2.9 |
| .compute_variables | - list(list(float)) | - list(list(float)) | ▷ section 4.2.10 |
| .homeostasis | - list(list(float)) | - self | ▷ section 4.2.11 |
| .allostasis | - list(float)<br>- fun: ∼ -> ∼<br>- fun: ∼ -> ∼ | - self | ▷ section 4.2.12<br>▷ section 4.2.11 |
| .spontaneous_reaction | - list(float)<br>- fun: ∼ -> ∼ | - self | ▷ section 4.2.13 |
| .tensor_pre_action | - self | - self | ▷ section 4.2.14 |
| .merge_base | - list(list(float))<br>- fun: ∼ -> ∼<br>- string | - self | ▷ section 4.2.15<br>▷ section 4.2.17 |
| .divide_base | - list(list(float))<br>- fun: ∼ -> ∼<br>- string | - self | ▷ section 4.2.16<br>▷ section 4.2.18 |
| .fusion | - list(float)<br>- Operad | - self | ▷ section 4.2.17 |
| .fission | - list(float)<br>- Operad<br>- float | - self | ▷ section 4.2.18<br>▷ section 4.2.19 |
| .compose | - fun: ∼ -> ∼ | - self | ▷ section 4.2.19 |

**4.2.3. Description of .__init__ (method).** This section describes the code and the functionalities of the method .__init__. The method is equipped with the following two input variables, including one that is optional.

| .__init__ | | |
|---|---|---|
| Inputs | Types | Specifications |
| cell | Cell | necessary |
| *innercells | list(SuperCell) | optional |

The method possesses one action, which we describe below through sevaral examples.

| Action | |
|---|---|
| Condition | Always |
| Description | The idea behind the method is to construct a tree structure encoding a super cell in the sense of [1, Conv. 2.2 & Def. 2.3]. The cell encoding the root of the tree is stored in `self.cell` while the list containing the children of a parent cell are stored in the object `self.innercells`.<br>The method goes as follows. First, the method initializes the object `self.level` to zero; the object `self.compose_false` to `False`; and the object `self.pre_action` to an empty list. The method also gives the value contained in the variable `cell` to the object `self.cell` and stores the Boolean value of the logical test<br>`(len(innercells) == 0) or (innercells[0] == [])`<br>in `self.is_leaf`. Note that a super cell may or may not have an object `innercells`.<br>If the value of `self.is_leaf` is `True`, then the object `.innercells` is not allocated and the user cannot access the object. On the other hand, if the value of `self.is_leaf` is `False`, then the object exists and contains the item stored in `innercells[0]`, which is supposed to be a list of `SuperCell` items. Lastly, the method calls the function `.set_level(1)` on every `SuperCell` item of `self.innercells`. This has the consequence of incrementing the value of all the objects `.level` by 1 throughout the hierarchical structure of each `SuperCell` item. The method also gives to the object `self.depth` the value `max_depth+1`, where the term `max_depth` represents the maximum of the values stored in the objects `.depth` throughout the hierarchical structures of all the `SuperCell` items of `self.innercells` |

In the examples given below, we define various `SuperCell` items through the constructor and we check that the arguments passed to the constructor are stored in the appropriate variables. First, recall that the value of the object `.is_leaf` is `True` whenever a single input is given to the constructor or whenever the second input is an empty list.

```
>>> c1 = Cell(2,0.5,[0,0],[[1,2],[5,1],[1,8]])
>>> c2 = Cell(2,0.5,[0,0],[[2,0],[0,7],[4,2]])
>>> c = Cell(2,0.5,[0,0],[[8,5],[4,5]])
>>> sc1 = SuperCell(c1)
>>> print sc1.depth, sc1.level, sc1.is_leaf, sc1.compose_state
0 0 True False
>>> print sc1.pre_action
[]
>>> sc1.cell.stdout()
residual:  0.5
cytosol[1]:  0
cytosol[2]:  0
1-th organelle:  [1, 2]
2-th organelle:  [5, 1]
3-th organelle:  [1, 8]
>>> sc2 = SuperCell(c2,[])
>>> print sc2.depth, sc2.level, sc2.is_leaf, sc2.compose_state
0 0 True False
>>> print sc2.pre_action
[]
```

```
>>> sc2.cell.stdout()
residual:  0.5
cytosol[1]:  0
cytosol[2]:  0
1-th organelle:  [2, 0]
2-th organelle:  [0, 7]
3-th organelle:  [4, 2]
```

While the object `.pre_action` is initialized as an empty list, it can be filled by several methods of the class `SuperCell` to compute the expected signal and specialized signal of the super cell (see [1, Def. 2.12]). As suggested by its name, the object `.pre_action` refers to the *pre-action* of the underlying super cell, as defined in [1, Def. 2.9]. As a result, passing the value contained in the object `.pre_action` to the method `.cell.action` outputs the action of the super cell. Following [1, Def. 2.12], we also use the objects `.pre_action` to store the *expected signals* and *specialized signals* associated with the super cell – these signals are, by definition, pre-actions for particular shapes of inputs. The type of signals stored in each object `.pre_action` depends on the method previously called. For instance, the methods `.action` (see secton 4.2.8) and `.allostasis` (see section 4.2.12) can be used to fill the objects `.pre_action` with their corresponding expected signals. On the other hand, the methods `.action` (see secton 4.2.8) and `.spontaneous_reaction` (see section 4.2.13) can be used to fill the objects `.pre_action` with their corresponding specialized signals.

When the methods of the class `SuperCell` are used in accordance with the instructions of this documentation, the organelles of the `Cell` item contained in the object `.cell` should be equal to the contents of the directly-accessible child `Cell` items stored in `.innercells` (for the corresponding indices). Note that some `SuperCell` item do not possess the object `.innercells`, namely those `SuperCell` items whose object `.is_leaf` contains the value `True` – we refer to these cells as *leaves* [1, Conv. 2.2 & Def. 2.3]. To equip a `SuperCell` item with an object `.innercells`, it suffices to pass a list of `SuperCell` item to the second argument of the constructor, as shown below.

```
>>> sc = SuperCell(c,[sc1,sc2])
>>> print sc.depth, sc.level, sc.is_leaf, sc.compose_state
1 0 False False
>>> print sc.pre_action
[] 17 1000000000000000
>>> sc.cell.stdout()
residual:  0.5
cytosol[1]:  0
cytosol[2]:  0
1-th organelle:  [8, 5]
2-th organelle:  [4, 5]
```

**4.2.4. Description of `.set_level` (method).** This section describes the code and the functionalities of the method `.set_level`. The method is equipped with the following input variable.

| .set_level | | |
|---|---|---|
| Inputs | Types | Specifications |
| `level` | `list(SuperCell)` | necessary |

The method possesses one action, which we describe below through an example.

| Action | |
|---|---|
| Condition | Always |
| Description | The method passes the value contained in the input `level` to the object `self.level` and uses the function `.set_level(level+1)` on every `SuperCell` item contained in the list `self.innercells`. For a well-defined super cell whose root cell is at level `0`, this is equivalent to shifting the values of each object `.level` present in the hierarchical structure of the super cell by the value `level-1` |

The following example shows how to use the method `.set_level`.

```
>>> c1 = Cell(2,0.5,[0,0],[[1,2],[5,1],[1,8]])
>>> c2 = Cell(2,0.5,[0,0],[[2,0],[0,7],[4,2]])
>>> c = Cell(2,0.5,[0,0],[[8,5],[4,5]])
>>> sc1 = SuperCell(c1)
>>> sc2 = SuperCell(c2,[])
>>> sc = SuperCell(c,[sc1,sc2])
>>> print sc.level, sc1.level, sc2.level
0 1 1
>>> sc.set_level(3)
>>> print sc.level, sc1.level, sc2.level
3 4 4
```

**4.2.5. Description of `.reset_depth` (method).** This section describes the code and the functionalities of the method `.reset_depth`, which does not take any input. The method possesses one action, which we describe below through an example.

| Action | |
|---|---|
| Condition | Always |
| Description | The method resets the values of the objects `.depth` for all the `SuperCell` items contained in the hierarchical structure of `self`. The reset is done recursively as follows:<br>(1) if `self` is a leaf, then the value `0` is stored in `self.depth`<br>(2) if `self` is a not leaf, then the method calls the function `.reset_depth()` for every `SuperCell` items in the list `self.innercells` and the object `self.depth` is given the value `max_depth+1` where the term `max_depth` is the maximum of the values of the objects `.depth` present in the hierarchical structures of the `SuperCell` items contained in the list `self.innercells` |

The following example shows how to use the method `.reset_depth`.

```
>>> c1 = Cell(2,0.5,[0,0],[[1,2],[5,1],[1,8]])
>>> c2 = Cell(2,0.5,[0,0],[[2,0],[0,7],[4,2]])
>>> c = Cell(2,0.5,[0,0],[[8,5],[4,5]])
>>> sc1 = SuperCell(c1)
>>> sc2 = SuperCell(c2,[])
>>> sc = SuperCell(c,[sc1,sc2])
>>> sc.depth = 4
>>> sc1.depth = 4
>>> print sc.depth, print sc1.depth, print sc2.depth
4 4 0
>>> sc.reset_depth()
>>> print sc.depth, print sc1.depth, print sc2.depth
1 0 0
```

**4.2.6. Description of .stdout (method).** This section describes the code and the functionalities of the method .stdout. The method is equipped with the following input variable, which is optional.

| .stdout | | |
|---|---|---|
| Inputs | Types | Specifications |
| *vector | list(SuperCell) | optional |

The method possesses two actions, which we describe below through sevaral examples.

| Action 1 | |
|---|---|
| Condition | If *vector is empty |
| Description | The method displays the tree structure of self on the standard output. Every line of the display looks as follows<br>....[lvl] -> Cell[res]{ooo...o}<br>and represents a Cell item belonging to the tree structure of self. The integer lvl shows the level of the Cell item while the integer res is the integer truncation of its residual value (section 4.1.3). Each symbol o represents an organelle of the underlying Cell item. Conventionally, the lines associated with the children of a parent cell are always displayed below the line associated with the parent cell.<br>Finally, the method colors in purple the cells for which the associated object .compose_state is True in purple (for examples, see section 4.2.15 and section 4.2.16). |

The following example shows how to use the method .stdout with no input.

```
>>> c1 = Cell(2,0.5,[0,0],[[1,2],[5,1],[1,8]])
>>> c2 = Cell(2,0.5,[0,0],[[2,0],[0,7],[4,2]])
>>> c = Cell(2,0.5,[0,0],[[8,5],[4,5]])
>>> sc1 = SuperCell(c1)
>>> sc2 = SuperCell(c2,[])
>>> sc = SuperCell(c,[sc1,sc2])
>>> sc.stdout()
[0] -> Cell[0]{oo}
.[1] -> Cell[0]{ooo}
.[1] -> Cell[0]{ooo}
```

We now describe the second action.

| Action 2 | |
|---|---|
| Condition | If `*vector` is a list of length equal to `self.dimension` |
| Description | The method displays the tree structure of `self` on the standard output. Every line of the display can take two forms: either the form seen in Action 1, or the following form<br>`....[lvl] -> Cell[res]{ooo...o} [num]: agr ... [num]: agr`<br>which represents a `Cell` item for which the object `.is_leaf` contains the value `True`. As with Action 1, the integers `lvl` and `res` correspond to the level of the `Cell` item and the integer truncation of its residual value, respectively. The list of terms `[num]: agr` shows the list of outputs `agr` returned by the functions `self.cell.agreement(num,vector[0])` (see section 4.1.16) where `num` is the index of the organelle of the underlying `Cell` item. Conventionally, the lines representing the children of a parent cell are always displayed below the line associated with the parent cell.<br>Finally, the method colors in purple the cells for which the associated object `.compose_state` is `True` in purple (for examples, see section 4.2.15 and section 4.2.16). |

The following example shows how to use the method `.stdout` with an input. We will consider the same setting as in the previous example.

```
>>> sc.stdout([1,10])
[0] -> Cell[0]{oo}
.[1] -> Cell[0]{ooo} [0]:0.9344877349 [1]:0.292714272 [2]:0.9996953077
.[1] -> Cell[0]{ooo} [0]:0.099503719 [1]:0.9950371902 [2]:0.5339929913
```

**4.2.7. Description of `.compress` (method).** This section describes the code and the functionalities of the method `.compress`, which does not take any input. The method possesses one action, which we describe below through an example.

| Action | |
|---|---|
| Condition | Always |
| Description | The method returns the `Cell` item resulting from composing (see section 4.1.8) all the cells present in the hierarchical structure of `self` (recursively from the leaves to the root). |

The following example illustrates how to use the method `.compress`. In particular, we use the methods `.stdout` associated with the class `Cell` (see section 4.1.6) and the class `SuperCell` (see section 4.2.6).

```
>>> c1 = Cell(2,0.5,[0,0],[[1,2],[5,1],[1,8]])
>>> c2 = Cell(2,0.5,[0,0],[[2,0],[0,7],[4,2]])
>>> c = Cell(2,0.5,[0,0],[[8,5],[4,5]])
>>> sc1 = SuperCell(c1)
>>> sc2 = SuperCell(c2,[])
>>> sc = SuperCell(c,[sc1,sc2])
>>> sc.stdout([1,10])
[0] -> Cell[0]{oo}
.[1] -> Cell[0]{ooo} [0]:0.9344877349 [1]:0.292714272 [2]:0.9996953077
.[1] -> Cell[0]{ooo} [0]:0.099503719 [1]:0.9950371902 [2]:0.5339929913
```

```
>>> c = sc.compress()
>>> c.stdout()
residual:  1.5
cytosol[1]:  0
cytosol[2]:  0
1-th organelle:  [1, 2]
2-th organelle:  [5, 1]
3-th organelle:  [1, 8]
4-th organelle:  [2, 0]
5-th organelle:  [0, 7]
6-th organelle:  [4, 2]
>>> d = SuperCell(c)
>>> d.stdout([1,10])
[0] -> Cell[1]{oooooo} [0]:0.9344877349 [1]:0.292714272 [2]:0.9996953077
[3]:0.099503719 [4]:0.9950371902 [5]:0.5339929913
```

**4.2.8. Description of `.action` (method).** This section describes the code and the functionalities of the method `.action`. The method is equipped with the following three input variables, including one that is optional.

| .action | | |
|---|---|---|
| Inputs | Types | Specifications |
| `vector` | `list(SuperCell)` | necessary |
| `identity` | `fun: list(float) -> Cell` | necessary |
| `option = "expected"` | `string` | optional |

The method possesses two actions, which we describe below through sevaral examples.

| Action 1 | |
|---|---|
| Condition | The input variable `option` contains the string `"specialized"` |
| Description | The method returns the action associated with the "specialized signal" [1, Def. 2.12] of the super cell `self` on the input `vector` relative to the function `identity`. To do so, the method first stores the recursive collection of specialized signals in the corresponding objects `.pre_action`. Each list `self.pre_action` is constructed through the following recursive process: (where every index `i` ranges from `0` to `len(self.cell.organelles)-1`):<br>(1) If `self.is_leaf` is `True`, then the element `self.pre_action[i]` is given the action of the cell `identity(self.cell.organelles[i])` on `vector` (see section 4.1.13);<br>(2) If `self.is_leaf` is `False`, then the element `self.pre_action[i]` is given the action of the super cell `self.innercells[i]` on `vector` relative to the function `identity`.<br>Then, the method `self.action` returns a `list` item that corresponds to the output of the function `self.cell.action` on the list `self.pre_action`. |

In the following two examples, we use the method `.stdout` (see section 4.2.6) with the constructor of the class `Operad` (see section 4.3.3), for a dimension equal to 2, and its associated method `.identity` of `Operad` (see section 4.3.4).

```
>>> c1 = Cell(2,0.5,[0,0],[[1,2],[5,1],[1,8]])
>>> c2 = Cell(2,0.5,[0,0],[[2,0],[0,7],[4,2]])
>>> c = Cell(2,0.5,[0,0],[[8,5],[4,5]])
>>> sc1 = SuperCell(c1)
>>> sc2 = SuperCell(c2,[])
>>> sc = SuperCell(c,[sc1,sc2])
>>> sc.stdout([1,10])
[0] -> Cell[0]{oo}
.[1] -> Cell[0]{ooo} [0]:0.9344877349 [1]:0.292714272 [2]:0.9996953077
.[1] -> Cell[0]{ooo} [0]:0.099503719 [1]:0.9950371902 [2]:0.5339929913
>>> operad = Operad(2)
>>> print sc.action([1,10],operad.identity,"specialized")
[0.14971941638608305, 2.248877665544332]
```

In the following example, we see that the method `.compress` (see section 4.2.7) can change the output of the method `.action`, as shown below.

```
>>> c = sc.compress()
>>> d = SuperCell(c)
>>> d.stdout([1,10])
[0] -> Cell[1]{oooooo} [0]:0.9344877349 [1]:0.292714272 [2]:0.9996953077
[3]:0.099503719 [4]:0.9950371902 [5]:0.5339929913
>>> print d.action([1,10],operad.identity,"specialized")
[0.28114478114478114, 4.9326599326599325]
```

We now describe the second action of the method.

| Action 2 | |
|---|---|
| Condition | The input variable `option` is not specified and/or contains a string that is not `"specialized"` |
| Description | The method returns the action associated with the "expected signal" [1, Def. 2.12] of super cell `self` on the input `vector` relative to the function `identity`. To do so, the method first stores the recursive collection of expected signals associated with the cells of the super cell in the corresponding objects `.pre_action`. Each list `self.pre_action` is constructed through the following recursive process: (where every index `i` ranges from `0` to `len(self.cell.organelles)-1`):<br>(1) If `self.is_leaf` is `True`, then the element `self.pre_action[i]` stores the list `vector` (see section 4.1.13);<br>(2) If `self.is_leaf` is `False`, then the element `self.pre_action[i]` is given the action of the super cell `self.innercells[i]` on `vector` relative to the function `identity`.<br>Then, the method `self.action` returns a list item that corresponds to the output of the function `self.cell.action` on the list `self.pre_action`. |

The following example uses the same items as those used to illustrate Action 1. As in the previous example, we compare the action of a super cell and its 'compressed' version.

```
>>> print sc.action([1,10],operad.identity)
[0.2141414141414142, 2.7525252525252526]
>>> print d.action([1,10],operad.identity)
[0.3939393939393939, 6.060606060606061]
```

**4.2.9. Description of `.base` (method).** This section describes the code and the functionalities of the method `.base`. The method is equipped with the following input variable.

| .base | | |
|---|---|---|
| Inputs | Types | Specifications |
| identity | fun: list(float) -> Cell | necessary |

The method possesses one action, which we describe below through sevaral examples.

| Action | |
|---|---|
| Condition | Always |
| Description | The method returns the basal pruning of the tree structure of self (see [1, Def. 2.10]), namely the super cell that only consists of the root cell of self and its associated children (formal or not). This means that the output is either the truncation of the tree structure of self at level 1 or the formal extension of self into a SuperCell item of height 1 if self is a leaf.<br>More specifically, the method outputs a SuperCell item whose object .cell is equal to the Cell item in self.cell and whose object .innercells is of the following form:<br>(1) If self.is_leaf is True, then .innercells[i] is the leaf super cell SuperCell(identity(self.cell.organelles[i])) for every index i ranging from 0 to len(self.innercells)-1.<br>(2) If self.is_leaf is False, then .innercells[i] is the leaf super cell SuperCell(self.innercells[i].cell) for every index i ranging from 0 to len(self.innercells)-1. |

The following example illustrates how to use the method .base. We make use of the method .stdout (see section 4.2.6), which we use to compare the input with the output of the method .base. Note that our examples usess the constructor of the class Operad (see section 4.3.3), for a dimension equal to 2, and its associated method .identity of Operad (see section 4.3.4). First, we give an example where the method .base returns the formal extension of a leaf into a SuperCell of height 1 by replacing its organelles with identity cells.

```
>>> c1 = Cell(2,0.5,[0,0],[[1,2],[5,1],[1,8]])
>>> c2 = Cell(2,0.5,[0,0],[[2,0],[0,7],[4,2]])
>>> c = Cell(2,0.5,[0,0],[[8,5],[4,5]])
>>> sc1 = SuperCell(c1)
>>> operad = Operad(2)
>>> f = sc1.base(operad.identity)
>>> f.stdout()
[0] -> Cell[0]{ooo}
.[1] -> Cell[0]{o}
.[1] -> Cell[0]{o}
.[1] -> Cell[0]{o}
```

We now illustrate the case where the method .base truncates the tree structure of a super cell at level 1. For simplicity, we use the method .copy, described in section 4.1.5, to build a 2-level super cell sd as follows.

```
>>> sc2 = SuperCell(c2,[])
>>> sc = SuperCell(c,[sc1,sc2])
>>> d = c.copy()
>>> c3= c2.copy()
>>> sc3 = SuperCell(c3)
>>> sd = SuperCell(d,[sc,sc3])
```

```
>>> sd.stdout()
[0] -> Cell[0]{oo}
.[1] -> Cell[0]{oo}
..[2] -> Cell[0]{ooo}
..[2] -> Cell[0]{ooo}
.[1] -> Cell[0]{ooo}
```

The truncation of `sd` by the method `.base` is shown below through the method `.stdout` (section 4.2.6).

```
>>> se = sd.base(operad.identity)
>>> se.stdout()
[0] -> Cell[0]{oo}
.[1] -> Cell[0]{oo}
.[1] -> Cell[0]{ooo}
```

**4.2.10. Description of `.compute_variables` (method).** This section describes the code and the functionalities of the method `.compute_variables`. The method is equipped with the following input variable.

| `.compute_variables` | | |
|---|---|---|
| Inputs | Types | Specifications |
| `gradient_descent` | `list(list(float))` | necessary |

The method possesses one action, which we describe below through an example.

| Action | |
|---|---|
| Condition | Always |
| Description | The method returns four lists `mu_var`, `alpha_var`, `kappa_var`, and `lambda_var` corresponding to the tuples $\mu$, $\alpha$, $\kappa$ and $\lambda$ needed to defined the cell $\mathsf{left}(c)(\alpha,\kappa,\lambda)$ and the collection $\mathsf{right}(c)(\mu,\alpha,\kappa)$ (see [1, Conv. 1.19], section 4.1.10 and section 4.1.11). In particular, these four lists correspond to the type of lists used to define and solve the homeostasis problem induced by $\lambda$ (see [1, Def. 1.39] and/or the description below)<br>First, the method stores in `mu_var` the list of values `self.innercells[i].cell.residual` for every index `i` ranging from `0` to `len(self.cell.organelles)` and constructs the list `alpha_var` as the list of integers `len(self.innercells[i].cell.organelles)` for every index `i` ranging from `0` to `len(self.cell.organelles)`.<br>Before explaining the the method constructs the two lists `lambda_var` and `kappa_var`, it is worth mentioning that the latter is the solution of the homeostasis problem induced by the former (see [1, Def. 1.39]). In other words, the list `lambda_var` is to define the organelles of a re-factorization of the basal junction of `self` (see 4.2.9) while the list `kappa_var` contains the new cytosols for the child cells of the resulting junction. |

| | |
|---|---|
| Description | In fact, the coefficient `lambda_var[i][u]` is expected to be relatively close to the value `self.cell.organelles[i][u]`. The amount by which `self.cell.organelles[i][u]` differs from corresponds to the value contained in `gradient_descent[i][u]`, which is expected to be small compared to `self.cell.organelles[i][u]` (as with any gradient-descent-based method). If the value contained in `gradient_descent[i][u]` is negative (implying that the cell $\mathsf{left}(c)(\alpha,\kappa,\lambda)$ is not well-defined: meaning that the associated object `.well_defined` is `False`), then the method `.compute_variables` changes the value of `gradient_descent[i][u]` to `0`. Then, the method computes `lambda_var` as the list containing the difference<br>`self.cell.organelles[i][u] - gradient_descent[i][u]`<br>for every pair `(i,u)` of valid indices. The modification of `gradient_descent` is important for the method `.homeostasis` (see section 4.2.11) in order to create cells $\mathsf{left}(c)(\alpha,\kappa,\lambda)$ and $\mathsf{right}(c)(\mu,\alpha,\kappa)$ that are well-defined .<br>Regarding `kappa_var`, each coefficient `kappa_var[i]` correspond to a list computing according to the vector formula of [1, Def. 1.41]. Specifically, this means that each coefficient `kappa_var[i][u]` contains the value<br>`lambda_var[i][u]`<br>`- self.innercells[i].cell.K[u]`<br>`+ self.innercells[i].cell.cytosol[u]`<br>for every index `u` ranging from `0` to `self`.dimension-`1`. |

The following example illustrates how to use the method `.compute_variables`. In particular, we make use of the method `.base` (see section 4.2.9), which allows us to interpret a leaf `SuperCell` item (see section 4.2.3) as a tree of cells of height 1, and we the method `.identity` associated with the class `Operad`.

```
>>> c = Cell(3,0.5,[12,5,8],[[7,2,1],[10,5,1],[1,7,8],[1,2,5]])
>>> sc = SuperCell(c)
>>> operad = Operad(3)
>>> base = sc.base(operad.identity)
>>> gradient_descent = [[.1,3,.1],[.1,.2,.1],[.1,.05,.2],[-2,.1,.1]]
>>> mu_var, alpha_var, kappa_var, lambda_var = base.compute_variables(
gradient_descent)
>>> print mu_var
[1, 1, 1, 1]
>>> print alpha_var
[0, 0, 0, 0]
>>> print kappa_var
[[-0.09999999999999964, 0, -0.09999999999999998], [-0.09999999999999964,
-0.20000000000000018, -0.09999999999999998], [-0.09999999999999998,
-0.04999999999999982, -0.20000000000000018], [2, -0.10000000000000009,
-0.09999999999999964]]
>>> print lambda_var
[[6.9, 2, 0.9], [9.9, 4.8, 0.9], [0.9, 6.95, 7.8], [3, 1.9, 4.9]]
```

**4.2.11. Description of `.homeostasis` (method).** This section describes the code and the functionalities of the method `.homeostasis`. The method is equipped with the following input variables.

| .homeostasis | | |
|---|---|---|
| Inputs | Types | Specifications |
| gradient_descent | list(list(float)) | necessary |
| identity | fun: list(float) -> Cell | necessary |

The method possesses one action, which we describe below through an example.

| Action | |
|---|---|
| Condition | Always |
| Description | The method finds a re-factorization, in the sense of [1, Def. 1.39], for the basal junction [1, Def. 2.10] of the tree structure of `self` [1, Conv. 2.2]. To do so, the method loops on the outputs of the method `.homeostasis` until its outputs generate a well-defined cell factorization.<br>More specifically, the method stores the basal junction of `self` in a variable `cd` by using the function `self.base(identity)` (see section 4.2.9). Then, the method uses the function<br>`cd.compute_variables(gradient_descent)`<br>to compute candidate parameters $\mu$, $\alpha$, $\kappa$ and $\lambda$ that allow us to re-factorize the junction `cd` into a junction consisting of the root cell $\mathsf{left}(c,d)(\alpha,\kappa,\lambda)$ and the list of child cells $\mathsf{right}(c,d)(\mu,\alpha,\kappa)$, as defined in [1, Conv. 1.19]). Recall that the data $\mathsf{left}(c,d)(\alpha,\kappa,\lambda)$ and $\mathsf{right}(c,d)(\mu,\alpha,\kappa)$ can be obtained through the methods `left` (section 4.1.10) and `right` (section 4.1.11). The other role of `cd.compute_variables` is to find the invalid perturbations values of `gradient_descent` and to set them to 0.<br>The method `.homeostasis` then re-computes (if necessary) the solutions $\mathsf{left}(c,d)(\alpha,\kappa,\lambda)$ and $\mathsf{right}(c,d)(\mu,\alpha,\kappa)$ until they all define well-defined `Cell` items.<br>When the cells present in the tree structure of `self` (and in particular the cells making the junction `cd`) are well-defined, the method is guaranteed to terminate (Indeed, if `gradient_descent` is only made of zeros, then the junction `cd` is a valid output for `.homeosasis`). |

The following example illustrates how to use the method `.homeostasis`. We make use of the class `Operad` (see section 4.3.3) and its associated method `.identity` (see section 4.3.4).

```
>>> c = Cell(3,0.5,[12,5,8],[[7,2,1],[10,5,1],[1,7,8],[1,2,5]])
>>> sc = SuperCell(c)
>>> operad = Operad(3)
>>> sc.cell.stdout()
residual:  0.5
cytosol[1]:  12
cytosol[2]:  5
cytosol[3]:  8
1-th organelle:  [7, 2, 1]
2-th organelle:  [10, 5, 1]
3-th organelle:  [1, 7, 8]
4-th organelle:  [1, 2, 5]
>>> gradient_descent = [[.1,3,.1],[.1,.2,.1],[.1,.05,.2],[-2,.1,.1]]
>>> sc.homeostasis(gradient_descent,operad.identity)
>>> sc.cell.stdout()
residual:  0.5
```

```
cytosol[1]:  10.3
cytosol[2]:  5.35
cytosol[3]:  8.5
1-th organelle:  [6.9, 2, 0.9]
2-th organelle:  [9.9, 4.8, 0.9]
3-th organelle:  [0.9, 6.95, 7.8]
4-th organelle:  [3, 1.9, 4.9]
```

**4.2.12. Description of `.allostasis` (method).** This section describes the code and the functionalities of the method `.allostasis`. The method is equipped with the following input variables.

| `.allostasis` | | |
|---|---|---|
| Inputs | Types | Specifications |
| `vector` | `list(float)` | necessary |
| `identity` | `fun: list(float) -> Cell` | necessary |
| `gamma` | `fun: Cell * list(list(float)) -> list(list(float))` | necessary |

The method possesses one action, which we describe below through an example.

| Action | |
|---|---|
| Condition | Always |
| Description | The method computes the algorithmic step described in [1, Sec. 2.3]. Recall that this step consists in mimicking a gradient-descent-like optimization of the parameters of the super cell `self` in order to minimize the values of the algebra operators (section 4.1.14) associated with each cells present in the hierarchical structure of `self`.<br>The method first stores the "expected signals" [1, Def. 2.12] of `self` relative to the list `vector` in the corresponding objects `.pre_action` (also, see section 4.2.8).<br>Then, for each cell in the hierarchical structure of `self` accessible via each object `.cell` (section 4.2.3), the method computes the allostatic differential [1, Def. 1.47] through the method `.allostasis` (section 4.1.15), namely the value<br>`dU_de = self.cell.algebra_operator(self.pre_action).`<br>The method then uses the allostatic differential `dU_de` and the function `gamma` to compute the allostatic matrix [1, Def. 2.18] associated with `self.cell`: this is done by creating a `list(list(float))` item `gradient_descent` whose coefficient `gradient_descent[i][u]` is the multiplication<br>`gamma(self.cell,i,self.pre_action)[i][u] * dU_de`<br>for every index `i` ranging from `0` to `len(self.cell.organelles)-1` and every index `u` ranging from `0` to `self.cell.dimension-1`. Finally, the matrix `gradient_descent` is given to the method `self.homeostasis` (section 4.2.11) together with the input `identity` in order to make each junction of `self` 'reach homeostasis' (In the sense of [1, Def. 2.11]). |

The following example illustrates how to use the method `.homeostasis`. We make use of the class `Operad` (see section 4.3.3), its associated method `.identity` (see section 4.3.4) and the function `usf.gamma` (see section 3.1.4), which we pass to the input parameter `gamma` .

We start our examples with a super cell that possess the same paramters as those used in the examples of section 3.1.3 and section 3.1.4, except for the amount given to the object

.residual. Note that a higher value for .residual allows the method .allostasis to update more parameters as it is less likely to produce ill-defined re-factorizations of the super cell through the use of the method .homeostasis (see [1, Def. 2.15], section 4.2.10, and section 4.2.11 for more information on well-definedness and the re-factorization process).

```
>>> operad = Operad(5)
>>> c = Cell(dimension = 5, residual = 1000, cytosol = [-2,3,5,0,0],
organelles = [[1,2,5,3,7], [5,0,0,0,4],[25,51,8,1,52]])
>>> sc = SuperCell(c)
>>> sc.cell.stdout()
residual:  1000
cytosol[1]:  -2
cytosol[2]:  3
cytosol[3]:  5
cytosol[4]:  0
cytosol[5]:  0
1-th organelle:  [1, 2, 5, 3, 7]
2-th organelle:  [5, 0, 0, 0, 4]
3-th organelle:  [25, 51, 8, 1, 52]
```

We now use the function usf.gamma (section 3.1.4) to generate a “gamma parameter” for the method .allostasis.

```
>>> brightness = [.25,.5,.75]
>>> profiles = [[(.0,.9),(.0,.5),(.0,.25)],[(.0,.4),(.0,.6),(.0,.5)]]
>>> scores = [(.5,1),(.8,1)]
>>> gamma = usf.gamma(3,20,brightness,profiles,scores)
>>> sc.allostasis([10,1,2,5,15],operad.identity,gamma)
[brightness] 0.8 0.4 0.2
True False

[brightness] 0.4 0.4 0.4
False True

[brightness] 0.6 0.4 0.4
False False

[Gamma parameters][0]
79.6021925779
1000.0
23.2126583827
[Allostasis matrix][1]
6.26672163138
323.214148404
2.41792613144
[Allostasis matrix][2]
-8.36162731541
-36.2360630527
-0.427461634702
```

The displays appearing after the green labels tell us about the average values of the gamma parameters and the average values of the coefficients of the allostatic matrix. The values displayed after the label [Allostasis matrix][2] are the average values of the values selected by the method .homeostasis for a valid update of the parameters of the super cell.

In our case, calling the method `.allostasis` has updated the super cell as follows.

```
>>> sc.cell.stdout()
residual:  1000
cytosol[1]:  -11.6441450451
cytosol[2]:  11.8732941391
cytosol[3]:  5
cytosol[4]:  0
cytosol[5]:  -224.354909108
1-th organelle:  [8.92395297628549, 2, 5, 3, 40.884183600753644]
2-th organelle:  [5, 0, 0, 0, 185.18031526336028]
3-th organelle:  [26.720192068820307, 42.12670586090116, 8, 1,
61.29041024378717]
```

After the update, we can see that the organelles of the super cell are more specialized in certain dimensions, which can make the next update more difficult for random inputs. We illustrate this in the next example, in which the organelles of the super cell all satisfy one of the contrast profiles associated with the function `usf.gamma`, but do not pass the agreement test (see section 3.1.3 and section 3.1.4 for more detail on contrast profiles and agreement tests).

```
>>> sc.allostasis([3,12,20,15,1],operad.identity,gamma)
[brightness] 0.2 0.2 0.2
True True

[brightness] 0.2 0.2 0.2
True True

[brightness] 0.6 0.4 0.2
True False

[Gamma parameters][0]
0.0
0.0
0.0
[Allostasis matrix][1]
0
0
0
[Allostasis matrix][2]
0
0
0
```

As is shown below, the parameters of the super cell have not changed.

```
>>> sc.cell.stdout()
residual:  1000
cytosol[1]:  -11.6441450451
cytosol[2]:  11.8732941391
cytosol[3]:  5
cytosol[4]:  0
cytosol[5]:  -224.354909108
1-th organelle:  [8.92395297628549, 2, 5, 3, 40.884183600753644]
2-th organelle:  [5, 0, 0, 0, 185.18031526336028]
```

```
3-th organelle:  [26.720192068820307, 42.12670586090116, 8, 1,
61.29041024378717]
```

Now, we give another example showing what happens when the super cell does not have enough “energy” in its object `.residual`. We use the same parameters as those used in the first example given above, except for the value give to `.residual`, which is set to 200.

```
>>> c = Cell(dimension = 5, residual = 200, cytosol = [-2,3,5,0,0],
organelles = [[1,2,5,3,7], [5,0,0,0,4],[25,51,8,1,52]])
>>> sc = SuperCell(c)
>>> sc.cell.stdout()
residual:  200
cytosol[1]:  -2
cytosol[2]:  3
cytosol[3]:  5
cytosol[4]:  0
cytosol[5]:  0
1-th organelle:  [1, 2, 5, 3, 7]
2-th organelle:  [5, 0, 0, 0, 4]
3-th organelle:  [25, 51, 8, 1, 52]
```

We now call the method `.allostasis` again. Because we have not changed the organelles of the super cell, we obtain the same displays regarding the brightness profiles satisfied by these organelles.

```
>>> gamma = usf.gamma(3,20,brightness,profiles,scores)
>>> sc.allostasis([10,1,2,5,15],operad.identity,gamma)
[brightness] 0.8 0.4 0.2
True False

[brightness] 0.4 0.4 0.4
False True

[brightness] 0.6 0.4 0.4
False False

[Gamma parameters][0]
79.6021925779
1000.0
23.2126583827
```

As can be seen, the result of applying the method `.allostasis` for such the new value of `.residual` has the consequence of canceling the expected updates for the first and second organelles (see the average updates after the label `[Allostasis matrix][2]` below)

```
[Allostasis matrix][1]
6.26672163138
323.214148404
2.41792613144
[Allostasis matrix][2]
0
0
1.77465882782
```

As shown below, the parameters of the super cell have been changed accordingly.

```
>>> sc.cell.stdout()
residual:  200
cytosol[1]:  -2
cytosol[2]:  11.8732941391
cytosol[3]:  5
cytosol[4]:  0
cytosol[5]:  0
1-th organelle:  [1, 2, 5, 3, 7]
2-th organelle:  [5, 0, 0, 0, 4]
3-th organelle:  [25, 42.12670586090116, 8, 1, 52]
```

**4.2.13. Description of `.spontaneous_reaction` (method).** This section describes the code and the functionalities of the method `.spontaneous_reaction`. The method is equipped with the following input variables.

| `.spontaneous_reaction` | | |
|---|---|---|
| Inputs | Types | Specifications |
| `vector` | `list(float)` | necessary |
| `identity` | `fun: list(float) -> Cell` | necessary |

The method possesses one action, which we describe below through an example.

| Action | |
|---|---|
| Condition | Always |
| Description | The present method extends the method `.spontaneous_reaction` defined for `Cell` items (section 4.1.12) to `SuperCell` items. In addition, the method carries out two other tasks: it stores the specialized signals [1, Def. 2.12] of `self` generated by the input `vector` in the corresponding objects `pre_action` (also, see section 4.2.8) and it implements the 'cleaning' operation described in [1, Def. 2.16], which means that it makes the `SuperCell` item `self` reach homeostasis (see [1, Def. 2.11 & Rem. 2.17]). Here, 'reaching homeostatis' means that the content of any `Cell` item of the form `*.innercells[i].cell`, where `*` represents a `SuperCell` item in the hierarchical structure of `self`, equals the list contained in the object `*.cell.organelles[i]` (see [1, Def. 2.16: item 2.3]).<br>To do so, the method calls the function `*.cell.spontaneous_reaction()` in every super cell `*` present in the hierarchical structure of `self`, and then stores the specialized signals generated by the input `vector` in the corresponding objects `*.pre_action`.<br>After each computation of a specialized signal, the method uses the parameters of the updated child super cells (stored in `.innercells`) to compute the value of the objects `.cell.organelles`, `*.cell.Sorg`, `*.cell.Sorg` and `*.cell.SK` (section 4.1.3). In addition, the method uses the assignments<br>`*.cell.organelles[i] = *.innercells[i].cell.K`<br>throughout the hierarchical structure of `self` to put the returned `SuperCell` item `self` in a homeostatic state (see [1, Def. 2.11 & Rem. 2.17]). |

The following example shows the action of the method `.spontaneous_reaction` on a `SuperCell` item. As can be seen, the cytosol of the three cell `c1`, `c2` and `c` are turned into lists of zeros and the residuals are augmented accordingly (see section 4.1.12).

```
>>> c1 = Cell(2,0.5,[4,2],[[1,2],[5,1],[1,8]])
>>> c2 = Cell(2,0.5,[1,2],[[2,0],[0,7],[4,2]])
>>> c = Cell(2,0.5,[8,7],[[8,5],[4,5]])
>>> sc1 = SuperCell(c1)
>>> sc2 = SuperCell(c2,[])
>>> sc = SuperCell(c,[sc1,sc2])
>>> operad = Operad(2)
>>> sc.stdout([4,5])
[0] -> Cell[0]{oo}
.[1] -> Cell[0]{ooo} [0]:0.977802414 [1]:0.7657048647 [2]:0.8523227286
.[1] -> Cell[0]{ooo} [0]:0.6246950475 [1]:0.7808688094 [2]:0.90795938
>>> sc.spontaneous_reaction([4,5],operad.identity)
>>> sc.cell.stdout()
residual:  15.5
cytosol[1]:  0
cytosol[2]:  0
1-th organelle:  [7, 11]
2-th organelle:  [6, 9]
```

We can display the description of the child super cells of cs to see that the cytosolic content have been cleaned up and that the objects .residual have been filled with the sum of their associated cytosolic contents.

```
>>> sc.innercells[0].cell.stdout()
residual:  6.5
cytosol[1]:  0
cytosol[2]:  0
1-th organelle:  [1, 2]
2-th organelle:  [5, 1]
3-th organelle:  [1, 8]
>>> sc.innercells[1].cell.stdout()
residual:  3.5
cytosol[1]:  0
cytosol[2]:  0
1-th organelle:  [2, 0]
2-th organelle:  [0, 7]
3-th organelle:  [4, 2]
```

**4.2.14. Description of .tensor_pre_action (method).** This section describes the code and the functionalities of the method .tensor_pre_action, which does not take any input. The method possesses one action, which we describe below through an example.

| Action | |
|---|---|
| Condition | Always |
| Description | In the same way as [1, Prop. 1.54] computes a barycenter of pre-actions (see [1, Rem. 1.55]), the present method computes the barycenter of the pre-actions associated with the basal junction of self, where we take the barycentric weights to be the outputs of the function self.cell.content_proportions() (see section 4.1.20).<br>Specifically, the output of the method is a list(float) item, call it barycenter, of length self.cell.dimension, whose element barycenter[u] is the sum of the terms<br>self.pre_action[i][u] * prop[i]<br>in which prop[i] is the i-th output of the function self.cell.content_proportions(), namely the ratio<br>self.cell.Sorg[i]/float(self.cell.SK),<br>over every index i ranging from 0 to len(self.cell.organelles)-1. |

The following example shows how to use the method .tensor_pre_action. Note that to use this method, we need to pre-fill the objects .pre_action of the super cell with lists of values. Ideally, these lists would correspond to one of the two types of signals associated with the super cell, namely either its expected signals or its specialized signals (see [1, Def. 2.12]). Below, we use the method spontaneous_reaction to fill the objects .pre_action with specialized signals. We use the same items as those used in section 4.2.13.

```
>>> c1 = Cell(2,0.5,[4,2],[[1,2],[5,1],[1,8]])
>>> c2 = Cell(2,0.5,[1,2],[[2,0],[0,7],[4,2]])
>>> c = Cell(2,0.5,[8,7],[[8,5],[4,5]])
>>> sc1 = SuperCell(c1)
>>> sc2 = SuperCell(c2,[])
>>> sc = SuperCell(c,[sc1,sc2])
>>> operad = Operad(2)
>>> sc.stdout([4,5])
[0] -> Cell[0]{oo}
.[1] -> Cell[0]{ooo} [0]:0.977802414 [1]:0.7657048647 [2]:0.8523227286
.[1] -> Cell[0]{ooo} [0]:0.6246950475 [1]:0.7808688094 [2]:0.90795938
>>> sc.spontaneous_reaction([4,5],operad.identity)
>>> print sc.pre_action
[[1.0246913580246915, 2.391975308641975], [1.2444444444444445,
2.5555555555555556]]
>>> print sc.tensor_pre_action()
[1.1245791245791246, 2.4663299663299663]
```

**4.2.15. Description of .merge_base (method).** This section describes the code and the functionalities of the method .merge_base. The method is equipped with the following three input variables, including one that is optional.

| .merge_base | | |
|---|---|---|
| Inputs | Types | Specifications |
| list_of_organelles | list(list(float)) | necessary |
| tensor | fun: list(SuperCell) -> SuperCell | necessary |
| order = "non-sorted" | string | optional |

The method possesses one action, which we describe below through sevaral examples.

<table>
<tr><th colspan="2">Action</th></tr>
<tr><td>Condition</td><td>Always</td></tr>
<tr><td>Description</td><td>The method .merge_base implements the operation described in [1, Def. 2.26]. The idea is to extend the method .merge (section 4.1.17) associated with the class Cell to the basal junction of self (see section 4.2.9). Put simply, the method merges a subset of the organelles of the root cell of self into a single organelle and uses the input function tensor to distribute the child super cells, stored in self.innercells, accordingly (see picture below). Note that .merge_base does not act on leaf super cells (see section 4.2.3).

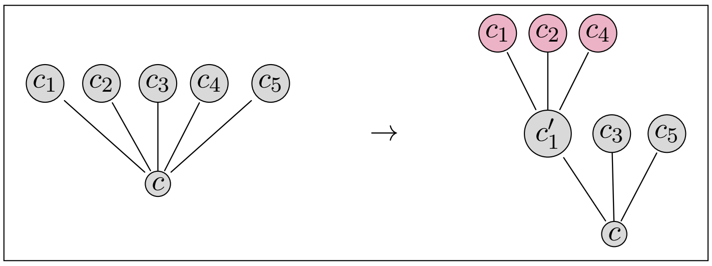

Now, to be more specific, the subset of indices of the organelles of self to be merged is given by the list list_of_organelles. Similarly to the method .merge, the list list_of_organelles is sorted before use if the input variable order does not contain the string "sorted". After this sorting step, the method merges the corresponding organelles of self.cell by calling the function

self.cell.merge(list_of_organelles,"sorted").

The method then makes sure that the merging of these organelles is consistent with the whole tree structure of self. This is done by creating two new lists merged_innercells and new_innercells for the merged and untouched organelles. The method fills the list merge_innercells with all the SuperCell items of the form self.innercells[i] for which the index i belongs to list_of_organelles. If we denote by j the smallest index of list_of_organelles, then the element new_innercells[j] is given the SuperCell item tensor(merged_innercells). For any other index, the list new_innercells contains the super cells of the form self.innercells[i] whose index i, ranging from 0 to len(self.innercells)-1, is not in list_of_organelles.<br>
The method also fills the list new_innercells[j].pre_action in the same way as the list new_innercells was filled, but where we replace the list self.innercells with the list self.pre_action.<br>
Finally, the method .merge_base terminates with the following actions:<br>
(1) it sets the level of new_innercells[j] to self.level+1 by calling the method .set_level(self.level+1) (section 4.2.4).<br>
(2) it assigns the list of super cells new_innercells to self.innercells;<br>
(3) it recomputes self.pre_action as the output of the function new_innercells[j].tensor_pre_action() (see section 4.2.14);</td></tr>
</table>

The following example illustrates the use of the method .merge_base with the methods .pseudo_tensor (section 4.3.7) and .merging_tensor (section 4.3.8) associated with the class Operad. In addition, we use the method .action (section 4.2.8) to pre-fill each object pre_action with a non-empty list of length 2, for every super cell present in the hierarchical structure of the super cell sc.

```
>>> c = Cell(2,0.5,[4,2],[[1,2],[5,1],[1,8],[4,4],[0,7]])
>>> operad = Operad(2)
>>> children = list()
>>> for i in range(5):
...     children.append(SuperCell(operad.identity(c.organelles[i])))
>>> sc = SuperCell(c,children)
>>> sc.action([0,0],"specialized")
>>> sc.stdout()
[0] -> Cell[0]{ooooo}
.[1] -> Cell[0]{o}
.[1] -> Cell[0]{o}
.[1] -> Cell[0]{o}
.[1] -> Cell[0]{o}
.[1] -> Cell[0]{o}
```

Below, we use `.merging_tensor`, which sets the objects `.compose_state` of each merged super cell to `True`, as shown by the purple display of the method `.stdout`.

```
>>> sc.merge_base([0,1,3],operad.merging_tensor,"sorted")
>>> sc.stdout()
[0] -> Cell[0]{ooo}
.[1] -> Cell[0]{ooo}
..[2] -> Cell[0]{o}
..[2] -> Cell[0]{o}
..[2] -> Cell[0]{o}
.[1] -> Cell[0]{o}
.[1] -> Cell[0]{o}
```

Alternatively, we can use the method `.pseudo_tensor`, which does not change the value of the objects `.compose_state`. Below, we give the output of the previous code if we used the method `.pseudo_tensor` instead of `.merging_tensor`.

```
>>> sc.merge_base([0,1,3],operad.merging_tensor,"sorted")
>>> sc.stdout()
[0] -> Cell[0]{ooo}
.[1] -> Cell[0]{ooo}
..[2] -> Cell[0]{o}
..[2] -> Cell[0]{o}
..[2] -> Cell[0]{o}
.[1] -> Cell[0]{o}
.[1] -> Cell[0]{o}
```

**4.2.16. Description of `.divide_base` (method).** This section describes the code and the functionalities of the method `.divide_base`. The method is equipped with the following input variables, including one that is optional.

| .divide_base | | |
|---|---|---|
| Inputs | Types | Specifications |
| list_of_organelles | list(list(float)) | necessary |
| tensor | fun: list(SuperCell) -> SuperCell | necessary |
| order = "non-sorted" | string | optional |

The method possesses one action, which we describe below through sevaral examples.

<table>
<tr><th colspan="2">Action</th></tr>
<tr><td>Condition</td><td>Always</td></tr>
<tr><td>Description</td><td>The method `.divide_base` implements the operation described in [1, Def. 2.25]. The idea is to extend the method `.divide` (section 4.1.18) associated with the class `Cell` to the basal junction of `self` (see section 4.2.9). Put simply, the method separates a subset of the organelles of the root cell of `self` from its complement subset in two different cells and uses the input function `tensor` to distribute the child super cells, stored in `self.innercells`, accordingly (see picture below).<br>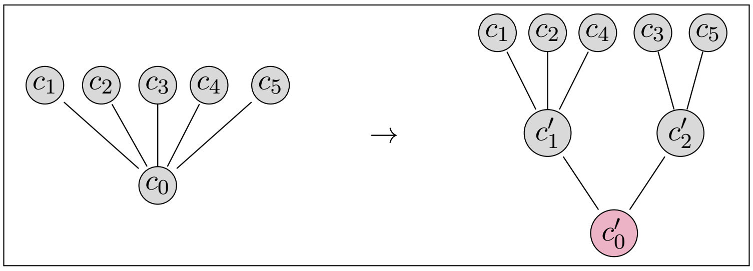<br><br>Now, to be more specific, the subset of indices of the organelles of `self` to be isolated from the other organelles is given by the list `list_of_organelles`. The method isolates the corresponding organelles in different cells `c_1` and `c_2` by calling the function<br>`self.cell.divide(list_of_organelles).`<br>The method then makes sure that the merging of these organelles is consistent with the whole tree structure of `self`. This is done by creating two new lists `new_innercells_1` and `new_innercells_2` of child super cells for the newly generated cells `c_1` and `c_2`. The list `new_innercells_1` contains the super cells of the form `self.innercells[i]` whose index `i`, ranging from `0` to `len(self.innercells)-1`, is in `list_of_organelles`, while the list `new_innercells_2` contains the super cells of the form `self.innercells[i]` whose index `i` is not in `list_of_organelles`. The resulting super cells are defined in the obvious way as follows:<br>`sc_1 = SuperCell(c_1,new_innercells_1)`<br>`sc_2 = SuperCell(c_2,new_innercells_2).`<br>The method fills up the lists `sc1.pre_action` and `sc2.pre_action` in the same way as the lists `new_innercells_1` and `new_innercells_1` are filled, but where we replace the lists `self.innercells_1` and `self.innercells_2` with the lists `self.pre_action`.<br>Finally, the method `divide_base` computes `self.pre_action` as the list of length 2 containing the outputs of the function `sc_1.tensor_pre_action()` and `sc_2.tensor_pre_action()` (see section 4.2.14);</td></tr>
</table>

The following example illustrates the use of the method `.divide_base` with the methods `.pseudo_tensor` (section 4.3.7) and `.dividing_tensor` (section 4.3.9) associated with the class `Operad`. In addition, we use the method `.spontaneous_reaction` (section 4.2.13) to store in each object `.cytosol` a list of zero values (as required by the method `.divide` (section 4.1.18) and pre-fill each object `pre_action` with a non-empty list of length 2, for every super cell present in the hierarchical structure of the super cell `sc`.

```
>>> c = Cell(2,0.5,[4,2],[[1,2],[5,1],[1,8],[4,4],[0,7]])
>>> operad = Operad(2)
>>> sc = SuperCell(c)
>>> sc.spontaneous_reaction([0,0],operad.identity)
>>> sc.stdout()
[0] -> Cell[6]{ooooo}
```

Below, we use the method `.dividing_tensor` for the input `tensor` associated with the method `.divid_top`. This has the consequence of setting the objects `.compose_state` of each divided super cell to `True`, as indicated by the purple display returned by the method `.stdout`.

```
>>> sc.divide_base([0,1,3],operad.dividing_tensor)
>>> sc.stdout()
[0] -> Cell[0]{oo}
.[1] -> Cell[3]{ooo}
.[1] -> Cell[3]{oo}
```

Alternatively, we can use the method `.pseudo_tensor`, which does not change the value of the objects `.compose_state`. Below, we give the output of the previous code if we used the method `.pseudo_tensor` instead of `.merging_tensor`.

```
>>> sc.divide_base([0,1,3],operad.pseudo_tensor)
>>> sc.stdout()
[0] -> Cell[0]{oo}
.[1] -> Cell[3]{ooo}
.[1] -> Cell[3]{oo}
```

**4.2.17. Description of `.fusion` (method).** This section describes the code and the functionalities of the method `.fusion`. The method is equipped with the following input variables, including one that is optional.

| `.fusion` | | |
|---|---|---|
| Inputs | Types | Specifications |
| `vector` | `list(float)` | necessary |
| `operad` | `Operad` | necessary |
| `filtering = [1.5,0]` | `list(float)` | optional |

The method possesses one action, which we describe below through an example.

| Action | |
|---|---|
| Condition | Always |
| Description | The method implements the algorithmic step described in [1, Sec. 2.5]. It mainly consists in recursively applying the method `.merge_base` (section 4.2.15) on all the non-leaf cells of `self` (see [1, Conv. 2.2 & Def. 2.3]) – leaf cells are excluded to keep the number of organelles of the super cell constant (in the sense of [1, Def. 2.7]). To do so, the method first calls the function `self.spontaneous_reaction(vector,operad.identity)` (see section 4.2.13) to clean every super cell present in the hierarchical structure of `self` from its cytosolic content. Then, the method computes a list `mer` of candidate organelles to be merged within each super cell by calling the function `*.cell.proposed_clustering(*.pre_action,"merging",filtering)` (see section 4.1.22), where `*` denotes any super cell present in the hierarchical structure of `self`. If the list `mer` is empty or includes all the organelles, then the fusion event is canceled. Otherwise, the method calls the function `self.merge_base(mer,operad.merging_tensor)`, which merges the organelles of whose indices belong to the list `mer`. Note that the merging is done relative to the method `.merging_tensor`, which is associated with the input `operad` (see section 4.2.15 and section 4.3.8).<br>Finally, the method ues the function `*.reset_depth()` to reset the depth of every super cell in the hierarchical structure of `self`. |

The following example illustrates the use of the method `.fusion` with the class `Operad`.

```
>>> c = Cell(2,0.5,[4,2],[[1,2],[5,1],[1,8],[4,4],[0,7]])
>>> operad = Operad(2)
>>> c1 = SuperCell(Cell(2,0.5,[1,1],[[10,1],[40,2],[10,5]]))
>>> c2 = SuperCell(Cell(2,0.5,[1,1],[[10,1],[40,2],[35,2]]))
>>> c3 = SuperCell(Cell(2,0.5,[1,1],[[10,1],[40,2],[100,5]]))
>>> c4 = SuperCell(Cell(2,0.5,[1,1],[[1,15],[0,25],[40,12]]))
>>> c5 = SuperCell(Cell(2,0.5,[1,1],[[80,1],[0,2],[27,1]]))
>>> sc = SuperCell(c,[c1,c2,c3,c4,c5])
>>> sc.stdout([10,50])
[0] -> Cell[6]{oooooo}
.[1] -> Cell[2]{ooo} [0]:0.292714272 [1]:0.2448393108 [2]:0.6139406135
.[1] -> Cell[2]{ooo} [0]:0.292714272 [1]:0.2448393108 [2]:0.2517386496
.[1] -> Cell[2]{ooo} [0]:0.292714272 [1]:0.2448393108 [2]:0.2448393108
.[1] -> Cell[2]{ooo} [0]:0.9914542955 [1]:0.9805806756 [2]:0.4696129729
.[1] -> Cell[2]{ooo} [0]:0.2083571163 [1]:0.9805806756 [2]:0.232274682
>>> sc.fusion([10,50],operad)
```

```
Searching cliques in the following graph:
[0, 2, 2, 1, 2]
[0, 0, 2, 1, 2]
[0, 0, 0, 1, 2]
[0, 0, 0, 0, 1]
[0, 0, 0, 0, 0]
Maximal weight = 2
Edges with maximal weight = [[0, 1], [0, 2], [0, 4], [1, 2], [1, 4], [2,
4]]
Merging occurring due to organelle(s):  [0, 1, 2, 4] at level = 0
>>> sc.stdout([10,50])
[0] -> Cell[6]{oo}
.[1] -> Cell[0]{oooo}
..[2] -> Cell[2]{ooo} [0]:0.292714272 [1]:0.2448393108 [2]:0.6139406135
..[2] -> Cell[2]{ooo} [0]:0.292714272 [1]:0.2448393108 [2]:0.2517386496
..[2] -> Cell[2]{ooo} [0]:0.292714272 [1]:0.2448393108 [2]:0.2448393108
..[2] -> Cell[2]{ooo} [0]:0.2083571163 [1]:0.9805806756 [2]:0.232274682
.[1] -> Cell[2]{ooo} [0]:0.9914542955 [1]:0.9805806756 [2]:0.4696129729
>>> sc.innercells[0].innercells[0].cell.stdout()
residual:  2.5
cytosol[1]:  0
cytosol[2]:  0
1-th organelle:  [10, 1]
2-th organelle:  [40, 2]
3-th organelle:  [10, 5]
>>> sc.innercells[0].innercells[1].cell.stdout()
residual:  2.5
cytosol[1]:  0
cytosol[2]:  0
1-th organelle:  [10, 1]
2-th organelle:  [40, 2]
3-th organelle:  [35, 2]
>>> sc.innercells[0].innercells[2].cell.stdout()
residual:  2.5
cytosol[1]:  0
cytosol[2]:  0
1-th organelle:  [10, 1]
2-th organelle:  [40, 2]
3-th organelle:  [100, 5]
>>> sc.innercells[0].innercells[3].cell.stdout()
residual:  2.5
cytosol[1]:  0
cytosol[2]:  0
1-th organelle:  [80, 1]
2-th organelle:  [0, 2]
3-th organelle:  [27, 1]
```

**4.2.18. Description of `.fission` (method).** This section describes the code and the functionalities of the method `.fission`. The method is equipped with the following input variables, including one that is optional.

| .fission | | |
|---|---|---|
| Inputs | Types | Specifications |
| `vector` | `list(float)` | necessary |
| `operad` | `Operad` | necessary |
| `filtering = [1.5,0]` | `list(float)` | optional |

The method possesses one action, which we describe below through an example.

| Action | |
|---|---|
| Condition | Always |
| Description | The method implements the algorithmic step described in [1, Sec. 2.5]. It mainly consists in recursively applying the method `.divide_base` (section 4.2.16) on all the cells of `self` – including leaf cells [1, Conv. 2.2 & Def. 2.3]. To do so, the method first calls the function<br>`self.spontaneous_reaction(vector,operad.identity)`<br>(see section 4.2.13) to clean every super cell present in the hierarchical structure of `self` from its cytosolic content. Then, the method computes a list `div` of candidate organelles to be isolated from the other organelles within each super cell by calling the function<br>`*.cell.proposed_clustering(*.pre_action,"division",filtering)`<br>(see section 4.1.22), where `*` denotes any super cell in the hierarchical structure of `self`. If the list `div` is empty or includes the indices of all the organelles, then the fission event is canceled. Otherwise, the method calls the function `self.divide_base(div,operad.dividing_tensor)`, which isolates the organelles whose indices belong to the list `div`. Note that the division is done relative to the method `.dividing_tensor`, which is associated with the input `operad` (see section 4.2.16 and section 4.3.9).<br>Finally, the method uses the function `*.reset_depth()`s to reset the depth of every super cell in the hierarchical structure of `self`. |

The following example illustrates the use of the method `.fission` with the class `Operad`.

```
>>> c = Cell(2,0.5,[4,2],[[1,2],[5,1],[1,8],[4,4],[0,7]])
>>> operad = Operad(2)
>>> c1 = SuperCell(Cell(2,0.5,[1,1],[[10,1],[40,2],[10,5]]))
>>> c2 = SuperCell(Cell(2,0.5,[1,1],[[10,1],[40,2],[35,2]]))
>>> c3 = SuperCell(Cell(2,0.5,[1,1],[[10,1],[40,2],[100,5]]))
>>> c4 = SuperCell(Cell(2,0.5,[1,1],[[1,15],[0,25],[40,12]]))
>>> c5 = SuperCell(Cell(2,0.5,[1,1],[[80,1],[0,2],[27,1]]))
>>> sc = SuperCell(c,[c1,c2,c3,c4,c5])
>>> sc.stdout([10,50])
[0] -> Cell[6]{ooo}
.[1] -> Cell[2]{ooo} [0]:0.292714272 [1]:0.2448393108 [2]:0.6139406135
.[1] -> Cell[2]{ooo} [0]:0.292714272 [1]:0.2448393108 [2]:0.2517386496
.[1] -> Cell[2]{ooo} [0]:0.292714272 [1]:0.2448393108 [2]:0.2448393108
.[1] -> Cell[2]{ooo} [0]:0.9914542955 [1]:0.9805806756 [2]:0.4696129729
.[1] -> Cell[2]{ooo} [0]:0.2083571163 [1]:0.9805806756 [2]:0.232274682
>>> sc.fission([10,50],operad)
```

```
Warning:  in SuperCell._fission:  total fission canceled at level 1
Warning:  in SuperCell._fission:  total fission canceled at level 1
Warning:  in SuperCell._fission:  total fission canceled at level 1
Searching cliques in the following graph:
[0, 1, 0]
[0, 0, 0]
[0, 0, 0]
Maximal weight = 1
Edges with maximal weight = [[0, 1]]
Division occurring due to organelle(s):  [0, 1] at level = 1
Warning:  in SuperCell._fission:  total fission canceled at level 1
Searching cliques in the following graph:
[0, 0, 0, 0, 0]
[0, 0, 0, 0, 1]
[0, 0, 0, 1, 0]
[0, 0, 0, 0, 0]
[0, 0, 0, 0, 0]
Maximal weight = 1
Edges with maximal weight = [[1, 4], [2, 3]]
Division occurring due to organelle(s):  [2, 3] at level = 0
>>> sc.stdout([10,50])
[0] -> Cell[0]{oo}
.[1] -> Cell[3]{oo}
..[2] -> Cell[2]{ooo} [0]:0.9904048953 [1]:0.9835249384 [2]:0.7321129014
..[2] -> Cell[0]{oo}
...[3] -> Cell[1]{oo} [0]:0.9914542955 [1]:0.9805806756
...[3] -> Cell[1]{o} [0]:0.4696129729
.[1] -> Cell[3]{ooo}
..[2] -> Cell[2]{ooo} [0]:0.292714272 [1]:0.2448393108 [2]:0.6139406135
..[2] -> Cell[2]{ooo} [0]:0.292714272 [1]:0.2448393108 [2]:0.2517386496
..[2] -> Cell[2]{ooo} [0]:0.2083571163 [1]:0.9805806756 [2]:0.232274682
>>> sc.innercells[0].innercells[0].cell.stdout()
residual:  1.25
cytosol[1]:  0
cytosol[2]:  0
1-th organelle:  [1, 17]
2-th organelle:  [1, 64]
3-th organelle:  [25, 18]
>>> sc.innercells[0].innercells[1].innercells[0].cell.stdout()
residual:  1.25
cytosol[1]:  0
cytosol[2]:  0
1-th organelle:  [1, 15]
2-th organelle:  [0, 25]
>>> sc.innercells[0].innercells[1].innercells[1].cell.stdout()
residual:  1.25
cytosol[1]:  0
cytosol[2]:  0
1-th organelle:  [40, 12]
```

**4.2.19. Description of .compose (method).** This section describes the code and the functionalities of the method .compose. The method is equipped with the following input variable.

| .compose | | |
|---|---|---|
| Inputs | Types | Specifications |
| identity | fun: list(float) -> Cell | necessary |

The method possesses one action, which we describe below through an example.

| Action | |
|---|---|
| Condition | Always |
| Description | The method implements the algorithmic step described in [1, Sec. 2.4] – the idea is to compose, within their respective parent cells, every super cell whose object .compose_state contains True. An important feature of the method is to proceed in a way that does not break the super cell structure (which is possible – see section [1, Sec. 2.4]).<br>Before describing the method is details, we can broadly describe the method as follows: it uses the function *.cell.compose(k,c) for every parent cell * and child cell c whose associated object .compose_state contains True (see section 4.1.8). During this composition process, the method also takes care of preserving the association between the pre-actions and their corresponding super cells.<br>More specifically, the method proceed as follows:<br>(1) if self.is_leaf contains the value True, then the method returns self.<br>(2) if self.is_leaf contains the value False, then the method checks whether all super cells in the object self.innercells are such that their associated objects .is_leaf and .compose_state contain the value True.<br>(2.1) If this is the case, then self.innercells only contains leaves (section 4.2.3) and the method composes all these leaf SuperCell within self, making the resulting SuperCell item self a leaf.<br>(2.2) Otherwise, the method composes the cells associated with the super cells of the list self.innercells within self by making sure to equip each leaf SuperCell of self.innercells with formal child super cells consisting of the super cells<br>SuperCell(identity(comp_i.cell.organelles[j]))<br>where comp_i is the output of the recursive call of the method .compose(identity) for the leaf super cell self.innercells[i]. |

| Description | Note that because the root cell of the underlying tree structure of `self` does not have a parent cell, the composition of the root cell never occurs, which usually has the consequence of making the hierarchical structure of `self` grow over time. The growth of the tree caused by these exceptions is illustrated below in the context of a series of calls of the method `.fission` (section 4.2.18) directly followed by calls of the method `.compose`. We display in brackets the outputs of the method `.fission` if they are different from the outputs of the method `.compose`. 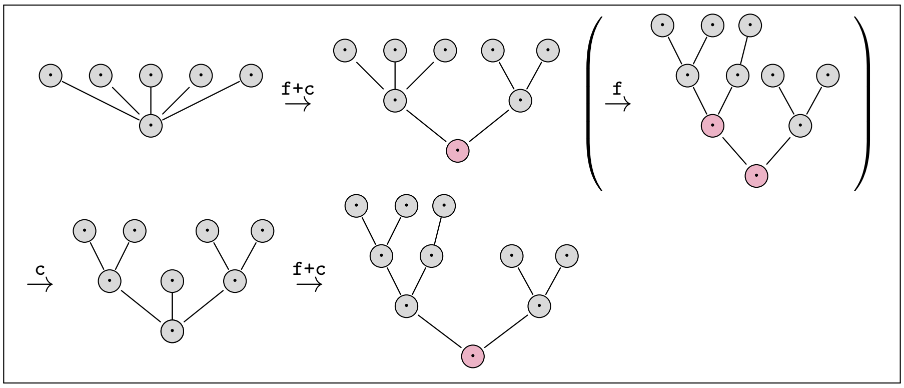 |
|---|---|

The following example combines the examples used in section 4.2.17 and section 4.2.18. We first construct a super cell of level 2 that possesses internal super cells whose objects `.compose_state` are `True`.

```
>>> c = Cell(2,0.5,[4,2],[[1,2],[5,1],[1,8],[4,4],[0,7]])
>>> operad = Operad(2)
>>> c1 = SuperCell(Cell(2,0.5,[1,1],[[10,1],[40,2],[10,5]]))
>>> c2 = SuperCell(Cell(2,0.5,[1,1],[[10,1],[40,2],[35,2]]))
>>> c3 = SuperCell(Cell(2,0.5,[1,1],[[10,1],[40,2],[100,5]]))
>>> c4 = SuperCell(Cell(2,0.5,[1,1],[[1,15],[0,25],[40,12]]))
>>> c5 = SuperCell(Cell(2,0.5,[1,1],[[80,1],[0,2],[27,1]]))
>>> sc = SuperCell(c,[c1,c2,c3,c4,c5])
>>> sc.stdout([10,50])
[0] -> Cell[6]{ooo}
.[1] -> Cell[2]{ooo} [0]:0.292714272 [1]:0.2448393108 [2]:0.6139406135
.[1] -> Cell[2]{ooo} [0]:0.292714272 [1]:0.2448393108 [2]:0.2517386496
.[1] -> Cell[2]{ooo} [0]:0.292714272 [1]:0.2448393108 [2]:0.2448393108
.[1] -> Cell[2]{ooo} [0]:0.9914542955 [1]:0.9805806756 [2]:0.4696129729
.[1] -> Cell[2]{ooo} [0]:0.2083571163 [1]:0.9805806756 [2]:0.232274682
>>> sc.fusion([10,50],operad)
Searching cliques in the following graph:
[0, 2, 2, 1, 2]
[0, 0, 2, 1, 2]
[0, 0, 0, 1, 2]
[0, 0, 0, 0, 1]
[0, 0, 0, 0, 0]
Maximal weight = 2
Edges with maximal weight = [[0, 1], [0, 2], [0, 4], [1, 2], [1, 4], [2,
4]]
Merging occurring due to organelle(s):  [0, 1, 2, 4] at level = 0
>>> sc.stdout([10,50])
```

```
[0] -> Cell[6]{oo}
.[1] -> Cell[0]{oooo}
..[2] -> Cell[2]{ooo} [0]:0.292714272 [1]:0.2448393108 [2]:0.6139406135
..[2] -> Cell[2]{ooo} [0]:0.292714272 [1]:0.2448393108 [2]:0.2517386496
..[2] -> Cell[2]{ooo} [0]:0.292714272 [1]:0.2448393108 [2]:0.2448393108
..[2] -> Cell[2]{ooo} [0]:0.2083571163 [1]:0.9805806756 [2]:0.232274682
.[1] -> Cell[2]{ooo} [0]:0.9914542955 [1]:0.9805806756 [2]:0.4696129729
>>> sc.fission([10,50],operad)
Warning:  in SuperCell._fission:  total fission canceled at level 2
Warning:  in SuperCell._fission:  total fission canceled at level 2
Warning:  in SuperCell._fission:  total fission canceled at level 2
Warning:  in SuperCell._fission:  total fission canceled at level 2
[0, 0, 0, 1]
[0, 0, 0, 0]
[0, 0, 0, 0]
[0, 0, 0, 0]
Maximal weight = 1
Edges with maximal weight = [[0, 3]]
Division occurring due to organelle(s):  [0, 3] at level = 1
Searching cliques in the following graph:
[0, 1, 0]
[0, 0, 0]
[0, 0, 0]
Maximal weight = 1
Edges with maximal weight = [[0, 1]]
Division occurring due to organelle(s):  [0, 1] at level = 1
Warning:  in SuperCell._fission:  total fission canceled at level 0
>>> sc.stdout([10,50])
[0] -> Cell[6]{oo}
.[1] -> Cell[0]{oo}
..[2] -> Cell[0]{oo}
...[3] -> Cell[2]{ooo} [0]:0.292714272 [1]:0.2448393108 [2]:0.6139406135
...[3] -> Cell[2]{ooo} [0]:0.2083571163 [1]:0.9805806756 [2]:0.232274682
..[2] -> Cell[0]{oo}
...[3] -> Cell[2]{ooo} [0]:0.292714272 [1]:0.2448393108 [2]:0.2517386496
...[3] -> Cell[2]{ooo} [0]:0.292714272 [1]:0.2448393108 [2]:0.2448393108
.[1] -> Cell[0]{oo}
..[2] -> Cell[1]{oo} [0]:0.9914542955 [1]:0.9805806756
..[2] -> Cell[1]{o} [0]:0.4696129729
```

We now show the action of the method `.compose()` on the super cell `cs` – note that the output is a 1-level tree structure, which corresponds to the same type of structure we started with (see the first display `cs.stdout([10,50])`).

```
>>> sc.compose(operad.identity)
>>> sc.stdout([10,50])
[0] -> Cell[6]{oooo}
.[1] -> Cell[5]{oooooo} [0]:0.292714272 [1]:0.2448393108 [2]:0.6139406135
[3]:0.2083571163 [4]:0.9805806756 [5]:0.232274682
.[1] -> Cell[5]{oooooo} [0]:0.292714272 [1]:0.2448393108 [2]:0.2517386496
[3]:0.292714272 [4]:0.2448393108 [5]:0.2448393108
.[1] -> Cell[1]{oo} [0]:0.9914542955 [1]:0.9805806756
.[1] -> Cell[1]{o} [0]:0.4696129729
```

#### 4.3. Description of Operad (class)

**4.3.1. Introduction.** This section introduces the reader to the code of the class Operad, which possesses the following external dependencies.

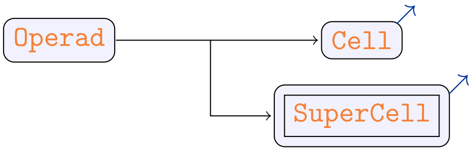

The main goal of the class Operad is to model the features of the colored operad described in [1, Sec. 3]. Before presenting the features of the class, recall that, mathematically, this colored operad can be defined as the map $\mathcal{O}_n^N : (\mathbb{R}_+^N)^n \times \mathbb{R}_+^N \to \mathbf{Set}$ sending every tuple $(x_1,\dots,x_n,y)$ in $(\mathbb{R}_+^N)^n \times \mathbb{R}_+^N$ to the set $\mathbb{R}$. Every element 0 in a set of the form $\mathcal{O}_1^N(a,a) = \mathbb{R}$ is called the *identity* on $a$ and the sum operation of the form

$$\mathcal{O}_n^N(x_1,\dots,x_n,y) \times \prod_{i=1}^{n} \mathcal{O}_{m_i}^N(z_{i,1},\dots,z_{i,m_i},x_i) \to \mathcal{O}_{m_1+\dots+m_n}^N(z_{1,1},\dots,z_{n,m_n},y)$$

is called a *composition*. In addition, the operad is equipped with tensor operations given by the addition operations of the following form:

$$\otimes : \mathcal{O}_n^N(x_1,\dots,x_n,y) \times \mathcal{O}_m^N(x_1',\dots,x_m',y') \to \mathcal{O}_{n+m}^N(x_1,\dots,x_n,x_1',\dots,x_m',y+y').$$

In this library, the class Operad is equipped with 1 object, recording the dimension $N$ of the objects of the operad, and 2 types of methods, which are listed below:

▷ a set of 3 methods meant to generate cells and super cells:

- `__init__`: initializes the dimension of the objects of the operad;
- `identity`: creates an identity cell, as defined in [1, Def. 1.9]);
- `generate`: generate a super cell with random values (section 4.2).

▷ a set of 4 methods giving a variety of tensor structures [1, Sec. 1.7]:

- `tensor`: returns a tensor of cells, as defined in [1, Conv. 1.50 & Ex. 1.51] .
- `pseudo_tensor`: returns the non-composed version of a tensor (see [1, Ex. 1.52]).
- `merging_tensor`: updates the objects `.compose_state` associated with the child cells of the root of a super cell according to [1, Def. 2.26].
- `dividing_tensor`: updates the object `.compose_state` associated with the root of a super cell according to [1, Def. 2.25].

**4.3.2. Structure.** The following tables give a preview of the class Operad. The table given below describes the various dependencies of the class.

| Dependencies | |
|---|---|
| Superclass ancestry | Module section |
| object | N/A |
| Statistics | |
| ▷ Importable objects: 1 | |
| ▷ Non-importable objects: 0 | |
| ▷ Importable methods: 7 | |
| ▷ Non-importable methods: 0 | |

The following table gives a description of the single importable object of the class.

| Objects | | |
|---|---|---|
| Name | Type | Related sections |
| .dimension | int | ▷ section 4.3.3 |

Finally, the following table gives a description of the 7 importable methods of the class:

| Methods | | | |
|---|---|---|---|
| Name | Input types | Output types | Related sections |
| .__init__ | - int | - Cell | ▷ section 4.3.3 |
| .identity | - list(float) | - Cell | ▷ section 4.3.4 |
| .tensor | - list(Cell) | - Cell | ▷ section 4.3.6 |
| .pseudo_tensor | - list(SuperCell) | - SuperCell | ▷ section 4.3.7 |
| .merging_tensor | - list(SuperCell) | - SuperCell | ▷ section 4.3.8 |
| .dividing_tensor | - list(SuperCell) | - SuperCell | ▷ section 4.3.9 |
| .generate | - int<br>- int<br>- fun: void -> float | - SuperCell | ▷ section 4.3.3 |

**4.3.3. Description of .__init__ (method).** Writing in progress – see cl_ope.py for more information.

**4.3.4. Description of .identity (method).** Writing in progress – see cl_ope.py for more information.

**4.3.5. Description of .generate (method).** Writing in progress – see cl_ope.py for more information.

**4.3.6. Description of .tensor (method).** Writing in progress – see cl_ope.py for more information.

**4.3.7. Description of .pseudo_tensor (method).** Writing in progress – see cl_ope.py for more information.

**4.3.8. Description of .merging_tensor (method).** Writing in progress – see cl_ope.py for more information.

**4.3.9. Description of .dividing_tensor (method).** Writing in progress – see cl_ope.py for more information.

### Presentation of the module `cellint.py`

#### 5.1. Description of `intcyt` (function)

This section describes the code of the function `intcyt`. The function is equipped with the following input variable, where we denote by $\alpha$ a polymophic type and we use the symbol `X` as a shortcut for the type `list(list(float))`.

| `intcyt` | | |
|---|---|---|
| Inputs | Types | Specifications |
| `operad` | `list(Operad)` | necessary |
| `supercell` | `list(SuperCell)` | necessary |
| `index` | `list(int)` | necessary |
| `events` | `list`($\alpha$) | necessary |
| `vector` | `list(float)` | necessary |
| `gamma` | `fun: Cell * X -> X` | necessary |
| `filtering = [1.5,1.5,0]` | `list(float)` | optional |

The function possesses one action, which is discussed in the tutorial of section 2.

| Action 1 | |
|---|---|
| Condition | Always |
| Description | This function implements the algorithm of [1, Sec. 2] – see section 2 of this documentation for a detailed description. |
